# Selection on many loci drove the origin and spread of a key innovation

**DOI:** 10.1101/2023.02.13.528213

**Authors:** Sean Stankowski, Zuzanna B. Zagrodzka, Martin D. Garlovsky, Arka Pal, Daria Shipilina, Diego Garcia Castillo, Alan Le Moan, Erica Leder, James Reeve, Kerstin Johannesson, Anja M. Westram, Roger K. Butlin

## Abstract

Key innovations are fundamental to biological diversification, but their genetic architecture is poorly understood. A recent transition from egg-laying to live-bearing in *Littorina* snails provides the opportunity to study the architecture of an innovation that has evolved repeatedly in animals. Samples do not cluster by reproductive mode in a genome-wide phylogeny, but local genealogical analysis revealed numerous genomic regions where all live-bearers carry the same core haplotype. Associated regions show evidence for live-bearer-specific positive selection, and are enriched for genes that are differentially expressed between egg-laying and live-bearing reproductive systems. Ages of selective sweeps suggest live-bearing alleles accumulated gradually, involving selection at different times in the past. Our results suggest that innovation can have a polygenic basis, and that novel functions can evolve gradually, rather than in a single step.

## Main text

Evolution is a gradual process, but occasionally results in sudden changes in form and function that allow organisms to exploit new ecological opportunities (*1, 2*). These game-changing traits—including flight, vision, and the bearing of live offspring—are known as ‘key innovations’ (*2*–*5*). Key innovations are all around us, and have catalyzed the diversification of many groups (*1*). Despite their significance, we know surprisingly little about the origins and genetic architecture of most innovations (*1*). This is because most originated deep in the past, making it difficult to disentangle causal loci from the countless genetic changes that accumulated up to the present.

A recent transition in female reproductive mode offers a rare opportunity to study the genetic architecture of an innovation that has evolved many times across the animal kingdom (*6*). We focus on a clade of intertidal gastropods (Genus *Littorina*), where the ancestral state is to lay a large egg-mass but one species gives birth to live young (Fig. 1A, fig. S1) (*7, 8*). Egg-layers have a gland that embeds egg-capsules into a protective jelly. In the live-bearer, *L. saxatilis*, this structure has evolved into a brood pouch where embryos develop inside the mother. Live-bearing is a recent innovation in the littorinidae and is thought to allow snails to reproduce in areas where eggs are exposed to harsh conditions (*8*). This is reflected in the much broader ecological and geographic distribution of *L. saxatilis* compared with the two egg-laying sister species, *L. arcana and L. compressa* (*8*) (Fig. 1B and 1C). Egg-laying and live-bearing species have adapted in parallel to contrasting environments (*8, 9*), partly decoupling reproductive mode from other axes of phenotypic divergence (Fig. 1B). There is also evidence for occasional hybridization between egg-layers and live-bearers (*10, 11*). These features provide an opportunity to identify and study the genetic changes underlying the live-bearing innovation.

**Figure 1.**
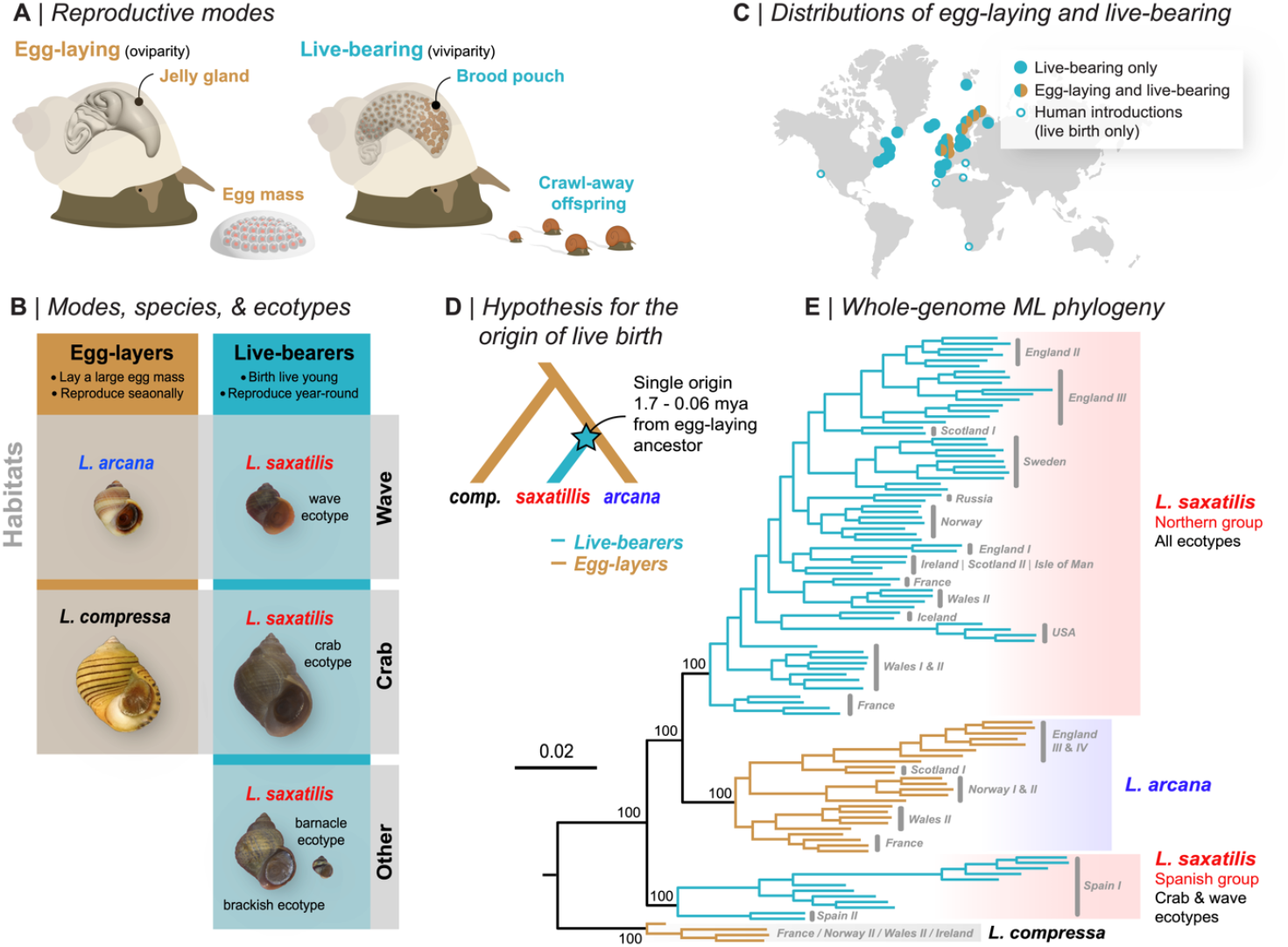
Variation in reproductive mode in *Littorina*. (A) Anatomical differences between modes (B) Egg-layers reproduce during a limited breeding season, while live-bearers release offspring year-round. The two egg-layers share their habitats with ecotypes of the live-bearer, *L. saxatilis*. (C) Approximate distributions of the modes, highlighting the broader distribution of live birth. (D) Existing hypothesis for the origin of live birth. (E) ML phylogenetic tree based on whole-genome sequences (108 individuals and 18.5 million variable sites). Bootstrap support for key nodes is shown.

## Live-bearing snails do not form a monophyletic group

We used whole-genome sequences from 108 individuals to test the existing hypothesis of a single origin of live-bearing (Fig. 1D) (*7*). Surprisingly, live-bearers formed two separate clades in a phylogenetic tree: one containing all *L. saxatilis* from Spain (hereafter ‘Spanish *saxatilis*’), and another including all other *L. saxatilis* (‘Northern *saxatilis*’) that was sister to egg-laying *L. arcana* (Fig. 1E). The discordance between evolutionary relationships and reproductive mode (also seen in PCAs, fig. S6) has several possible explanations, including two genetically independent transitions between egg-laying and-live birth. However, given the close relationships of these taxa, a single origin could have been followed by the sharing of causal alleles between lineages via gene exchange and selection (*12*). For example, live-bearing could first have evolved in Spain, whereupon causal alleles spread to the north, introgressing into the genetic background of the resident egg-laying lineage. In this case, we would expect genealogies for loci causing live birth to be strongly discordant from the genome-wide tree, with samples grouping by reproductive mode (*9*).

## Topology weighting reveals rampant genealogical discordance and loci associated with reproductive mode

With this expectation in mind, we used topology weighting (Fig. 2A) to identify genomic regions associated with reproductive mode. For each genomic window, topology weighting calculates the degree of monophylly toward three possible taxon subtrees (Fig. 2C, fig. S8): (*i*) the background topology, Tb, observed in our genome-wide analysis, (ii) the reproduction topology, Tr, where samples cluster by reproductive mode, and (iii) the control topology, Tc, which is of no specific interest except that it provides a control for distinguishing incomplete lineage sorting (ILS) from other processes that cause genealogical discordance. We used non-overlapping 100-SNP windows (mean size 5.8 kb, fig. S7), and calculated topology weights (*13*) for each window by sampling 10,000 subtrees (Fig. 2A). We took the novel approach of analyzing the joint distribution of topology weights in a ternary plot (Fig. 2A) and used simulations to understand how different factors shape the ternary distribution of weights (Fig. 2B; Supplementary text, figs. S9—S19; tables S3 & S4).

**Figure 2.**
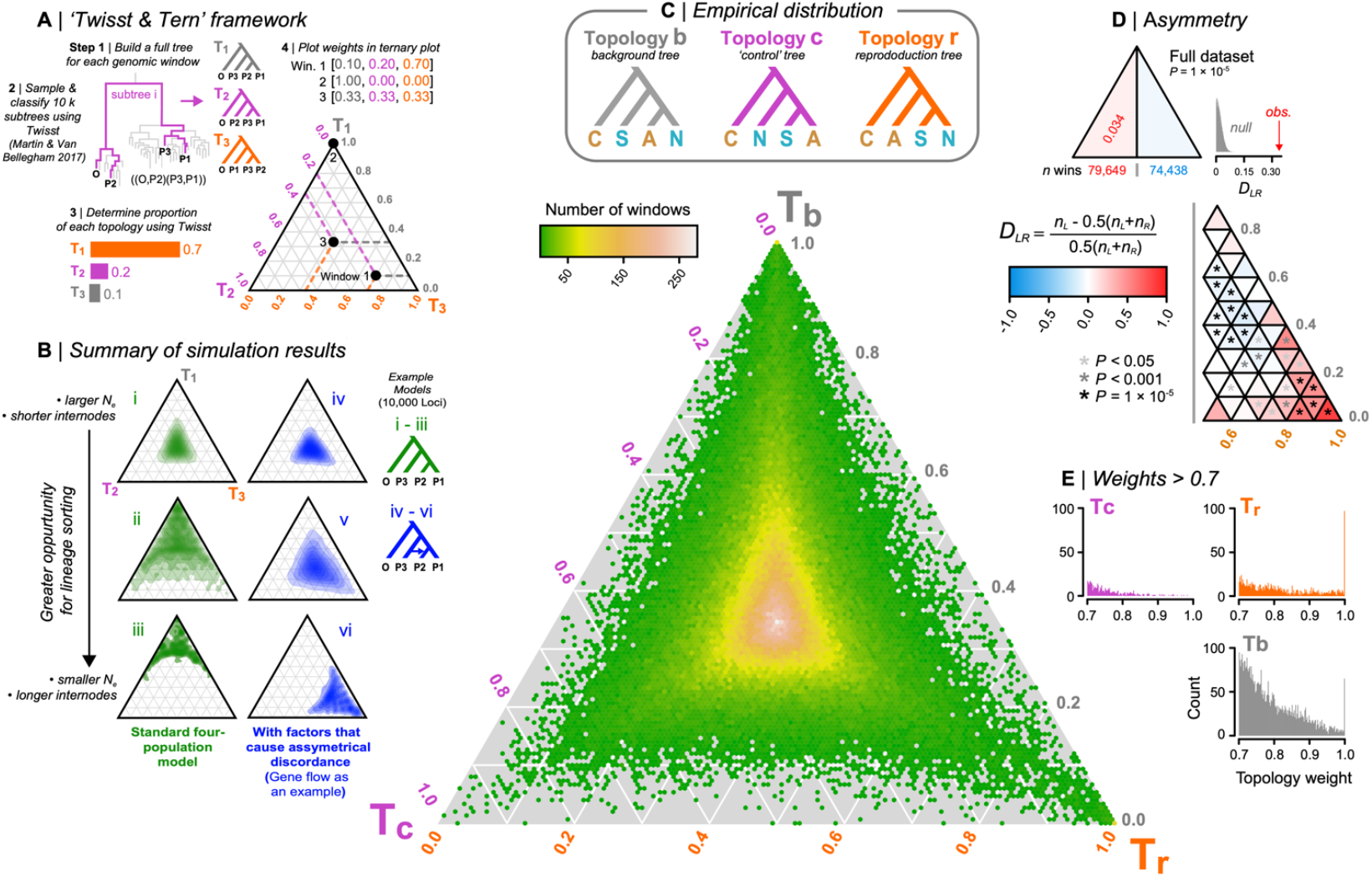
Topology weighting reveals genomic regions associated with reproductive mode. (A) For each genomic window, we inferred a full tree including all haplotypes, and then sampled and classified 10k ‘subtrees’ by randomly picking one haplotype per group. Topology weights are the proportions of each topology among all subtrees. Windows were then plotted in a ternary plot based on their topology weights. (B) Simulated distributions of weights. A greater opportunity for lineage sorting (i - iii) biases the distribution toward the topology that matches the demographic history. Incomplete lineage sorting yields genealogies that are a better fit to one of the discordant trees, but the distribution is always symmetrical between the left and right half triangles. Additional factors, including gene flow, create a bias toward one of the discordant genealogies (panels iv - vi). (C) Possible topologies and the empirical distribution of weights for the 154,971, 100 SNP windows; C: *compressa*, A: *arcana*, S: Spanish *saxatilis*, N: Northern *saxatilis*. Hexagonal bins are colored according to window count. (D) Counts of windows in the left and right half triangles, with the asymmetry quantified using *D*_*LR*_. Further division into sub-triangles reveals left-right asymmetry throughout the distribution. Asterisks indicate significant asymmetry between corresponding left- and right-sided sub-triangles. (E) Distributions of weights > 0.7.

We expected the empirical distribution of weights to be biased toward Tb, because lineage sorting results in concordance between the demographic history and underlying gene trees (*14*) (Fig. 2B, Supplementary text). However, the observed bias was only slight (Tb = 0.380, Tc = 0.310, Tr = 0.308), with just 62 of ∼155,000 genomic regions perfectly fitting Tb (*i*.*e*., Tb = 1) (Fig. 2C). Instead, the bulk of the distribution fell close to the center of the triangle, revealing extensive ILS due to rapid diversification relative to the effective population size (*14, 15*). Thus, although well-supported statistically, the genome-wide tree is a very poor predictor of evolutionary relationships at any given genomic region.

We found substantial left-right asymmetry in the distribution of weights (Fig. 2D). Such a bias is not expected to arise from ILS, because there is an equal chance that a given gene tree will more closely resemble either alternative topology (Fig. 2B, supplementary text) (*14*). We detected asymmetry using a new statistic, *D*_*LR*_ (Fig. 2D, fig. S19). A genome-wide test, performed by calculating *D*_*LR*_ between the two halves of the triangle, revealed a 3.4% excess of windows shifted toward the control topology (*D*_*LR*_ = 0.034, permutation test p = 1e-5). *D*_*LR*_ calculated between analogous left- and right-side sub-triangles, revealed that this asymmetry was driven by an excess of trees with a small bias toward Tc (Fig. 2D, table S5). Further exploration showed that this bias is due to several previously identified chromosomal inversions, where one arrangement is more common in Spanish *L. saxatilis* and *L. arcana*, and the other is more common in *L. compressa* and Northern *L. saxatilis* (*D*_*LR*_ for regions outside inversions = −0.007, p = 0.074) (figs. S21—S24, table S6, Supplementary text).

Much stronger asymmetry was observed between the extreme left and right sub-triangles, corresponding to windows that strongly fit one of the alternative topologies (Fig. 2D). However, the asymmetry was in the opposite direction to the genome-wide pattern, with a large excess of windows strongly biased toward the reproduction tree compared with the control tree (Tr > 0.7 = 1151 windows vs. 461 for Tc; *D*_*LR*_ = −0.43, p = 1e-5). A total of 88 windows perfectly fit the reproduction topology (*i*.*e*., Tr = 1), compared with 0 windows that perfectly fit the control topology (*D*_*LR*_ = 1.00, p = 1e-5; Fig. 2E, table S5).

## Evidence for live-bearer specific positive selection

Although neutral gene flow can generate strong asymmetry under some circumstances, we are unable to explain the observed Tr bias without invoking natural selection (supplementary text). We found strong additional evidence for live-bearer-specific positive selection in these regions. First, window-based estimates of nucleotide diversity (π) in live-bearers decreased substantially with increasing Tr weight (Fig. 3A). We found no such relationship in egg-layers. Eighty-four of the 88 (95%) perfectly associated regions showed reduced π in live-bearers (mean π_live-bearer_ = 0.0029 vs π_Egg-layer_ = 0.0065; paired Wilcoxon test, p = 1.313e-15, Fig. 3A, fig. S25), consistent with selection having purged diversity from live-bearing haplotypes (*16*). Although this pattern could in principle result from a live-bearer-specific demographic bottleneck, we can rule this out because live-bearers and egg-layers have similar levels of genome-wide diversity (mean π live-bearer = 0.0065 vs. π egg-layer = 0.0062; fig. S26). Further, relationships between π and the other weights (Ts and Tc) were weak, and similar for both groups, confirming that reduced π in live-bearers is specific to Tr rather than being a general feature of windows with extreme weights (fig. S27). The site-frequency spectra (SFS) and sample-size-corrected estimates of private alleles for perfectly associated regions provide further evidence for selection (Fig 3B & C; figs. S28—S30; tables S9 and S10): the live-bearer SFS was strongly skewed toward rare variants (Tajima’s D = −1.89, 95% CIs -1.77 – -2.01; fig. S29), the majority of which (80%) were private to the group. Both results are expected during the phase when diversity is recovered by mutation after a selective sweep (*17*). In contrast, the SFS for egg-layers was much closer to the neutral expectation (Tajima’s D = −0.24, 95% CIs −0.037 – −0.437), with polymorphic sites being 2.14 times more abundant in egg-layers after accounting for the difference in sample size.

**Figure 3.**
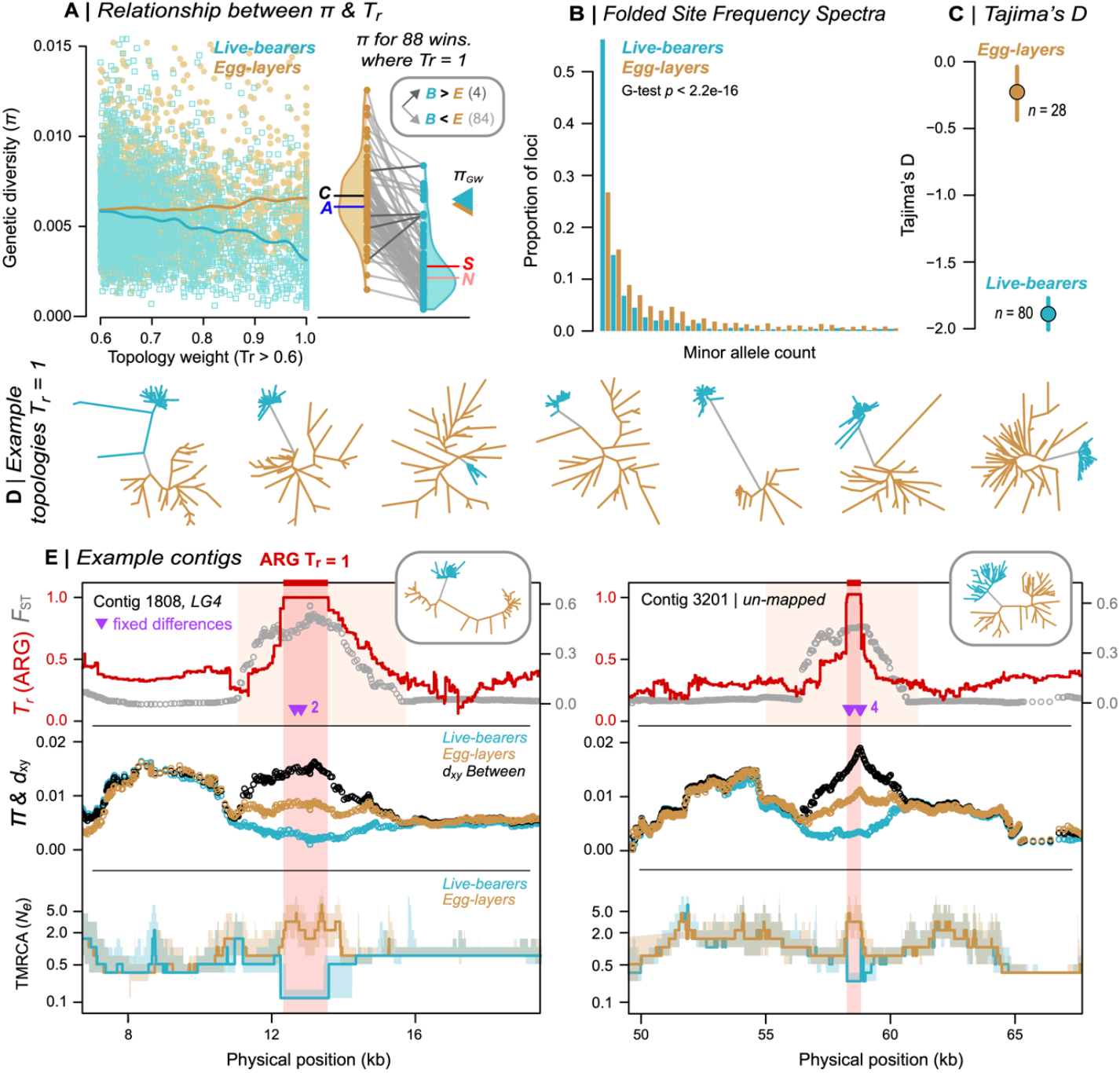
Evidence for positive selection on haplotypes associated with live birth. (A) Relationship between π and Tr for both reproductive modes. Triangles on right: genome-wide π. Violin plots: distributions of π for windows where Tr = 1. Letters show mean values of π for egg-layers and live-bearers. (B) Folded SFS for each mode in perfectly associated regions, projected at the same sample size for comparison. (C) Estimates of Tajima’s D with 95% CIs for perfectly associated regions. (D) Unrooted trees for example windows where Tr = 1. (E) Variation across two example contigs that contain a window where Tr = 1 (span of the orange box). The tree associated with each region is shown. Top panel: *F*_*ST*_ between egg-layers and live-bearers in 3kb sliding windows (30 bp step). TrARG shows the results of topology weighting applied to marginal trees obtained from inferred ancestral recombination graphs (ARGs). Purple arrows show fixed differences between modes. Middle panel: π and d_xy_ in sliding windows. Bottom panel: traces of time to the most recent common ancestor (TMRCA) obtained from ARGs. Bold lines: median estimates; Envelopes: 95% CIs. The red box shows the inferred length of the core haplotype block associated with live birth.

We characterized footprints of selection within contigs to more accurately estimate the number and size of candidate regions (Fig. 3E). The 88 perfectly associated windows mapped to 50 contigs in our genome assembly (mean 1.7 ± sd 1.5 windows per contig; table S8). Associated regions were narrow, mostly spanning less than 20 kb (mean 12 kb ± sd 14.4 kb). Sliding-window analysis of each contig generally revealed clear peaks of allele frequency differentiation (*F*_*ST*_) and sequence divergence (*d*_*xy*_) between the groups, as well as valleys of nucleotide diversity (π) in live-bearers (Fig. 3E; fig. S33). We also inferred ancestral recombination graphs (ARGs) for selected contigs to refine candidate regions (Fig. 3E). Unlike the trees for windows of arbitrary size and position, each marginal tree in an ARG corresponds to an inferred non-recombining segment of the genome (*18*). Thus, by applying topology weighting to the sequence of marginal trees, we were able to identify more precisely the segment of genome retained by all live bearing samples following the sweep. In both cases, the core live-bearing haplotype spanned less than 2 kb. Live-bearers showed much shallower coalescence in these regions than egg-layers, as expected following a selective sweep (Fig. 3E).

## Mode-associated regions are widespread and enriched for genes that are differentially expressed between reproductive systems

The assignment of contigs to a genetic map revealed that reproductive-mode-associated windows are widespread across the genome, rather than co-localizing to one or a few genomic regions (Fig. 4a; table S11). As expected for a polygenic trait, the number of mode-associated windows on each LG was strongly predicted by LG size (Tr > 0.7, r = 0.79, p < 0.0001; Tr > 0.9, r = 0.71, p < 0.005). Associated windows were also widespread within linkage groups, in some cases with strong associations near opposite ends of the same LG (Fig. 4B).

**Figure 4.**
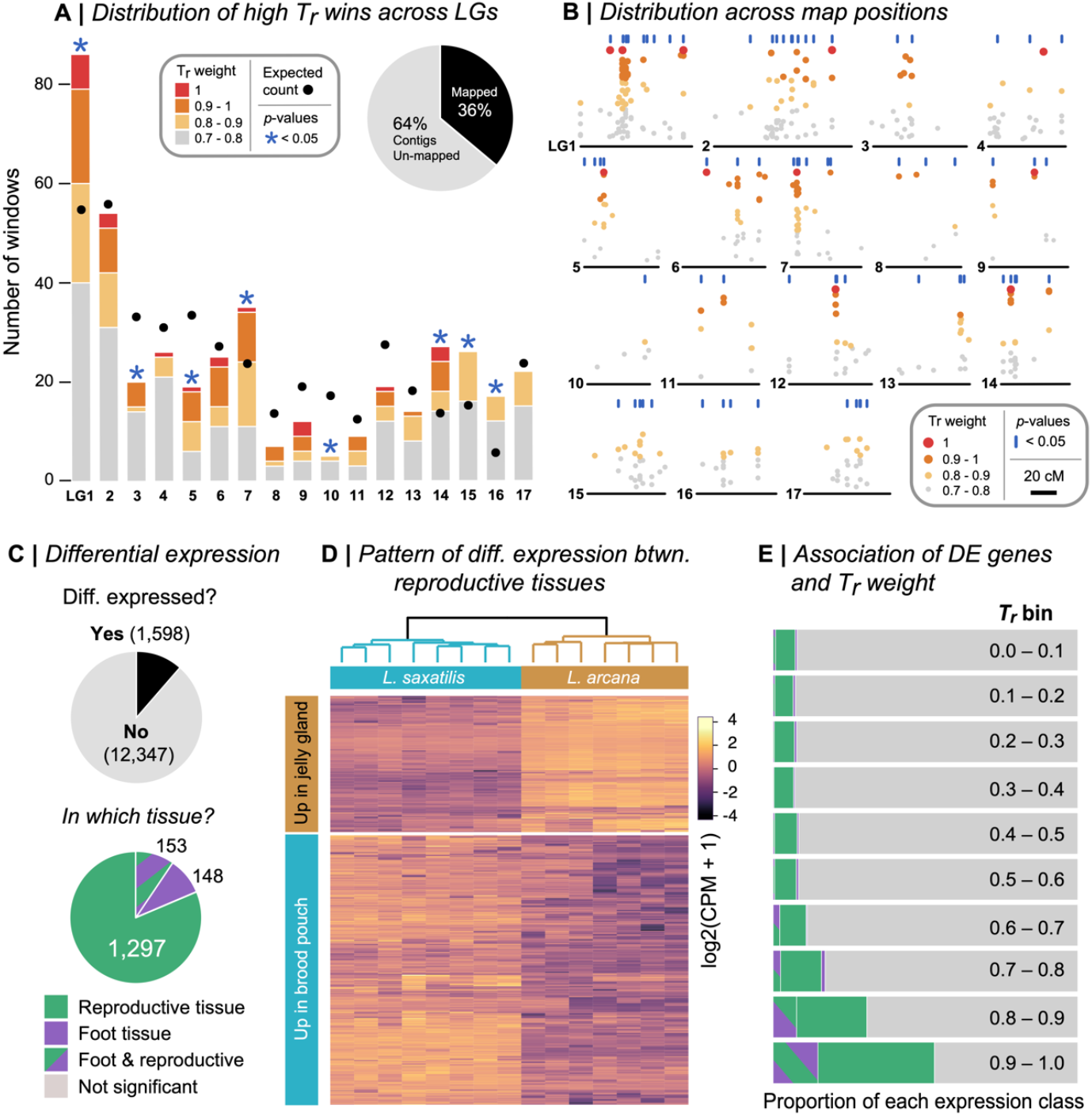
Candidate regions are widespread across the genome and enriched for genes that are differentially expressed between reproductive systems. (A) The number of high Tr windows (Tr > 0.7) assigned to each of the 17 *L. saxatilis* LGs. The circles show the expected number given the total assigned of windows to each LG. Asterisks indicate when the observed number is unlikely to be recovered by chance (p < 0.05). (B) Distribution of high Tr windows across LGs. Vertical blue lines indicate map positions that are enriched for high Tr windows. (C) Number of genes that showed differential expression (DE) and the number of DE genes in each expression class. (D) Clustering of reproductive tissue libraries based on patterns of expression. (E) The proportion of genes in each DE class after binning each gene according to the Tr weight.

Candidate regions also showed strong enrichment of genes that are differentially expressed between live-bearing and egg-laying reproductive tissues. To identify differentially expressed genes (DEGs), we collected reproductively mature female *L. arcana* and Northern *L. saxatilis* from a single location to control for environmental effects, and compared transcriptomes from whole reproductive systems (brood pouch vs jelly gland) and a non-reproductive control tissue (foot). We identified 1,598 DEGs, the majority of which showed differential expression between the reproductive tissues (1,297) (Fig. 4C, fig. S36). Of these, 66.1% (858) showed higher expression in the brood pouch of live bearers (Fig. 4D). To test for the enrichment of DEGs in regions associated with reproductive mode, we binned each DEG according to the Tr score of its associated genomic region (Fig. 4E). We found that the proportion of reproductive mode DEGs strongly increased with increasing Tr weight (Spearman’s rho = 0.903, p = 9e-04; table S12)

Gene ontology analysis and functional annotation suggest that the transition to live-birth involved genes with diverse functions. Separate GO analyses conducted on a sequence-based gene set (574 genes in regions where Tr > 0.7) and expression-based gene set (1,450 reproductive mode DEGs) yielded 37 enriched gene ontology terms, including transmembrane transport, calcium ion binding, and ion channel activity (Fig. S37). We examined the putative functions of the 27 genes found in both sets in more detail (table S13). These included genes putatively associated with antibacterial activity (lectin L6-like protein; higher expression in brood pouch), the synthesis of mucin-type oligosaccharides (GALNT10-like; higher expression in brood pouch), the formation of structural tissue (IFB-like and CMP-like, both higher expression in brood pouch), and two secretary genes involved in egg-mass production in another marine snail (both with lower expression in brood pouch).

## Conclusions

Our analyses show that live-bearing, a key innovation, is associated with selection on many loci, as in the only comparable analysis in *Zootoca* lizards (*19*). Although our genome-wide analysis hinted at two independent origins of live-bearing, the alleles associated with this trait have a single origin and have spread across space and genetic background. We found evidence that selection has acted on differences in gene expression, driving the origin of the live-bearing brood pouch. Other associated loci may underpin physiological changes that contribute to the difference in mode, such as differences in embryo retention-time (*20*), or may underlie other adaptations the became beneficial as live-bearing spread. Approximate estimates of the timing of selective sweeps at live-bearing loci, based on the accumulation of private mutations (*T* = π_w_/2*μ*), span a broad range from ∼20 k to 200 k generations before present, with a median time of 70 k generations BP (roughly 35k years BP, assuming 2 generations per year) (fig. S38). Thus, our results suggest that alleles associated with live-bearing accumulated gradually over the last 100 k years. These findings are relevant to the long-standing debate about the genetic basis of evolutionary novelty. Because key innovations are not visible to selection before they arise, models of saltational evolution invoke large-effect macromutations to explain their sudden appearance (*21*). However, our results show that innovation can have a polygenic basis, and suggest that novel functions can arise as the end product of sustained quantitative evolution, rather than in a single evolutionary step.

## Acknowledgments

Juan Galindo, Mauricio Montaño-Rendón, Natalia Mikhailova, April Blakeslee, Einar Arnason and Petri Kemppainen provided samples. Richard Turney, Graciela Sotelo, Jenny Larsson, Thomas Broquet and Stéphane Loisel helped with sample collection and processing. We thank Sci Ani for providing and allowing us to modify the cartoons shown in Fig. 1. Mark Dunning helped develop bioinformatic pipelines. Rui Faria, Hernan Morales and Vitor Sousa provided helpful discussion. Matthew Hahn, Jon Slate, Mark Ravinet, Joost Raeymaekers, Aaron Comeault and Nick Barton gave feedback on a draft version of the manuscript.

## Author contributions

Conceptualization: SS, AMW, KJ, RKB

Analysis: SS, ZBZ, MG, AP, DS, DGC, EL, JR, ALM

Writing – original draft: SS, RKB

Writing – review & editing: SS, ZBZ, MG, AP, DS, DGC, TB, ALM, EL, JR, KJ, AMW, RKB

## Data and materials availability

Sequence data are available on the short-read archive (SRA). All other data and analysis scripts are available on Github at https://github.com/seanstankowski/Littorina_reproductive_mode.

## List of supplementary materials

Materials and Methods

Supplementary Text Figs. S1 to S37

Tables S1 to S13

References (*18-62*)

