## Supplement for "Selection on many loci drove the origin and spread of a key innovation"

**This PDF file includes:**

Materials and Methods
Supplementary Text
figs. S1 to S38
tables S1 to 14

### Materials and Methods

Study system

The three species of *Littorina* examined here all reproduce by direct development (*i.e.,* they do not have larval dispersal), but differ in their mode of female reproduction (*8*). Egg-laying species (the ancestral reproductive mode) have an anatomical structure called a jelly gland (Fig. 1A, fig. S1A), that embeds egg capsules into a sticky mass that is attached to the substrate. Once laid, the egg mass can be as large as the snail that laid it and can contain hundreds of fertilized eggs with synchronized development occurring outside of the mother (Fig.1A, fig. S1B). In contrasts, in *L. saxatilis*, the only species of the genus that is live-bearing (the derived reproductive strategy), have an anatomical structure called a brood pouch, where fertilized eggs are deposited, pass through a larval stage, metamorphose, hatch and develop before finally creeping out of the mother (Fig.1A, fig. S1C) (Fig.1A, fig. S1C). In fully mature *L. saxatilis*, the contents of the brood pouch usually contain a complete continuum of embryonic development (*22*), from very early embryos through to fully mobile miniature snails that crawl out of the mother (Fig.1A, fig. S1D). Crawl-away offspring experience no maternal care once they leave the mother (*8*).

The jelly gland and brood pouch are homologous anatomical structures despite the striking functional differences between them (*8*). In reproductively immature females, it is not possible to reliably determine whether the structure will develop into a brood pouch or jelly glad. However, as the snails develop, the differences become very clear, and are easy to distinguish when snails are in reproductive condition.

In addition to the differences in their reproductive organs, egg-laying and live-bearing snails show differences in their timing of reproduction throughout the year (38, 39). The egg layers *L. arcana* and *L. compressa* reproduce during a defined breeding season beginning in the early autumn and peaking in the winter months (37, 38). In summer, few individuals can be found in reproductive status. The jelly glands of females atrophy outside the breeding season, and most males 'shed' their penis, and re-grow it before the next breeding season. In contrast with this seasonal reproductive behavior, *L. saxatilis* mate and reproduce year-round (37, 38). These differences in mating behavior can be seen as part of a difference in the ecological niche occupied by egg-layers and live-bearers (37, 38).

Although live birth is a recent innovation, the live-bearing species, *L. saxatilis*, has a much larger geographic distribution than any egg laying species of *Littorina* (*8*) (Fig. 1C). *L. arcana* and *L. compressa* are only found in the UK, Ireland, the northern coast of France and in parts of Norway and Russia (*8*). The distributions of both egg-layers, *L. arcana* and *L. compressa* are very similar, with the two species almost always found in the same geographic regions. By contrast, the distribution of *L. saxatilis* is much broader. Natural populations can be found in all areas where egg-layers are present, but also in many areas where they are absent (*8*). This includes the Iberian Peninsula, Sweden, Iceland, Greenland, the northeastern coast of the USA, and Russia. Other populations, thought to be human introductions, have been found in Morocco (Africa), South Africa, the Bay Area on the west coast of the USA, and Venice (Italy) (*8*).

Where the egg-layers *L. arcana* and *L. compressa* are present, they are almost always found in sympatry with *Littorina saxatilis* (*8*). For example, in areas that are exposed to strong wave action (fig. S2, wave habitat), *L. arcana* is syntopic with the wave ecotype of *L. saxatilis*, with both species often inhabiting the same rock crevices. In areas where the risk of crab predation is high (fig. S2, crab habitat), *L. compressa* is syntopic with the crab ecotype *of L.*

*saxatilis*. Because of these common selection pressures, sympatric egg-laying and live-bearing snails have converged on very similar phenotypes that are typically observed across gradients of crab-wave exposure (*23*). For example, in the wave habitat, *L. arcana* and *L. saxatilis* are so similar that mature females can only be identified by dissection and males cannot be reliably identified without diagnostic genetic markers (*8*).

Sample collection and identification

We obtained samples of *Littorina saxatilis, L. arcana* and *L. compressa* from 18 locations across the northern hemisphere, attempting to capture as much of their geographic distribution and phenotypic variation as possible (table S1; fig. S3). Samples were either collected by the authors or were kindly donated by experienced collectors (See acknowledgments). A detailed table of sample locations with coordinates, sample numbers, and species and ecotype designations can be found in table S1.

After collection, live samples were dissected either by the authors or by experienced collectors to determine sex, species and reproductive status based on internal anatomy according to Reid (*8*). Because *L. saxatilis* and *L. arcana* can only be reliably distinguished based on the female reproductive system, we only sequenced reproductively mature females at sites where the species are known to co-occur. Foot tissue was stored in 99% ethanol prior to the extraction of DNA.

DNA extraction, library preparation and sequencing

DNA was extracted from a small piece of foot tissue (2 x 2 mm) using a CTAB protocol (*24*) and checked for quality using gel electrophoresis, and concentration using a fluorometer at the University of Sheffield, UK. TrueSeq DNA Nano gel-free sequencing libraries (350 bp insert) were prepared by Edinburgh Genomics at the University of Edinburgh and sequenced on a HiSeq X (150 PE) to an expected depth of 15 x coverage.

Filtering and mapping of reads, variant calling and post-call filtering

*Associated code:*
Variant calling
https://github.com/seanstankowski/Littorina_reproductive_mode/tree/main/variant%20calling
Post-call filtering
https://github.com/seanstankowski/Littorina_reproductive_mode/tree/main/post%20call%20filtering

Sequencing adaptors were removed and sequences were trimmed for low quality using Trimmomatic (*25*) with the parameters recommended by the authors. Raw reads were filtered for quality and mapped to the *Littorina saxatilis* V.1 reference genome as described in Westram et al. (*26*) using the BWAmem algorithm (*27*). PCR duplicates were marked using the bammarkduplicates tool in biobambam2 (*28*). Read groups were assigned using the AddOrReplaceReadGroups tool in Picard toolkit (Broad Institute).

Variant calling was performed using GATK 4.2.5.0 (*29*) by executing steps documented in the short variant discovery pipeline. Briefly, HaplotypeCaller was used to call SNPs and indels and produce a gVCF file for each sample using the --include-non-variant-sites flag to facilitate the calling of invariant sites. To make this feasible with a large, highly fragmented reference genome, we performed this step on subsets of 1000 assembly contigs to produce 389 gVCFs for each individual. GenotypeGVCFs was then used to perform joint genotyping across the samples, to produce a multi-sample VCF for each subset of contigs. The multi-sample VCFs were then concatenated together using bcftools to produce a complete VCF including all variants and individuals.

After variant calling, we filtered the call set using bcftools (*30*). We applied soft and hard filters to the data to remove poorly sequenced and low-quality sites (table S2). First, we applied a set of hard filters to remove sites below a certain threshold based on the Fisher Strand (FS > 60.0), Strand Odds Ratio (SOR>3), RMS Mapping Quality (MQ<40), Mapping Quality Rank Sum Test (MQRankSum < -12.5), Quality by depth (QD<2.0) and Read Position Rank Sum Test (ReadPosRankSum<-8.0'). We also dropped (*i*) sites within 5 base pairs of an indel and complex variants, (*ii*) multi-allelic SNPs, and sites with 2.5 times the mean depth (> 3750 reads). The corresponding change in the number of sites after each step is available in table S2.

Before deciding on final filters for the depth, genotype quality and missing data, we applied a preliminary round of soft filters with fairly relaxed criteria (table S2). Specifically, we set genotypes with fewer than 5 reads (FMT/DP < 5) and a genotype quality less than 20 (FMT/GQ < 20') to missing, and then removed all sites with more than 20 % missing data.

Because the probability of calling a variant site increases with the per-site coverage, we calculated and compared the distributions of depth per site for variant and invariant sites separately (fig. S4A). This analysis revealed that the distribution of depth was quite different for the variant and invariant sites at lower depths, because sites with fewer reads are more likely to be called as invariant. Specifically, we observed a relative excess of invariant sites at depths between 750 – 2000 reads across all samples (fig. S4A). This was highlighted by plotting the ratio of invariant to variant sites across total depth. This ratio was extremely high and variable at lower total depths, though decreased precipitously as the total depth per site increased. Eventually, the relationships reached a stable plateau we take to represent the 'true' ratio of invariant to variant sites (fig. S4B).

Informed by this result, we (*i*) dropped all sites with a minimum total depth below 1500 (INFO/DP<1500), (*ii*) set all genotypes with fewer than 10 reads to missing, and then (*iii*) dropped all sites with more than 10% missing data. This drastically improved the depth distributions for variant and invariant sites such that the bias towards calling invariant sites at lower depths was substantially reduced (fig. S4C). The final data set included 18,537,198 SNPs and 310,294,741 invariant positions.

Genome-wide phylogenetic inference and principal components analysis

*Associated code:*
https://github.com/seanstankowski/Littorina_reproductive_mode/tree/main/genomeWide_relationships

We used RAxML v8 (*31*) to reconstruct the evolutionary relationships among the 108 samples from a concatenated alignment of all 18,537,198 SNPs and an ascertainment bias correction to account for invariant sites (Fig. 1E, fig. S5). The VCF file was converted to *phylip* format using the script *vcf2phylip* (*32*). The ML phylogenetic analysis was conducted using the HPC-PTHREADS-SSE3 of RAxML with the GTRGAMMA model. Five runs of RAxML were conducted, and each produced an identical topology. We tested the support for each node using 100 bootstrap replicates and with the rapid bootstrapping algorithm. The topology was rooted with *L. compressa* (*7*) and drawn with Figtree 1.4.4 (*33*).

In addition to the tree-based approach, we used principal components analysis (PCA) to infer relationships between the samples based on genome-wide data (fig. S6). We used the same dataset as described for the phylogenetic analysis except that we only analysed markers with <

5% missing data (16,658,016 SNPs). We used *plink* v. 1.9 (*34*) to convert the VCF file into plink format, and then conducted four separate PCAs. The first 2 PCAs were conducted with the full set of individuals, but with different numbers of markers: one dataset included all SNPs, while the other was thinned to reduce LD, yielding 5,742,754 SNPs. The LD thinning was performed in plink by removing SNPs that had an $r^2$ value greater than 0.1 with any other SNP within a 50-SNP sliding window that was advanced by 10 SNPs each time.

<u>Local tree building and topology weighting</u>

*Associated code:*
Tree building and topology weighting
https://github.com/seanstankowski/Littorina_reproductive_mode/tree/main/Twisst_popgenWins
Drawing of unrooted high $T_r$ trees from Newick format
https://github.com/seanstankowski/Littorina_reproductive_mode/tree/main/down_stream_analyses_and_ploting/scripts_for_drawing_unrooted_trees

We used local tree building and topology weighting to characterize patterns of genealogical variation across the genome, following a custom pipeline established by Martin and Van Belleghem (*13*). The following pipeline was run twice independently from the phasing all the way through to the topology weighting, and produced highly similar results.

We first phased our genotype data using Beagle 5.3 (*35*), with the settings recommended in the manual for a large outbred population. Each assembly contig was then divided into non-overlapping genomic windows that each contained 100 SNPs. This yielded 154,971 genomic windows, with a mean physical length of 5.8 kb ± s.d. 5.3 kb (fig. S7). A neighbor joining tree was then constructed for each genomic region using the program *Phyml* (*36*). We assumed a GTR model and inferred trees using the settings suggested by Martin and Van Belleghem (*13*). Selected trees were illustrated using the *R* package *ggtree* (*37*) using the unrooted layout.

Topology weighting was then performed on each of the 154,971 trees using the program *Twisst* (*13*). Topology weights are values that describe the degree of monophyly in a large genealogy in light of the possible monophyletic taxon-level relationships that could be observed. Consider a large genealogy for a single non-recombining locus with 100 sampled individuals, but only four taxa: 1, 2, 3, 4 (Fig. 2A, fig. S8). Although there is an inordinate number of possible unrooted subtrees that one can observe given the large number of tips, there are only 3 possible unrooted topologies that can be observed if we randomly sample a subtree from the full tree that includes only one tip of each taxon: ((1,2)(3,4)), ((1,3)(2,4)), ((1,4)(2,3)). If the topology of many random four-taxon subtrees (say 10,000) is determined by iterative sampling of the full tree, then the fraction of each subtree gives a quantitative description of the degree of monophylly toward each of the three possible taxon subtrees.

Topology weighting was performed using the Newick tree files as the input, with the following taxon-level topologies specified for calculating the weights (Fig. 2C). Given that we have four taxa in our genome-wide phylogeny (*L. compressa* = C; Spanish *L. saxatilis* = S, *L. arcana* = A, and Northern *L. saxatilis* = N), the three possible unrooted taxon topologies are: Tb = ((c,s)(a,n)), Tr = ((c,a)(s,n)) and Tc = ((c,n)(s,a)) (Fig. 2A). Tb—the background topology—is consistent with the taxon-level relationships that we observed in our genome-wide tree (but note that the full topology will not necessarily be exactly the same as background tree because relationships may differ within each of the four groups). whereas Tr and Tc represent possible alternative relationships. In Tr—the reproduction tree—the taxa group according to their reproductive mode, whereas in Tc—the 'control' tree—the taxa do not cluster as expected based on their geographic relationships or any known aspect of their biology. We refer to Tc as the control topology, *sensu* Martin et al. (*38*), because it provides a useful control for distinguishing incomplete lineage sorting from other processes that can generate discordant gene trees.

Because each of the 154,971 trees is large, consisting of 2*n* = 216 tips, we calculated the weights based on a fixed number of subtrees rather than performing an exhaustive search of all possible quartets. We sampled 10,000 subtrees, which has been shown to give very narrow limits on the true topology weights (*13*).

#### Plotting of topology weights in a ternary plot

Rather than plotting the topology weights across the genome (*13*), we took the novel approach of analyzing the joint distribution of topology weights within a ternary plot (Fig. 2A). The ternary plot has been used to study concordance in phylogenetic studies (*39*), and is also a natural framework for analyzing the distribution of weights in a tree with four taxa, as it makes possible a graphical to representation of each genomic window as a single point in an equilateral triangle based on the three topology weights.

The three corners of the ternary plot—[1,0,0], [0,1,0], [0,0,1]—correspond with genomic windows that show taxon-level relationships that are consistent with one the three possible subtrees; that is, 100% of the sampled subtrees (in our case 10,000) perfectly match one of the three alternative trees, implying that samples from each of the four groups are monophyletic (Fig. 2A). In contrast, the very center of the ternary plot—[0.33,0.33,0,33]—corresponds with a genomic window where all three of the possible subtrees were sampled at equal frequency. Any other location in the ternary plot indicates a bias toward one of the subtrees, but with some resemblance to at least one of the other alternative topologies.

#### Exploring the ternary distribution with simulations

*Associated code:*
Simulations of topology weights in a ternary framework
https://github.com/DaSh-bash/LittorinaBrooding
Ternary analysis and plotting of simulated data
https://github.com/seanstankowski/Littorina_reproductive_mode/tree/main/MSPrime_sims

To understand how different evolutionary processes shape the distribution of topology weights in the ternary framework, we simulated genealogies for non-recombining haplotypes in a four-taxon framework using MSprime 1.2.0 (*40*, *41*). We specified a demography where four descendent populations—O, P3, P2, P1—are produced by population splits at three points going backwards in time (table S3): T1, which splits population P12 to give rise to descendent populations P1 and P2; T2, which gives rise to the populations P12 and P3; and T3, where the common ancestor of all the descendent population splits, giving rise to lineages O and P123. The size of each population ($X$) is set to $N_eX$ haploid sequences, and uni- or bi-directional migration can occur between a single pair of ingroup taxa (*i.e.*, between P2 and P3). Simulated genealogies were visualized in demesdraw 0.3.2 (*42*)

We varied these 12 parameters to produce 101 unique models that fall into the 9 scenarios outlined below (*a* – *h*, fig. S9; table S4). For each model, we simulated 10,000 coalescent trees, which were then output in Newick format using tskit 0.5.4 (*43*), and passed to the *Twisst* algorithm for topology weighting as described above. The topology weights were plotted in a ternary plot using the R library *Ternary*. The results of the simulations are shown in figs. S10-S18, and described in detail in the supplementary text.

a. *Uniform $N_e$ and split times:* We conducted 12 simulations where we varied the number of generations between the three splits, while keeping populations identical in size. These simulations range from very short durations between splits, mimicking a scenario with almost no demographic separation (T1 = 1, T2 = 2, T3 = 3), through to very long durations between splits relative to the $N_e$ ($T_1$ = 7290, $T_2$ = 14580, $T_3$ = 21870). $N_e$ was set to 500 for all populations and no migration was allowed between them
b. *Varying but equal split times for T2 and T3, with T1 set to 5k generations:* We conducted 8 simulations where we varied the durations between splits T2 and T3, but fixed T1 at 5000 generations to allow populations to diverge after all 3 splits had occurred. $N_e$ was set to 500 for all populations across all simulations. No migration was allowed between populations.
c. *Uneven split times:* We conducted 10 simulations where the number of generations between population splits varied in evenness. This was done by fixing the values of T1 and T3, and varying the time of T2. We used 2 combinations of T1 and T3 (90 and 270 and 405 and 1215) to see how the unevenness of splits interacted with variation in the duration of divergence. $N_e$ was set to 500 for all populations across all simulations with no migration between populations.
d. *Uniform variation in $N_e$:* We performed 10 simulations using a model with the same, equal split times but with different values of $N_e$. The split times used for all populations were $T_1$ = 90, $T_2$ = 180, $T_3$ = 270. We varied the $N_e$ from 5 to 5000, keeping it uniform for all populations. No migration was allowed between populations.
e. *Varying $N_e$ in one population:* We performed 12 simulations using a model with equal split times, but with the $N_e$ of one population (always P3) set to a value that was larger or smaller than all other populations where $N_e$ was always 500. This was done for 4 different sets of split times, ranging from short durations between splits ($T_1$ = 10, $T_2$ = 20, $T_3$ = 30), to very long durations ($T_1$ = 7290, $T_2$ = 14580, $T_3$ = 21870). Four different values of $N_e$ were tested for each scenario. No migration was allowed between populations.
f. *Unidirectional migration between P2 and P3:* We conducted 16 simulations with varying rates of migration between populations P2 and P3. We defined four histories with the same $N_e$ for all populations ($N_e$ = 500) and equal split times, but with duration between splits varying from very short ($T_1$ = 10, $T_2$ = 20 $T_3$ = 30) to very long ($T_1$ = 7290, $T_2$ = 14580, $T_3$ = 21870). For each of the four histories, we modelled four different rates of migration: $m$ = 0, 0.001, 0.010 or 0.100.
g. *Bidirectional migration between P2 and P3:* We conducted 16 simulations as described above for the unidirectional migration scenario, but with migration in both directions (P3 to P2 and P2 to P3).
h. *Unidirectional migration for 10% of the genome:* We conducted 12 simulations where migration only occurs for a fraction of the genome. We defined four histories with an $N_e$ of 500 for all populations and equal split times, but with the duration between splits varying between the two scenarios ($T_1$ = 100, $T_2$ = 200 $T_3$ = 300; $T_1$ = 405, $T_2$ = 945, $T_3$ = 1215). For each scenario, we modelled heterogenous migration by combining two simulations together. 90% of genealogies were simulated under a 'background' demography without gene flow, and the remaining 10% were simulated under the same model, but one of seven rates of migration from P3 to P2: $m$ = 0, 0.001, 0.005, 0.010, 0.05, 0.01, or 0.05.
i. *Ancestral structure:* We modeled ancestral structure by combining simulations of two scenarios together, following Martin et al. (*44*). Ninety percent of the genome was simulated under the 'background' demography (O(P3(P2,P1))), and the remaining 10% under the

'alternate' demography (O(P2(P3,P1))). Split T2 was set to be more ancient in the background to mimic a region of the genome that is polymorphic at particular loci. $N_e$ was set to 500 for all populations for all simulations. No migration was allowed between populations.

Quantifying asymmetry with the $D_{LR}$ statistic

*Associated code:*
Ternary analysis and plotting of simulated data
https://github.com/seanstankowski/Littorina_reproductive_mode/tree/main/MSPrime_sims

The results of simulations outlined above show that divergence under an idealized four-population model without migration produces a symmetrical distribution of topology weights between the left and right halves of the ternary plot (figs. S10 & 14; Supplementary text). This is because there is an equal chance that a given gene tree will more closely resemble either alternative topology under incomplete lineage sorting (ILS) (*14*). However, deviations from a simple four population model (*e.g.*, gene flow) can lead to a bias in the probability toward one alternative topology, leading to an asymmetrical distribution of topology weights (figs. S14 & S18; Supplementary text).

To measure genealogical bias, we developed a statistic that can be used to detect and quantify asymmetry in the ternary framework. $D_{LR}$, which is similar to Patterson's D statistic (*45*), can range from negative 1 to positive 1, and gives and an indication of the strength of the bias toward one alternative topology relative to the expectation of equality ($D_{LR} = 0$). A genome-wide estimate of $D_{LR}$ can be calculated directly from the topology weights as:

$$D_{LR} = \frac{(n_{\mathrm{T2} > \mathrm{T3}}) - \left(0.5\left((n_{\mathrm{T2} > \mathrm{T3}}) + (n_{\mathrm{T2} < \mathrm{T3}})\right)\right)}{0.5\left((n_{\mathrm{T2} > \mathrm{T3}}) + (n_{\mathrm{T2} < \mathrm{T3}})\right)} \qquad \text{(eq. 1)}$$

Where T2 and T3 are the two alternative topologies (see the notation for topologies in Fig. 2A), $n_{\mathrm{T2}} > n_{\mathrm{T3}}$ is the number of windows where T2 is greater than T3, and $n_{\mathrm{T3}} < n_{\mathrm{T2}}$ is the number of windows where T3 is greater than T2. Because these quantities indicate the number of windows in the left and right halves of the triangle, equation 1 can be expressed more intuitively as:

$$D_{LR} = \frac{n_L - (0.5 n_{\mathrm{T}})}{(0.5 n_{\mathrm{T}})} \qquad \text{(eq. 2)}$$

Where $n_L$ is the number of trees on the left side of the plot and $n_{\mathrm{T}}$ is the sum of the trees on the left and right sides. Under ILS the expected number of windows in each half of the plot is simply half of the total number of windows (*i.e.*, $0.5n_T$). Note that $n_{\mathrm{T}}$ can be a slightly different number from the total number of windows analyzed because some may fall precisely on the line that bisects the base of the ternary plot, such that they are neither left- nor right-sided (*i.e.*, when T2 = T3).

In addition to a genome-wide estimate (*i.e.,* comparing the full left- right- half triangles), symmetry can also be quantified for any arbitrary area of the of the distribution, provided that analogous sections from both sides of the ternary plot are compared. Detailed examples of genome-wide and partial estimates of $D_{LR}$ are shown in fig. S19.

*Advantages over existing site-based statistics*

Why use topology-based estimates of genealogical asymmetry when site-based methods already exist? The framework presented here has many advantages over existing methods.

- *No need for a defined outgroup*—Site-based statistics like Patterson's D require one or more appropriate outgroup taxa that are used to define alleles at each SNP as ancestral or derived. This is challenging (usually requires multiple outgroups for high confidence assignment), and assumes that there is no allele sharing due to gene flow or ancestral structure between the ingroup and outgroup taxa. This is not required in the framework presented here, as estimates of symmetry are inferred from how samples cluster in a tree topology.
- *Site-based statistics are low dimensional summaries of topologies*—Given that the underlying genealogies are what we care about, it makes more sense to study them directly, rather than variation at individual sites.
- *There is a movement away from site-based analyses to those based on genealogies*—Inference tools like Tsinfer (*46*), Relate (*47*), and ARGweaver (*48*) are making it possible to infer full genomic sequences of genealogical trees from genome-wide data. Unlike the arbitrary windows that are used in most genomic studies, the genomic spans of the trees inferred with these methods have biological meaning, as their boundaries are defined by recombination events that occurred in the past (*i.e.*, the windows are not arbitrary). Topology weighting can be performed directly on these marginal trees (as described in the section 'Inference of ancestral recombination graphs', pp. 26-27, and as show in Fig. 3E), and the $D_{LR}$ statistic can be calculated directly form the distribution of topology weights.
- *Can be calculated for systems with more than four taxa*—The $D_{LR}$ statistic can be extended to higher dimensions. Consider a tree with 5 taxa, instead of the 4 analyzed here. There are 15 possible unrooted topologies, including the one 'background' topology that matches the demographic history of the populations, and 14 alternative trees. In an idealized four population model, where only ILS generates discordant trees, we expect bias toward these 14 topologies to be equally likely for a randomly drawn locus. Thus, for each of the alternative genealogies, we expect it to be the best-fitting one for 1/14 ≈ 7.14% of loci, providing that the number of loci is large. The significance of the observed deviation from this fraction can be determined using a G-test (or similar).

Plotting and symmetry analysis of the empirical ternary distribution

*Associated code:*
Ternary analysis and plotting of empirical data
https://github.com/seanstankowski/Littorina_reproductive_mode/tree/main/down_stream_analyses_and_ploting/ternary_plot_figure

Because of the large number of genomic windows analysed, we plotted the empirical distribution of weights using a ternary histogram, where the triangle is divided up into equal-sized hexagons that are colored to indicate the number of windows that fall into that area of the ternary distribution. This was constructed in the R package ggtern (*49*) using the *geom_hex_tern* function.

We tested for asymmetry of the distribution between the left and right side of the diagrams by computing $D_{LR}$ as described above (eq. 1). We used a permutation test (99,999 permutations) to calculate the probability of obtaining the empirical estimate of $D_{LR}$ by chance, assuming that there is an equal probability of a window falling on either side. (Main text, Fig. 2D). To understand how symmetry varies more locally across the ternary distribution, we subdivided each side of the ternary plot into 45 sub-triangles (fig. S20), calculated $D_{LR}$ separately for each one, and used permutation tests to determine the significance of each estimate. Permutation tests were conducted in R using custom scripts.

Contributions of chromosomal inversions to asymmetry

*Associated code:*
PCA on inversion LGC 2.1.
https://github.com/ja-Reeve/Littorina_inversion_identification/blob/main/R_scripts/PCA_of_inversion_area.R.
Effects of inversions on asymmetry
https://github.com/seanstankowski/Littorina_reproductive_mode/tree/main/data/Master_data_file

Previous work has identified 17 large putative chromosomal inversions in *Littorina* (*50*), many of which segregate in the species studied here (Reeve et al. in prep.). Because inversions are large and experience reduced recombination, differences in the frequency of different orientations among the four clades may have a disproportionately large effect on the ternary distribution.

To determine the effect of known chromosomal inversions, we identified windows associated with each inversion using a previously published genetic map (*26*) and established coordinates for the inversion boundaries (*51*). We initially assigned each window to one of three groups (*i*), within an inversion, (*ii*) within a colinear region, and (*iii*) uncertain. The uncertain group, which contained a relatively small number of windows, includes windows that fall very near to inversion breakpoints, the precise locations of which are unknown. These windows were not considered in subsequent analyses.

To test for a general effect of inversions on asymmetry, we first calculated $D_{LR}$ only on windows in colinear regions (fig. S21). The result was then compared with the estimate obtained for the full dataset, which included all windows (Fig. 2D, fig. S21). We followed up by calculating $D_{LR}$ for each individual inversion separately, to quantify their individual contributions to the distribution of weights (fig. S22 & S23). For one inversion that showed fairly strong asymmetry, we used Principal Components Analysis in the R package *Adegenet* (*52*) to determine the frequency of the alternative arrangements in each of the four clades (fig. S24). This is possible because, within inversions, individual samples are expected to cluster into three distinct groups that correspond with the separate karyotypes (*53*). The results of these analyses are described in detail in the supplementary text.

Estimating diversity, divergence and differentiation in 100 SNP windows.

*Associated code:*
Calculation of population genetic stats
https://github.com/seanstankowski/Littorina_reproductive_mode/tree/main/Twisst_popgenWins/genomics_general-master
Statistical tests and plotting scripts
https://github.com/seanstankowski/Littorina_reproductive_mode/tree/main/down_stream_analyses_and_ploting/diversity_divergence_high_Tr_trees

We estimated a set of summary statistics in the same 100 SNP genomic windows that were used for tree construction and topology weighting (Fig. 3A, figs. S25—S27). This included nucleotide diversity within groups ($\pi$) and nucleotide divergence between groups ($d_{xy}$). The calculations were performed using the popgenwins.py script available at https://github.com/simonhmartin/genomics_general (26).

We used two different types of groupings for these calculations. First, we grouped samples by taxon, computing $\pi$ within *L. compressa*, Spanish *L. saxatilis*, *L. arcana*, and Northern *L. saxatilis*, and $d_{xy}$ between each pair of taxa. Second, we grouped samples by their reproductive mode, which meant pooling the egg-layers (*L. arcana* and *L. compressa*), and live-bearers (Spanish *L. saxatilis* and Northern *L. saxatilis*), and calculating statistics within and between these groups.

We used paired Wilcoxon signed-rank tests, performed in R, to compare median diversity between egg-layers and live-bearers. This was done for all windows to compare genome-wide levels of diversity, and then for windows perfectly associated with reproductive mode. Ridgeline plots and Violin plots of diversity were produced in R using ggplot2.

<u>Relationship between genetic diversity, differentiation and topology weights</u>

*Associated code:*
Statistical tests and plotting scripts
https://github.com/seanstankowski/Littorina_reproductive_mode/tree/main/down_stream_analyses_and_ploting/diversity_divergence_high_Tr_trees

We tested for the relationships between of π, $d_{xy}$ and all three of the topology weights (Tb, Tc, and Tr) for the 100 SNP windows that had weights greater than 0.6. To summarize general patters, we calculated a weighted average of π across the variation in the topology weights using the ksmooth function in R. Spearman rank correlations were computed between selected variables in R to estimate the strength and significance of any relationships.

<u>Calculation and analysis of the folded SFS</u>

*Associated code:*
Analysis of private alleles and folded SFS
https://github.com/seanstankowski/Littorina_reproductive_mode/tree/main/012plot_privateAlleles

To test for evidence of positive selection in regions of the genome associated with reproductive mode, we calculated the folded SFS for egg-layers and live-bearers by determining the distribution of the counts of minor alleles over all sites segregating within each group. For regions targeted by positive selection, we expect to see an SFS that is skewed toward an excess of rare variants (*17*), translating to a negative estimate of Tajima's D (*54*).

To do this, we first used BCFtools (*30*) to subset the full VCF file producing a reduced version that only contained regions of the 50 contigs where Tr = 1. We then used VCFtools (*55*) to convert the VCF into a matrix of genotypes using (--012 format). A visual representation of the genotype matrix can be seen in fig. S28. This already clearly shows reduced polymorphism in live-bearers compared with egg-layers, as well as a bias toward rare variants in live-bearers.

Separate folded SFS were calculated for four different groups of samples: all egg-layers ($n$ = 28), all live-bearers ($n$ = 80), all Northern *L. saxatilis* ($n$ = 68), and all Spanish *L. saxatilis* ($n$ = 12). The full SFS for each group is shown in fig. S29. We calculated Tajima's D (*54*) for each group, directly from the folded SFS using a custom script in R. We estimated 95 % confidence intervals around the estimates of Tajima's D by jackknife resampling using custom R scripts. The results of these analyses are available in fig. S29 & S30 and table S9.

We also used custom R scripts to compare the SFS for different groups (*e.g.*, live-bearers vs. egg-layers and Northern *L. saxatilis* vs. Spanish *L. saxatilis*) to determine if they were significantly different. Because the sizes of the SFS differed between the groups being compared, we created sample-size-matched spectra by down-sampling the larger group to match the sample size of the smaller group. This was done by randomly sampling $n$ individuals from the larger group without replacement. The size-matched SFSs were then compared with a G-test in R. The random subsampling and statistical comparison were repeated 10,000 times. We also generated a randomly chosen size-matched spectrum for live-bearers to present in the main text, to facilitate its visual comparison with the SFS for egg-layers (Fig. 3B).

#### Analysis of private alleles

Associated code:
Analysis of private alleles and folded SFS
https://github.com/seanstankowski/Littorina_reproductive_mode/tree/main/012plot_privateAlleles

Positive selection is also expected to impact patterns of allele sharing between groups harboring ancestral and derived mutations. For example, if a new beneficial mutation arises on a single ancestral sequence, positive selection will purge diversity from it (*16*). This will cause the most variation in the group harboring the ancestral allele to become private to that group. However, new mutations will eventually begin to arise on chromosomes that descended from the new selected allele, and these will be exclusive to the group where the new allele becomes fixed.

To obtain estimates for the fractions of private alleles, we used custom R scripts to divide samples into pairs of groups (*i.e.,* live-bearers and egg-layers; Spanish *L. saxatilis* and Northern *L. saxatilis*), and identify alleles that were private to each group. However, because the probability of discovering rare alleles depends on the number of genomes sampled, we corrected for sample size by randomly down-sampling the larger group to the sample size of the smaller one before estimating the number of private alleles. The estimation of private alleles was repeated for 10,000 size-matched samples to produce distributions of (*i*) the number of private alleles in each group, (*ii*) the proportion of private alleles in each group, and (*iii*) the difference in the proportion of private alleles between the groups. Inferences were made from the mean and degree of overlap of these distributions. The results of this analysis are presented in fig. S31 and table S10.

#### Calculation of $\pi$, $d_{xy}$, and $F_{ST}$ across contigs

*Associated code:*
Window-based summary statistics
https://github.com/seanstankowski/Littorina_reproductive_mode/tree/main/down_stream_analyses_and_plotting/variation_within_contigs

We conducted sliding window analysis to explore patterns of variation across the 50 contigs that showed a perfect association with live birth. Specifically, for each contig, we calculated nucleotide diversity within egg-layers and live-bearers ($\pi$), nucleotide divergence between egg-layers and live-bearers ($d_{xy}$), and genetic differentiation ($F_{ST}$) between egg-layers and live-bearers. All statistics were calculated within 3kb overlapping windows separated by a 30 bp step. The calculations were performed using the popgenwins.py script available at https://github.com/simonhmartin/genomics_general (*38*). Figures for the relevant stretches of each contig are provided in fig. S33.

#### Inference of Ancestral Recombination Graphs

*Associated code:*
ARG inference in ARGweaver
https://github.com/seanstankowski/Littorina_reproductive_mode/tree/main/ARGinference

The genomic windows used in analysis up to this point (both in this paper and in evolutionary genomic studies in general) are essentially arbitrary, chosen to contain the same number of variant positions (*i.e.,* 100 SNPs). Ideally, we would partition the genome in a way that coincides the historical coalescence and recombination events that generated our observed set of genomes.

Here we do this by inferring Ancestral Recombination Graphs (ARGs) from our genomic dataset. The ARG is a graph structure that depicts the entire ancestral history of a set of genomes—including each recombination and coalescence event—that contains the genealogies for each non-recombined genomic block. Thus, by inferring the ARG, we can obtain tree topologies for each inferred non-recombining region, and more precisely infer the genomic span of haplotypes that are perfectly associated with live-birth.

We inferred ARGs using the program ARGweaver (*48*). We focused on two 30kb genomic regions (Contig1808:1-30000 and Contig3201:45000-75000) that spanned areas perfectly associated with reproductive mode. We focused only on these regions because the method is computationally expensive, and these regions are very well sequenced and assembled which is necessary for meaningful ARG inference. Due to computational limitations, we also restricted the analysis to a subset of 50 individuals (25 egg-layers and 25 live-bearers; *i.e.*, 100 haploid samples), as suggested by the authors of ARGweaver. These individuals were chosen to give equal and broad representation across the four clades.

For both contigs, we converted the SNP information from the VCF format to *sites* format, which is compatible with ARGweaver. The *sites* format only includes information on genomic positions that vary within the 100 samples. Positions that were absent in the original VCF were masked from being incorrectly treated as invariant sites while inferring ARGs.

ARGweaver was executed on both contigs using the following parameters: $N_e = 20{,}000$, $\mu = r = 1.5 \times 10^{-8}$. ARGweaver simplifies ARG inference by forcing coalescence and recombination events to occur at discrete timepoints. Here we used the 30 specified time points shown in fig. S34, with the maximum possible TMRCA set to 20*Ne* (fig S1). ARGweaver was run for 10,000 iterations, out of which the first 5000 iterations were considered as burn-in after visual examination of the MCMC traces and therefore, excluded from subsequent inference. From each of the 5000 remaining iterations, we estimated TMRCA for all the samples, as well as within live-bearers and within egg-layers. These estimates were used to determine the median and confidence limits for the TMRCA shown in main text Fig. 3E.

In addition to estimating the TMRCA, we also performed topology weighting on the marginal trees from the ARG. This allowed a more precisely identification of regions of the genome that were perfectly associated with live-birth (*i.e.*, trees where Tr = 1). To do this, we extracted all marginal trees from the last MCMC iteration (*i.e.*, iteration 10,000) and performed topology weighting as described in an earlier section. We used the combined information from the topology weights and TMRCA to identify core haplotype blocks associated with live-birth, as described in Shipilina et al. (*18*). Specifically, we used the topology weights and estimates of TMRCA to identify unique edges in the ARG that were both ancestral only to all live-bearing samples (Tr = 1), and that were defined by a single a coalescence time. Block inference was performed with a custom R script (*18*).

<u>Assignment of high Tr windows to map positions and significance testing</u>

Associated code:
*Permutation test and plotting scripts*
https://github.com/seanstankowski/Littorina_reproductive_mode/tree/main/down_stream_analyses_and_ploting/Tr_scores_across_LGs

We assigned 100 SNP windows to a previously published genetic map (*26*) in order to understand how associated regions were distributed across the genome. We first determined how many high Tr windows were associated with each of the 17 *L. saxatilis* linkage groups. We then used the Pearson correlation coefficient to test for a relationship between the number of high Tr windows assigned to each linkage group and linkage group size, defined here as the total number of windows assigned to each LG. We used permutation tests (9,999 permutations) for each LG to determine if they contained more or fewer high Tr windows than expected by chance. Briefly, for an LG containing *n* windows, we randomly sampled *n* windows at random from the full set of windows in order to generate a null distribution of high Tr counts. Permutation tests were also used to test each of 1,481 positions in the genetic map for enrichment of high Tr windows using the same logic as for each LG.

<u>Differential expression analysis</u>

Associated code:
*Differential expression analysis*
https://github.com/MartinGarlovsky/littorina_reprod_rnaseq

Samples for RNA sequencing were collected from Anglesey, Wales (fig. S1; table S1) during two trips (referred to as 'early' and 'late' trips below) in late Autumn of 2018. Snails were transported live to the University of Sheffield immediately after collection and kept at ~4$^{o}$C. Samples were dissected within four days of collection and identified as described above. We snap-froze reproductive and foot tissue samples in liquid nitrogen and stored them at -80$^{o}$C until the RNA extraction.

We extracted total RNA using TRIzol reagent (Invitrogen), cleaned the samples with the Turbo DNA-free kit (Ambion) to remove any residual DNA, and purified them with the RNeasy MinElute Cleanup Kit (Qiagen). All protocols followed the manufacturer's instructions. RNA quality and quantity were evaluated using RNA screen tape analysis. We only retained samples with RIN numbers over 7.

Rather than sequencing tissues from single individuals, we created pools of each tissue by combining 800 nanograms of RNA extracted from four different individuals. This included (*i*) 8 pools of RNA extracted from live-bearing reproductive systems, (*ii*) eight pools of RNA extracted from egg-laying reproductive systems, (*iii*) 3 pools of foot tissue from live-bearing samples, and (*iv*) 3 pools of foot-tissue from egg-laying samples. The individuals in each of the six foot-tissue pools correspond with the same set of individuals in one of the reproduction-tissue pools, thus controlling for the effects of individual. Samples were sent to Edinburgh Genomics for library preparation (Illumina TruSeq stranded mRNA-seq preps) and sequencing (NovaSeq 100PE sequencing).

We assessed the quality of the raw reads with FastQC and conducted quality trimming and adapter clipping using Trimmomatic v0.38 (*25*). Specifically, Illumina adapters were removed together with leading and trailing low quality or N bases (below quality 3), trimmed when the average quality per base dropped below 15 when scanned with a 4-base wide sliding window and removed if the reads were shorter than 36. We used FastQC (*56*) again to check the improvement of read quality. Reads were mapped to the *L. saxatilis* reference using HISAT2 (*57*) using parameters recommended by the authors. Next, we assessed mapping quality using SAMtools (*58*) and used StringTie (*59*) to obtain transcript count data for each gene. All analyses were performed on the University of Sheffield HPC, SHARC.

We first filtered the dataset by removing genes with fewer than 20 reads among all libraries, yielding 40,736 gene IDs. We also only retained genes that had at least 5 counts per million (*60*) in three or more replicates across any species/tissue combination. This reduced the number to 13,945 genes that were used in subsequent analysis. We used DESeq2 (*35*) to fit a model to the expression data that included the main effects of tissue and species, and the tissue-by-species interaction. The model also included a covariate term to account for samples where only reproductive tissue was harvested (earlier collection), and those where both foot and reproductive tissue were collected (later collection). Based on this analysis, we identified genes that showed differential expression between (*i*) egg-laying and live-bearing reproductive tissues only, (*ii*) egg-laying and live-bearing foot tissues only, or (*iii*) between both types of egg-laying and live-bearing tissues (fig. S36). Genes were considered differentially expressed based on a log2-fold-change > 2 and p-value < 0.01 corrected for multiple testing.

<u>Functional annotation and gene ontology (GO) analysis</u>

We conducted a functional annotation and GO-term enrichment analysis, focusing on two partly overlapping sets of genes: (set A) those identified as differentially expressed between the egg-laying and live-bearing reproductive tissues and (set B) genes within genomic regions associated with reproductive mode in our genealogical analysis (Tr > 0.7) (fig. S37). The gene identifiers for the best hits of the oyster *Crassostrea virginica* (gca002022765v4 from Ensembl) were used for functional enrichment using ShinyGO v0.75c (*61*) as this was the only mollusc species available. A false discovery rate (FDR) cutoff of 0.05 was used for both enrichment analyses and other parameters were also left as default.

For the RNAseq data (set A), fasta sequences corresponding to the expressed transcript were obtained from the gtf file using GffRead (*62*). Out of 13797 transcripts that were considered expressed for the RNAseq analysis, a corresponding gene from *C. virginica* was found for 8785 transcripts. These genes were used as the background for the enrichment analysis. Of the 1297 genes that were differentially expressed between the reproductive tissues, 639 had a corresponding gene in *C. virginica*; functional enrichment analysis was performed using these genes

For the reproductive mode-associated regions (set B), BedTools intersect (*63*) was used to find any genes from the gtf file that were contained within the selection region, and the above procedure was then used to obtain the transcript sequence. BLAST version 2.13.0 was used to get the best gene match using blastx to the predicted proteins of *C. virginica*. Of the 1179 selection regions, 614 regions had no transcript overlapping the region. For the remaining 565 regions, there were 574 transcripts overlapping one or more regions. From these transcripts, 271 had a hit to a *C. virginica* protein and were used for functional enrichment analysis.

<u>Rough estimates of the ages of live-bearing alleles</u>

*Associated code:*
Window-based summary statistics
https://github.com/seanstankowski/Littorina_reproductive_mode/tree/main/dating_sweeps

We used theory outlined in Tavares et al. (*64*) to roughly estimate the age of sweeps at live bearing alleles. During a sweep, diversity is eliminated close to the selected site. In the case of a hard sweep, all ancestral diversity is eliminated from the swept region, but new mutations in the swept segment at rate $\mu$ will generate $\pi_w = 2\mu T$ at $T$ generations after the sweep. Thus, the time of the onset of the sweep can be estimated as $T = \pi_w/2\mu$. Diversity arising in other ways complicates the estimation, causing $T$ to be overestimated. For example, some ancestral variation may be retained if the beneficial allele was segregating on multiple haplotypes before the sweep (soft sweep). This means that $\pi_w$ would reflect mutations that arose both before and after the sweep. Similarly, gene flow and recombination between populations may introduce additional variation into a swept region that did not arise though new mutation. Finally, it is difficult to define precise sweep boundaries, and error in doing so would tend to inflate $\pi_w$.

To correct for these confounding sources of diversity, we estimated the age of each sweep (*i.e.*, the time in generations that each sweep began) by calculating $\pi_w$ from private alleles. This assumes that alleles private to live-bearers arose in the swept segment after the sweep, and that alleles shared between egg-layers and live-bearers reflect some confounding factor. However, this approach would over-estimate the age of a soft sweep if the private alleles were already private to haplotypes carrying the beneficial allele when the sweep began.

For each of the 50 regions associated with reproductive mode (fig. S28; table S8), we identified and then removed SNPs that were polymorphic in both egg-layers and live-bearers (see section 'Analysis of private alleles', p. 23). We then calculated $\pi_w$ for live-bearers as described above (see section 'Estimating diversity, divergence and differentiation in 100 SNP windows', p. 21). We corrected the denominator in the $\pi_w$ calculation for the removal of the shared sites.

For calculating $T$, we assumed a mutation rate of $\mu = 1.5\times10^{-8}$. We converted time in generations into years using two estimates of generation time for *Littorina saxatilis*: 2 generations per year and 1 generation per year (*65*, *66*). Estimates of time in generations varied roughly ten-fold, ranging from 23,266 to 227,094 generations before present, with a median of 69,790 generations before present (Fig. S38). In years, this equates to a minimum of 11,633 – 23,266 years before present, and maximum of 113,547 – 227,094 years before present, with a median of 34,895 – 69,790 years before present. The distribution of ages is strongly negatively skewed, with the majority of sweeps being initiated in the last 100,000 generations.

### Supplementary text

Detailed results from the ternary analysis of simulated topology weights

In this section, we interpret the results of ternary analysis of simulated topology weights, focusing on the effects that different parameters have on the extent of lineage sorting (ILS) and asymmetry of the distribution. Each scenario is first discussed separately, and some general conclusions are drawn at the end.

*Scenario a: Uniform $N_e$ and split times:* This scenario gives a clear picture of how variation in the time between population splits shapes the distribution of topology weights (fig. S10; table S4). With a very small number of generations between splits, most loci are distributed very near to the center of the ternary plot, indicating that we see roughly equal proportions of the three possible topologies from a large sample of subtrees. This is expected, because when the durations between splits are much less than the average time to coalescence (*i.e.*, $\ll 2N_e$ generations; here $2N_e = 1000$ generations), most gene copies will coalesce in the common ancestor to all of the populations, rather than in the lineage that is ancestral to each spilt (*14*, *15*). In other words, ILS (aka ‘deep coalescence’; (*14*)) is extensive and dominates the relationships among the individuals. As the time between splits increases, we begin to see a bias toward the top of the ternary distribution, indicating a tendency of gene trees to show a greater resemblance to species tree rather than the two alternative trees. However, we only see a strong shift toward the top of the plot as the time between splits approaches $2N_e$ generations. Only long after the average time to coalescence (*i.e.*, $\gg 2N_e$ generations) do we see perfect concordance between all gene trees and the species tree. This is also expected because the full distribution of coalescence times is expected to be very broad (*e.g.* the standard deviation of times is also $2N_e$ generations). Estimates of $D_{LR}$ for these simulations are all low, and consistent with values expected by chance ($p > 0.05$). Thus, there is no effect of time between splits on the symmetry of the distribution (table S4).

*Scenario b: Varying but equal split times for T2 and T3, with T1 set to 5k generations:* This scenario shows how the effects of the split time play out over longer evolutionary time frames (fig. S11; table S4). The key result here is that the effects of ILS cannot be resolved even if a long period of evolution occurs after the splits. Rather, additional time only reinforces the patterns of discordance, resulting in trees that perfectly fit one of the three alternative topologies. However, the fraction of each topology observed is a function of time between splits (see the pie charts in fig. S11). For example, when the times are short, (*i.e.*, when ILS is extensive) all three trees are observed at similar frequencies. However, as the time increases, the fraction of the two alternative trees decreases, such that an increasing fraction of the loci show the background topology. Estimates of $D_{LR}$ are not significantly different from 0 in any simulation, showing that a period of subsequent evolution has no effect on the symmetry of the distribution (table S4).

*Scenario c: Uneven split times:* This scenario shows the unevenness of split times shapes the distribution of weights. We can see, in both the ‘short’ and ‘long’ split time scenarios, that the evenness of split times has a small but pronounced effect on the concordance between gene-trees and the background demography (fig. S12; table S4). Specifically, lineage sorting is more complete when $T_2$ and $T_3$ occur deeper back in time. Thus, ILS has a more pronounced impact when splits occur closer to the present. Estimates of $D_{LR}$ show that the evenness of splits has no effect on the symmetry of the distribution (table S4).

*Scenario d: Uniform variation in $N_e$:* We know from theory that decreasing $N_e$ has the same effect on lineage sorting as increasing the time between splits (*15*). In other, words the probability of lineage sorting increases with decreasing $N_e$. This is very clear from this scenario, where simulations were conducted with the same split times, but different values of $N_e$ (fig. S13; table S4). At large values of $N_e$, the distribution of weights is centered in the middle of the ternary plot, but as the $N_e$ is decreased, the distribution shifts toward the apex of the triangle. As with variation in the split times, uniform variation in $N_e$ does not impact the symmetry of the distribution of weights ($D_{LR}$ not significantly different from 0; table S4).

*Scenario e: Varying $N_e$ in one population:* In this scenario, we alter the $N_e$ only in P3, making it either smaller or larger than the background $N_e$, which was fixed for all other lineages ($N_e$ = 500) (fig. S14; table S4). This has the effect of increasing or decreasing the probability that gene copies in that population will coalesce before the common ancestor to the ancestral population P123. When the $N_e$ for P3 is larger than the background (*i.e.*, $N_e$ = 5000), we see a stronger effect of ILS. However, when the $N_e$ is smaller than the background, we see little change in the extent of lineage sorting. Rather we tend to see an increase in variance of the topology weights resulting in more extreme values. This makes intuitive sense, because if all gene copies in P3 coalesce before $T_3$, then all individuals will form a monophyletic group, rather than being randomly scattered across a tree. This will ultimately cause a given gene tree to look more like one of the alternative subtrees, but the overall level of discordance is controlled by the rate of ILS experienced by the other groups. Changing the $N_e$ of one population has no effect on the symmetry of the distribution ($D_{LR}$ not significantly different from 0; table S4).

*Scenarios f & g: Uni- and bi-directional migration between P2 and P3:* In these scenarios, we simulated geneflow between P2 and P3 to see how it influences the distribution of weights. In the standard four population model, we generally expect gene copies in P1 and P2 to coalescence first. However, gene flow between non-sister taxa leads to the sharing of haplotypes between them, leading to shallower coalescence between P2 and P3. This effect is very clear in the simulations, as gene flow between P2 and P3 shifts the distribution of weights from the species tree toward the relevant alternative topology where these taxa cluster together (figs. S15 & S16; table S4). This effect is strongest in simulations where lineage sorting is more complete before the onset of gene flow, because, without a dominant topology to begin with, there is no bias in the distribution that gene flow can shift. This is most obvious when the split times are very short, and gene flow between P2 and P3 only leads to the exchange of variation that is already shared among all four groups. As expected, the estimates of $D_{LR}$ are substantial and highly significant for most simulations, reflecting the strong asymmetry between the left and right sides of the plots (table S4). In general, the effect of gene flow becomes clearer as the migration rate increases, but only up to a point that is determined by the extent of lineage sorting before the onset of migration. Similarly, gene flow in both directions (*i.e.*, P3 to P2 and P2 to P3) results in more pronounced symmetry at lower migration rates, as there is twice as much migration between the groups.

*Scenario h: Unidirectional migration for 10% of the genome:* In this scenario, we combine two simulations—one with gene flow and one without—in order to simulate heterogenous gene flow across the genome, as is expected when species boundaries are porous (fig. S17; table S4). The effects of gene flow on the 10% of loci (red points) is clear in all but one of the simulations, where the time between splits is smallest and where the rate of migration is lowest. Even in the context of the symmetrical background distribution consisting of 90% of loci, the asymmetry in the simulations is detectable with the $D_{LR}$ statistic which was highly significant (table S4). This highlights the potential utility of the ternary framework when it comes to detecting gene flow and introgression in genome-wide datasets.

Although neutral gene flow can generate strong asymmetry in some of these simulations, the pattern of asymmetry observed in our empirical data is not recovered in neutral simulations. First, windows that are a perfect fit for the alternative topology never became more abundant than those that perfectly fit the background tree (Table S14). In these simulations, we always observed a ratio of Tr:Tb much less than 1, with a maximum of 0.37 at an extreme migration rate of 50% between P3 and P2. In our observed data, we found a positive ratio, of 1.41, indicating that windows with Tr = 1 are more abundant than those with Tb = 1. Observing this result in a neutral model would require substantial gene flow (> 10%) for more than half of the genome. However, models with migration also fail to produce the otherwise flat-tailed distribution of topology weights for the alternative topology, as seen in Fig. 2E (for Tr and Tc). Rather, the distributions for the alternative tree Tr with migration always had a similar shape to that observed for the background tree (Fig 2E, as for Tb). Thus, we conclude that realistic levels of neutral gene flow for a small fraction of the genome cannot produce the patterns of asymmetry that we observe in our empirical data (i.e., a strong excess of the alternative tree compared with the background tree, but with an otherwise flat-tailed distribution of weights). We think that selection on a small fraction of shared genome is needed to create this pattern.

*Scenario i: Ancestral structure*: This scenario, which follows that used Martin et al. (*44*), is designed to model a region of the genome undergoing balancing selection or some other process that maintains polymorphism at particular loci in the ancestral lineage P123 (Fig. S18). In this scenario, 10% of loci evolve under a different history, which results in a bias toward the corresponding alternative topology. This is reflected in the non-zero estimates of $D_{LR}$ in all of the simulations.

*General conclusions*: Based on the results of these simulations, we can make some general conclusions regarding how different factors shape the ternary distribution of topology weights. First, more time between splits and smaller effective population sizes ($N_e$) increase the probability of lineage sorting. This shifts the distribution of weights toward the top corner of the ternary plot, which corresponds to the topology that matches the background demography. No combination of split times or effective sizes causes an asymmetrical distribution of topology weights between the left and right sides of the ternary plot. Gene flow and ancestral structure can both produce distributions of weights that are asymmetrical. The ability to detect gene flow with a genome-wide estimate of $D_{LR}$ will depend on the number of loci experiencing gene flow, the rate of migration (including whether it is uni- or -bidirectional), and the extent of lineage sorting before gene flow commences. Finally, the pattern of asymmetry that we observed toward Tr is not consistent with what we found in neutral simulations.

<u>Inversions explain the genome-wide bias toward Tc</u>

In the symmetry analysis in main text Fig. 2D, we show that there is a small but significant genome-wide bias toward the control topology (left side of ternary plot), which is caused by loci that show a slightly better fit toward Tc than the other two topologies.

We found that this effect could be explained by several large chromosomal inversions that segregate among these species of *Littorina* (23) (fig. S21). For each window, we used a genetic map to first determine whether it was located within a known chromosomal inversion (15,992), in a colinear region of genome (60,441), or in an area of uncertainty near an inversion breakpoint (5,197). We dropped windows near breakpoints to ensure that we only analyzed windows inside and outside of known inversions.

To reveal the general effect of inversions, we removed all associated windows and estimated $D_{LR}$ for the colinear regions only. This test revealed a very small estimate of $D_{LR}$ = -0.0072 (*i.e.*, indicating 0.72% more windows on the right side of the plot than expected) that was not significantly different from 0 (p = 0.074) (fig. S21). Thus, colinear regions of the genome fit the expectations for an idealized four population model (*i.e.*, neutral sorting with no gene flow between lineages) in that the two alternative topologies are present in roughly equal frequencies. In contrast the $D_{LR}$ for windows inside inversions was significantly positive ($D_{LR}$ = 0.23), indicating that there 23% more windows on the left side of the plot than expected (p = 6.67e-134).

To further understand the asymmetry, we dissected the effects of the inversions by examining each one separately. We found a broad range of $D_{LR}$ estimates ranging from not significantly different from 0 for LGC1.1 and LGC14.3 ($p > 0.05$), to greater than 0.8 for LGC2.1 (fig. S22; table S6). However, there were also some general patterns. First, of the 12 inversions that show significant asymmetry, only one (LGC17.1) showed a negative $D_{LR}$ (-0.218), indicating a slight bias toward Tr. Also, the biases toward Tc were quite substantial for several inversions. For example, 5 inversions showed an estimate > 0.4. Thus, it is clear that multiple chromosomal inversions contribute to the Tc bias.

Ternary distributions for the three inversions with the largest $D_{LR}$ values clearly show the left-hand bias of weights (fig. S23). They also show that the windows fall roughly in the area where we observe the Tc bias in the analysis of the full dataset.

Focusing on the inversion with the largest estimate of $D_{LR}$, we used PCA to determine the karyotype for each individual and then estimated the frequency of the two orientations in each clade. The PCA analysis revealed three fairly clear groups—left and right groups coinciding with the alternative homokaryotypes, and a central group of heterokaryotyes—as expected for a chromosomal inversion (*53*) (fig. S24). The resulting arrangement frequencies for each clade are as we would expect from the results of topology weighting. Contrary to what we would expect from the background phylogeny, the A arrangement is more common in *L. compressa* and Northern *L. saxatilis*, whereas the R arrangement is more common in *L. arcana* and Spanish *L. saxatilis* (fig. S24).

### Supplementary Figures and Tables

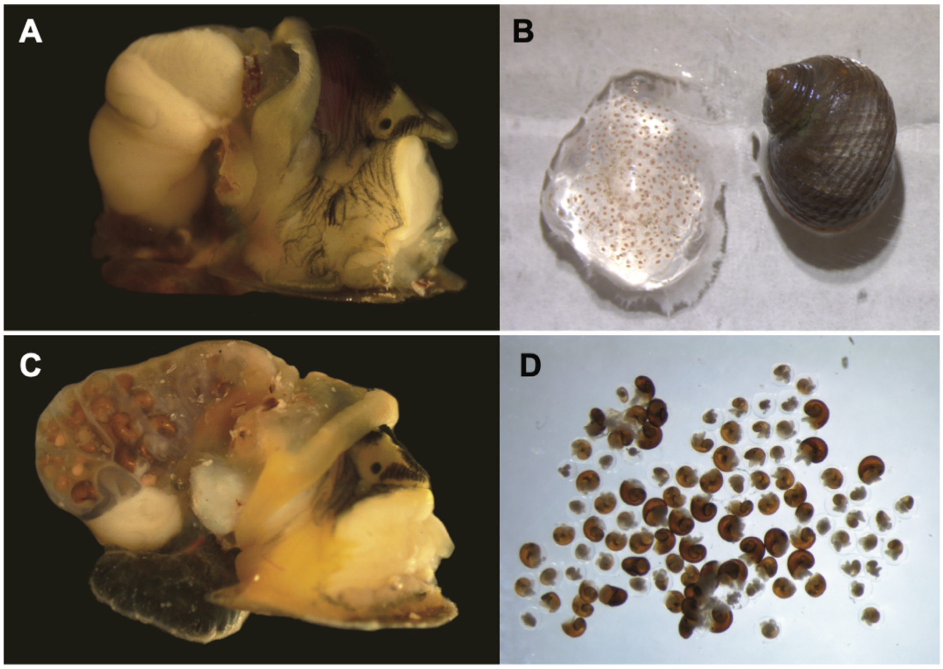

**Figure S1. Anatomical differences between egg-laying and brooding *Littorina*.** A) Photograph of a dissected *L. arcana* showing the jelly gland. B) An egg-mass laid by an individual of *Littorina arcana* in the lab at the University of Sheffield. C) A dissected *L. saxatilis* showing the brood pouch with embryos developing inside. D) The contents of a dissected brood pouch show the continuum of development that occurs within the mother, from embryos through to crawl-away offspring.

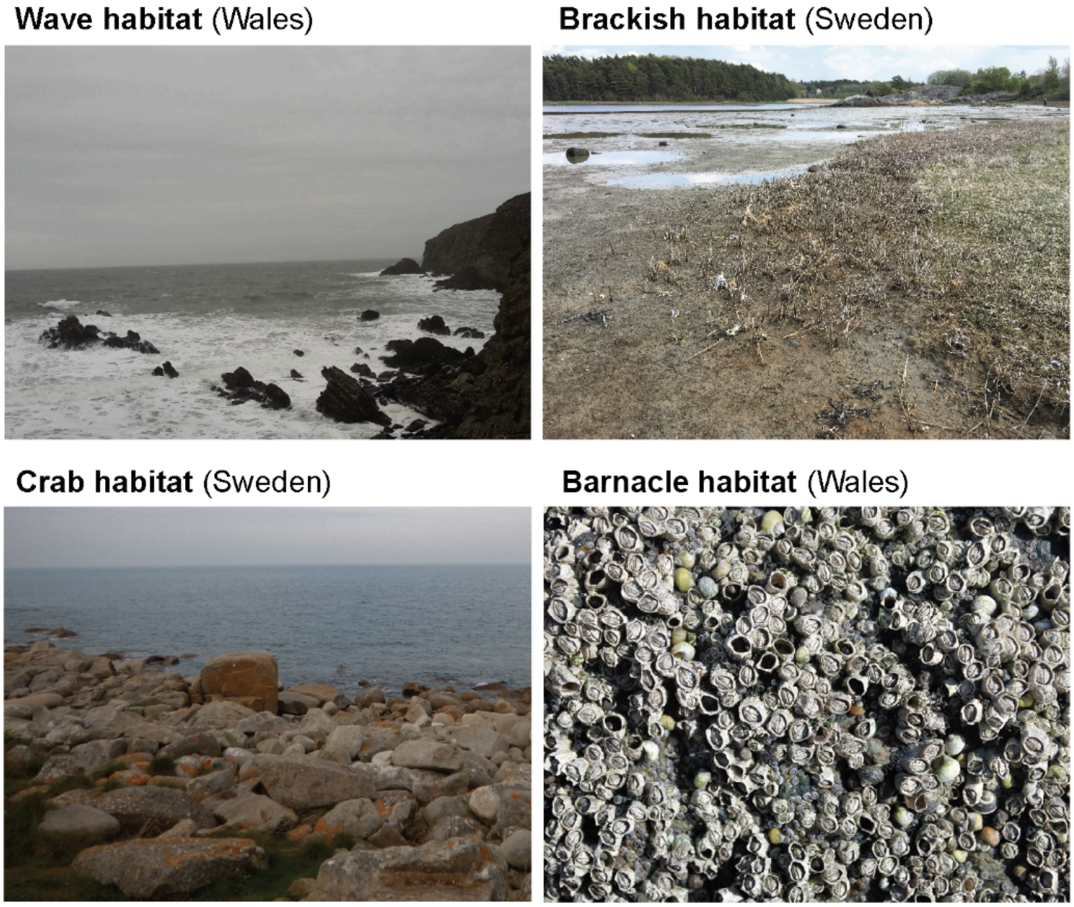

**Figure S2. Characteristic habitats associated with *Littorina* species and ecotypes.** Wave habitat near Holyhead, Wales. Brackish habitat near Tjärnö, Sweden. Crab habitat, near Tjärnö, Sweden. Barnacle habitat near Holyhead, Wales.

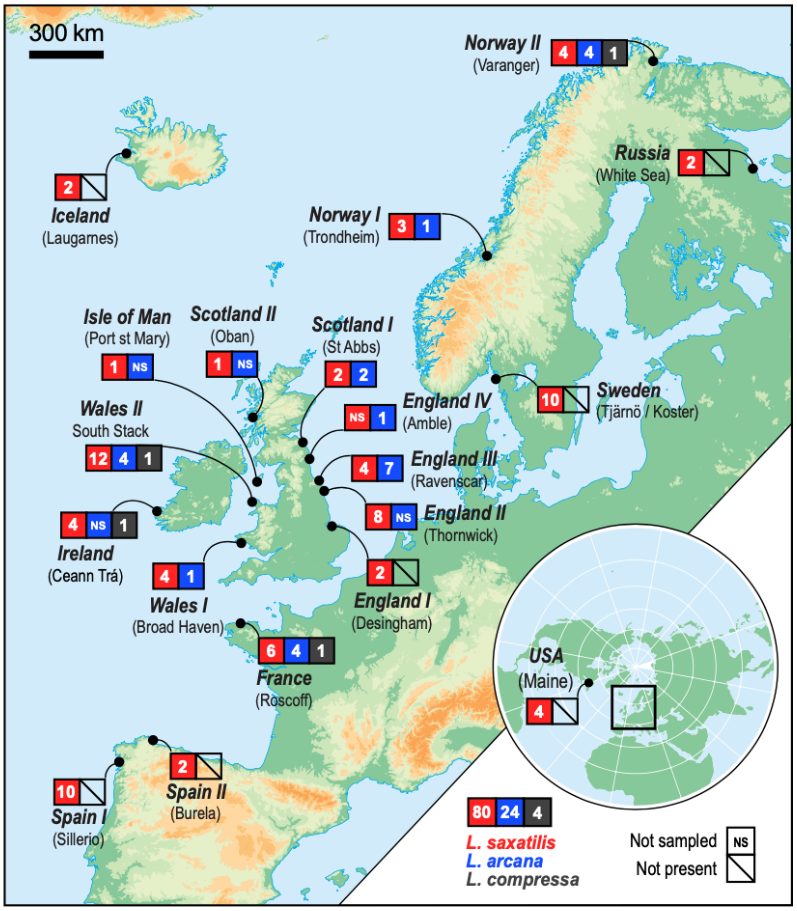

**Figure S3. Map of sampling locations with the number of each species sampled.** The number of each species sampled is given in the colored boxes (red = *L. saxatilis*, blue = *L. arcana* and gray = *L. compressa*). N.S. indicates that the species was present at the sample location but was not sampled in this study. Boxes with the strike-though indicate that *L. arcana* and *L. compressa* are not present. The locations correspond with those listed in Table S1.

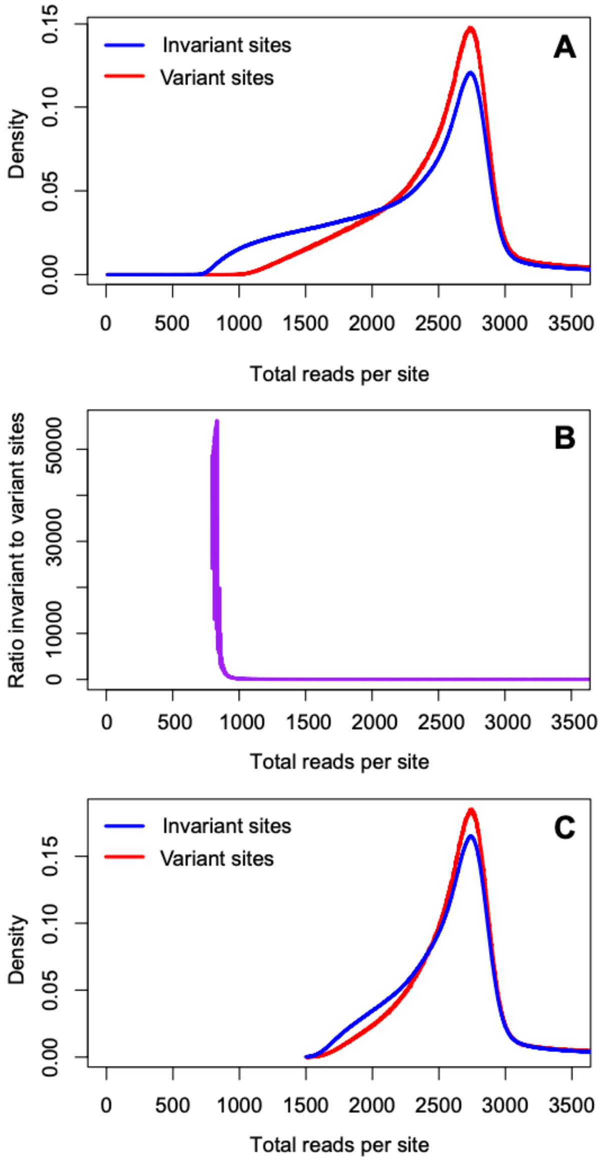

**Figure S4. Distribution of depth at different stages of filtering.** The blue and red lines in plots A and C show the count of sites in the dataset that have *n* total reads across all 108 individuals for invariant (blue line) and variant sites (red line). (A) The distributions after the initial filtering, which included the removal of genotypes with a depth lower than 5 and quality score less than 20. (B) The relationship between the ratio of variant to invariant sites and the total coverage per site. Note that this ratio is very high at low total coverage, because sites with fewer total reads are more likely to be called as invariant; The ratio of invariant to variant sites stabilizes around a total coverage of 1500. (C) The final distribution of the count of sites in the dataset that have a count of *n* total reads across all 108 individuals after removing all sites with a total depth less than 1500 reads over all 108 individuals and stricter soft filtering (table S2).

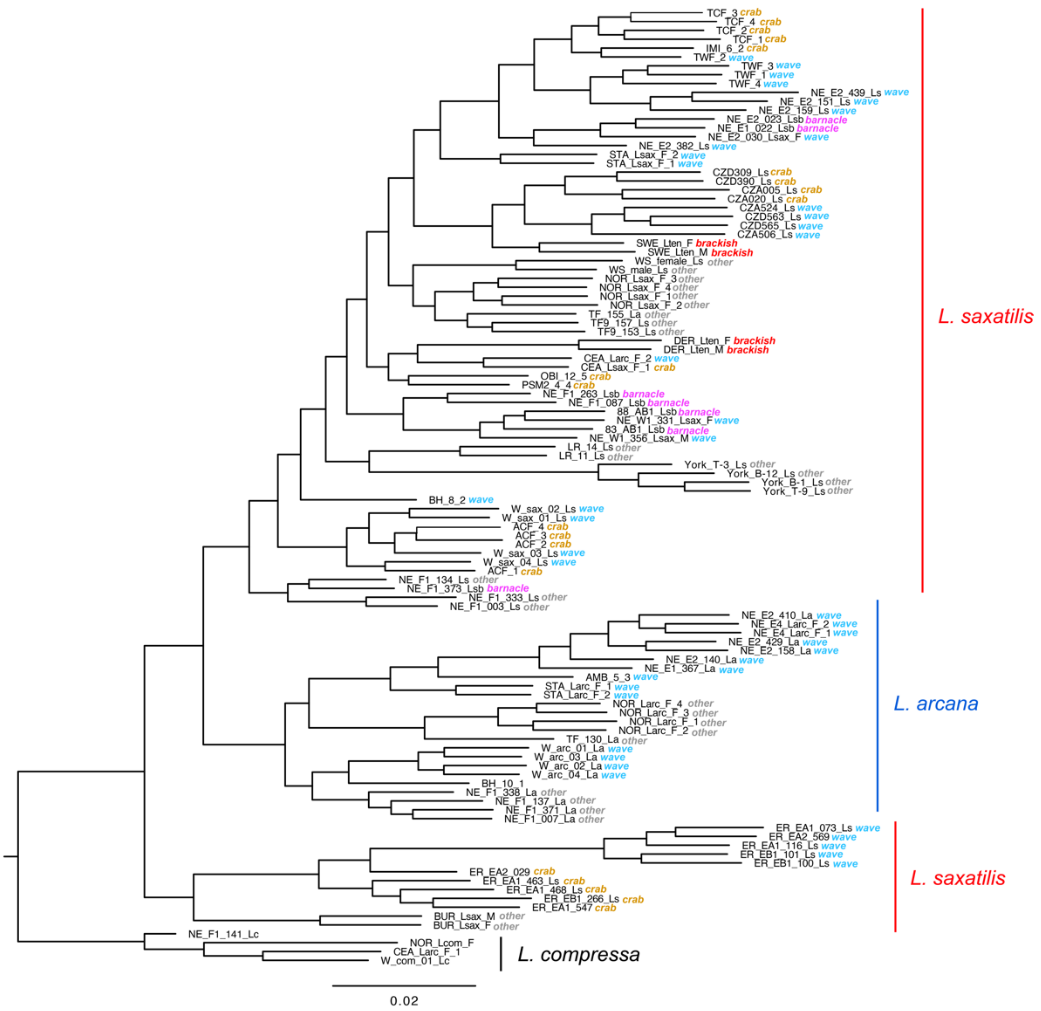

**Figure S5. ML phylogeny inferred from genome-wide data.** This tree is the same as the one in main text Fig. 1E, except that it includes tip labels and information about the ecotype of each sample. Sample IDs for each tip correspond to those in table S1. Colored labels indicate whether the ecotype of a sample was crab, wave, brackish, barnacle or other.

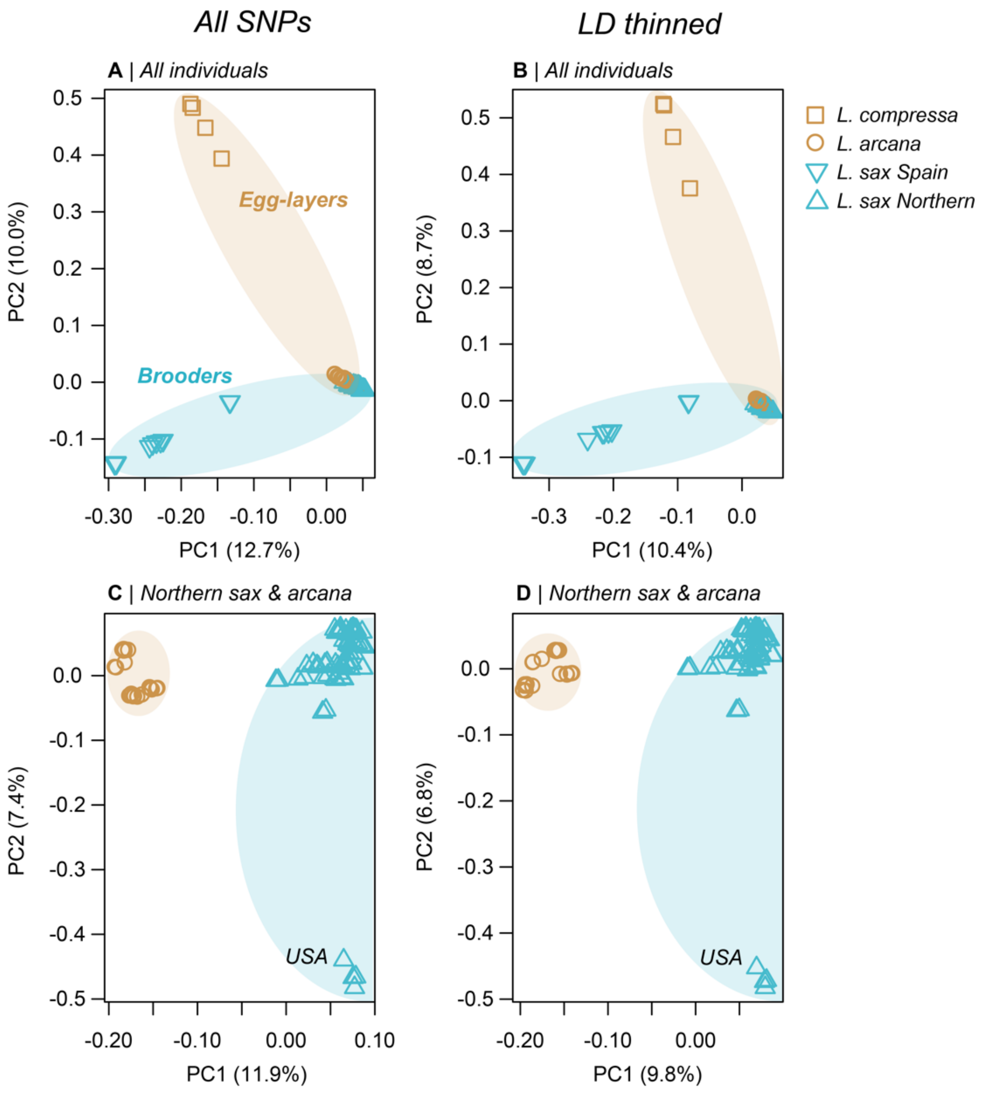

**Figure S6. The results of four separate principal components analyses (PCs 1 & 2)**. A) PCA conducted on all SNPs and all individuals. B) All individuals but with the SNP data pruned to reduce LD among linked sites. C) All SNPs, but only individuals of Northern *L. saxatilis* and *L. arcana*. D) With the LD thinned data and the reduced set of individuals. In the PCA with all individuals, the first 2 PCs recover the discordance between genome-wide relationships and reproductive mode that was observed in the ML phylogeny. The analyses of the full and LD pruned datasets produced very similar results. In the second set of PCAs, which contained only Northern *L. saxatilis* and *L. arcana*, PC1 clearly separates individuals by species, as was also observed in the phylogenetic analysis. PC2 separated North American samples of Northern *L. saxatilis* from the rest of the group. Overall, the PCA analyses provide further support for our main conclusion regarding the discordance between genome-wide variation and reproductive mode. Specifically, egg-laying and live-bearing samples are often more similar to one another than they are to samples that share the same mode of reproduction.

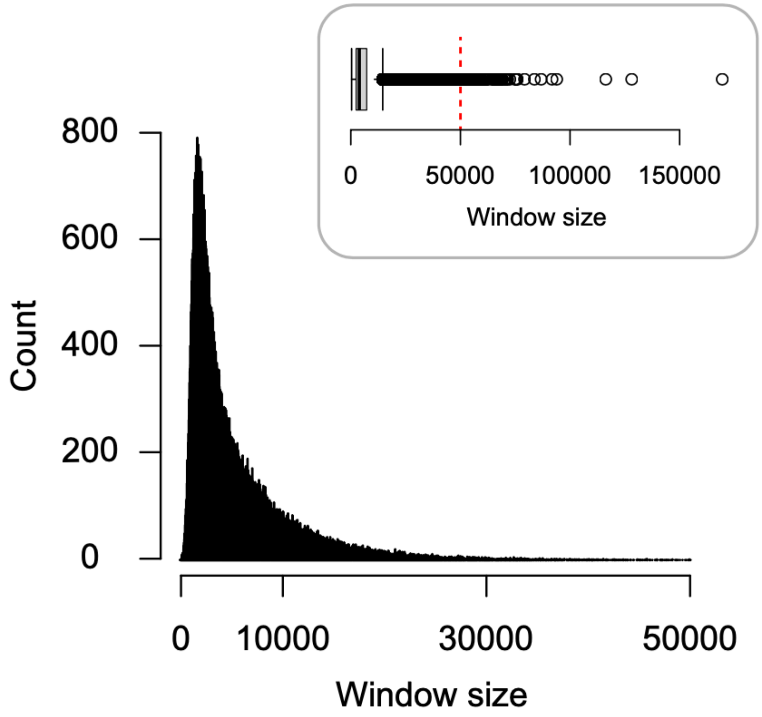

**Figure S7. Distribution of physical sizes (kbp) of the 100 SNP windows used for local tree construction and topology weighting.** The mean size is 5.8 kb ± s.d. 5.3 kb. The red dashed line in the top plot shows the margin of the main plot.

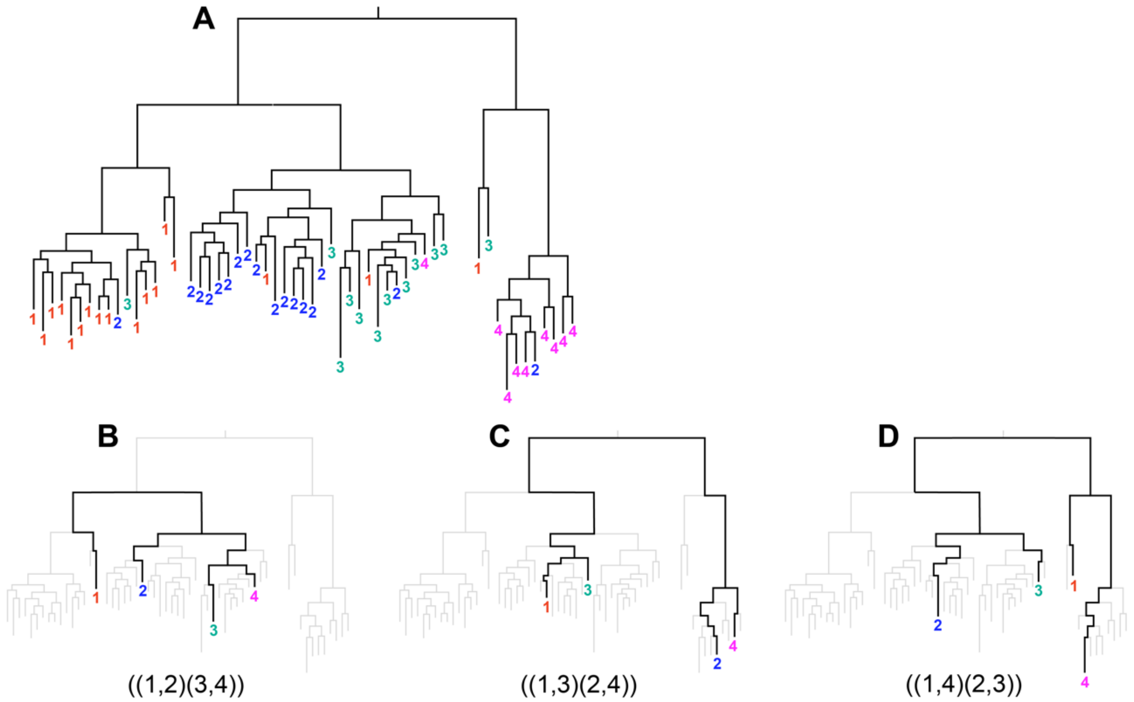

**Figure S8. Alternative subtrees sampled from a large genealogy.** (A) A hypothetical genealogy for a single non-recombining locus sampled from four species (1,2,3,4). The species do not form monophyletic clades, meaning that relationships between the species vary depending on the samples being considered. (B - D) Three example taxon subtrees sampled from the full tree showing the three possible unrooted topologies that can be observed in a tree with four taxa.

*a. Uniform $N_e$ and split times*

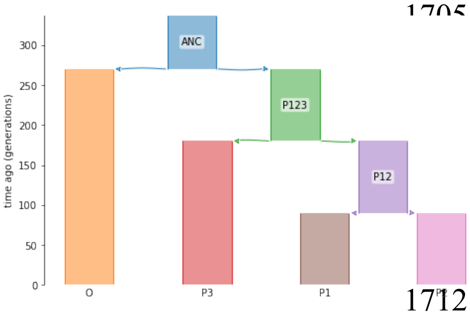

*b. Varying but equal split times for T2 and T3, with T1 set to 5k generations*

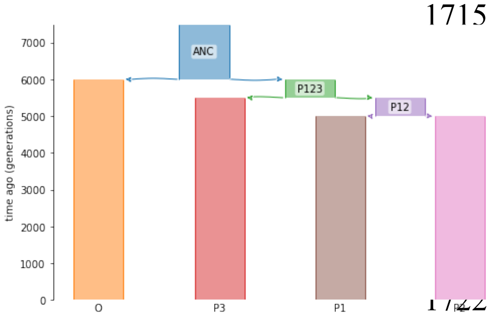

*c. Uneven split times*

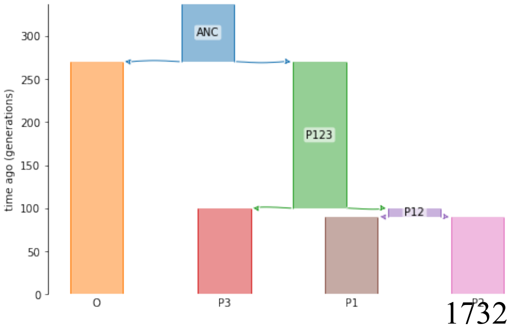

*e. Varying $N_e$ in one population*

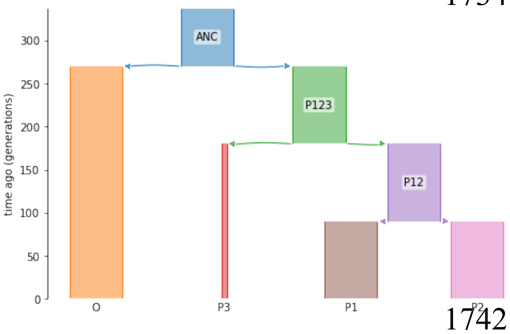

**Figure S9. Cartoons of the models used in the simulations**. Letters correspond with the scenarios described in the main text. Scenario d is not shown, but looks identical to scenario a. The figure continues onto the next page.

*f & g. Uni- and bi-directional migration between P2 and P3*

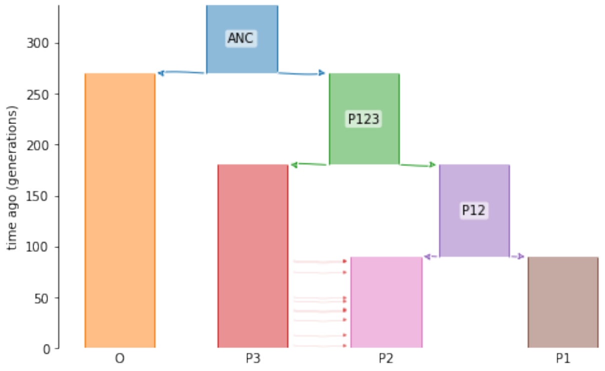

*h. Unidirectional migration for 10% of the genome*

*90% of loci* *10% of loci*

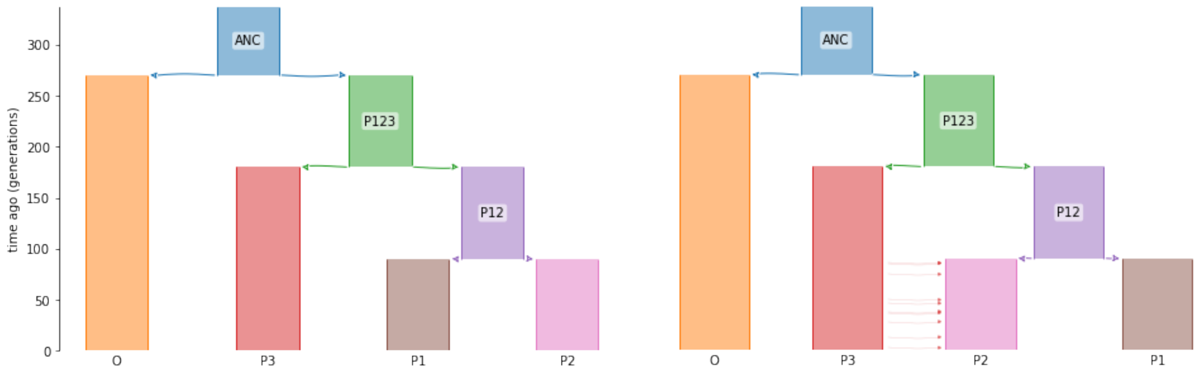

*i. Ancestral structure*

*90% of loci* *10% of loci*

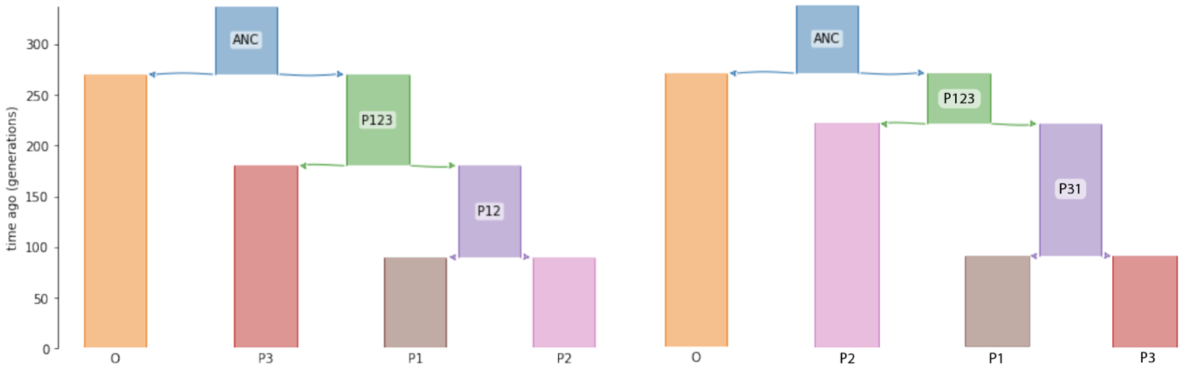

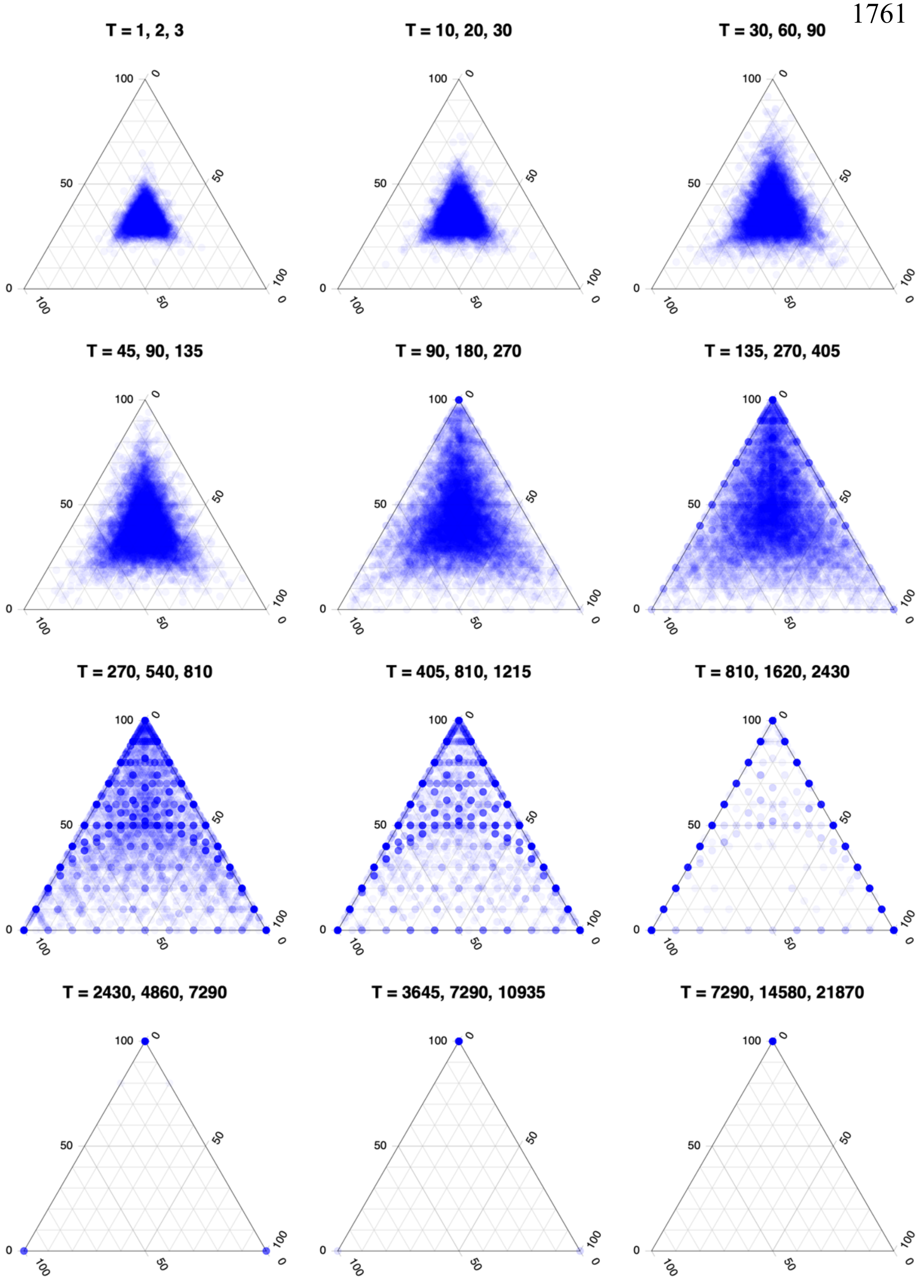

**Figure S10. Joint distribution of topology weights for simulations with different but even split times.** Each point is one of 10000 simulated loci. The time of the three splits ($T_1$, $T_2$, $T_3$) used for each simulation is given above each plot. All lineages have an $N_e$ of 500, with no migration between them.

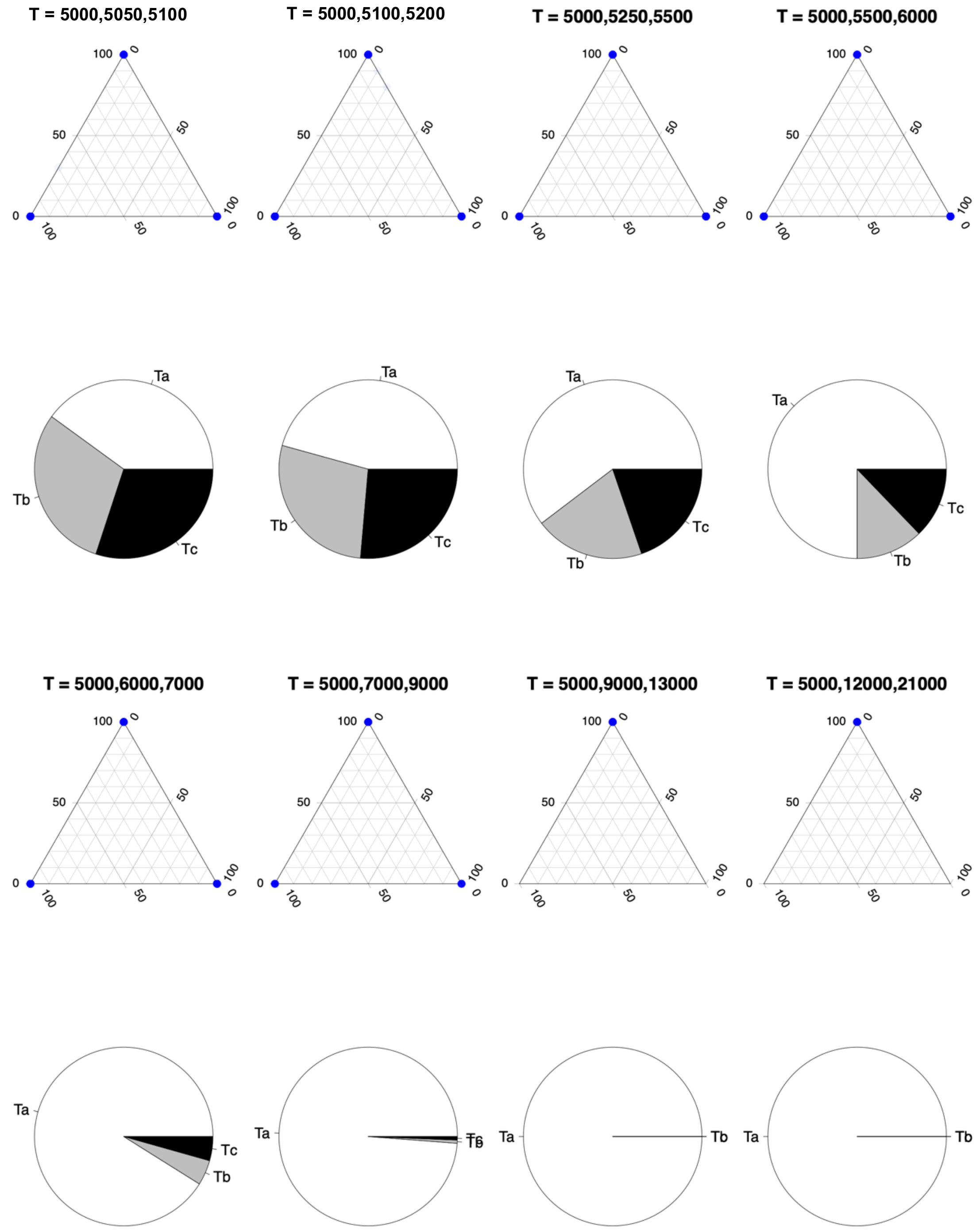

**Figure S11. Joint distribution of topology weights for simulations with different but even split times, followed by 5000 generations after the final split.** The time of the three splits ($T_1$, $T_2$, $T_3$) used for each simulation is given above each plot. All lineages have an $N_e$ of 500, with no migration between them. The pie charts under each triangle show the fraction of trees with a topology that perfectly match one of the three subtrees Ta (apex of the triangle), Tb (left corner of the triangle), or Tc (right corner) for that simulation.

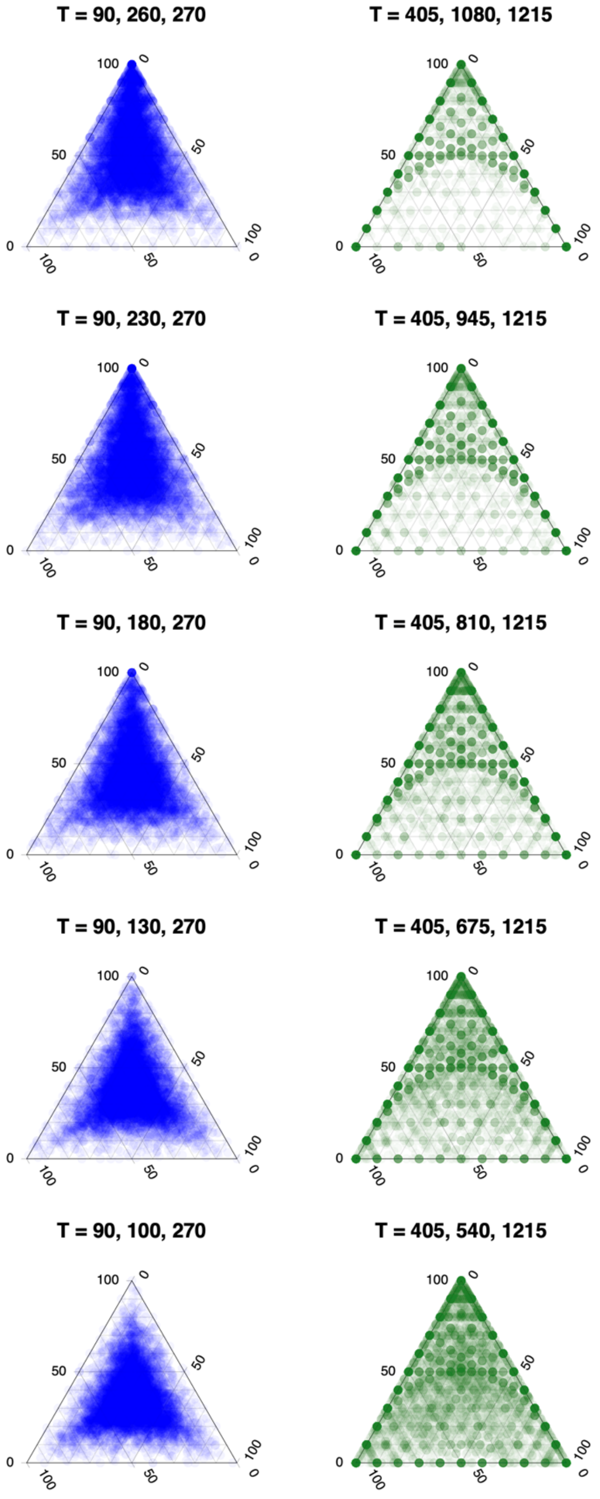

**Figure S12. Joint distribution of topology weights for simulations with uneven split times.** Each point is one of 10000 simulated loci. The columns show simulations with the same $T_1$ and $T_3$ splits, but with the $T_2$ varying between them. The split times used for all lineages are given above each plot. All simulations have an $N_e$ of 500, with no migration between lineages.

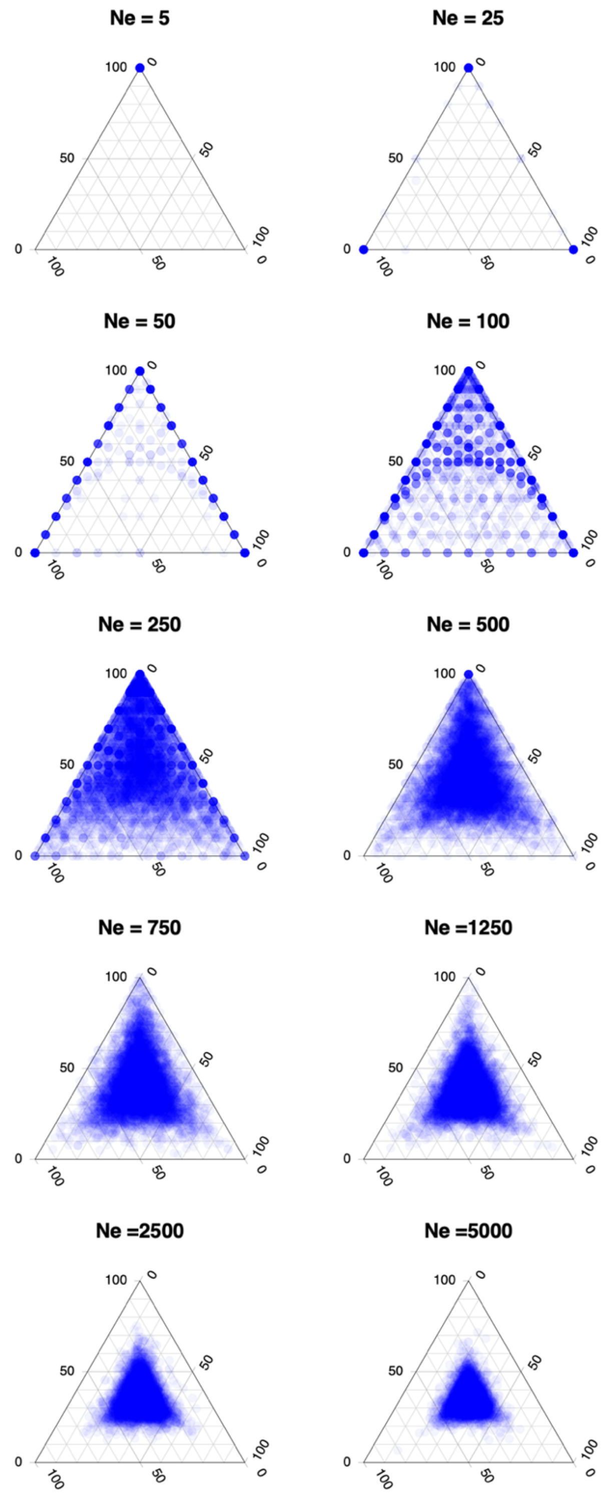

**Figure S13. Joint distribution of topology weights for simulations with different effective populations sizes.** Each point is one of 10000 simulated loci. The $N_e$ used for all lineages is given above each plot. All simulations have the slit times $T_1 = 90$, $T_2 = 180$, $T_3 = 270$, with no migration between lineages.

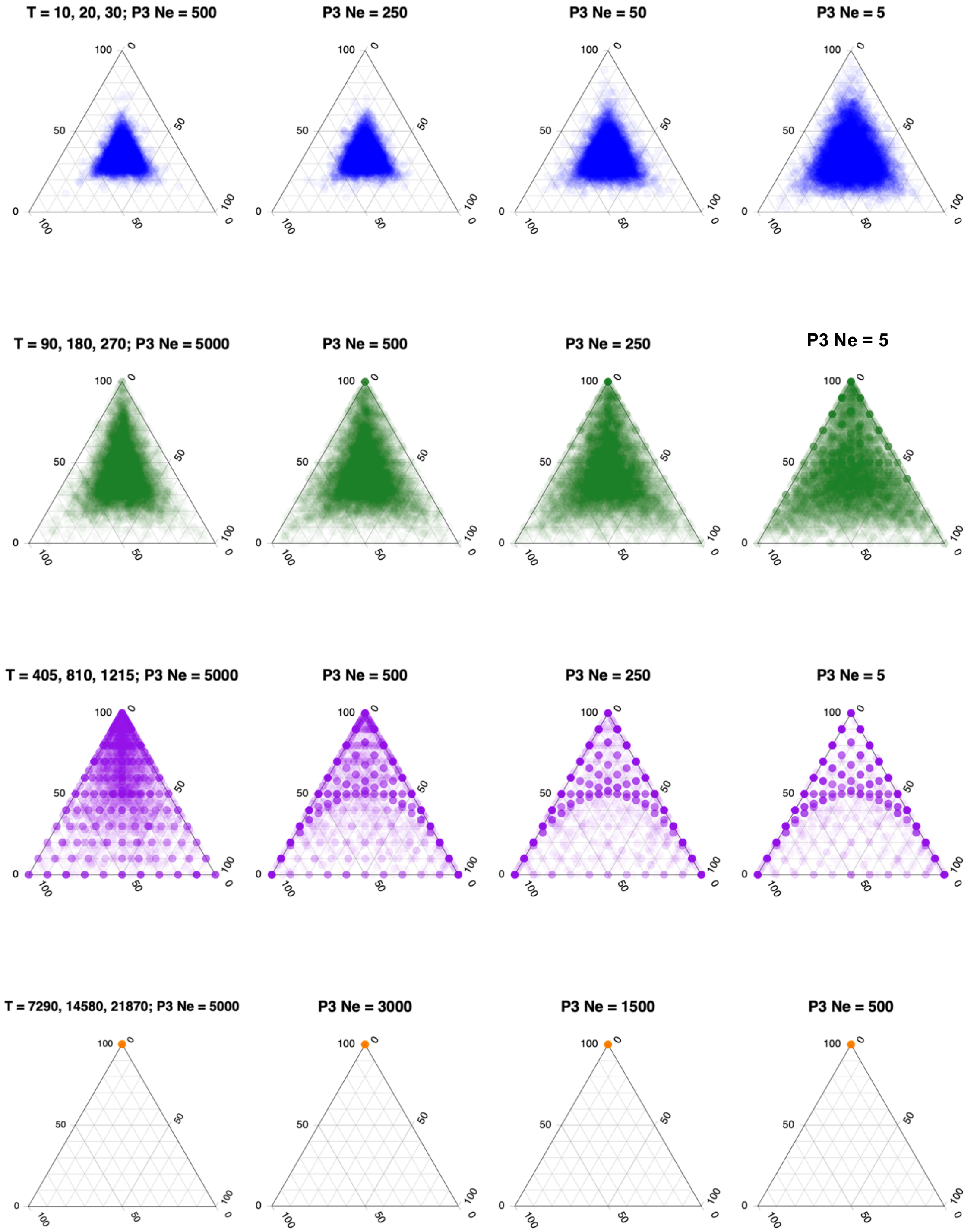

**Figure S14. Joint distribution of topology weights for simulations where population P3 has a different effective population size than O, P1, P2.** Each row shows four simulations with the same split times (T). The $N_e$ used for lineage P3 is given above each plot. The $N_e$ for all other populations is 500. The split times used for all lineages are given above each plot. No migration was allowed between lineages. Each point is one of 10000 simulated loci.

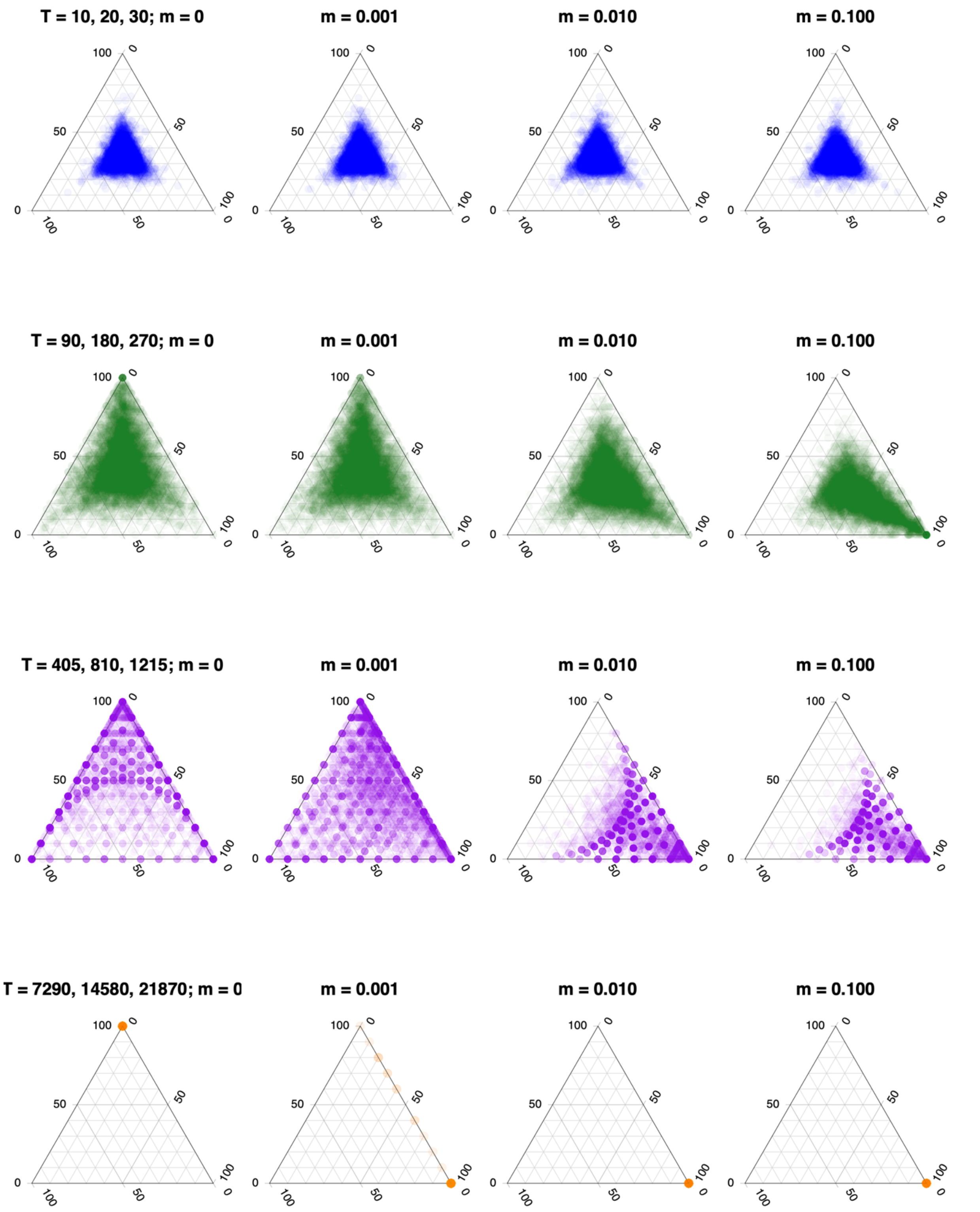

**Figure S15. Joint distribution of topology weights for simulations with different rates of unidirectional migration.** Each row shows four simulations with the same split times (T), but with different rates of migration from P2 to P3, as indicated above each plot (*m* = 0, 0.001, 0.010 or 0.100). All lineages have an $N_e$ of 500. Each point is one of 10000 simulated loci.

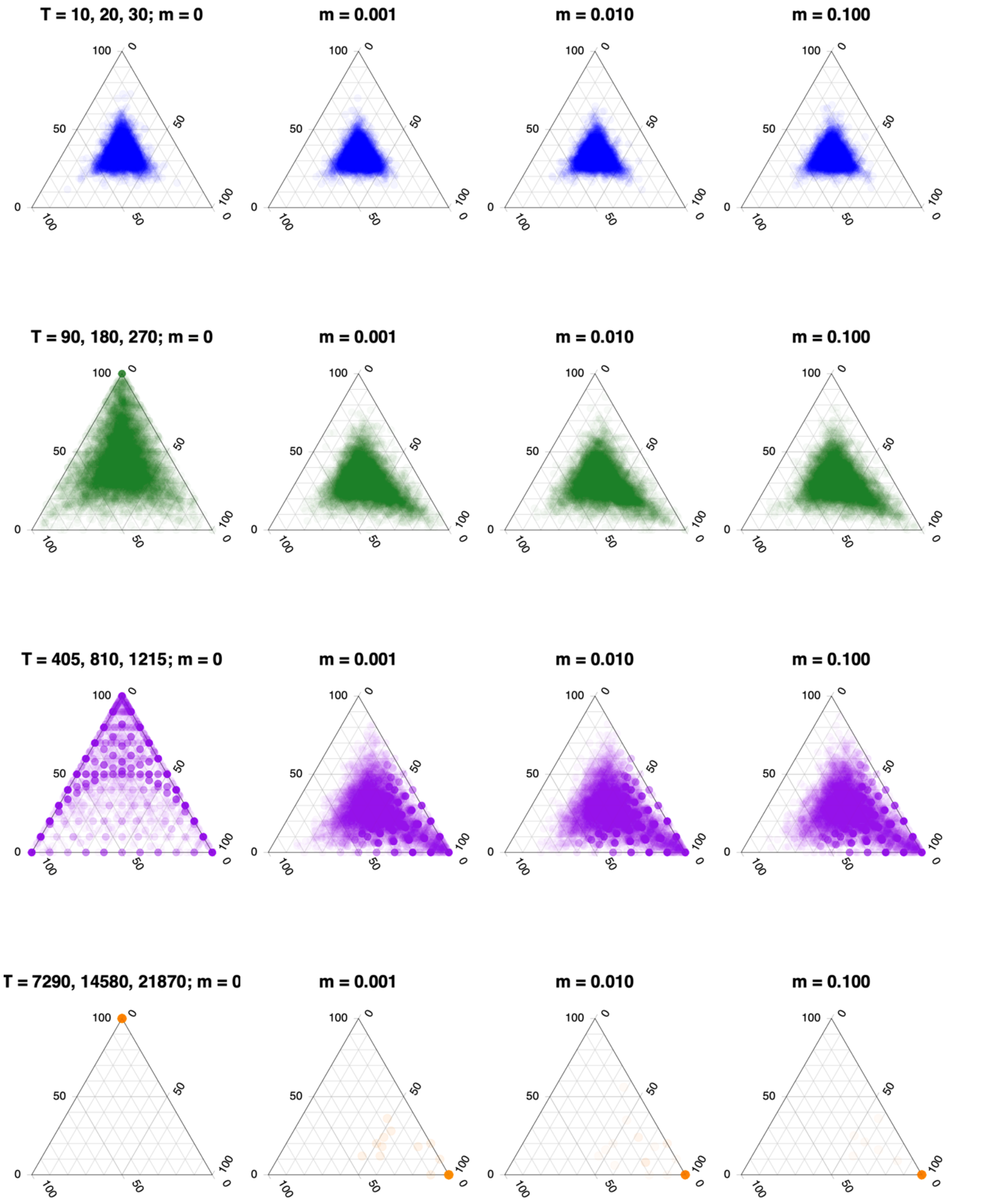

**Figure S16. Joint distribution of topology weights for simulations with different rates of bidirectional migration.** Each row shows four simulations with the same split times (T), but with different rates of migration from P2 to P3 and P3 to P2, as indicated above each plot ($m = 0$, 0.001, 0.010 or 0.100). All lineages have an $N_e$ of 500. Each point is one of 10000 simulated loci.

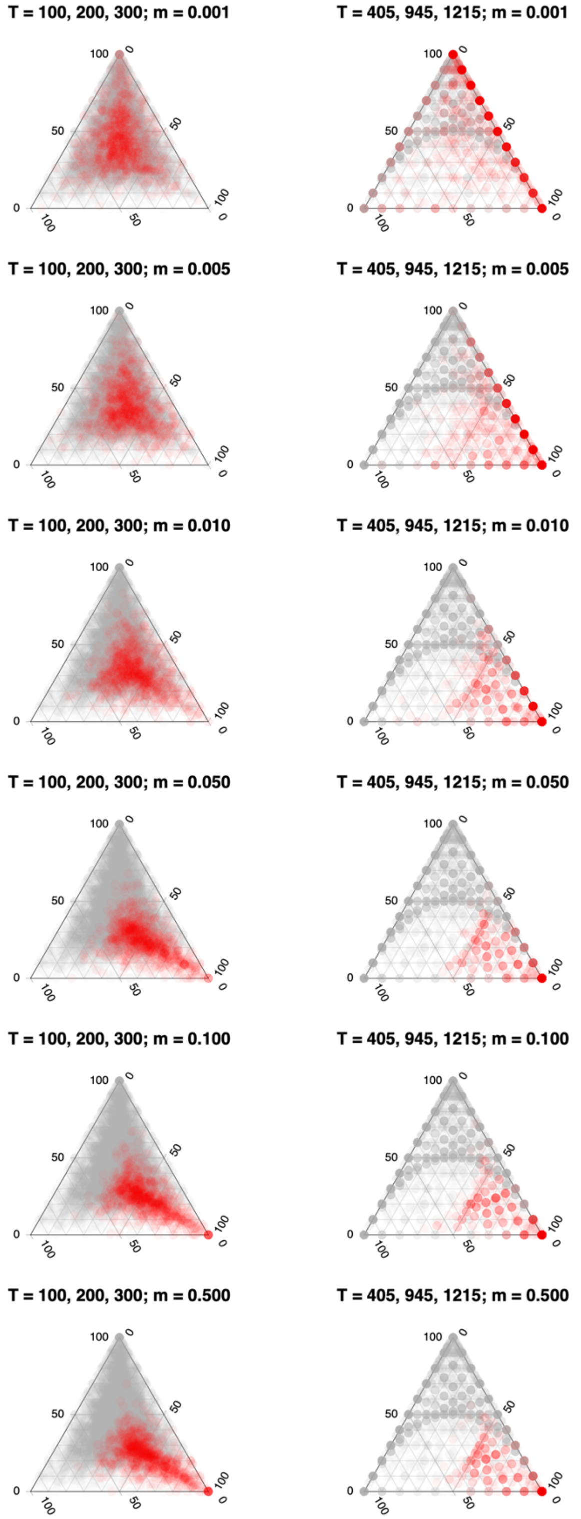

**Figure S17. Joint distribution of topology weights for simulations with gene flow for 10% of the genome.** Each column shows six simulations with the same split times (T), but with $m = 0$ for 90% of the genome (9000 blue dots) and $m = 0$, 0.001, 0.005, 0.010, 0.050, 0.100, or 0.500 for the 10% of the genome (1000 red dots). All lineages have an $N_e$ of 500.

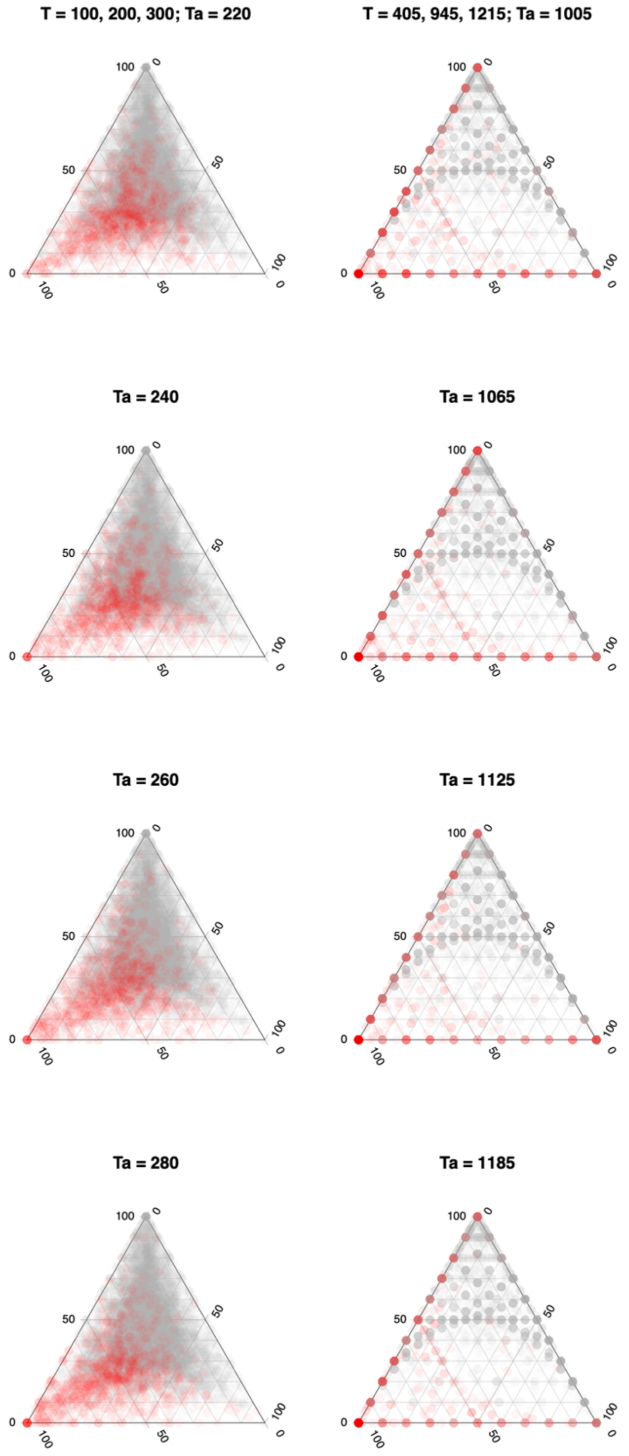

**Figure S18. Joint distribution of topology weights for simulations with ancestral structure.** Each column shows four simulations with the same split times for 90 % of genomic windows (gray dots). 10 percent of the genome evolves under an alternative demography (O,(P1(P2,P3))), with a higher split time for T2 to mimic a scenario with ancestral structure (red dots). The Ta values used are shown above each plot.

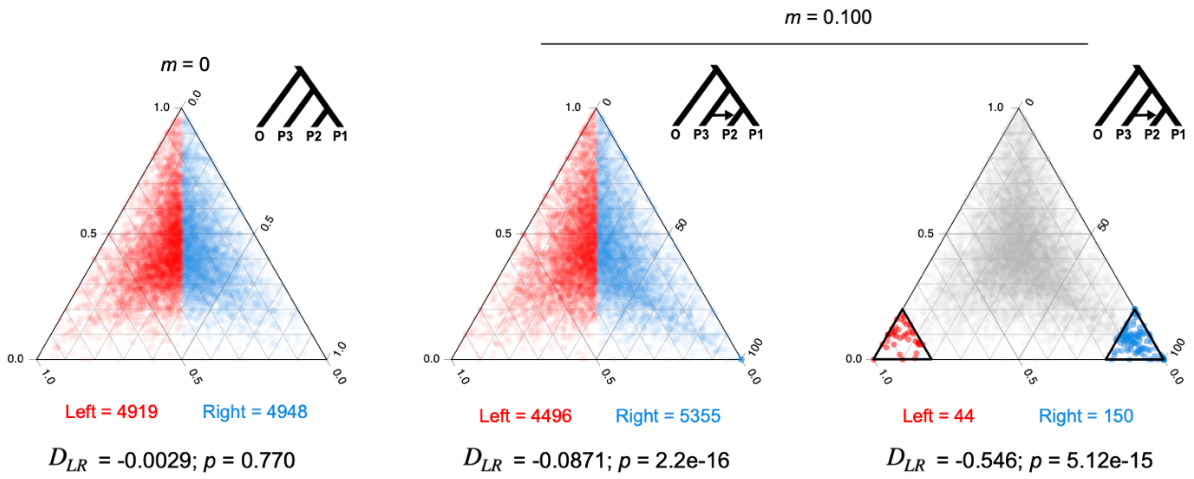

**Figure S19. Example of symmetry analysis of simulated topology weights.** The $D_{LR}$ statistic calculated for two different simulated distributions, each consisting of 10,000 non-recombining loci. The counts of the left- and right-sided windows, estimate of $D_{LR}$, and significance of difference from 0 are given below each plot. Left-sided windows are colored red and right-sided windows are blue. The left plot shows a distribution of topology weights calculated under an ideal four-population model with uniform effective sizes and split times (Scenario a; fig. S9). A genome-wide test for asymmetry is conducted by calculating $D_{LR}$ between the full left and right half triangles. The negative estimate of $D_{LR}$ (-0.0029) indicates that there is a small bias toward Tc, with 0.29% fewer windows in the left side of the plot than expected (*i.e.*, There is an equal probability of a point falling on the left or right side). However, the observed bias is not significantly different from the expectation of equality (G-test, p = 0.770). The two plots on the right side show a different distribution of weights, with 90% of the genome simulated under a model without migration but where migration occurs at a rate of 0.1 between P3 and P2 for 10 % of loci (Scenario h; fig. S9). In the middle plot, the genome-wide estimate of $D_{LR}$ (-0.0871) indicates an 8.7% excess of windows on the right side of the plot, which is much greater than expected by chance (G-test, p = 5.12e-15). In the right plot, $D_{LR}$ is estimated between the extreme left- and right-sided sub triangles. In this case, there is very strong and significant asymmetry, with 54% more windows on the right than we would expect by chance ($D_{LR}$ = -0.546, p = 5.12e-15).

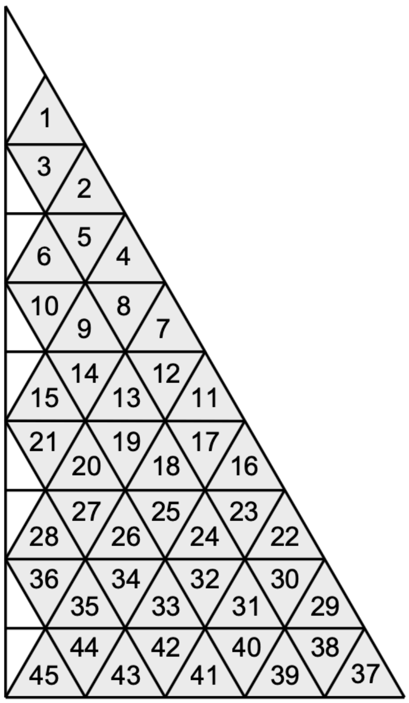

**Figure S20. Identities of the sub-triangles analysed in the fine scale symmetry analysis**. The numbers correspond to the sub-triangle IDs given in table S5 and main text Fig. 2D. The central sub-triangles of the ternary plot (here, seen as small white right-angled triangles on the left side of the plot) cannot be divided into triangles at the same scale so were not considered in the analysis.

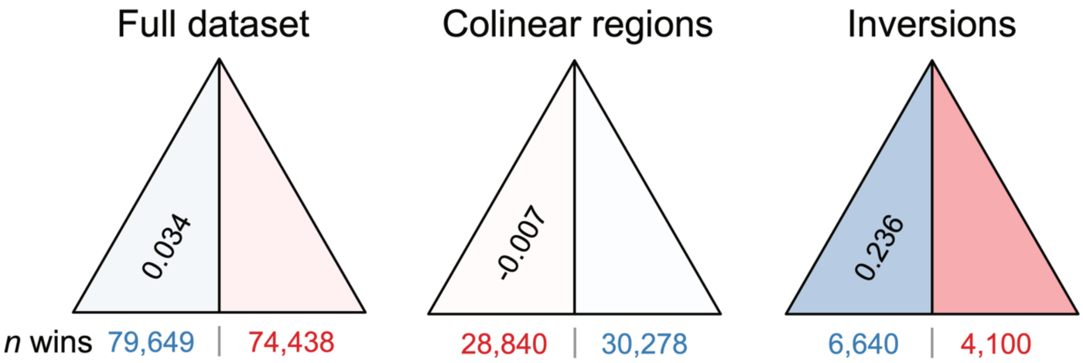

**Figure S21. Analysis of symmetry for different partitions of the genome.** Results for the full dataset, colinear regions, and chromosomal inversions are show. $D_{LR}$ is provided in the left-hand sub-triangle. The intensity of shading in each side is proportional to the excess (blue) or deficit (red) of windows relative to the expectation. The numbers of windows falling on each side are shown under the plot.

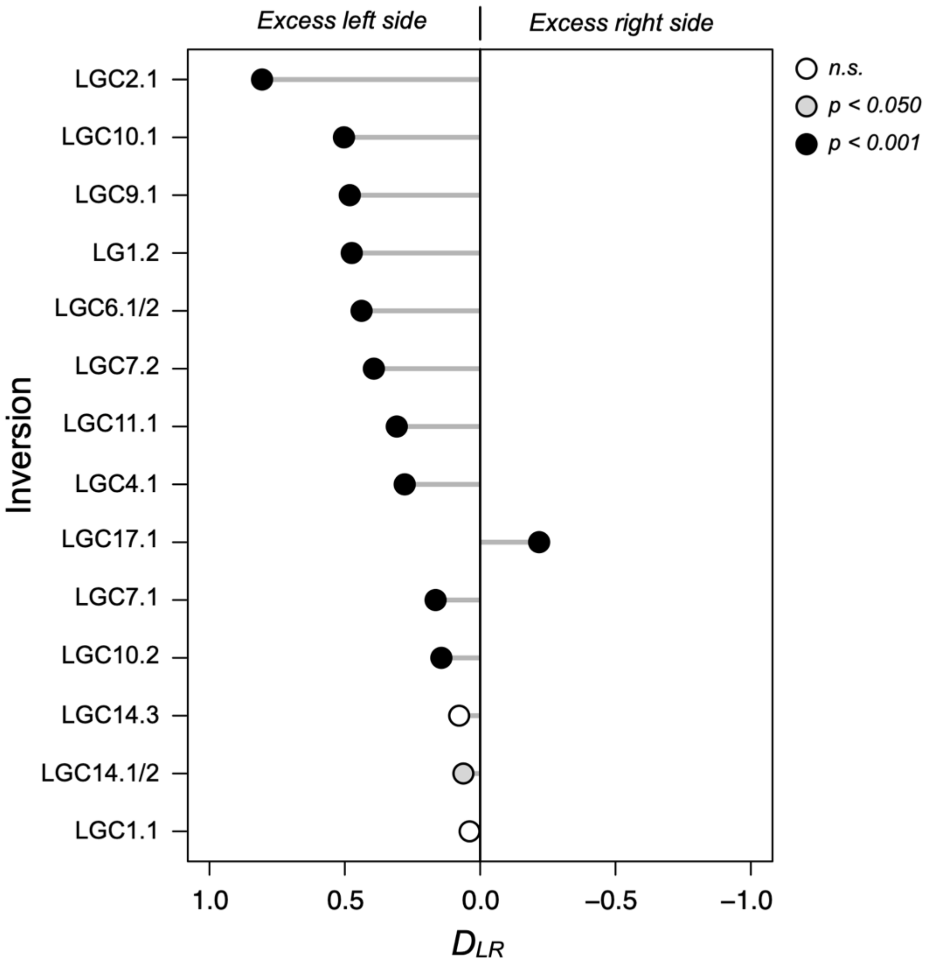

**Figure S22. Estimates of $D_{LR}$ for each chromosomal inversion.** The line running through the center of the plot indicates a $D_{LR}$ of 0. The X-axis runs from positive 1 to negative 1, as positive estimates of $D_{LR}$ indicate a bias of windows on the left side of the triangle, while negative estimates indicate a bias on the right side.

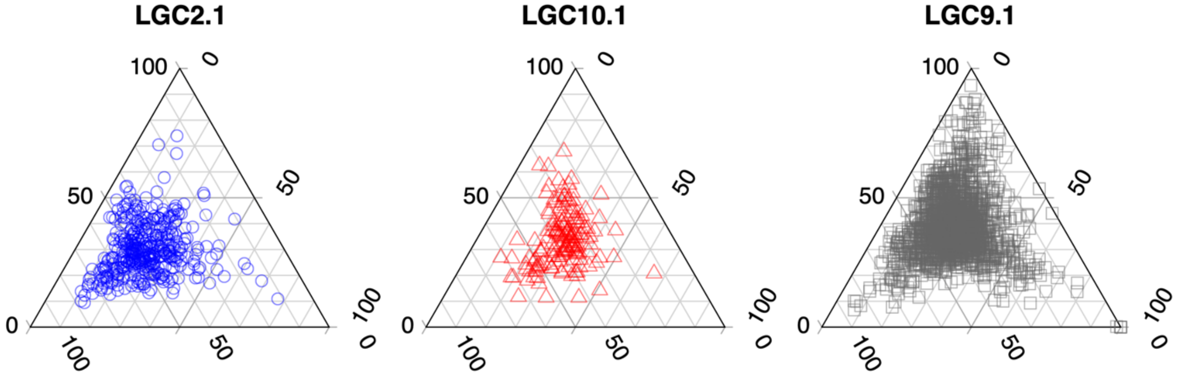

**Figure S23. Ternary plots for the three inversions that show the largest values of $D_{LR}$.** Each point represents a single genomic window.

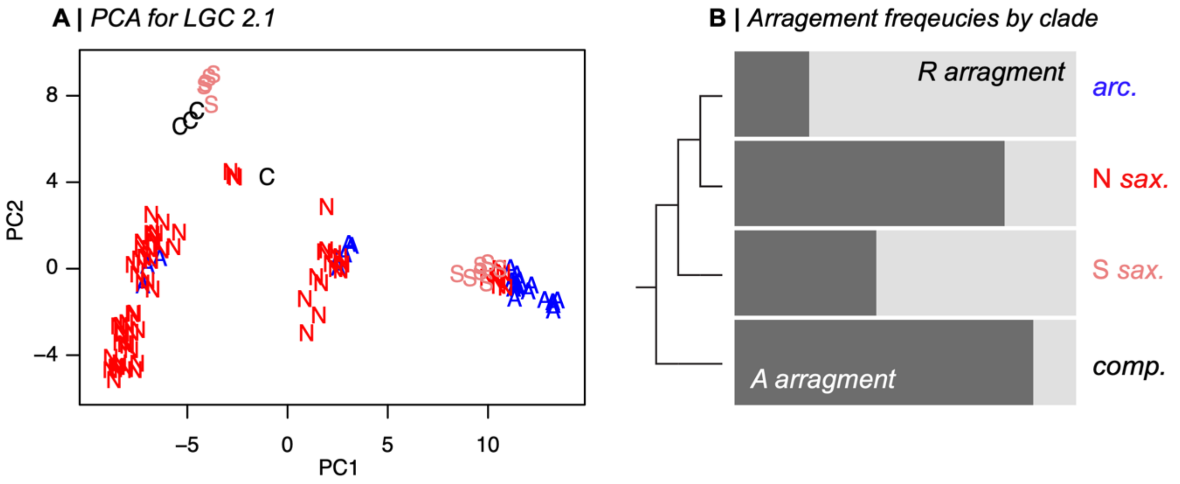

**Figure S24. PCA and arrangement frequencies for inversion LGC2.1.** (A) The PCA is conducted on the SNP genotype matrix for loci that fall inside the inversion. Samples correspond within clade names (A: *L. arcana*; N: Northern *L. saxatilis*; S: Spanish *L. saxatilis*: C: *L. compressa*) (B) Frequencies of the 2 arrangements (R and A) in each clade inferred from the clusters in the PCA.

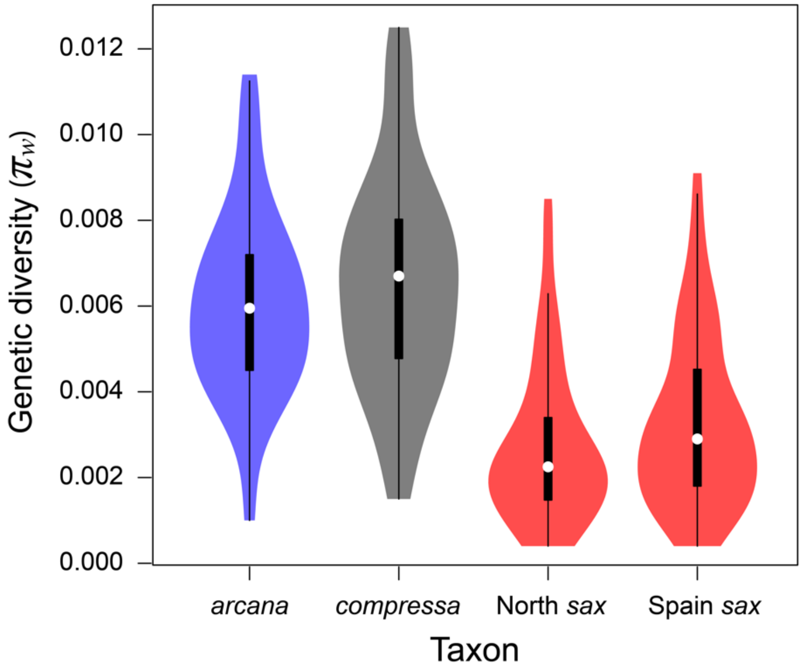

**Figure S25. Violin plots showing the distributions of $\pi_w$ for windows perfectly associated with reproductive mode.** The white point is the median, the thick black bar is the interquartile range and the thick black line is 1.5 times the interquartile range. This figure expands on main text Fig. 3A, where we show only the mean values for each clade; here we show the full distributions. These results clearly show that live-bearing clades show lower diversity than the egg-layers, emphasizing that the result is not just driven by lower diversity in one of the live bearing clades. They also reveal that Northern *L. saxatilis* shows slightly but significantly lower diversity than Spanish *L. saxatilis* in these regions (median $\pi$ for Northern = 0.0027 vs. 0.0033 for Spanish, Wilcoxon test, p = 1.49e-06).

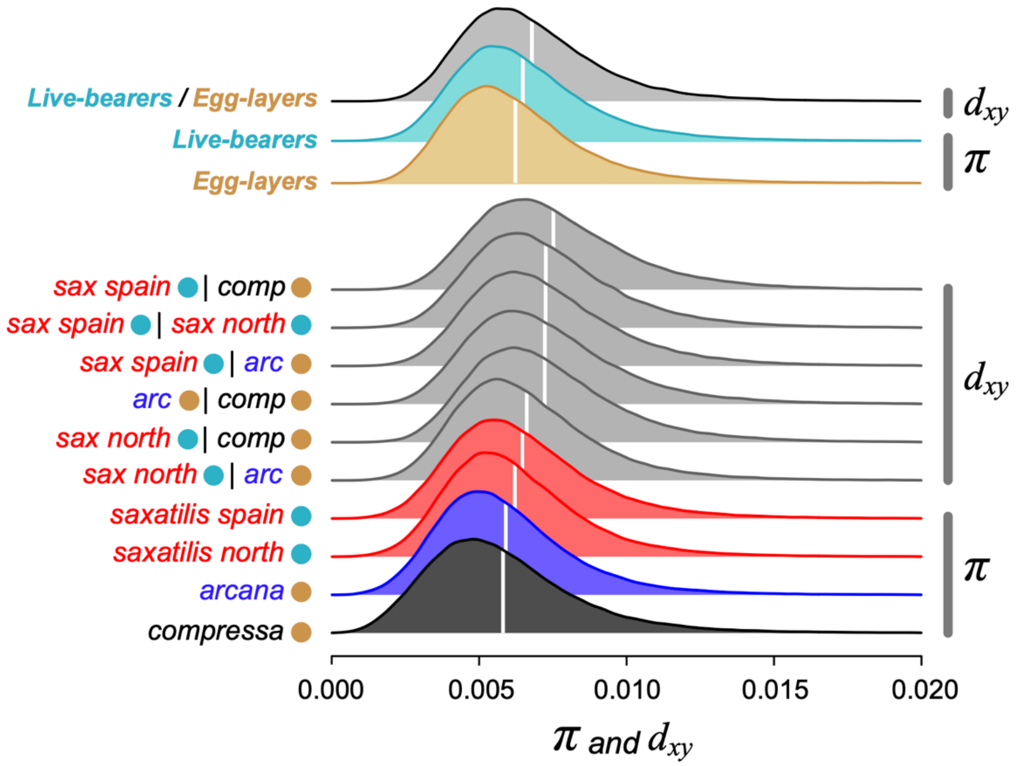

**Figure S26. Distribution of within-group diversity ($\pi$) and between-group divergence ($d_{xy}$) for 100 SNP windows.** The distribution is truncated at 0.02, to emphasize the main distributions rather than the long tail. The white lines show the mean estimate for each distribution. The cyan and gold circles indicate the reproductive mode of the taxon (live-bearing and egg-laying, respectively). This figure expands on Fig. 3A of the main text, where we show the median estimates of nucleotide diversity ($\pi$) within live bearers and egg-layers (*i.e.*, median $\pi$ over all 100 SNP windows). The full distributions here highlight that egg-layers and live-bearers have similar levels of genome-wide nucleotide diversity. In fact, a paired Wilcoxon test reveals that live-bearers have marginally higher diversity than egg-layers ($p < 2.2e\text{-}16$), making it even more unlikely that we would see consistently reduced diversity in high Tr windows in live-bearers by chance. Because this comparison is made between groups that are a mix of individuals from two different clades, we also examined diversity within each clade to ensure that our key results are not an artifact of the mixing of different groups. We found the same result, with the two egg-layers showing lower diversity than either of the live-bearers. Spanish *L. saxatilis* had the highest diversity, and *L compressa* the lowest.

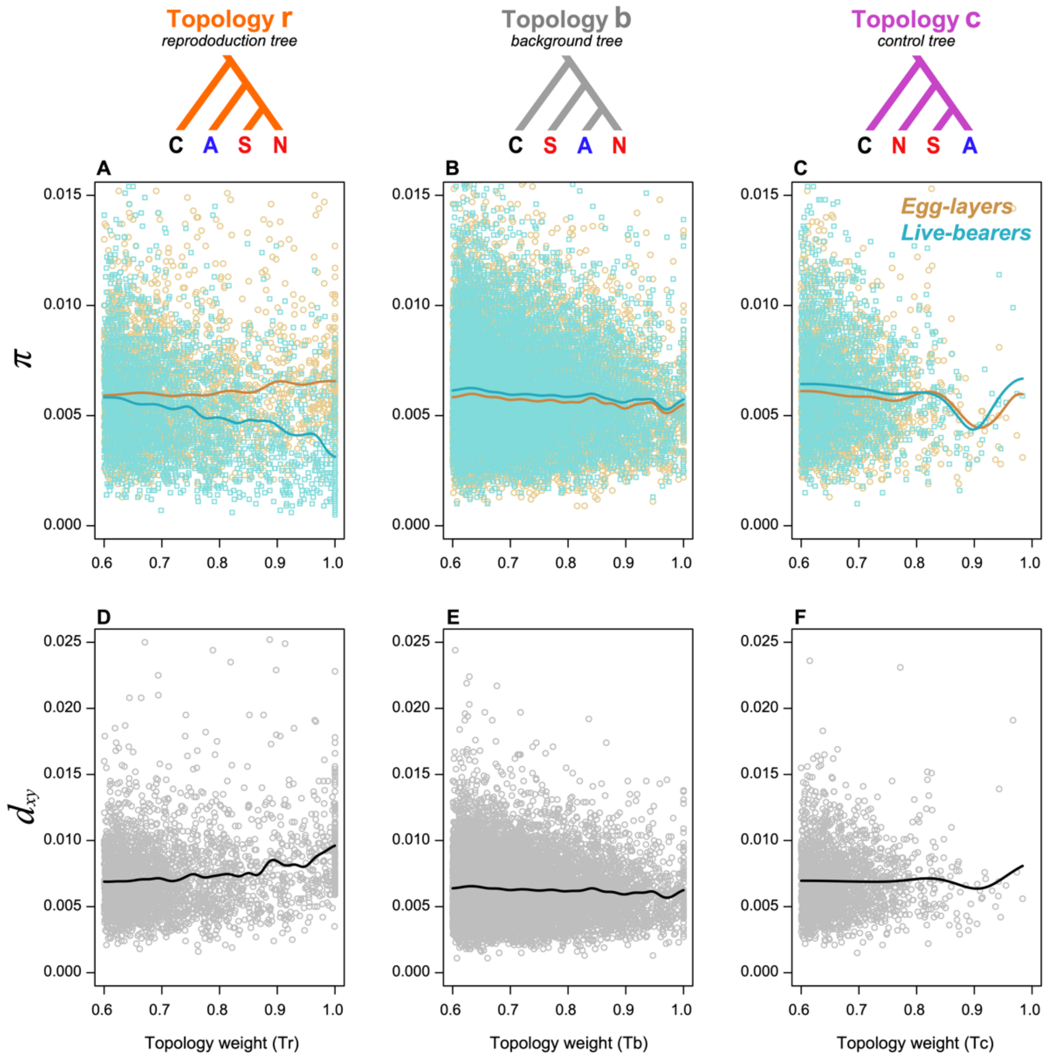

**Figure S27. Relationship between genetic diversity (π), divergence ($d_{xy}$) and topology weights for 100 SNP genomic windows.** Each point represents a single genomic window (egg-layers in gold, live-bearers in cyan, between modes in gray). The solid lines in the plot show the weighted moving average computed by kernel smoothing. This figure expands on main text Fig. 3A, by showing the relationships for all three topology weights. As mentioned in the main text, we see no obvious relationship between Tr and π for egg-layers (Spearman correlation rho = 0.043, p = 0.023). However, for live-bearers, we see that genetic diversity decreases with increasing Tr weight (Spearman correlation rho = -0.256, p < 2.2e-16). For the other two topologies, Tb and Tc, we do not see any notable difference in the relationship with π between egg-layers and live-bearers. This clearly shows that the decrease in diversity with increasing Tr weight in live-bearers is specific only to Tr. We also see that $d_{xy}$ tends to increase with Tr. This is most pronounced for windows perfectly associated with reproductive mode, which show a $d_{xy}$ that is significantly higher than windows where Tr < 1 (median $d_{xy}$ = 0.010 for perfectly associated windows vs. 0.0064 for the background, Wilcoxon test, p < 2.2e-16). but with no clear relationship for Tb or Tc (Tb: rho = -0.034, p = 0.016; Tc: rho = -0.038, p = 0.410).

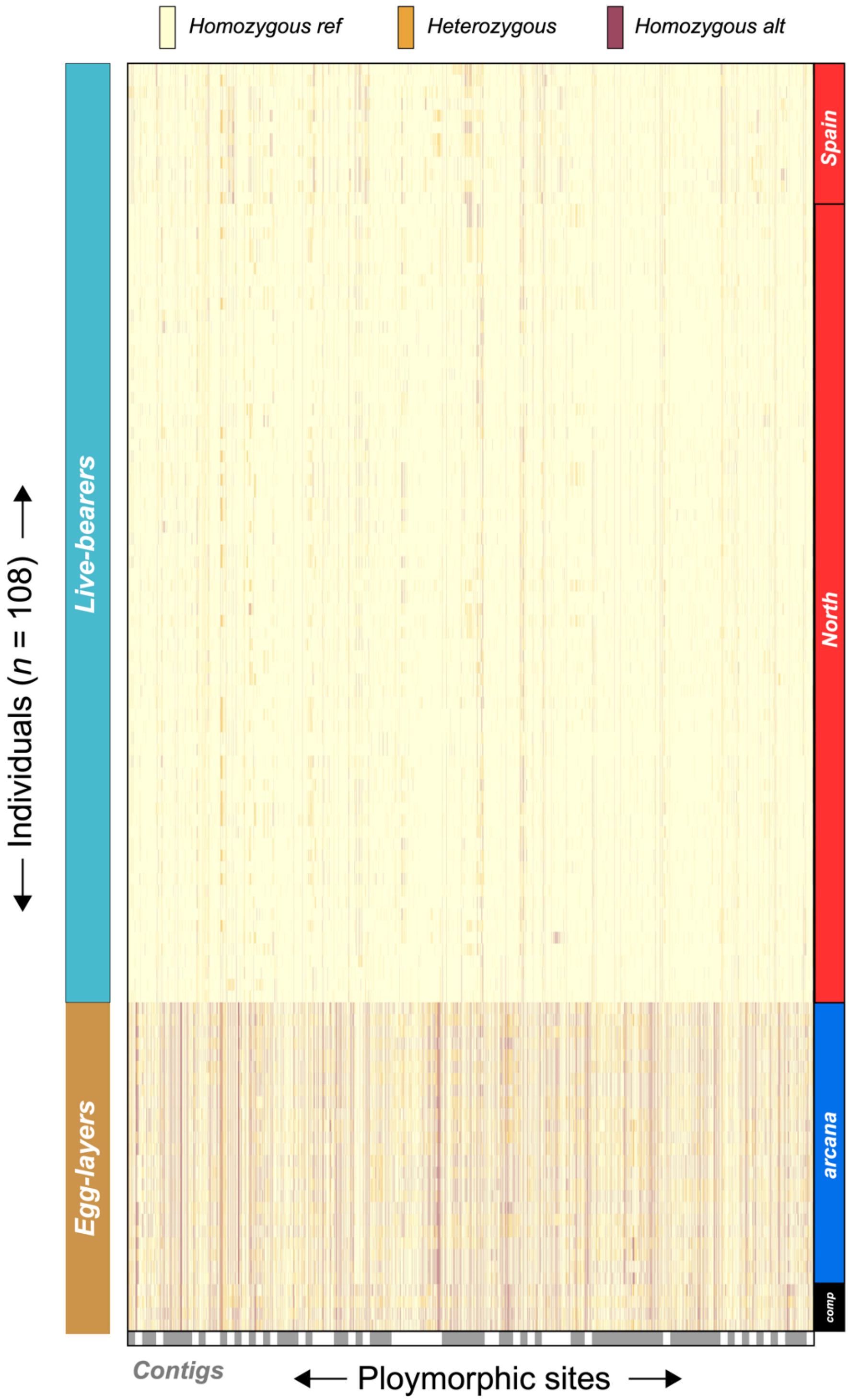

**Figure S28. Visualization of the genotype matrix for the 50 regions perfectly associated with live birth.** Each row corresponds to an individual and each column corresponds with a polymorphic site. Individuals are grouped according to their mode (left boxes) and clade in the genome wide phylogenetics analysis (right boxes). Sites are grouped by assembly contig (alternating gray and white boxes). The individual's diploid genotype at each site is colored according to the key at the top of the plot.

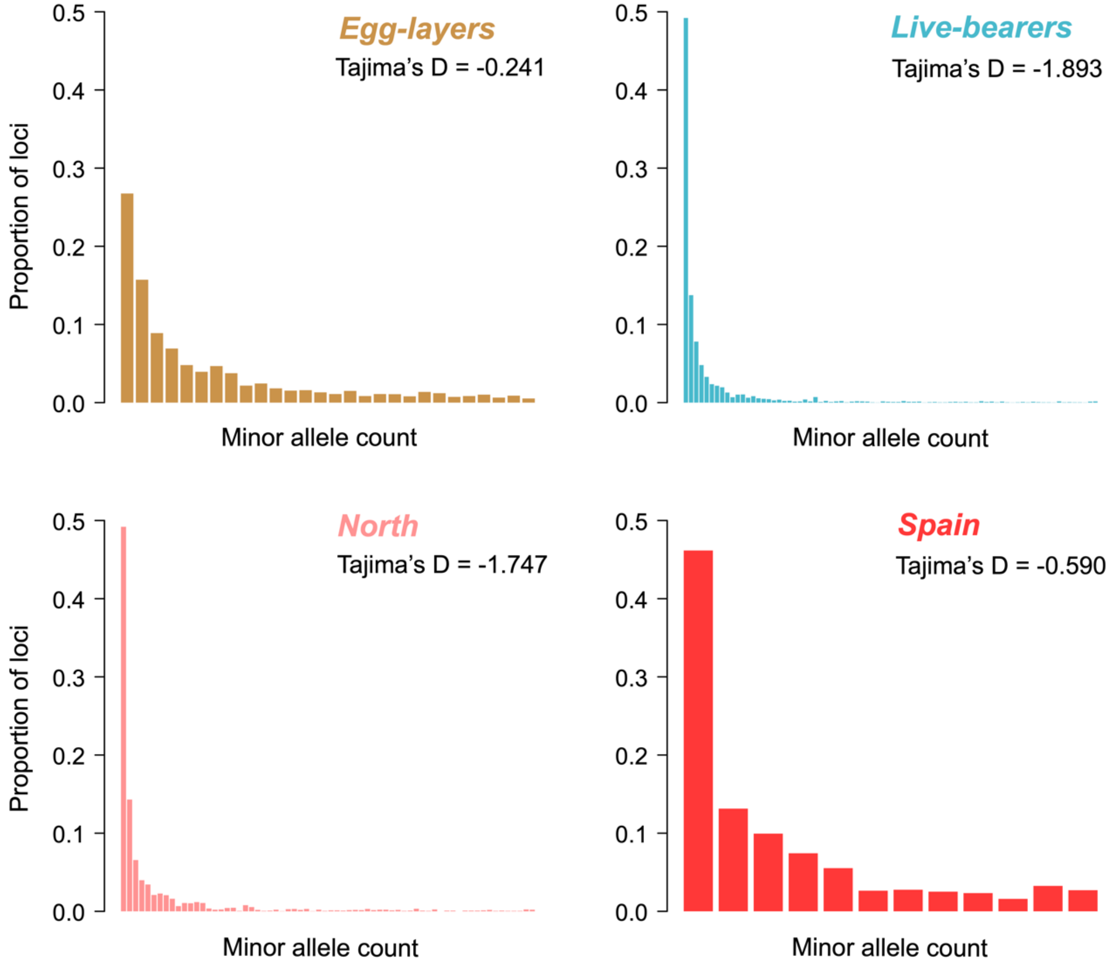

**Figure S29. Full folded site frequency spectra for loci within regions perfectly associated with reproductive mode.** Because the number of polymorphic loci differs between the groups the bars have been scaled to represent the proportion of loci associated with each minor allele count. Tajima's D for each group is also given. Sample sizes for each group: egglayers, $n = 28$; live-bearers $n = 80$, North, $n = 68$; Spain, $n = 12$. We compared the spectra for live-bearers and egg-layers and between Northern *L. saxatilis* and Spanish *L. saxatilis* by creating and comparing 10,000 sample-size matched SFS. In both cases, all 10,000 spectra were significantly different ($p < 0.05$.)

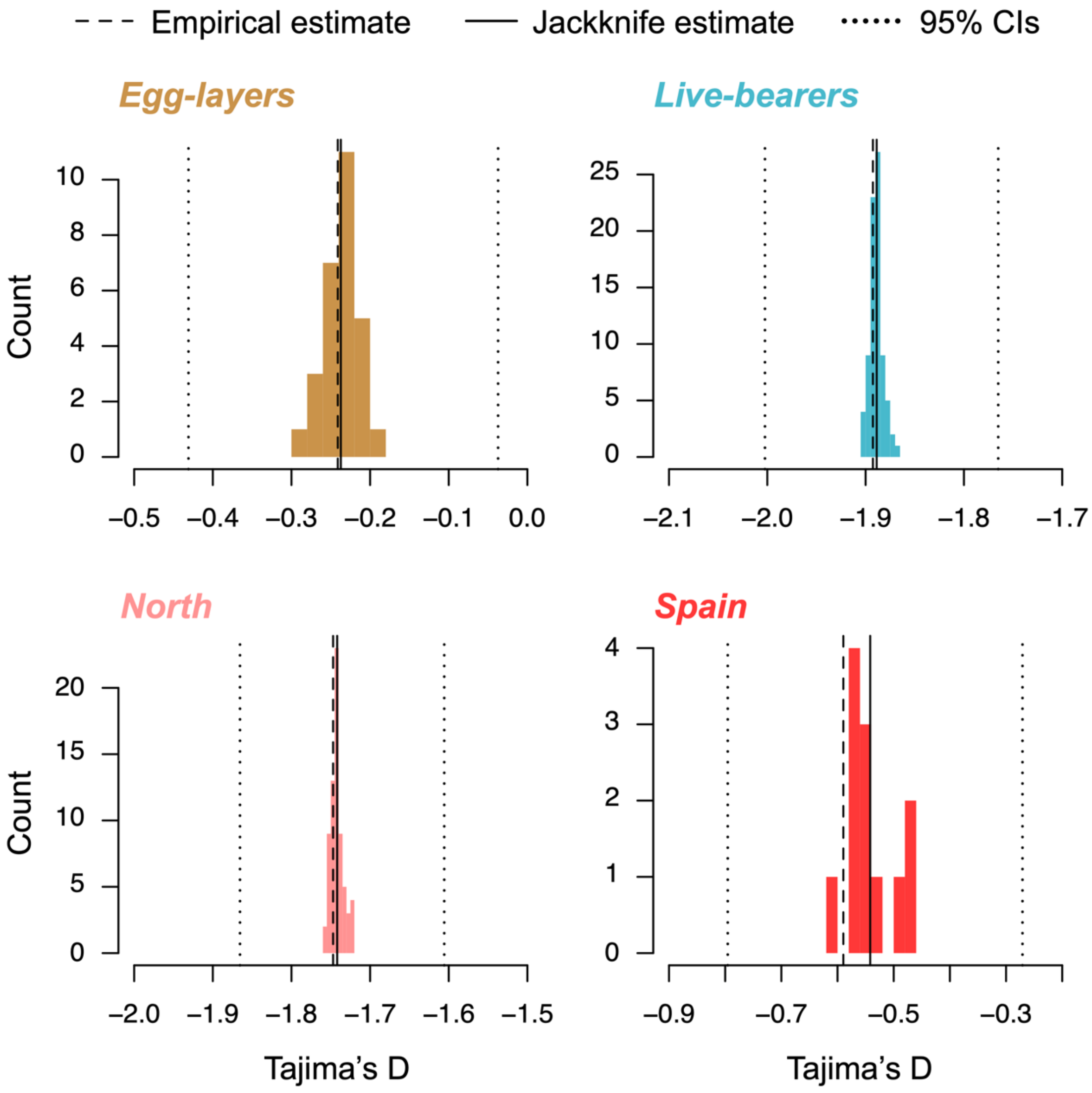

**Figure S30. Distributions of Tajima's D calculated for loci within regions perfectly associated with reproductive mode by jackknife resampling.** The colored histograms show the distributions of the $n - 1$ estimates of Tajima's D. The dashed line indicates the value of the empirical point estimate, and the solid line shows the mean of the $n - 1$ estimates. The dotted lines are the upper and lower 95% confidence intervals.

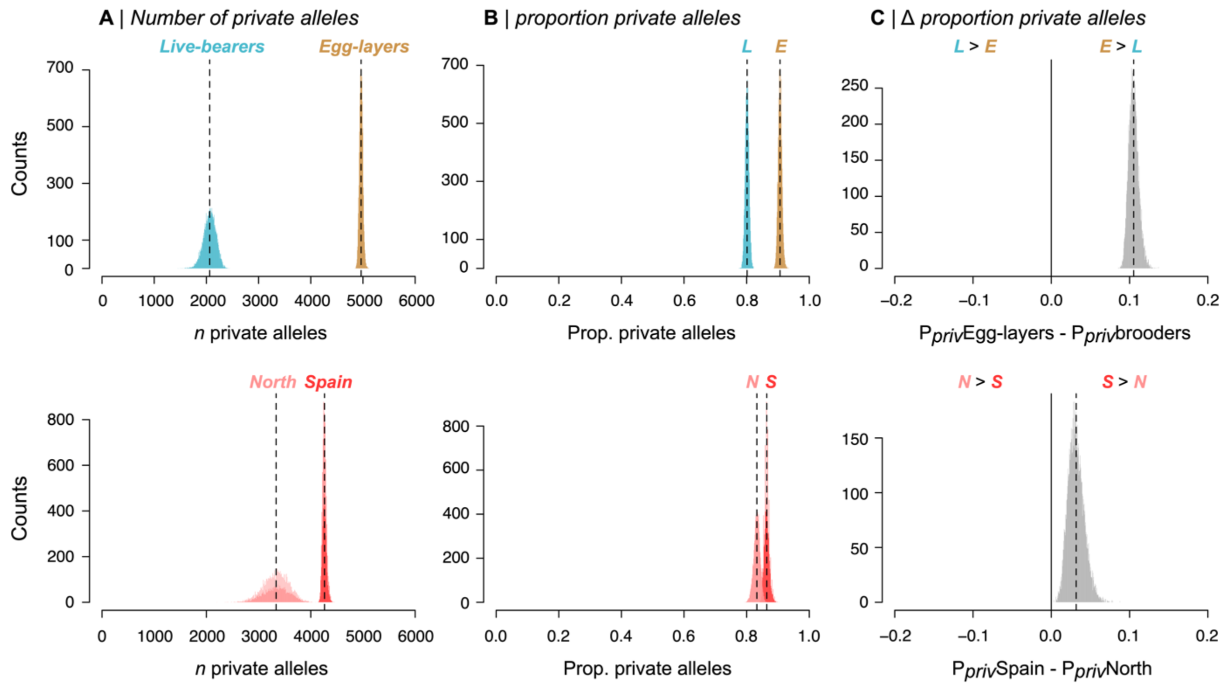

**Figure S31. Sample-size adjusted estimates of the number and proportion of private alleles per group for loci in regions perfectly associated with reproductive mode**. Estimates were obtained by down-sampling the larger group to the size of the smaller one without replacement, such that each plot shows the results for 10,000 random subsamples of the larger taxon. The dashed line shows the mean estimate for each group. A) the number of private alleles in live-bearers and egg-layers (live-bearers down sampled to 28). B) the proportion of private alleles ($P_{priv} = n_{private} / n_{polymorphic}$) in live-bearers and egg-layers. C) the difference in the proportion of private alleles between the egg-layers and live-bearers ($P_{priv}$ egg-layers – $P_{priv}$ live-bearers); the black solid line at position 0 indicates where the proportion of private alleles in each group is equal. The bottom row is the same as the top plot, except that loci with alleles that are private to the Spanish and Northern groups of live-bearers are being compared. This analysis clearly shows that egg-layers and live-bearers both harbor a large fraction of private alleles, though the proportion is higher in egg-layers. Spanish *L. saxatilis* also showed a slightly higher fraction of private alleles than Northern *L. saxatilis*.

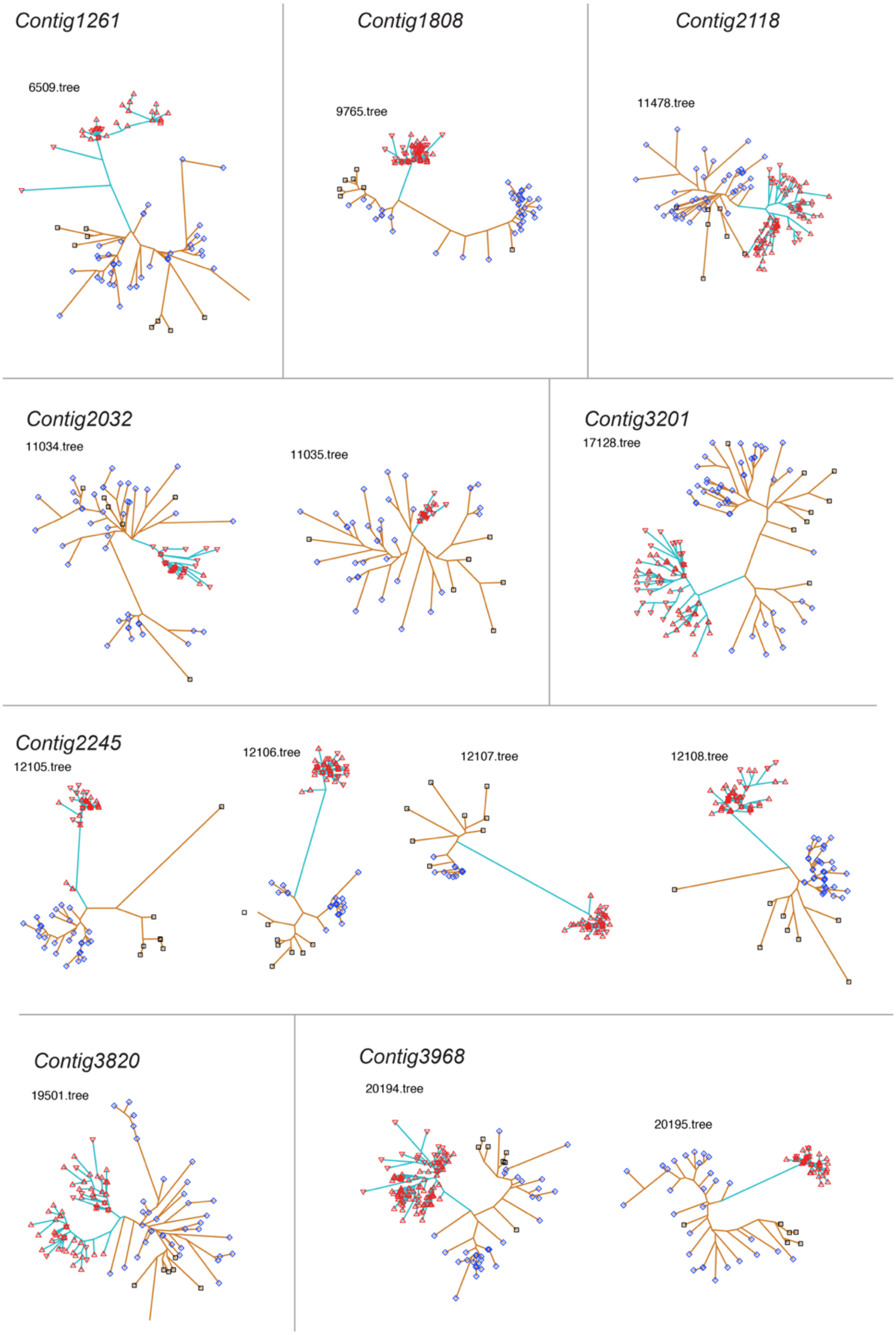

**Figure 32. Unrooted trees for all 88 windows where $T_r = 1$.** Clades are colored according to the reproductive mode (gold egg-layers, cyan live-bearers). Symbols indicate the taxon. North *L. saxatilis*, red up-facing triangles; Spain *L. saxatilis*, red down-facing triangles; *L. arcana*, blue diamonds; *L. compressa*, black squares. Trees on different contigs are separated by borders and ordered left to right or top to bottom. The figure continues over the following 5 pages.

2628

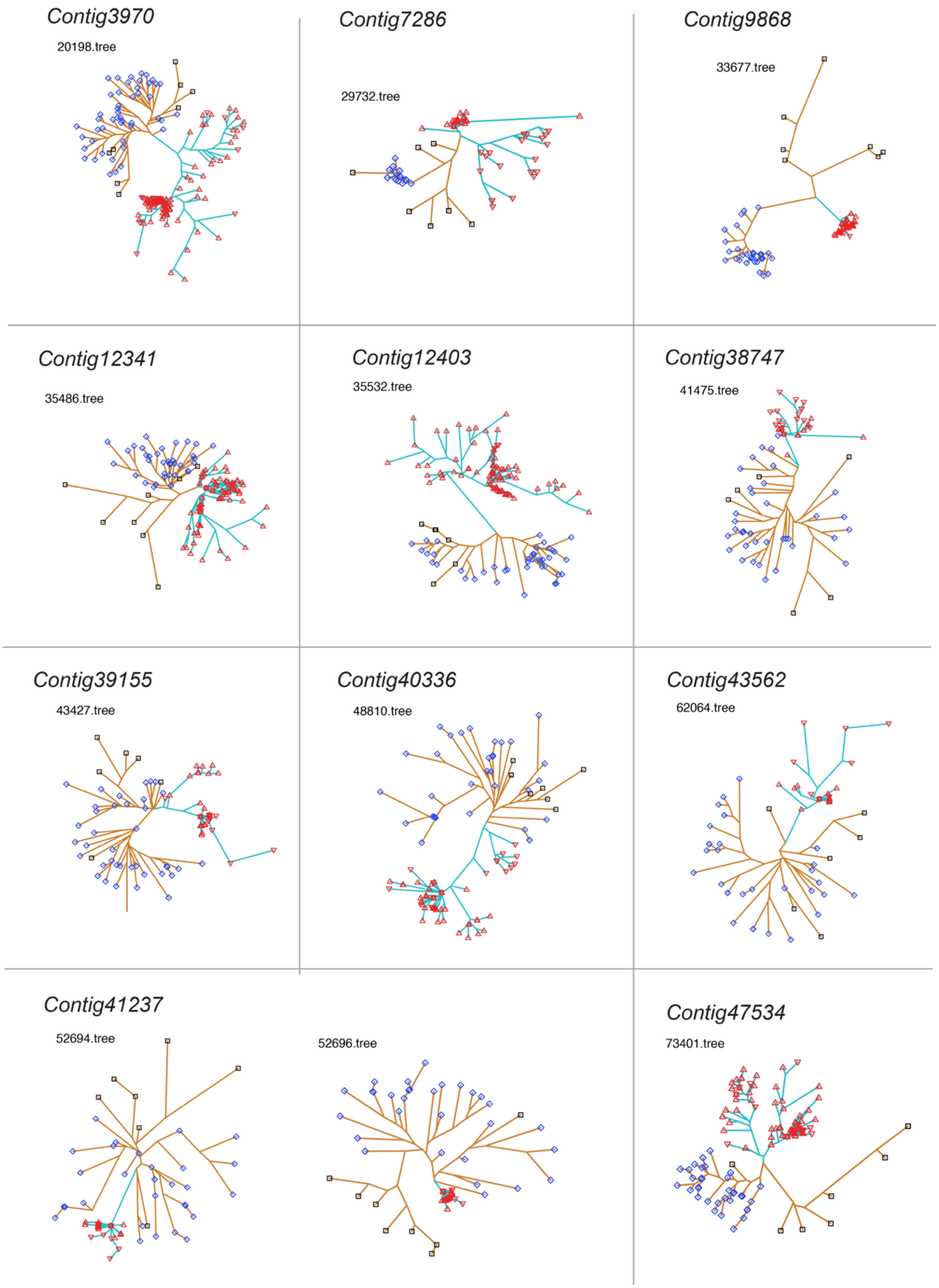
Contig3970
20198.tree
Contig7286
29732.tree
Contig9868
33677.tree
Contig12341
35486.tree
Contig12403
35532.tree
Contig38747
41475.tree
Contig39155
43427.tree
Contig40336
48810.tree
Contig43562
62064.tree
Contig41237
52694.tree
52696.tree
Contig47534
73401.tree

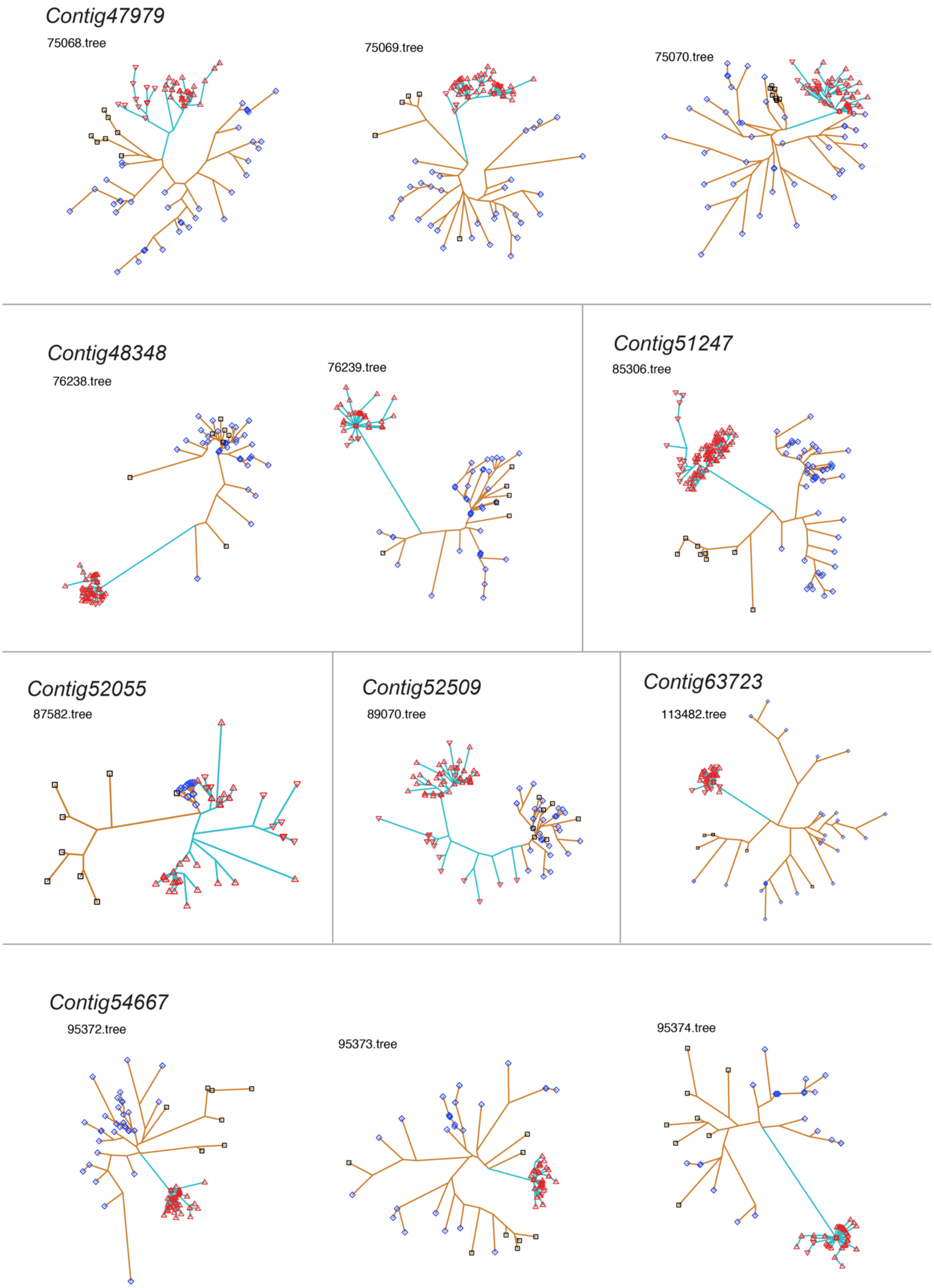
Contig47979
75068.tree
75069.tree
75070.tree
Contig48348
76238.tree
76239.tree
Contig51247
85306.tree
Contig52055
87582.tree
Contig52509
89070.tree
Contig63723
113482.tree
Contig54667
95372.tree
95373.tree
95374.tree

### Contig51247

106684.tree

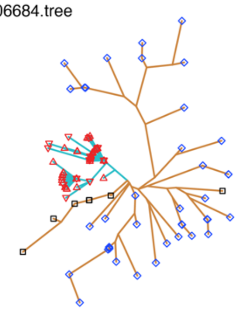

106685.tree

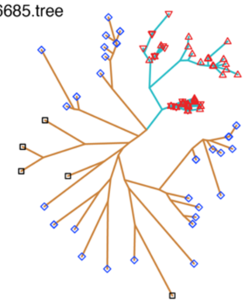

106686.tree

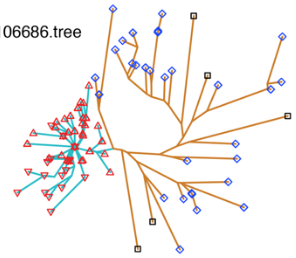

106688.tree

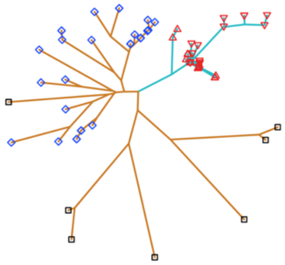

106689.tree

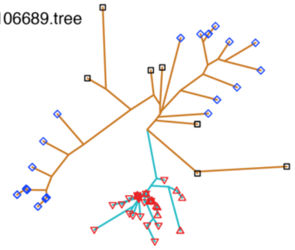

106690.tree

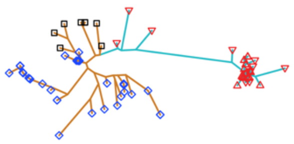

### Contig60406

107721.tree

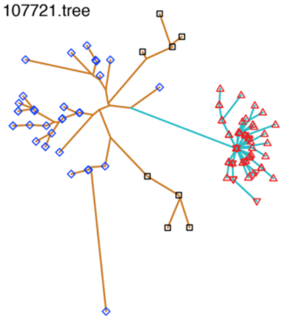

107723.tree

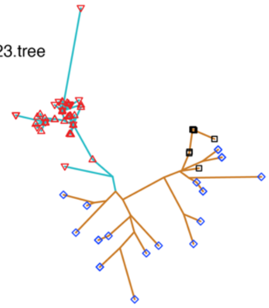

107726.tree

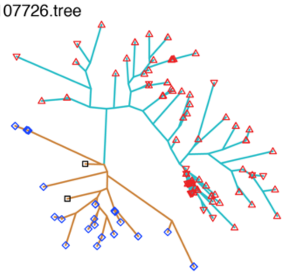

### Contig61156

109076.tree

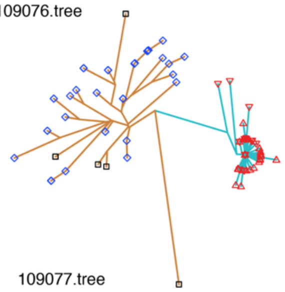

109077.tree

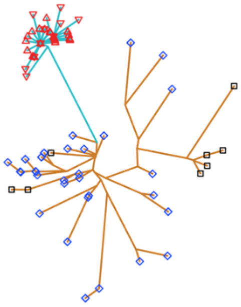

### Contig61890

110564.tree

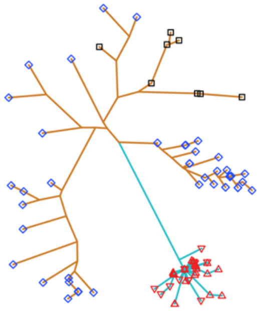

110565.tree

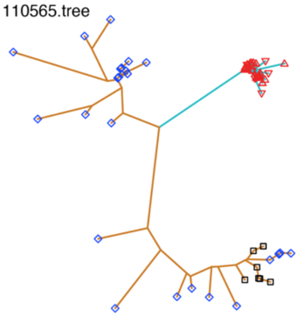

### Contig66365

117438.tree

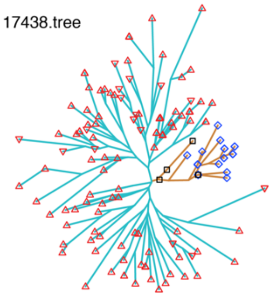

### Contig67554

119150.tree

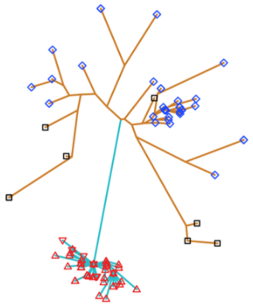

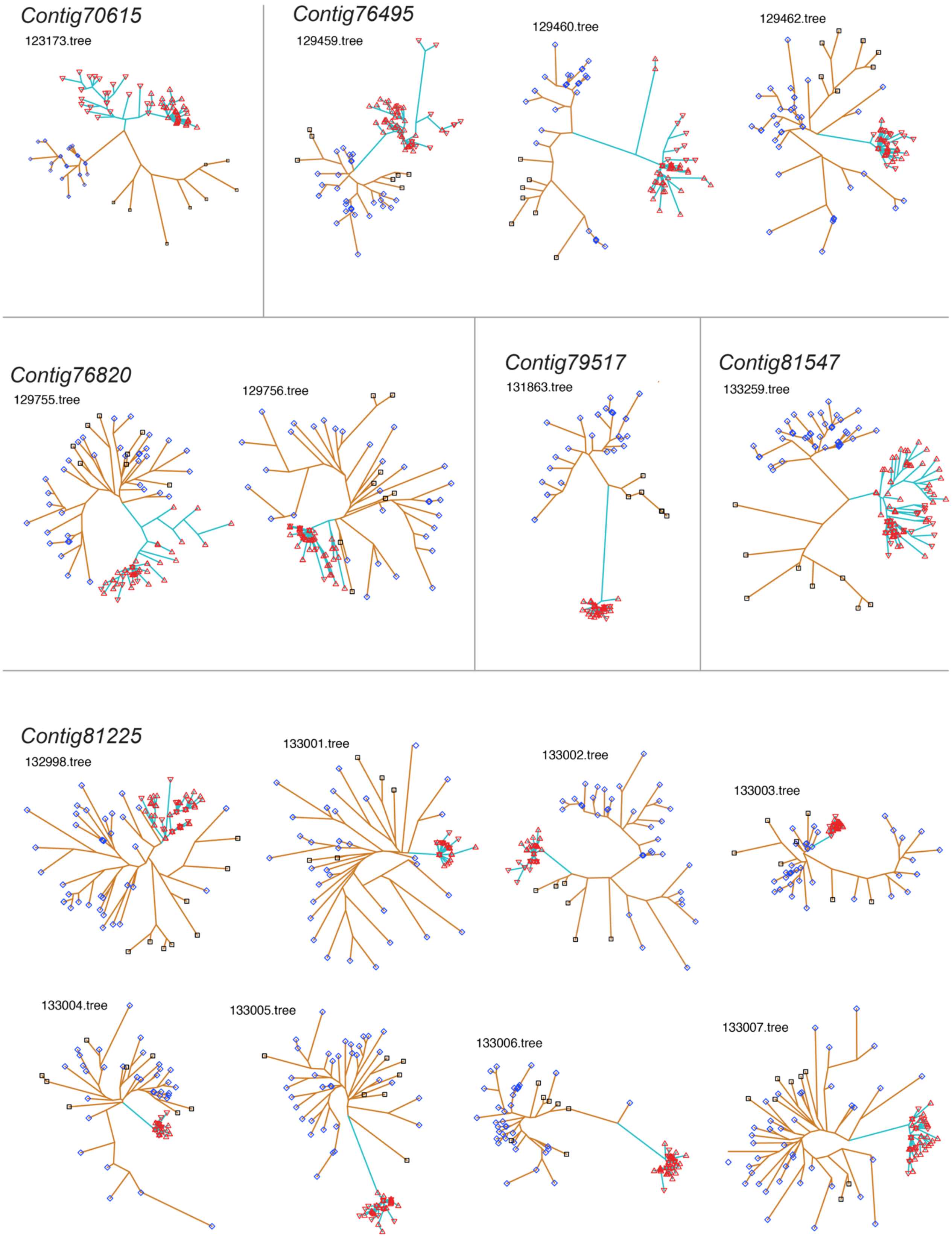
Contig70615
123173.tree
Contig76495
129459.tree
129460.tree
129462.tree
Contig76820
129755.tree
129756.tree
Contig79517
131863.tree
Contig81547
133259.tree
Contig81225
132998.tree
133001.tree
133002.tree
133003.tree
133004.tree
133005.tree
133006.tree
133007.tree

2631

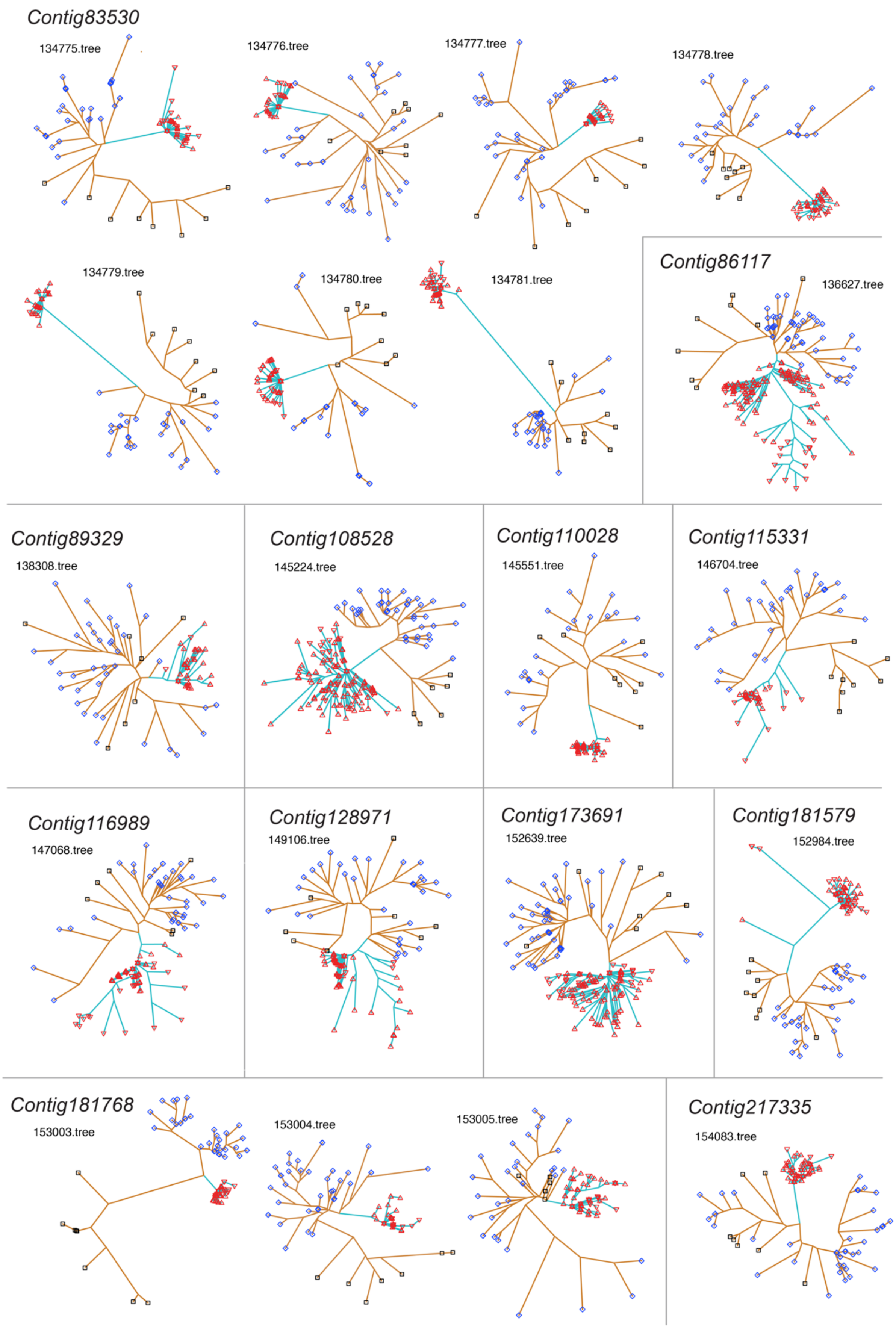
Contig83530
134775.tree
134776.tree
134777.tree
134778.tree
134779.tree
134780.tree
134781.tree
Contig86117
136627.tree
Contig89329
138308.tree
Contig108528
145224.tree
Contig110028
145551.tree
Contig115331
146704.tree
Contig116989
147068.tree
Contig128971
149106.tree
Contig173691
152639.tree
Contig181579
152984.tree
Contig181768
153003.tree
153004.tree
153005.tree
Contig217335
154083.tree

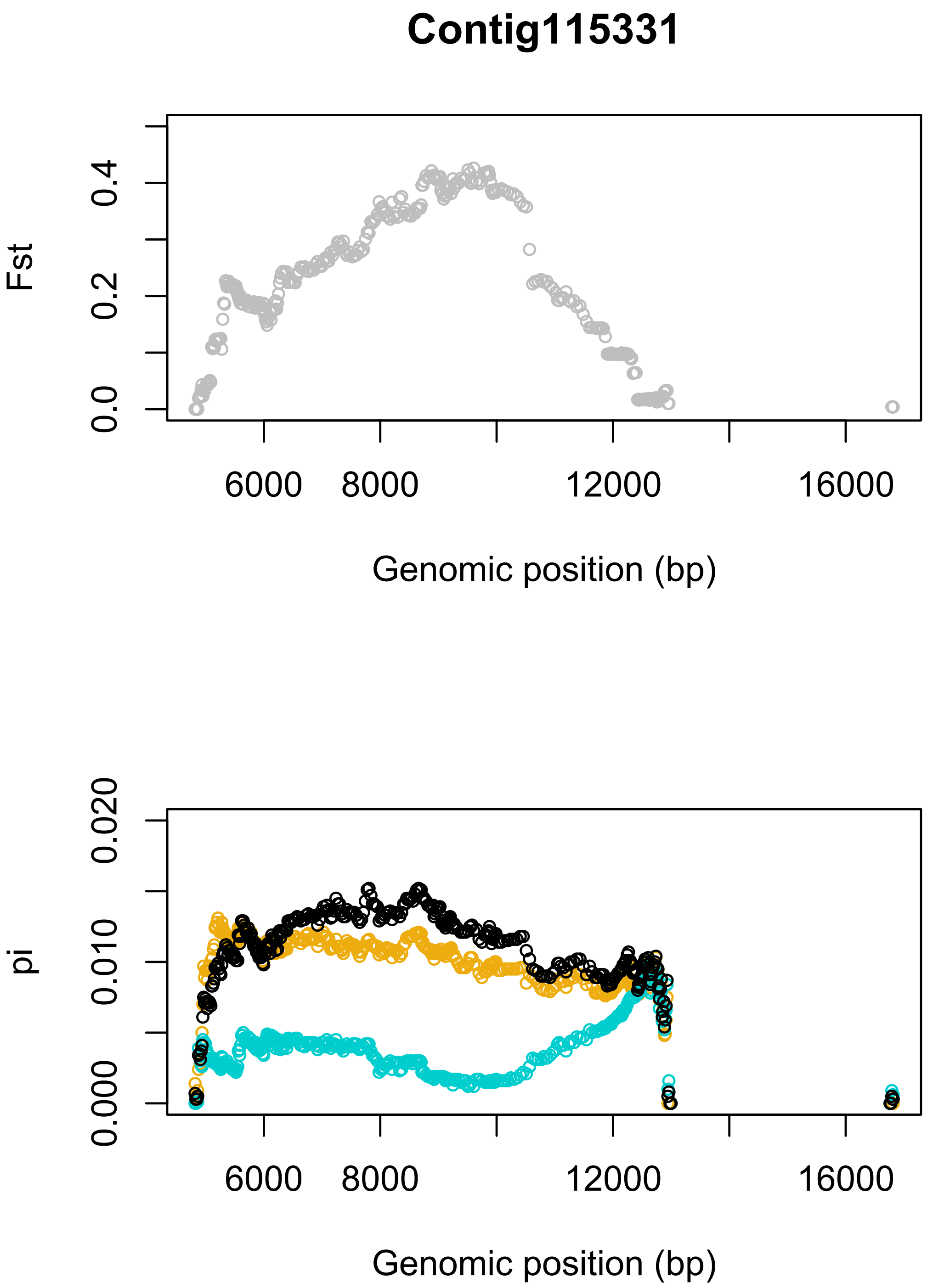

**Fig. S33. Patterns of genetic differentiation (*$F_{ST}$*) and nucleotide diversity (*$\pi$*) across regions where Tr = 1.** Top plot, $F_{ST}$ calculated in 3 kb sliding windows with a 30 bp step. Bottom plot, $\pi$ calculated in 3 kb sliding windows with a 30 bp step. Black line, $\pi_b$ between egg-layers and live-bearers; cyan line, $\pi_w$ within live-bearers; gold line, $\pi_w$ within egg-layers. The figure continues over the next 50 pages. Contigs are shown in the same order as in table S8.

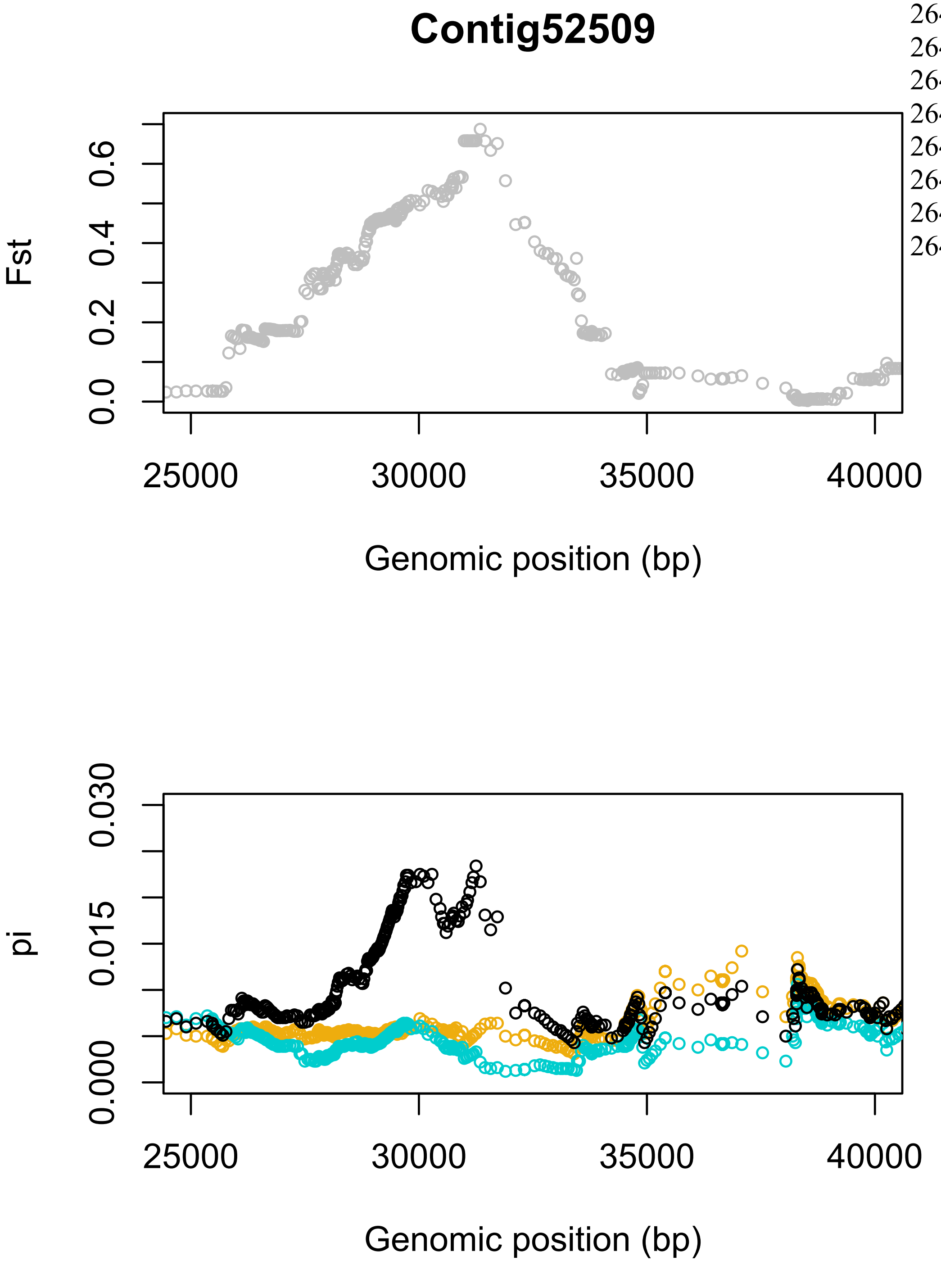
Contig52509
Fst
0.0
0.2
0.4
0.6
25000
30000
35000
40000
Genomic position (bp)
pi
0.000
0.015
0.030
25000
30000
35000
40000
Genomic position (bp)

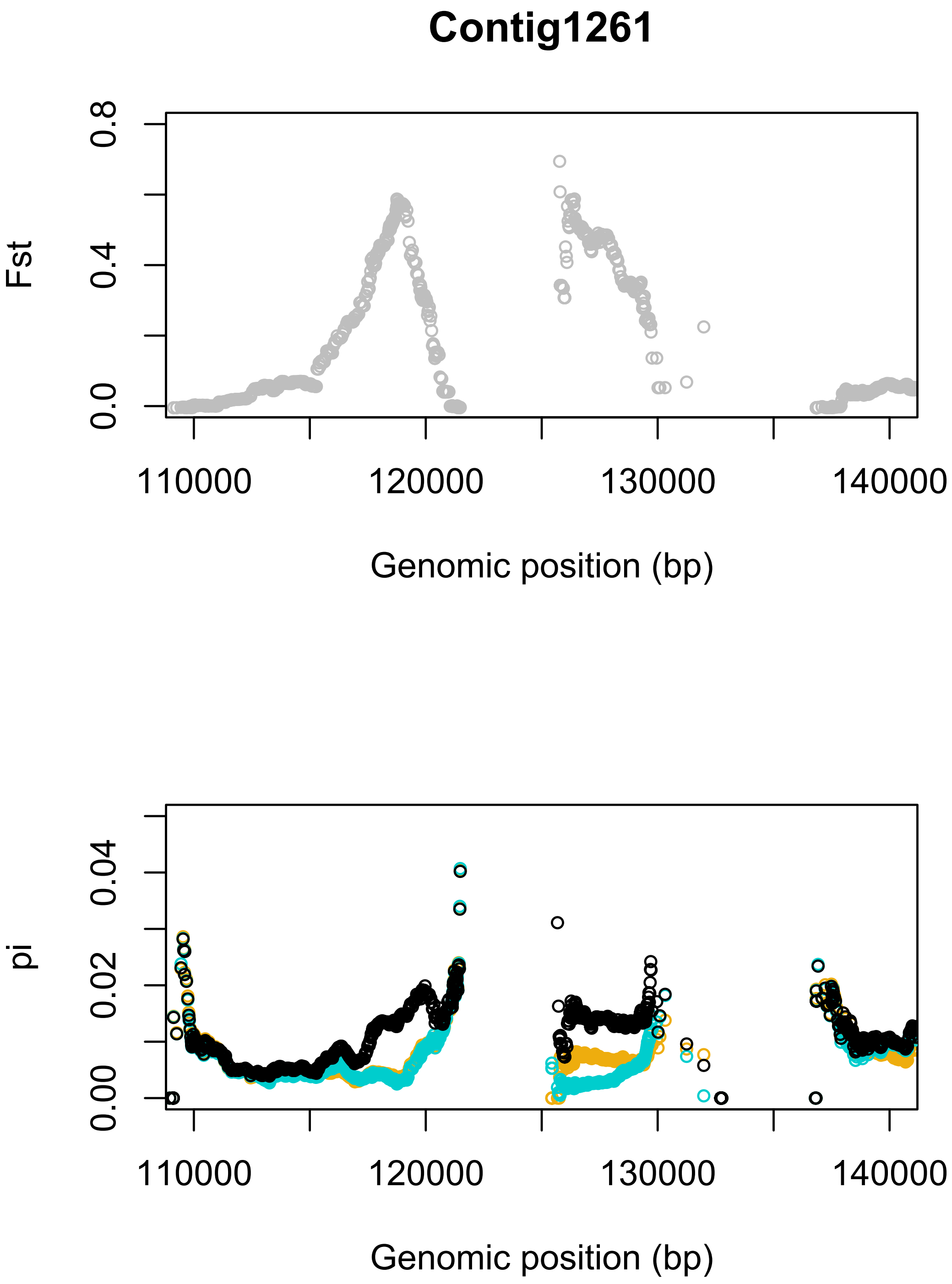
Contig1261
Fst
0.0
0.4
0.8
110000
120000
130000
140000
Genomic position (bp)
pi
0.00
0.02
0.04
110000
120000
130000
140000
Genomic position (bp)

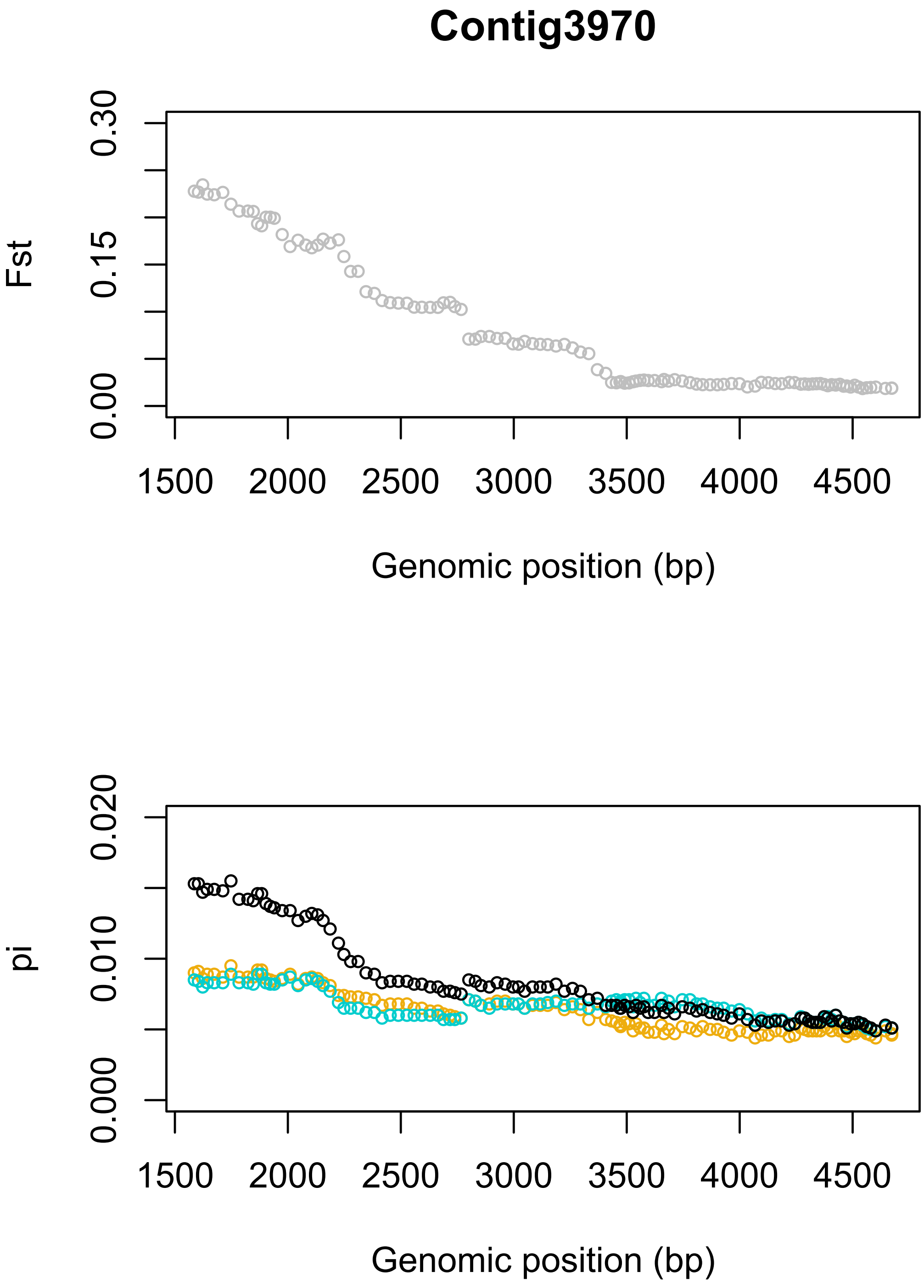
Contig3970
Fst
0.00
0.15
0.30
1500
2000
2500
3000
3500
4000
4500
Genomic position (bp)
pi
0.000
0.010
0.020
1500
2000
2500
3000
3500
4000
4500
Genomic position (bp)

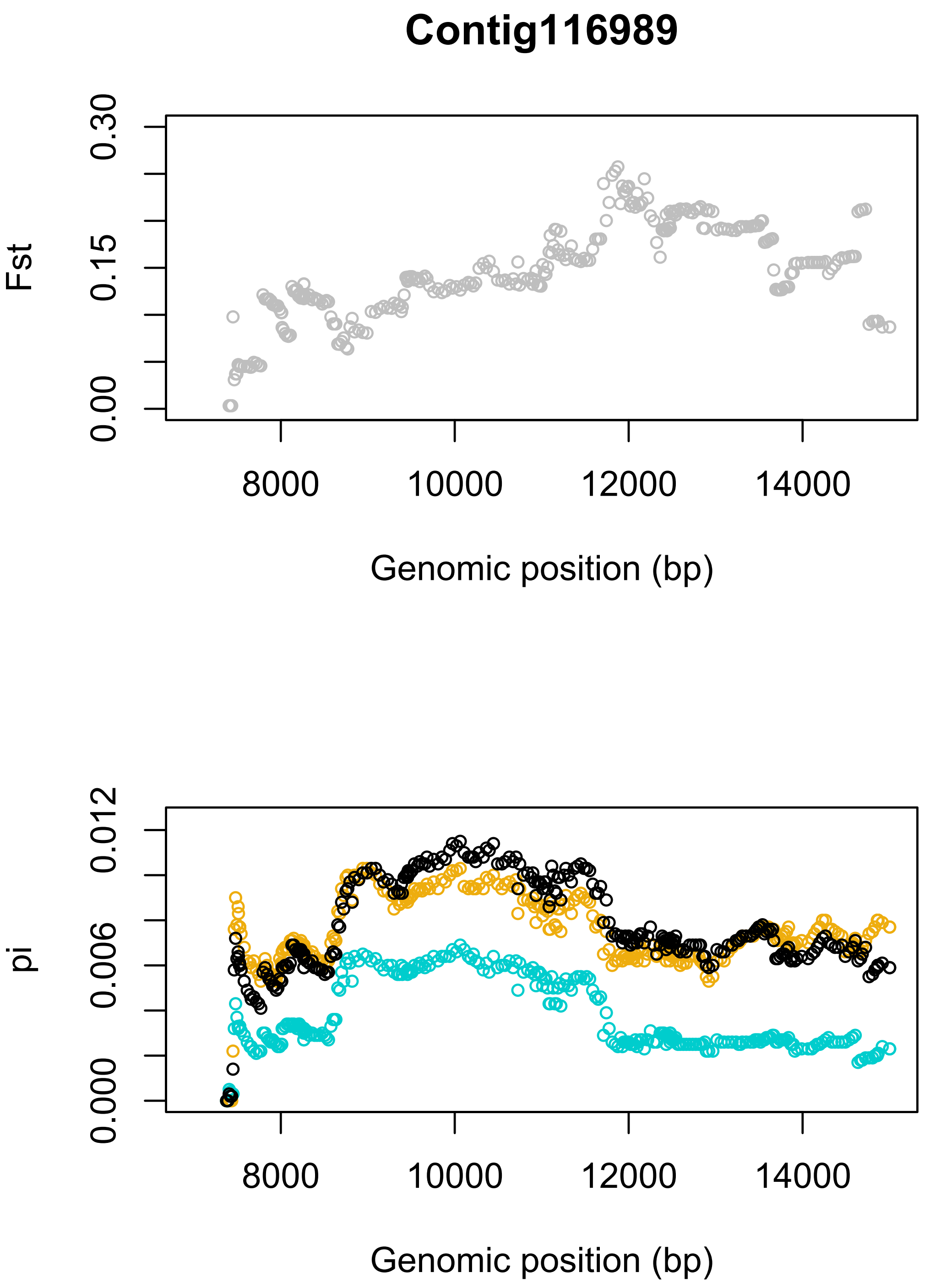
Contig116989
Fst
0.00
0.15
0.30
8000
10000
12000
14000
Genomic position (bp)
pi
0.000
0.006
0.012
8000
10000
12000
14000
Genomic position (bp)

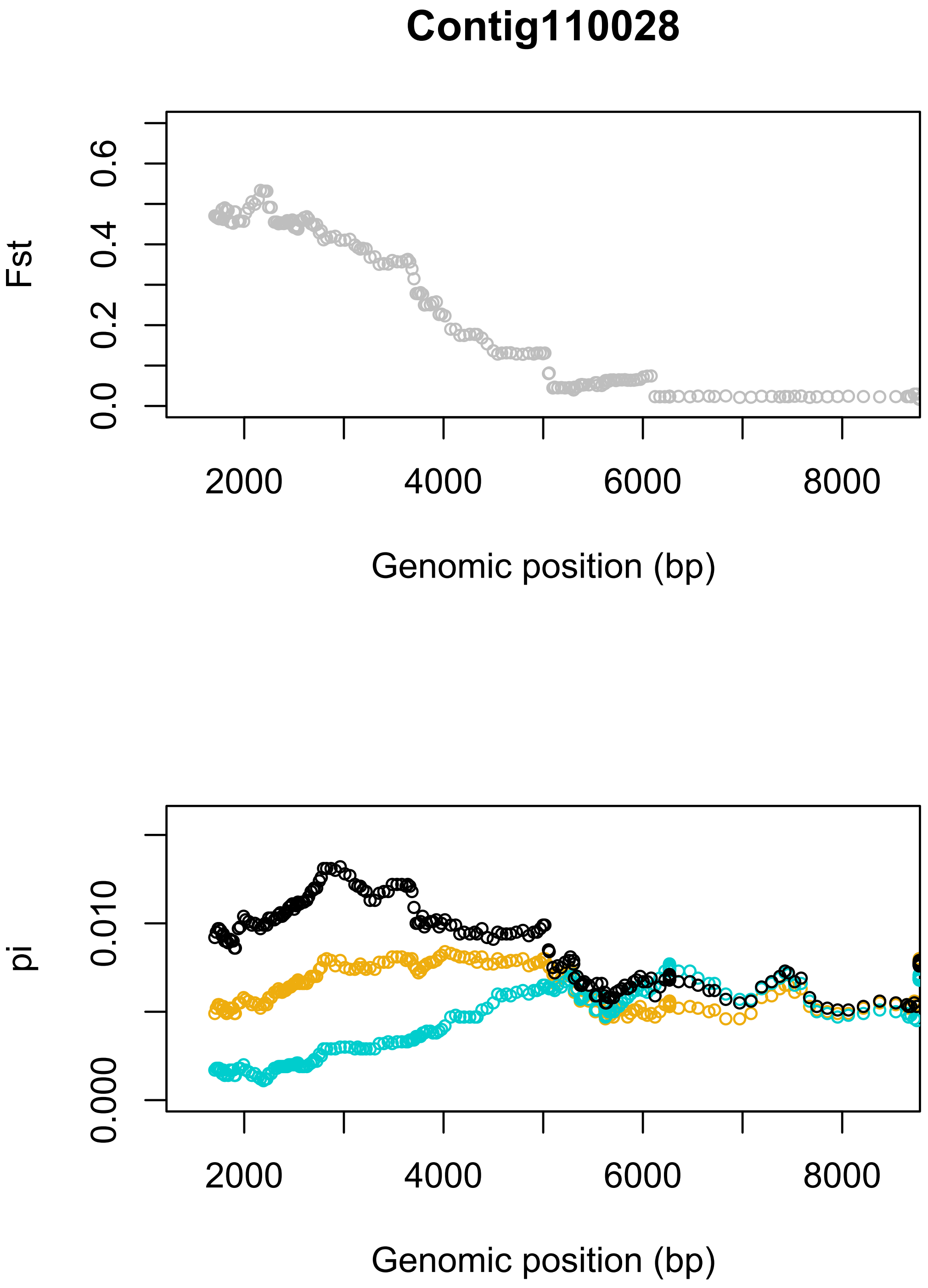
Contig110028
Fst
0.0
0.2
0.4
0.6
2000
4000
6000
8000
Genomic position (bp)
pi
0.000
0.010
2000
4000
6000
8000
Genomic position (bp)

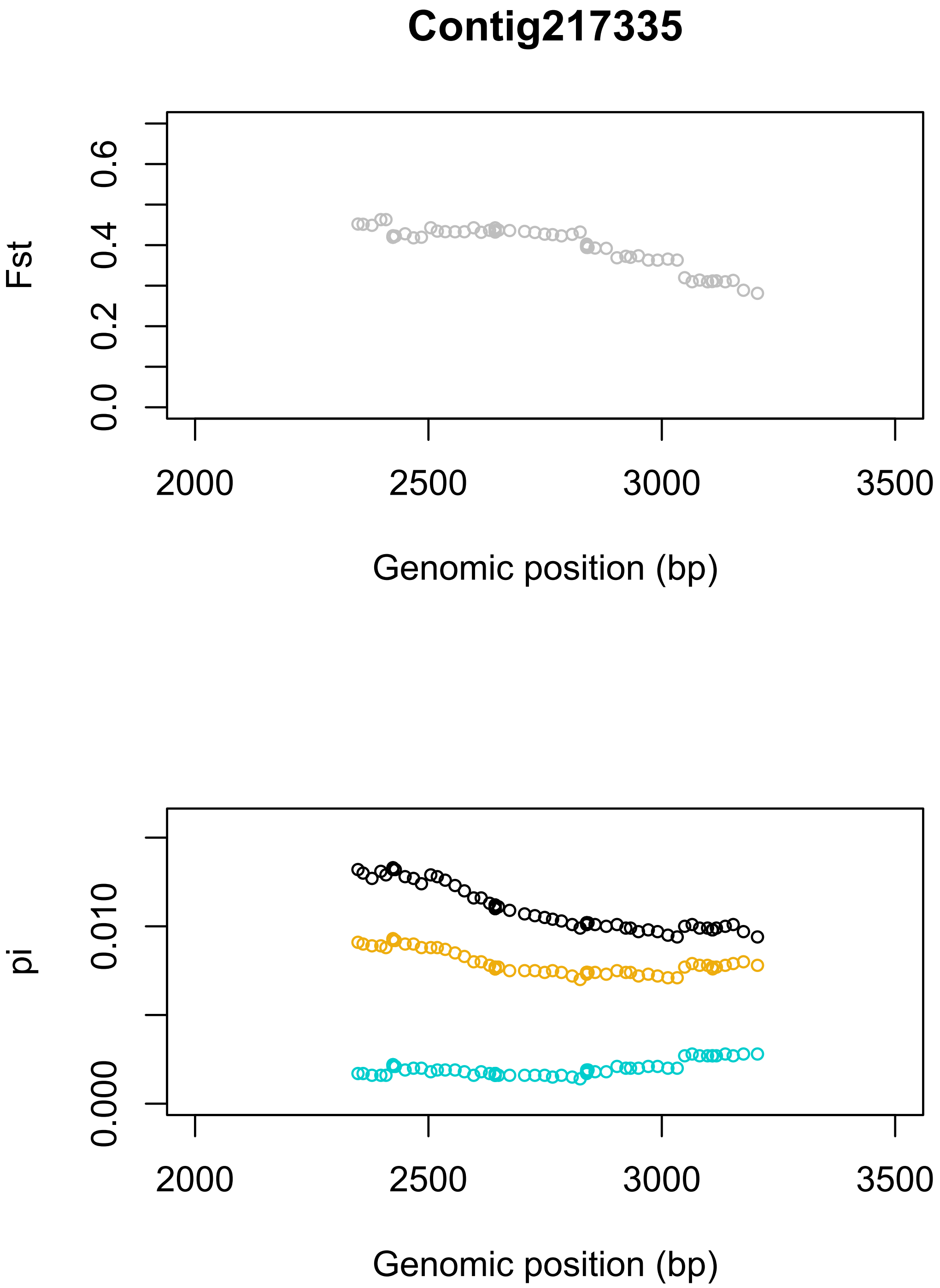

Contig217335
Fst
0.0
0.2
0.4
0.6
2000
2500
3000
3500
Genomic position (bp)
pi
0.000
0.010
2000
2500
3000
3500
Genomic position (bp)

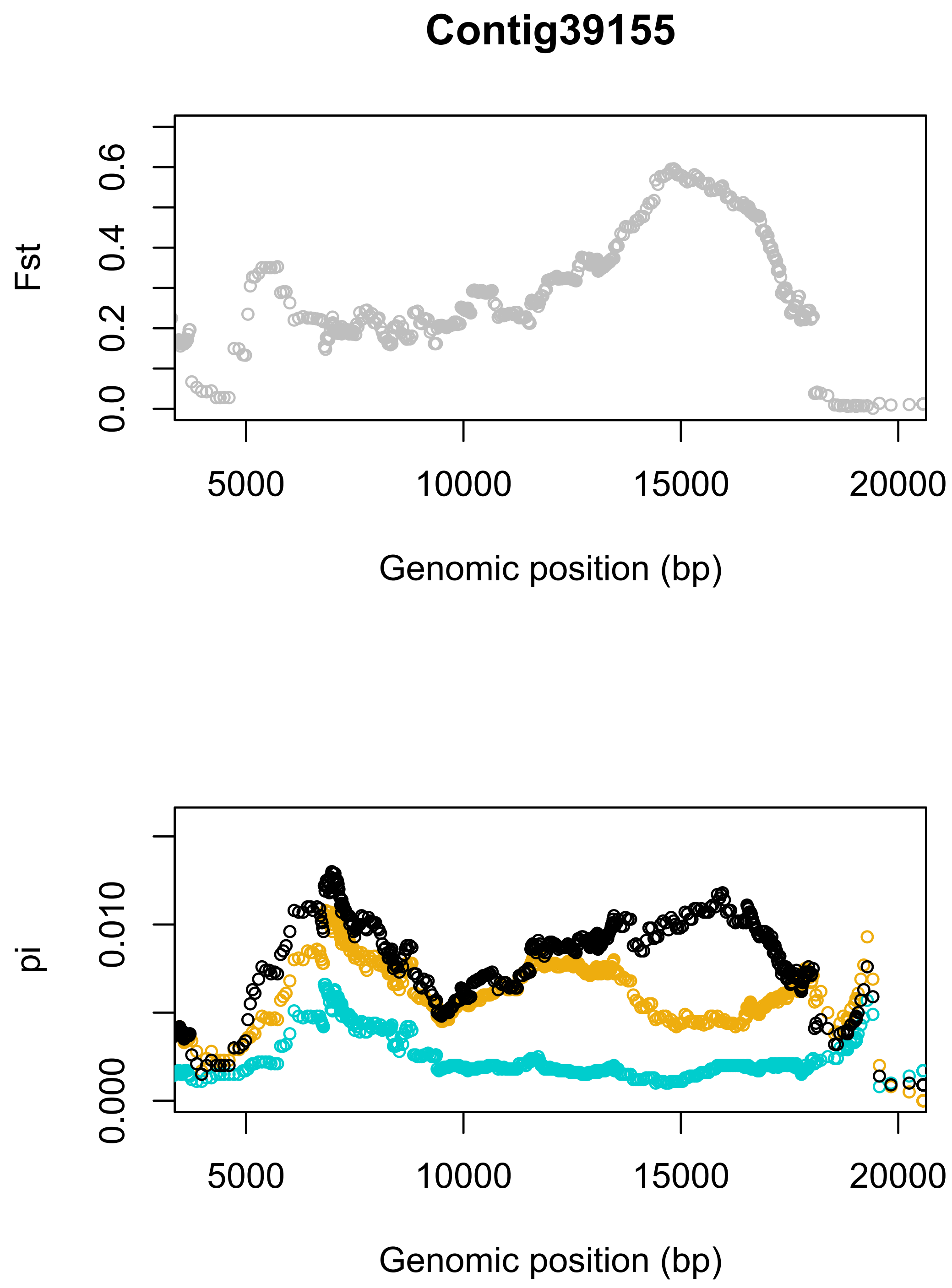

Contig39155
Fst
0.0
0.2
0.4
0.6
5000
10000
15000
20000
Genomic position (bp)
pi
0.000
0.010
5000
10000
15000
20000
Genomic position (bp)

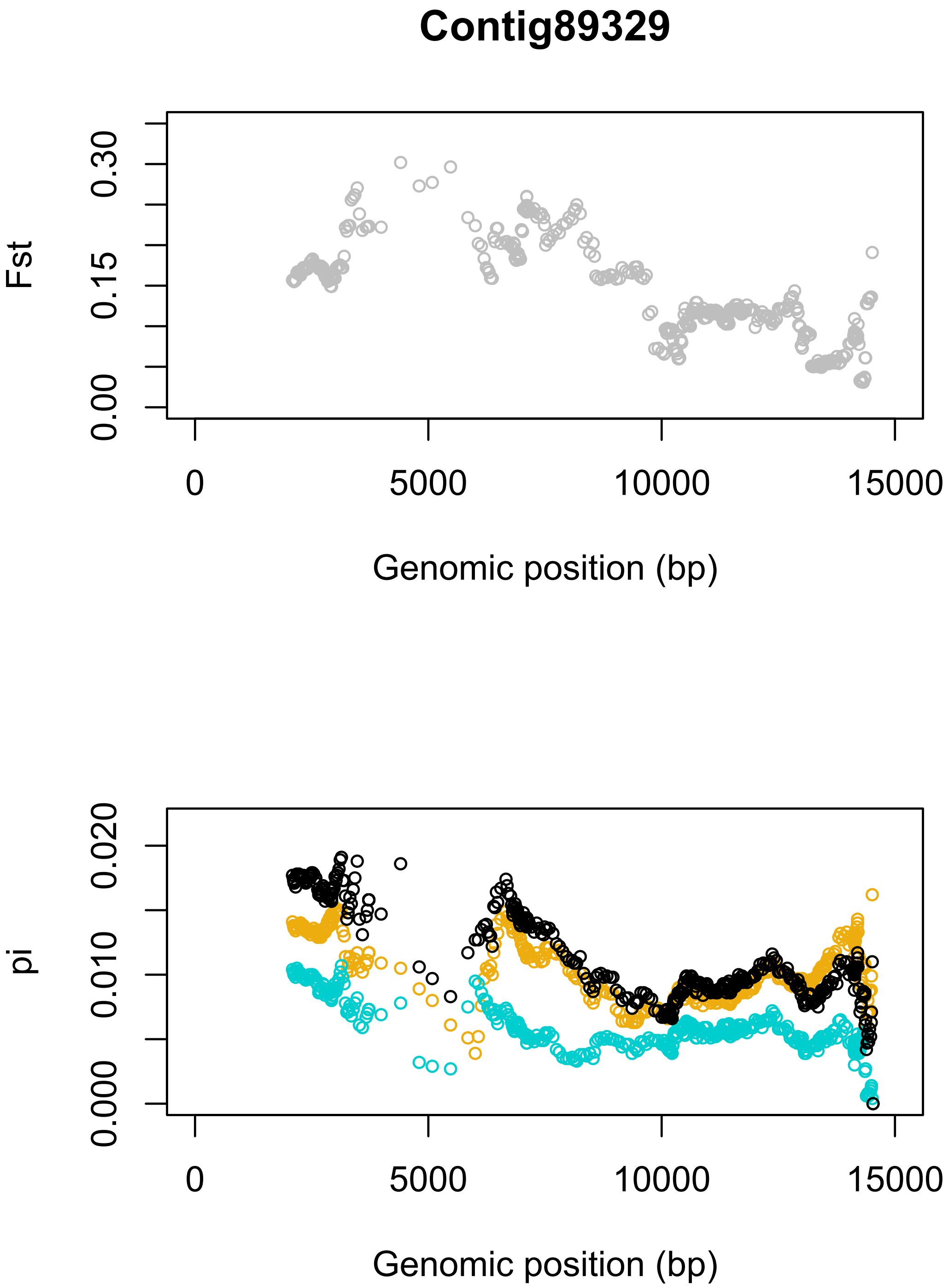
Contig89329
Fst
0.00
0.15
0.30
0
5000
10000
15000
Genomic position (bp)
pi
0.000
0.010
0.020
0
5000
10000
15000
Genomic position (bp)

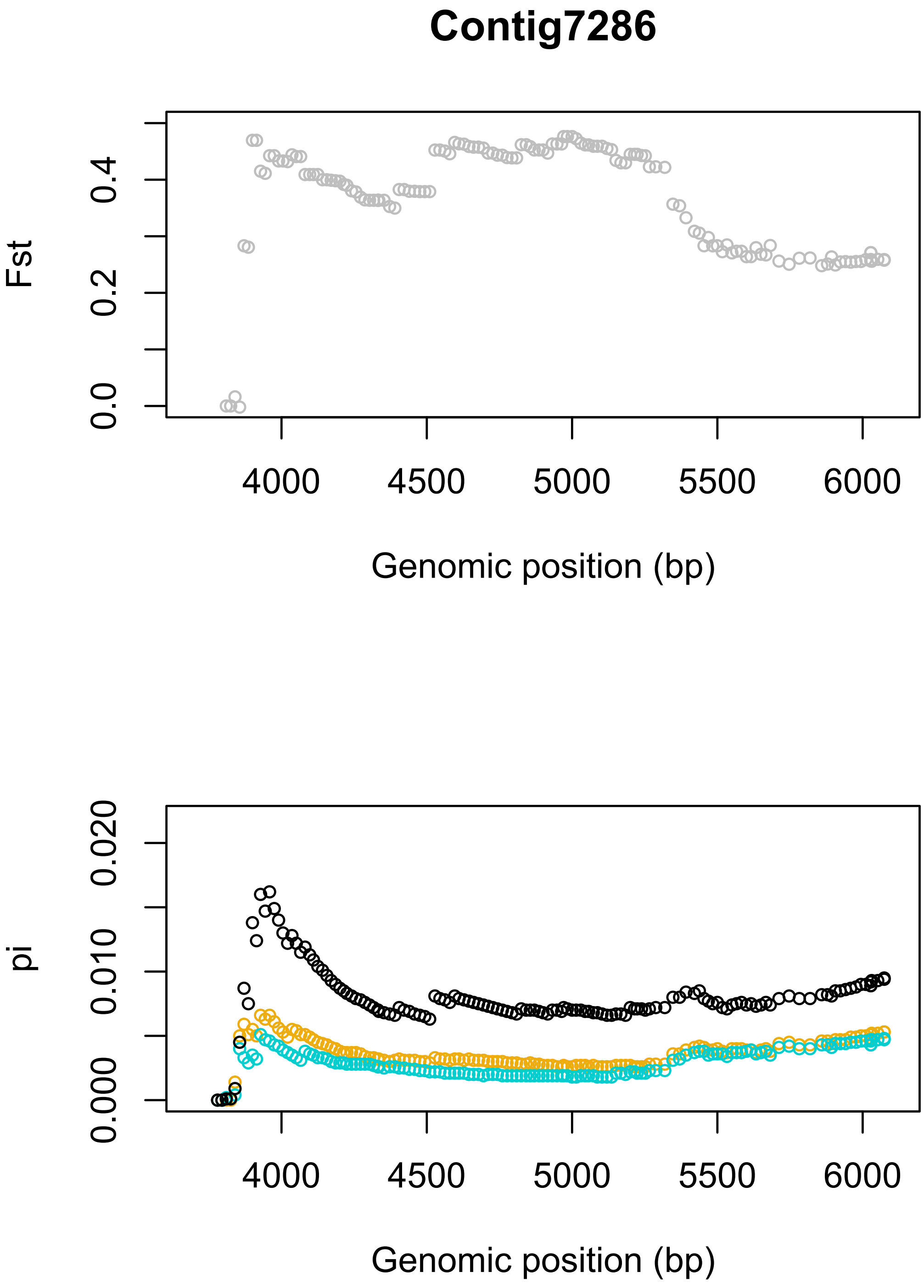
Contig7286
Fst
0.0
0.2
0.4
4000
4500
5000
5500
6000
Genomic position (bp)
pi
0.000
0.010
0.020
4000
4500
5000
5500
6000
Genomic position (bp)

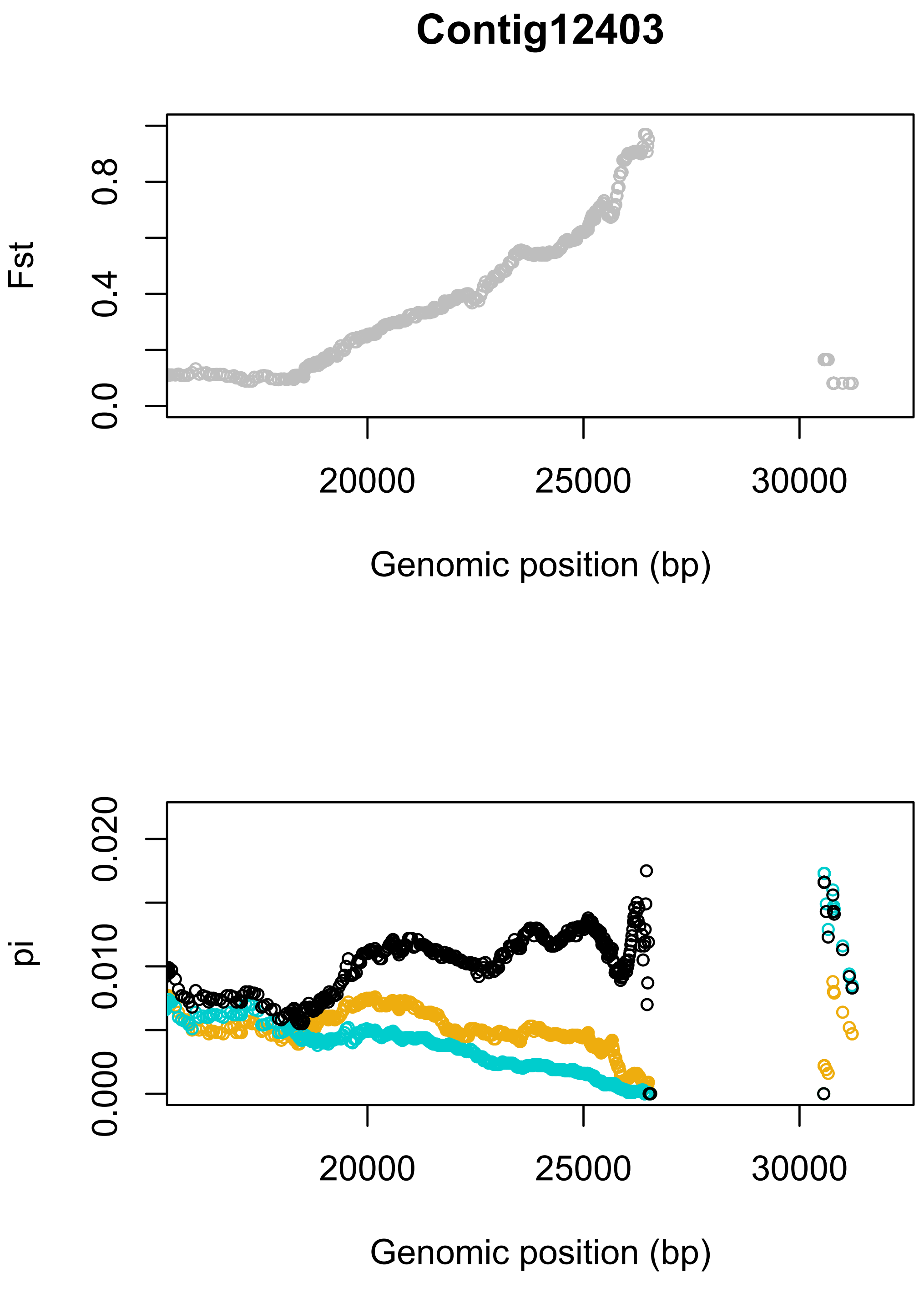
Contig12403
Fst
0.0
0.4
0.8
20000
25000
30000
Genomic position (bp)
pi
0.000
0.010
0.020
20000
25000
30000
Genomic position (bp)

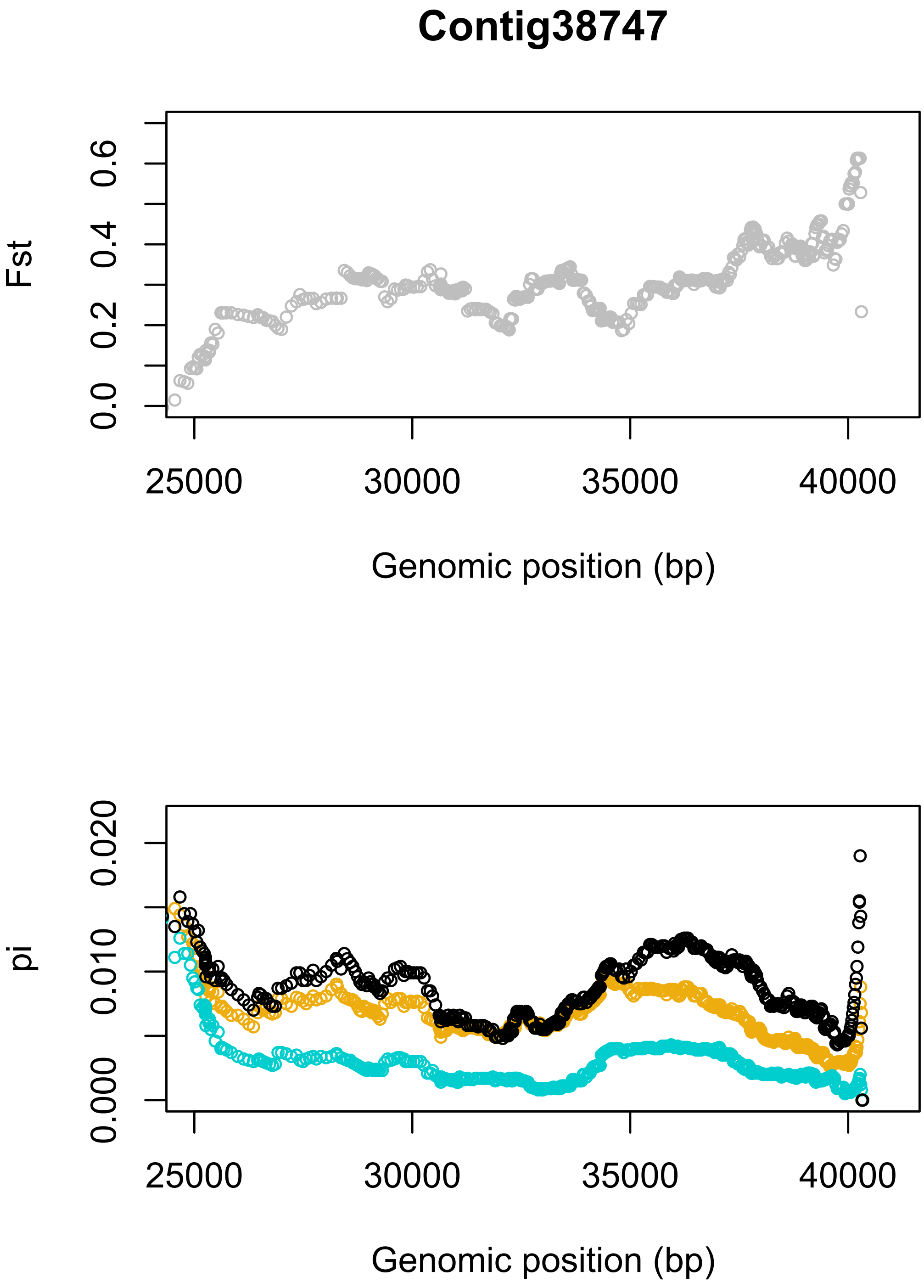
Contig38747
Fst
0.0
0.2
0.4
0.6
25000
30000
35000
40000
Genomic position (bp)
pi
0.000
0.010
0.020
25000
30000
35000
40000
Genomic position (bp)

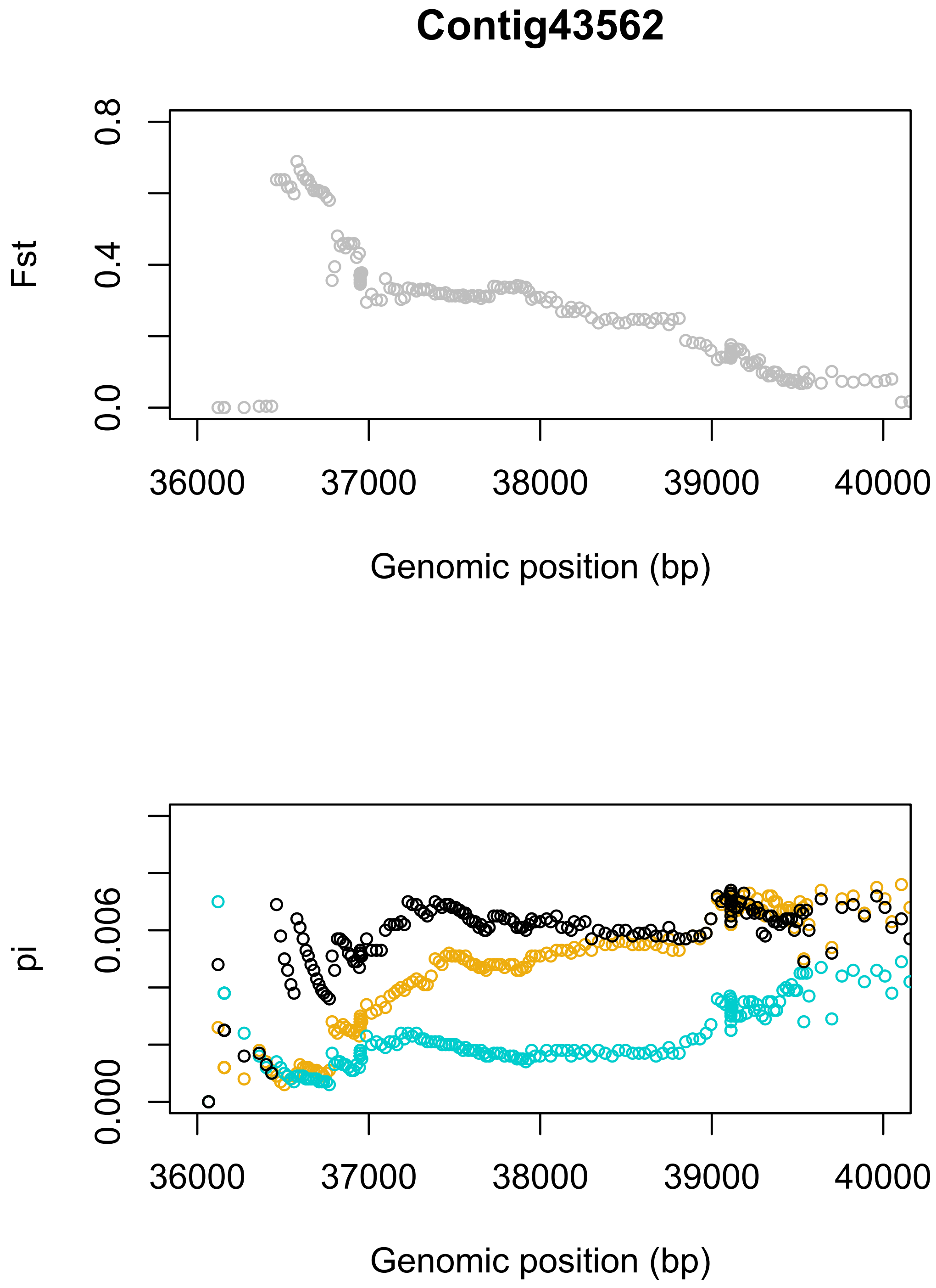
Contig43562
Fst
0.0
0.4
0.8
36000
37000
38000
39000
40000
Genomic position (bp)
pi
0.000
0.006
36000
37000
38000
39000
40000
Genomic position (bp)

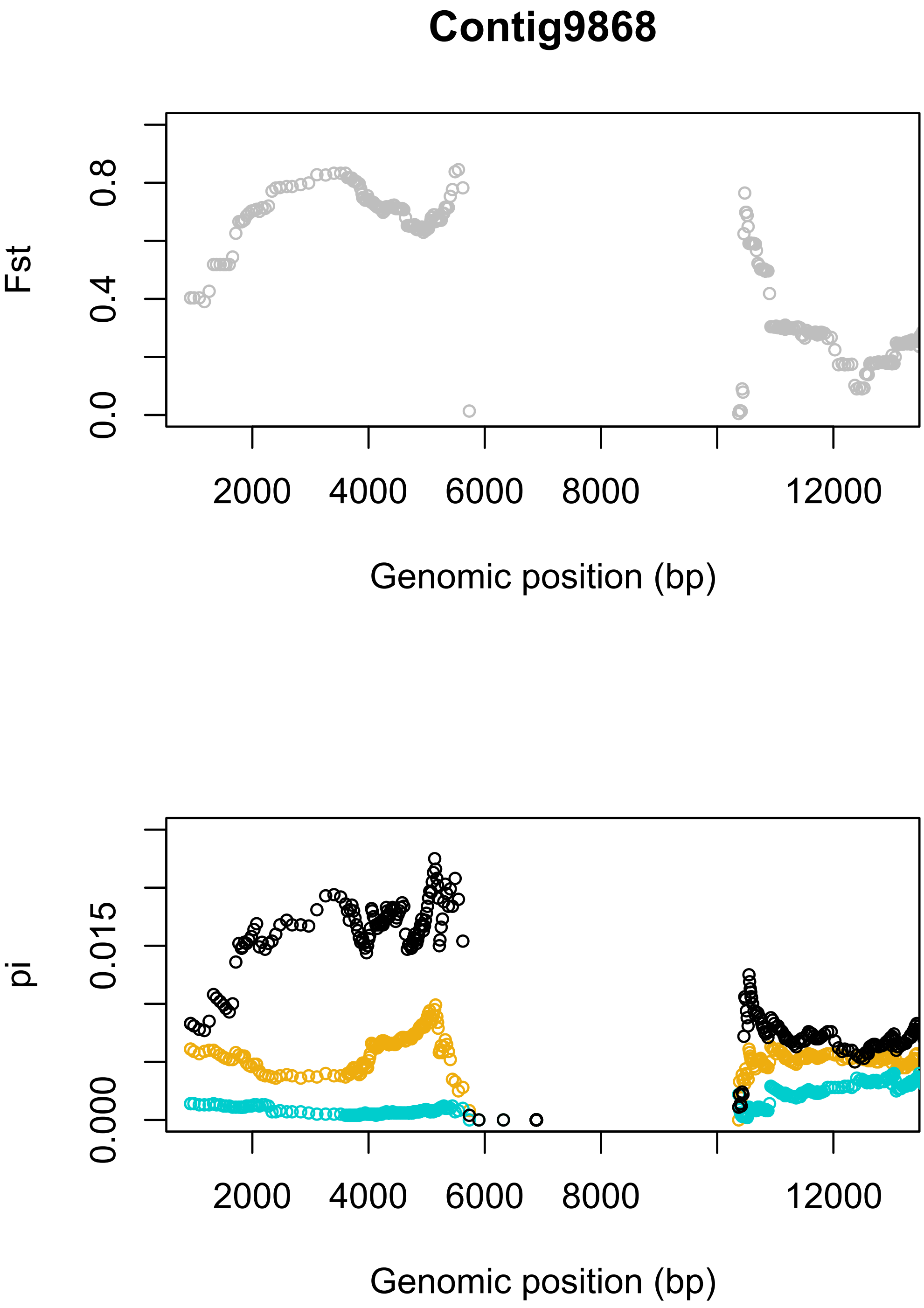
Contig9868
Fst
0.0
0.4
0.8
2000
4000
6000
8000
12000
Genomic position (bp)
pi
0.000
0.015
2000
4000
6000
8000
12000
Genomic position (bp)

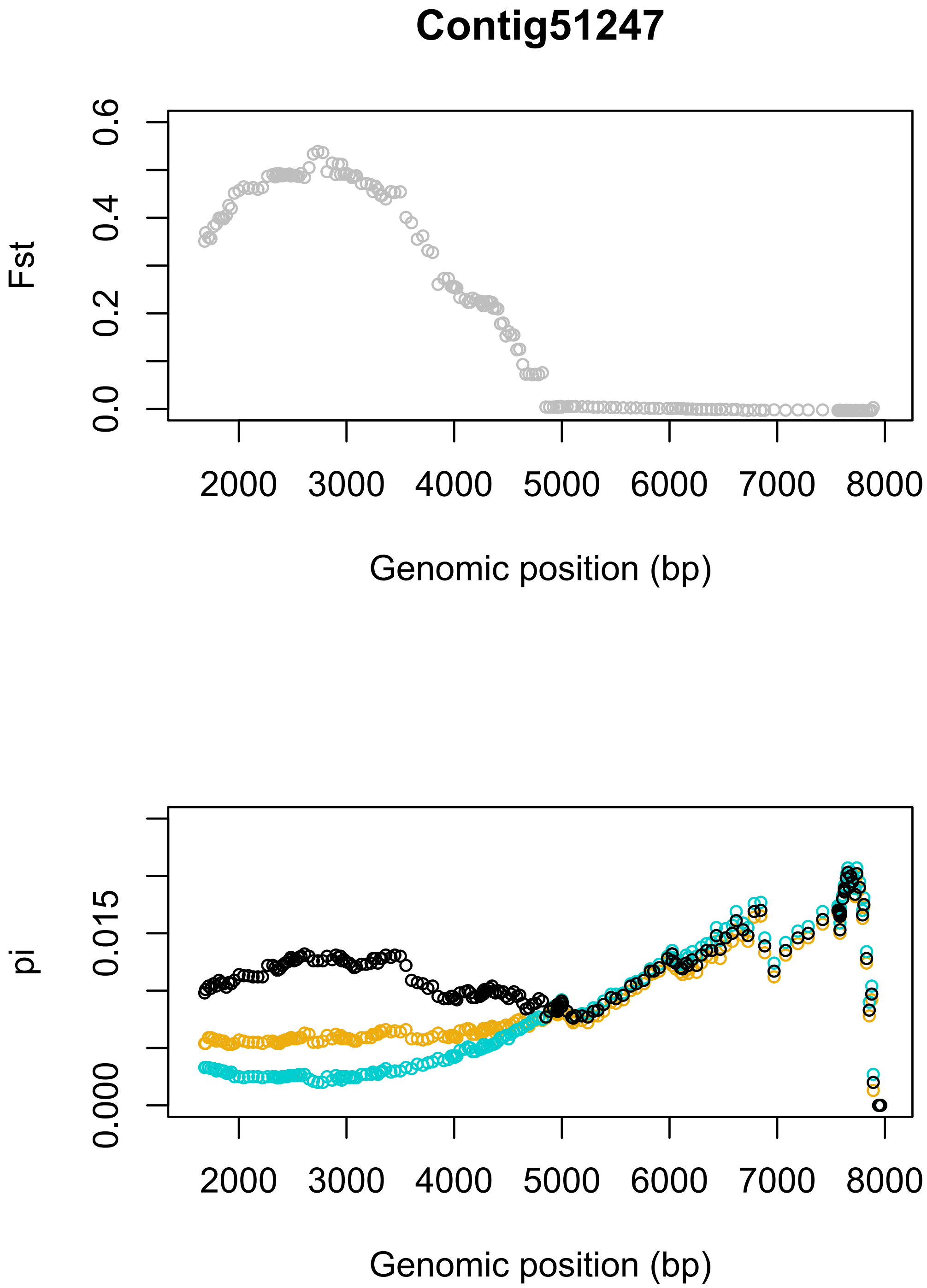
Contig51247
Fst
0.0
0.2
0.4
0.6
2000
3000
4000
5000
6000
7000
8000
Genomic position (bp)
pi
0.000
0.015
2000
3000
4000
5000
6000
7000
8000
Genomic position (bp)

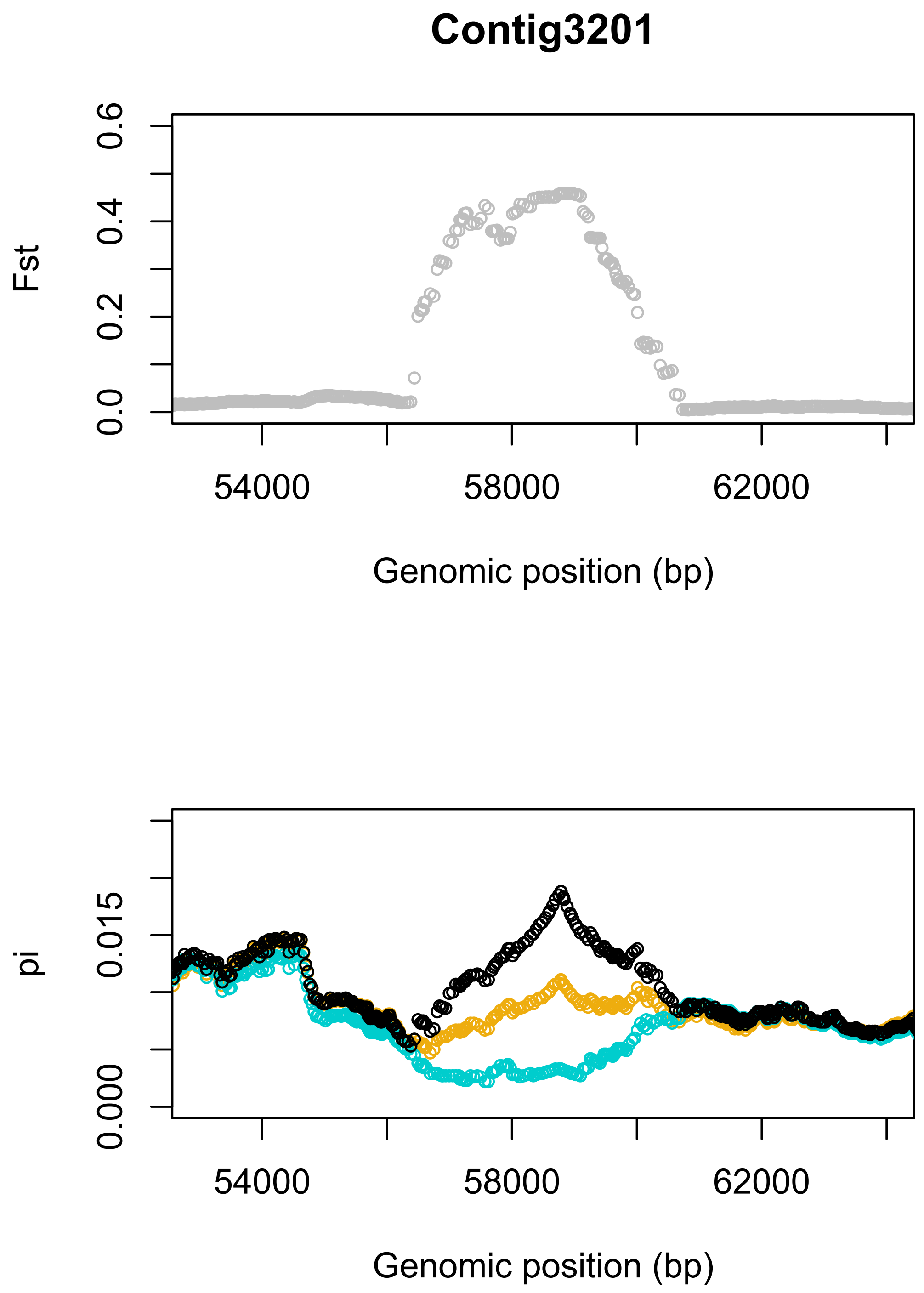
Contig3201
Fst
0.0
0.2
0.4
0.6
54000
58000
62000
Genomic position (bp)
pi
0.000
0.015
54000
58000
62000
Genomic position (bp)

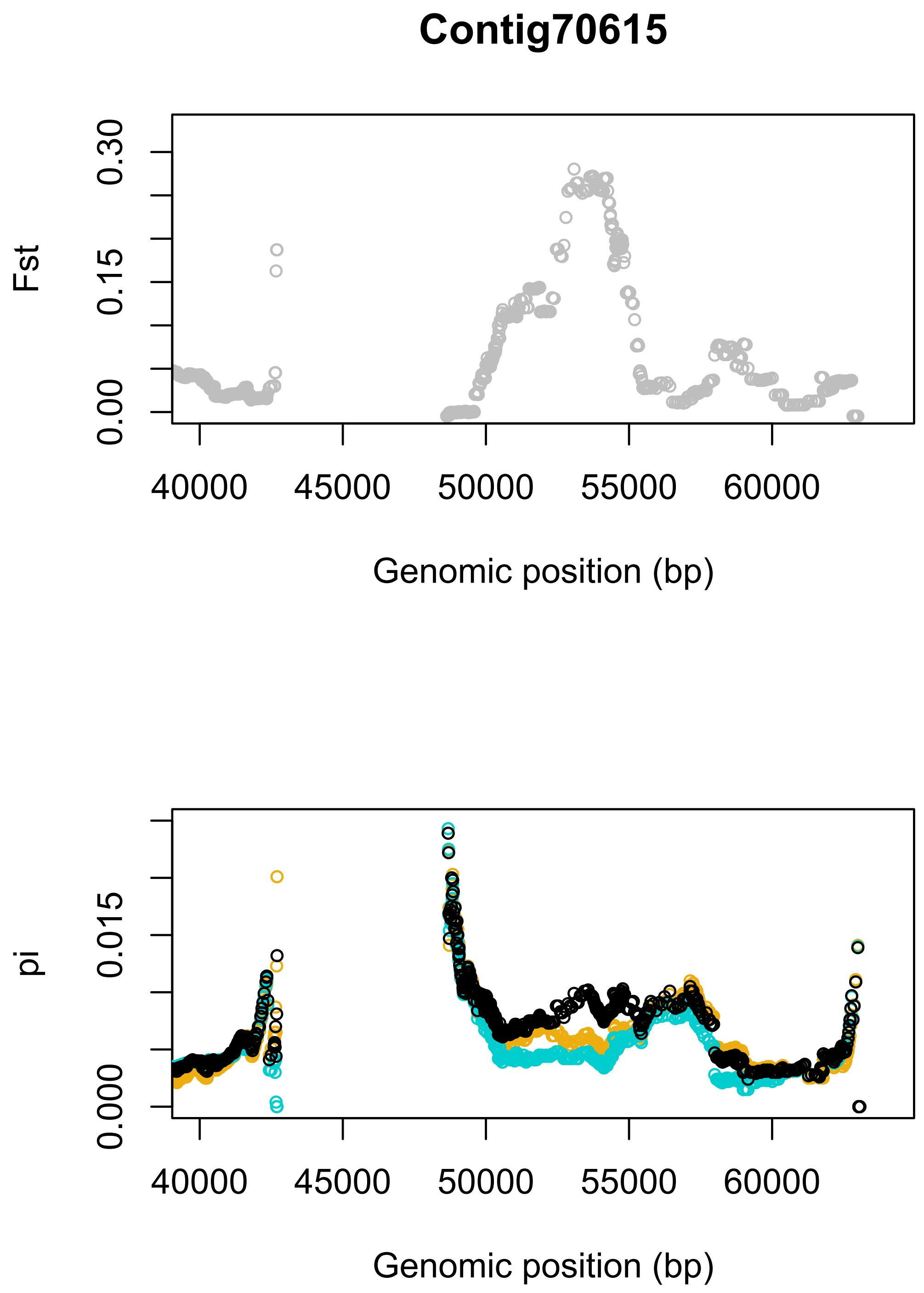

Contig70615
Fst
0.00
0.15
0.30
40000
45000
50000
55000
60000
Genomic position (bp)
pi
0.000
0.015
40000
45000
50000
55000
60000
Genomic position (bp)

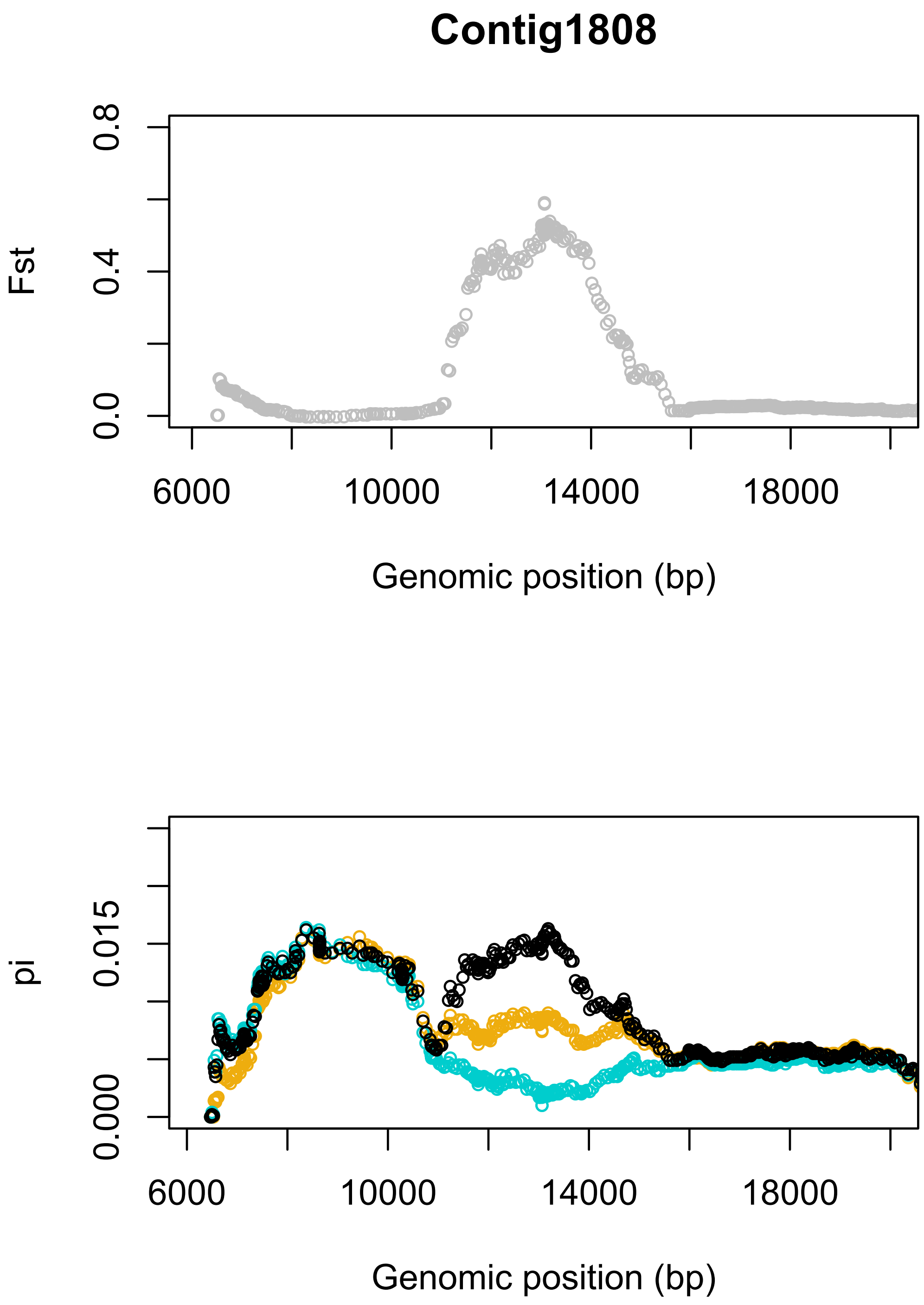
Contig1808
Fst
0.0
0.4
0.8
6000
10000
14000
18000
Genomic position (bp)
pi
0.000
0.015
6000
10000
14000
18000
Genomic position (bp)

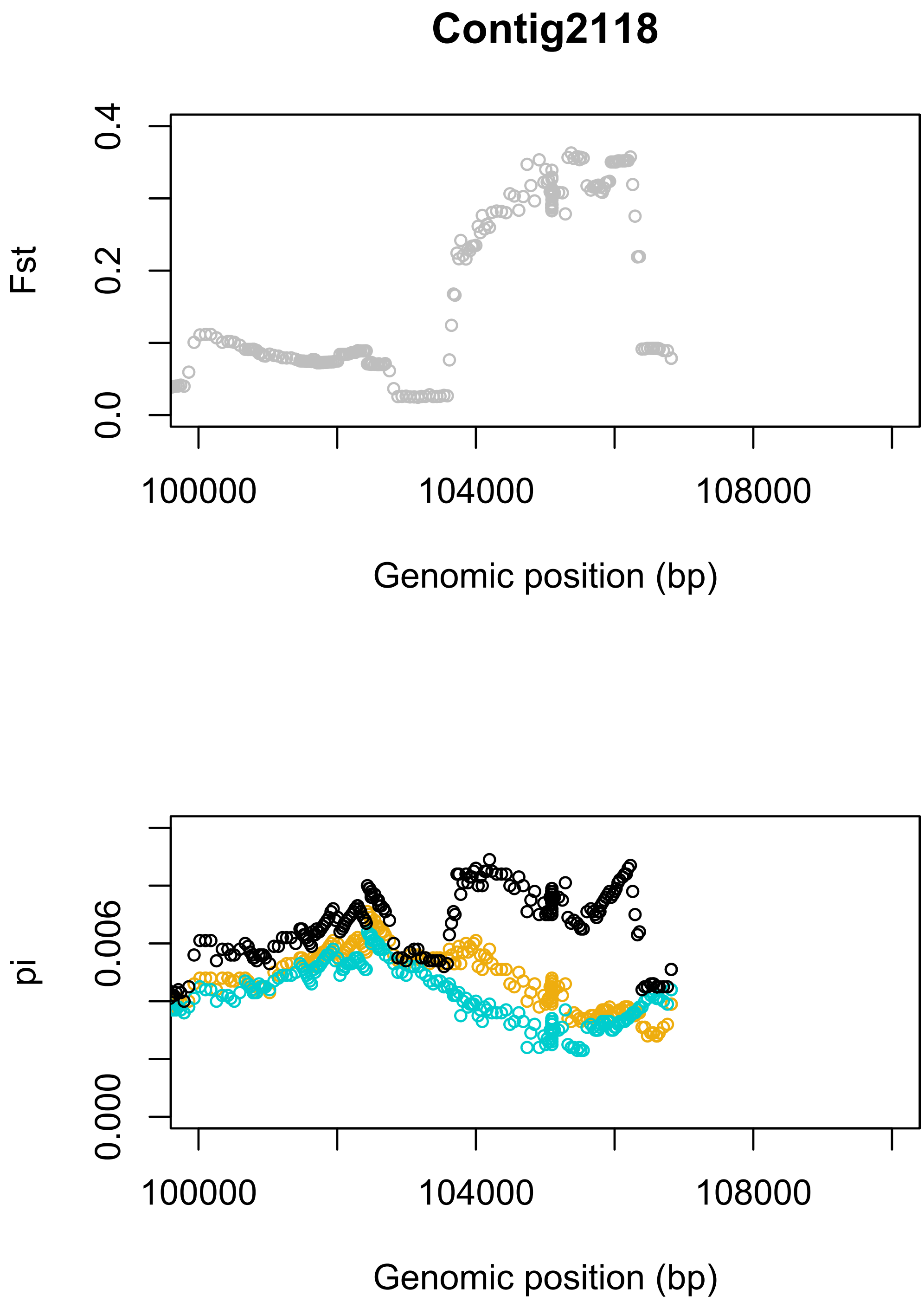
Contig2118
Fst
0.0
0.2
0.4
100000
104000
108000
Genomic position (bp)
pi
0.000
0.006
100000
104000
108000
Genomic position (bp)

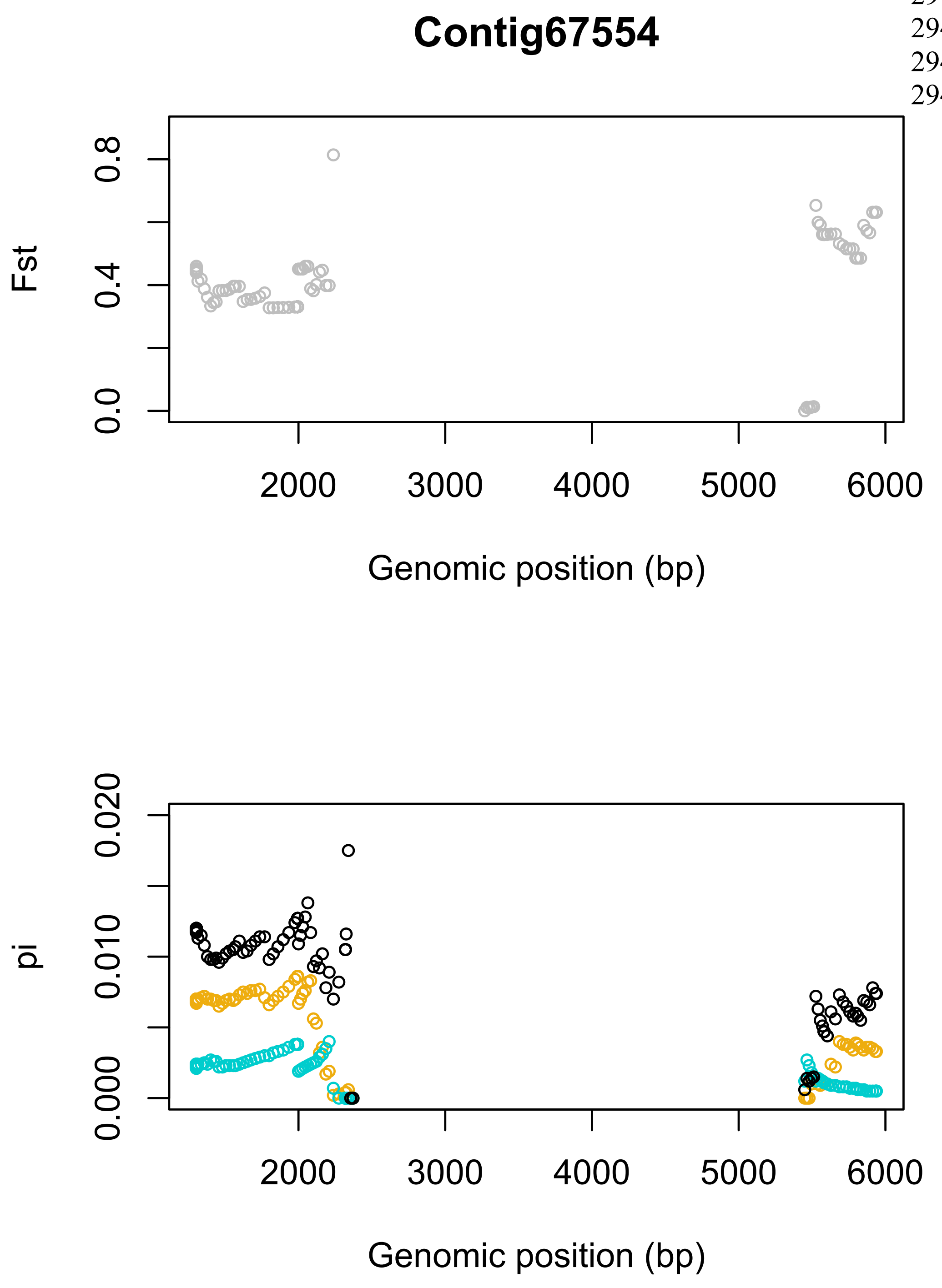
Contig67554
Fst
0.0
0.4
0.8
2000
3000
4000
5000
6000
Genomic position (bp)
pi
0.000
0.010
0.020
2000
3000
4000
5000
6000
Genomic position (bp)

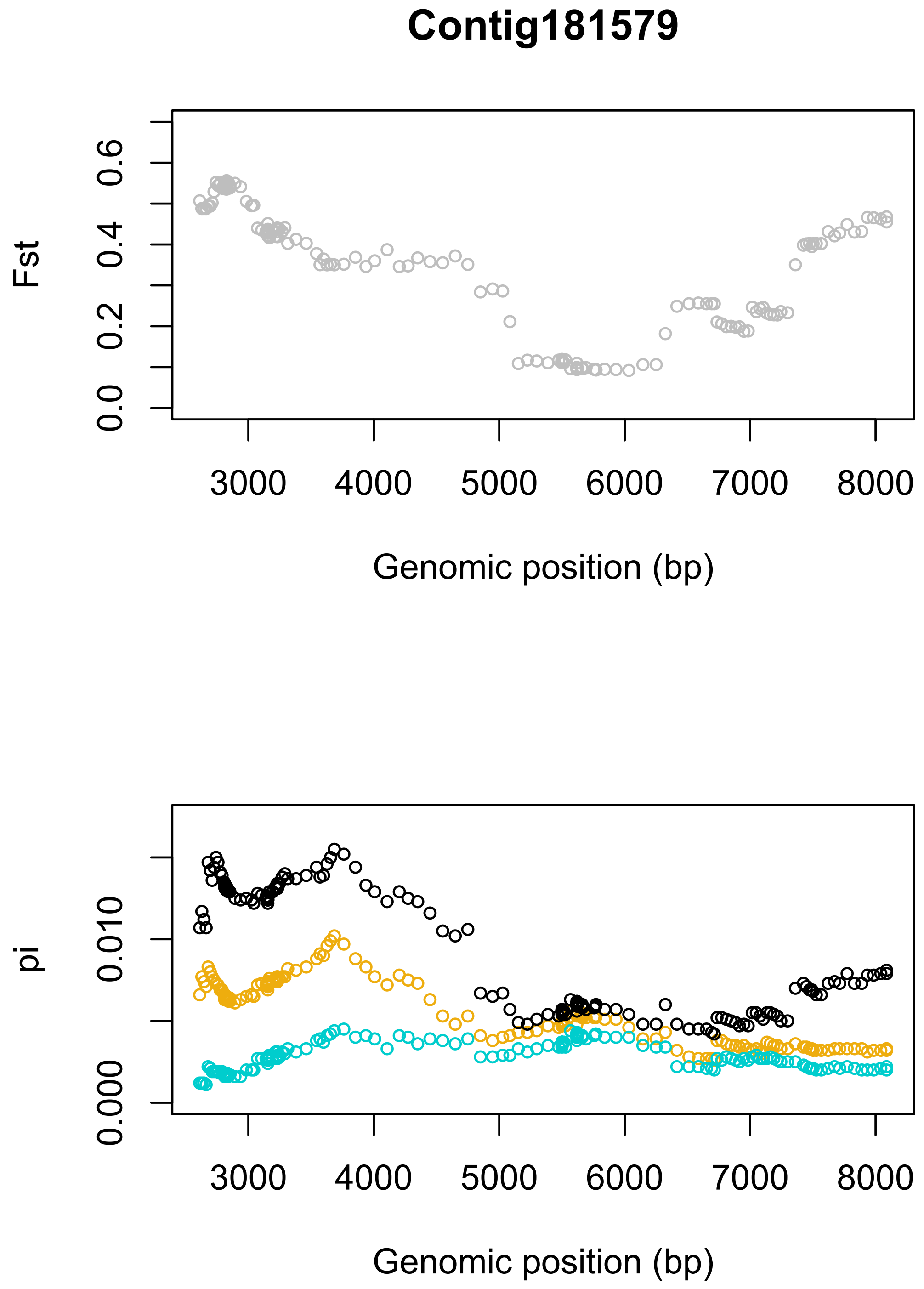

Contig181579
Fst
0.0
0.2
0.4
0.6
3000
4000
5000
6000
7000
8000
Genomic position (bp)
pi
0.000
0.010
3000
4000
5000
6000
7000
8000
Genomic position (bp)

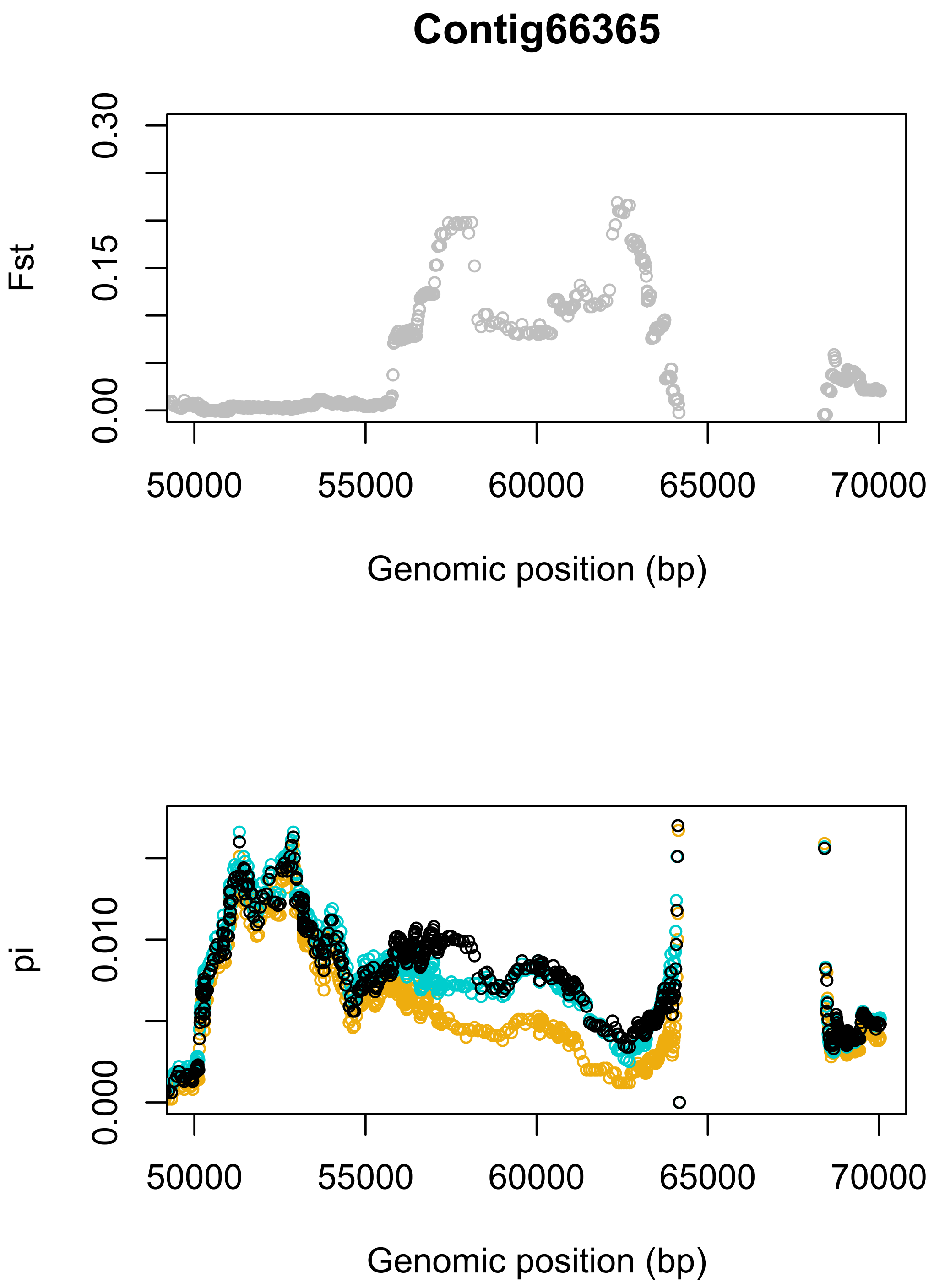
Contig66365
Fst
0.00
0.15
0.30
50000
55000
60000
65000
70000
Genomic position (bp)
pi
0.000
0.010
50000
55000
60000
65000
70000
Genomic position (bp)

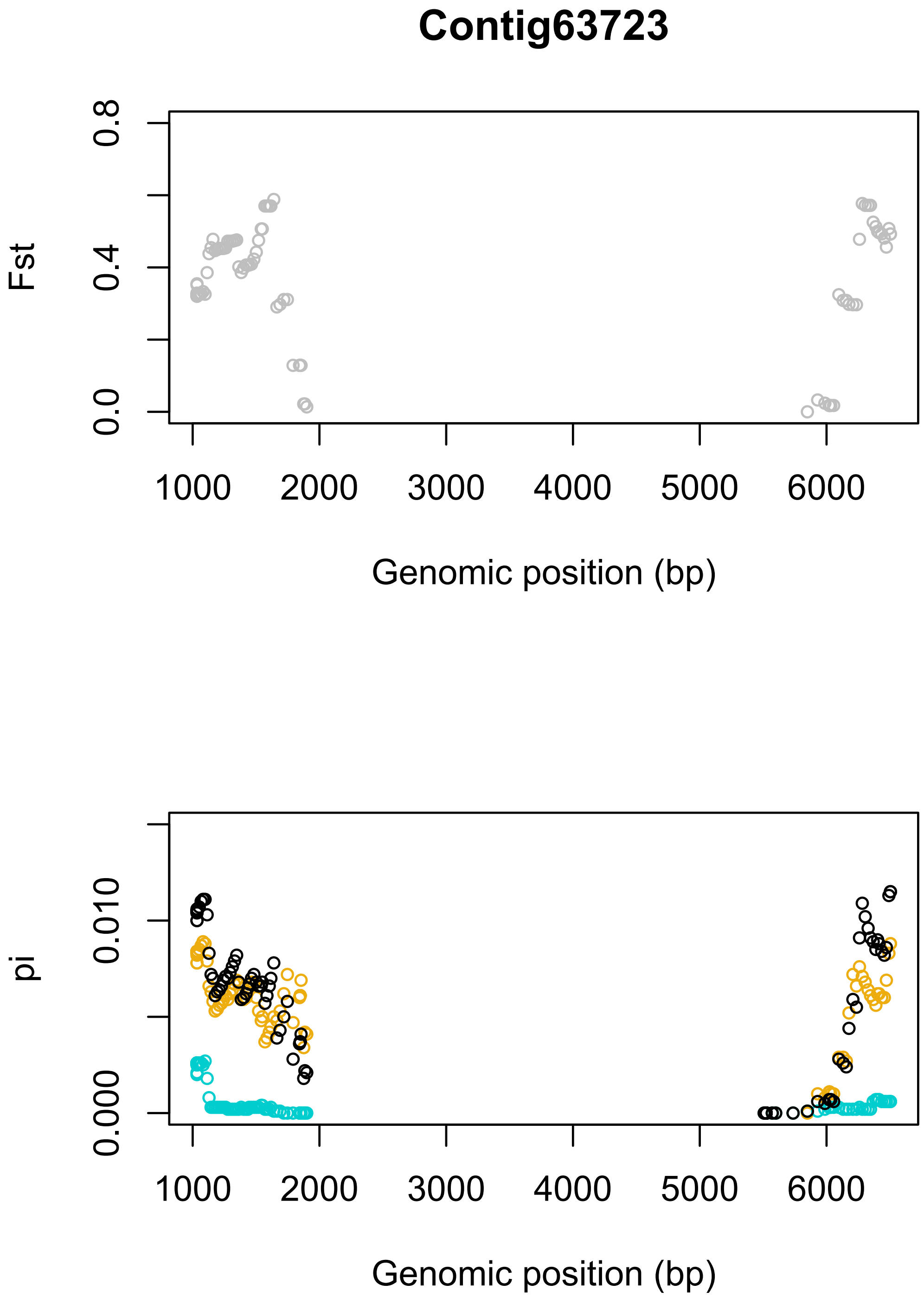
Contig63723
Fst
0.0
0.4
0.8
1000
2000
3000
4000
5000
6000
Genomic position (bp)
pi
0.000
0.010
1000
2000
3000
4000
5000
6000
Genomic position (bp)

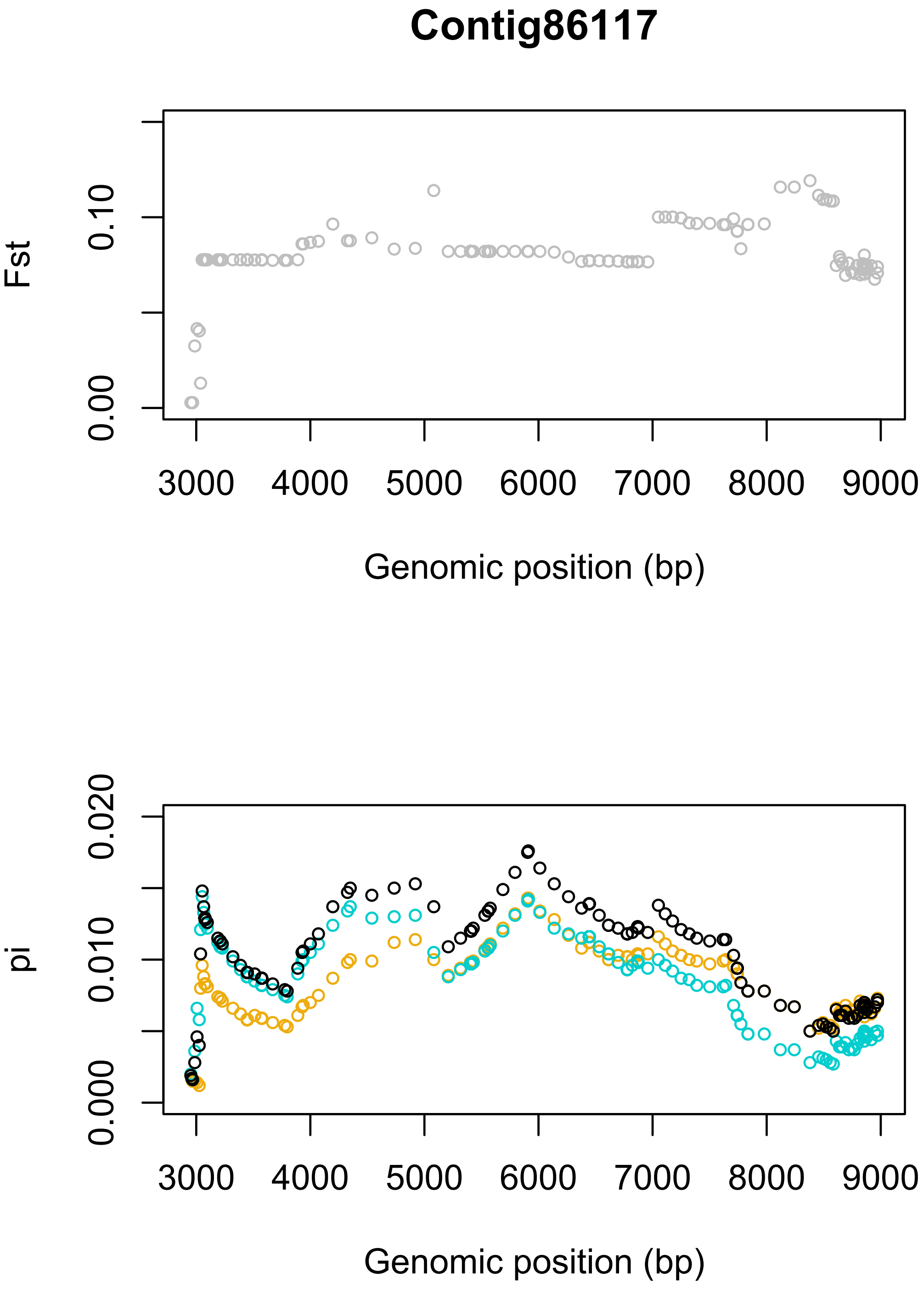
Contig86117
Fst
0.00
0.10
3000 4000 5000 6000 7000 8000 9000
Genomic position (bp)
pi
0.000
0.010
0.020
3000 4000 5000 6000 7000 8000 9000
Genomic position (bp)

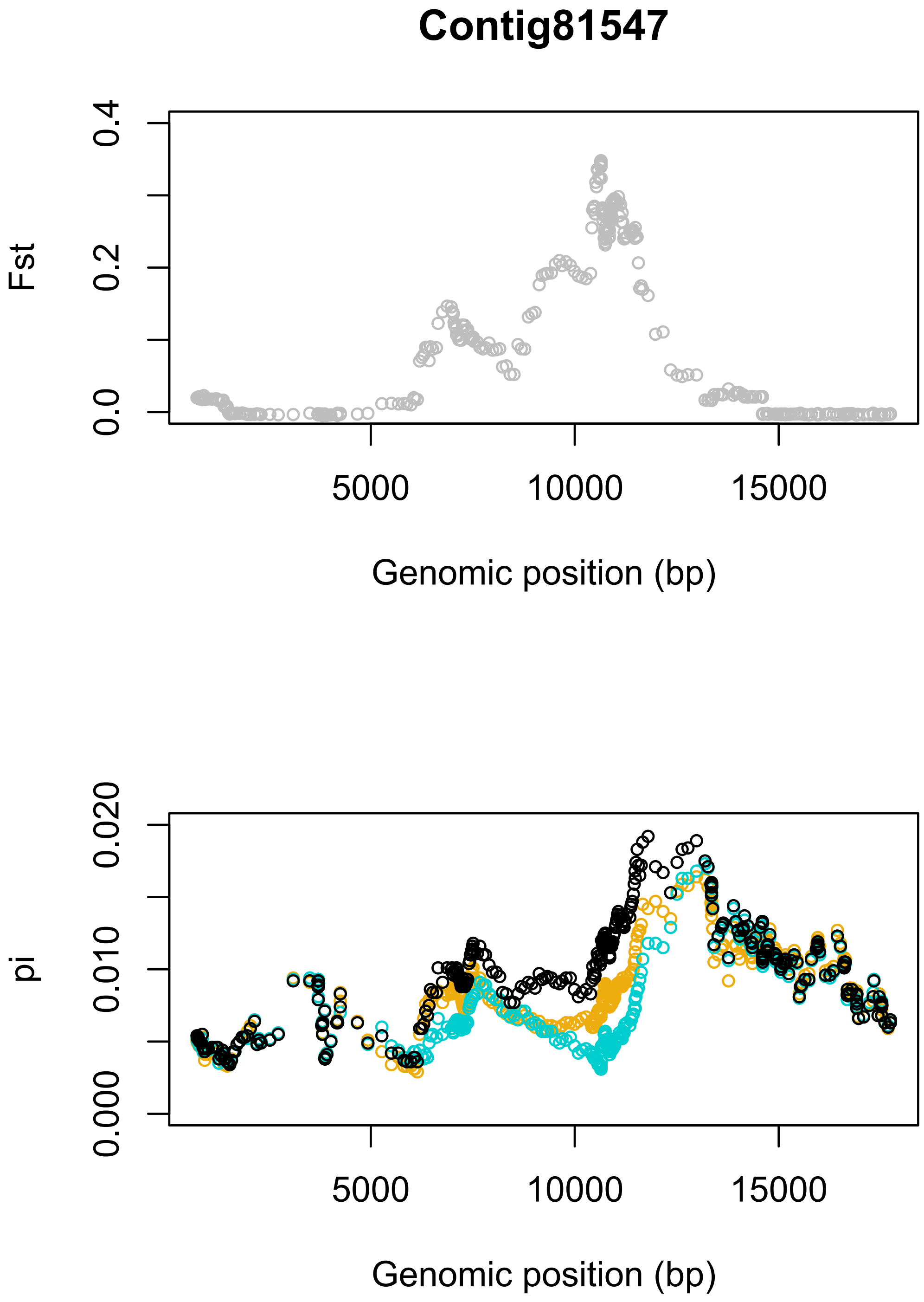
Contig81547
Fst
0.0
0.2
0.4
5000
10000
15000
Genomic position (bp)
pi
0.000
0.010
0.020
5000
10000
15000
Genomic position (bp)

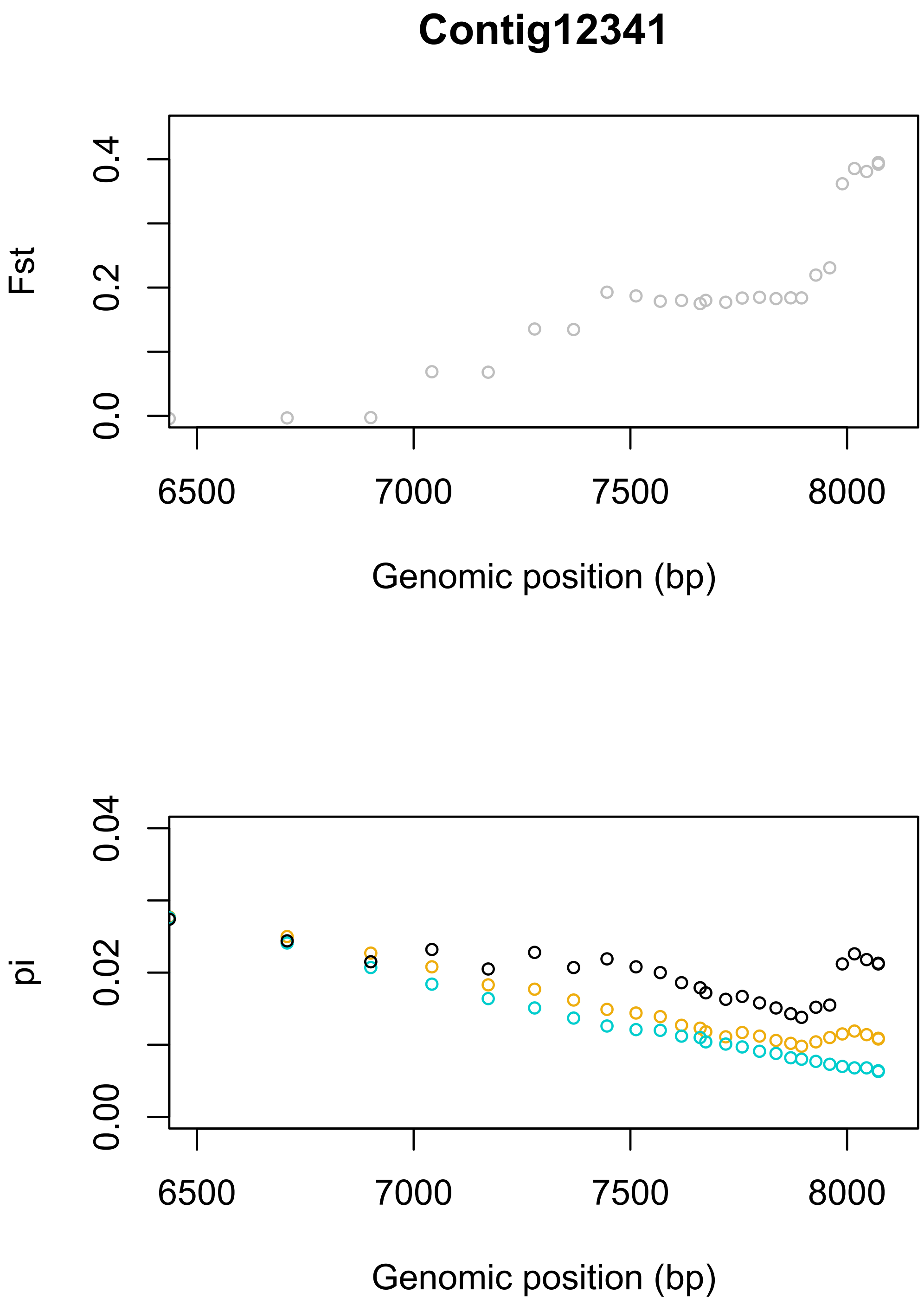
Contig12341
Fst
0.0
0.2
0.4
6500
7000
7500
8000
Genomic position (bp)
pi
0.00
0.02
0.04
6500
7000
7500
8000
Genomic position (bp)

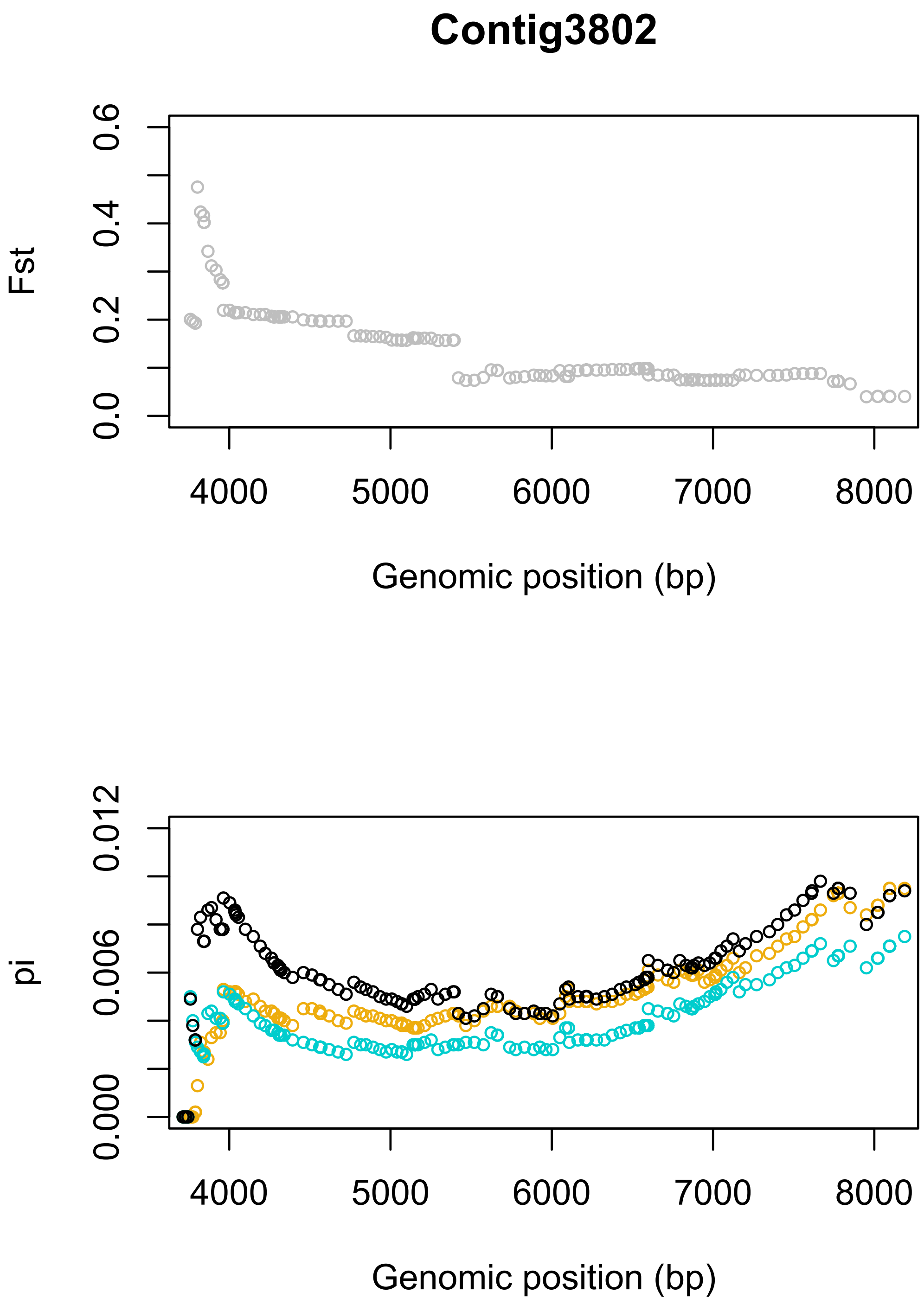
Contig3802
Fst
0.0
0.2
0.4
0.6
4000
5000
6000
7000
8000
Genomic position (bp)
pi
0.000
0.006
0.012
4000
5000
6000
7000
8000
Genomic position (bp)

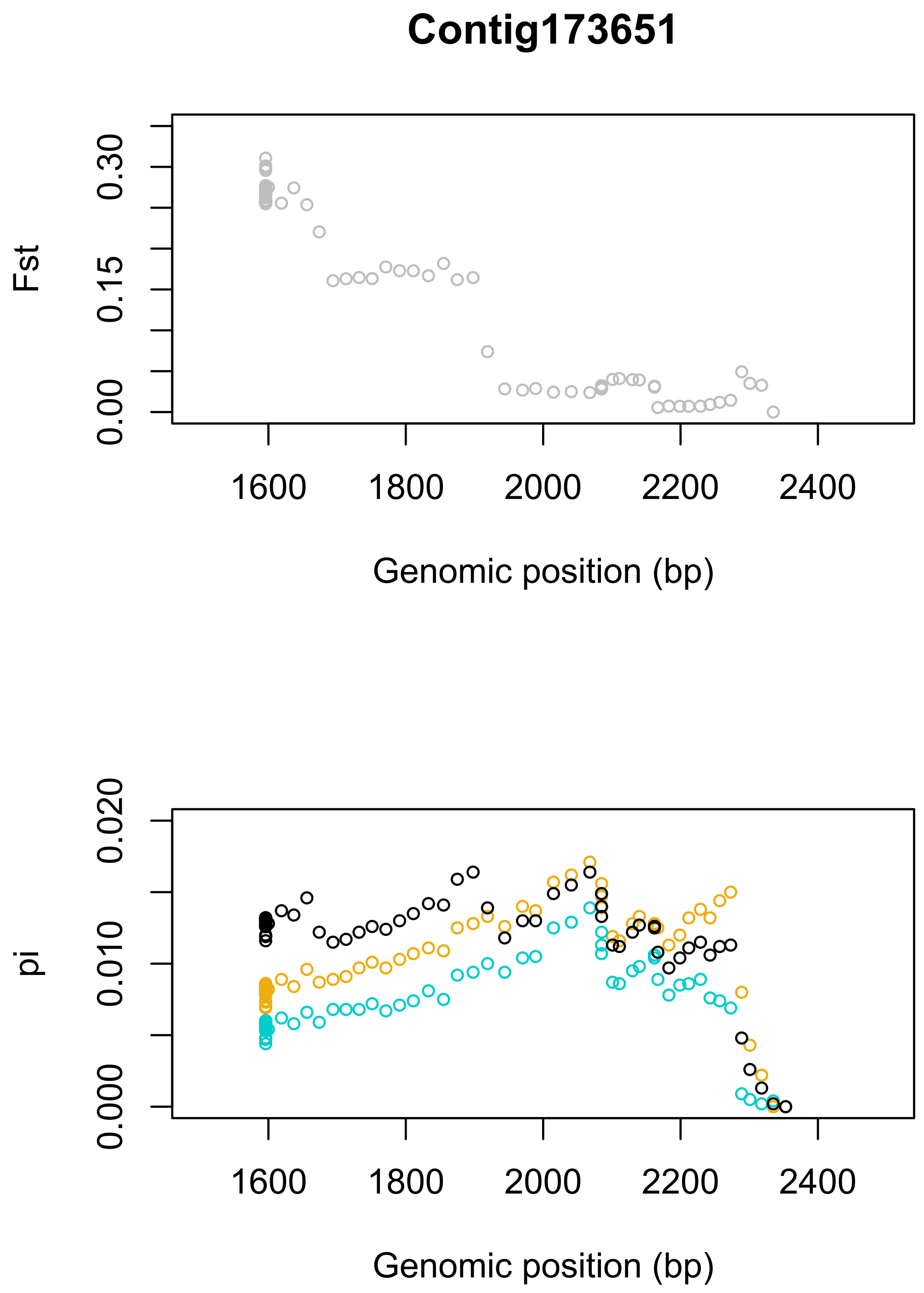
Contig173651
Fst
0.00
0.15
0.30
1600
1800
2000
2200
2400
Genomic position (bp)
pi
0.000
0.010
0.020
1600
1800
2000
2200
2400
Genomic position (bp)

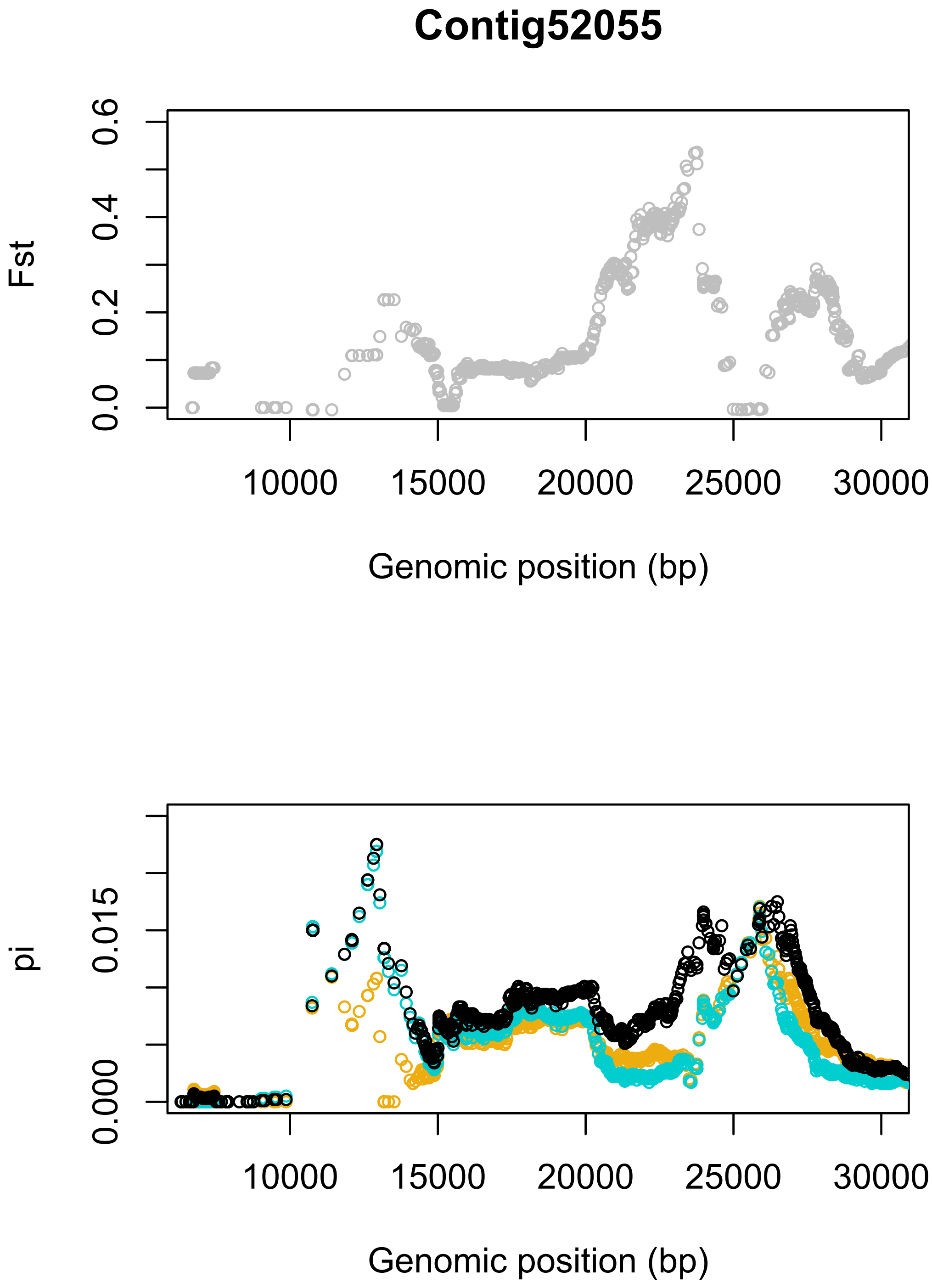
Contig52055
Fst
0.0
0.2
0.4
0.6
10000
15000
20000
25000
30000
Genomic position (bp)
pi
0.000
0.015
10000
15000
20000
25000
30000
Genomic position (bp)

**Contig128971**

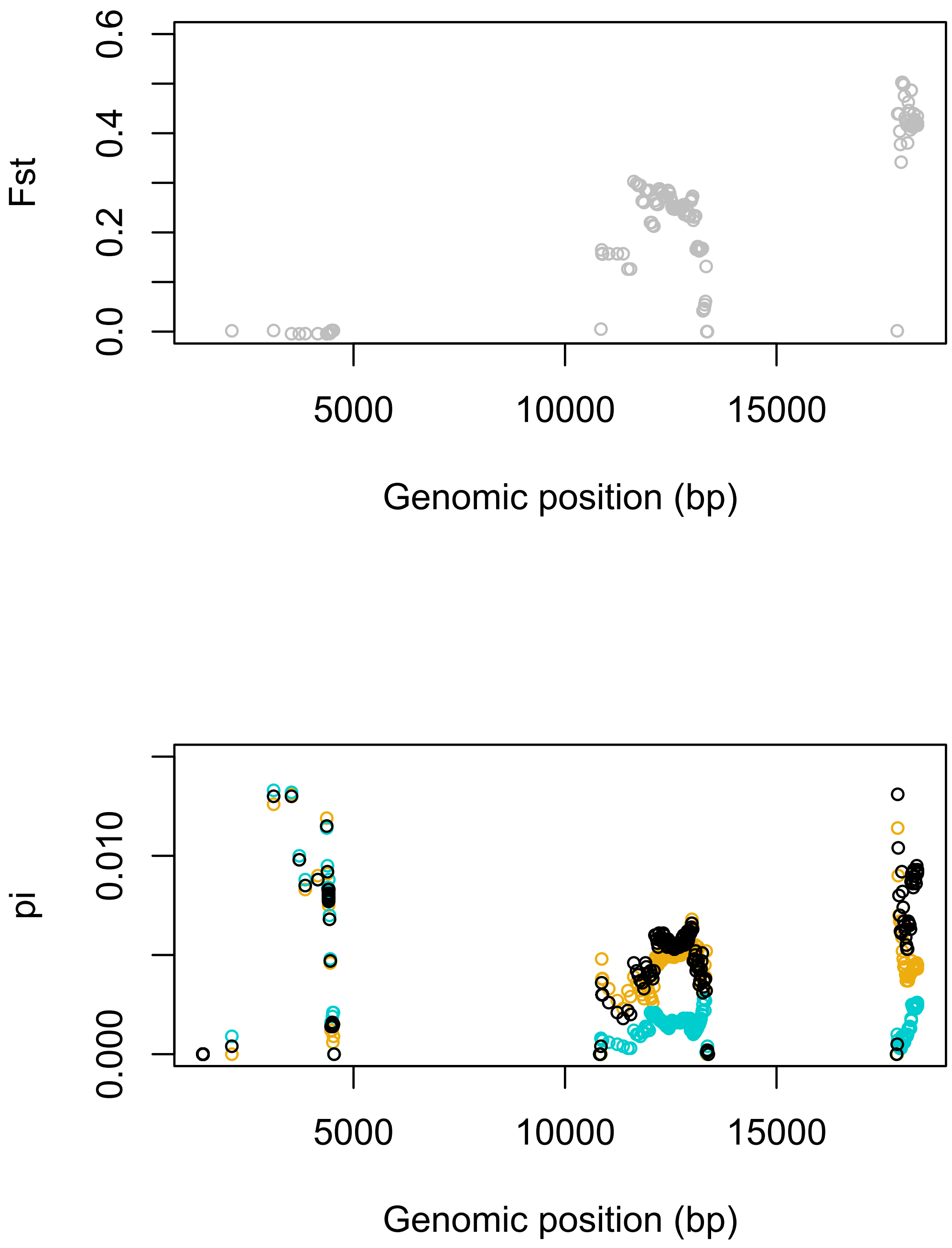

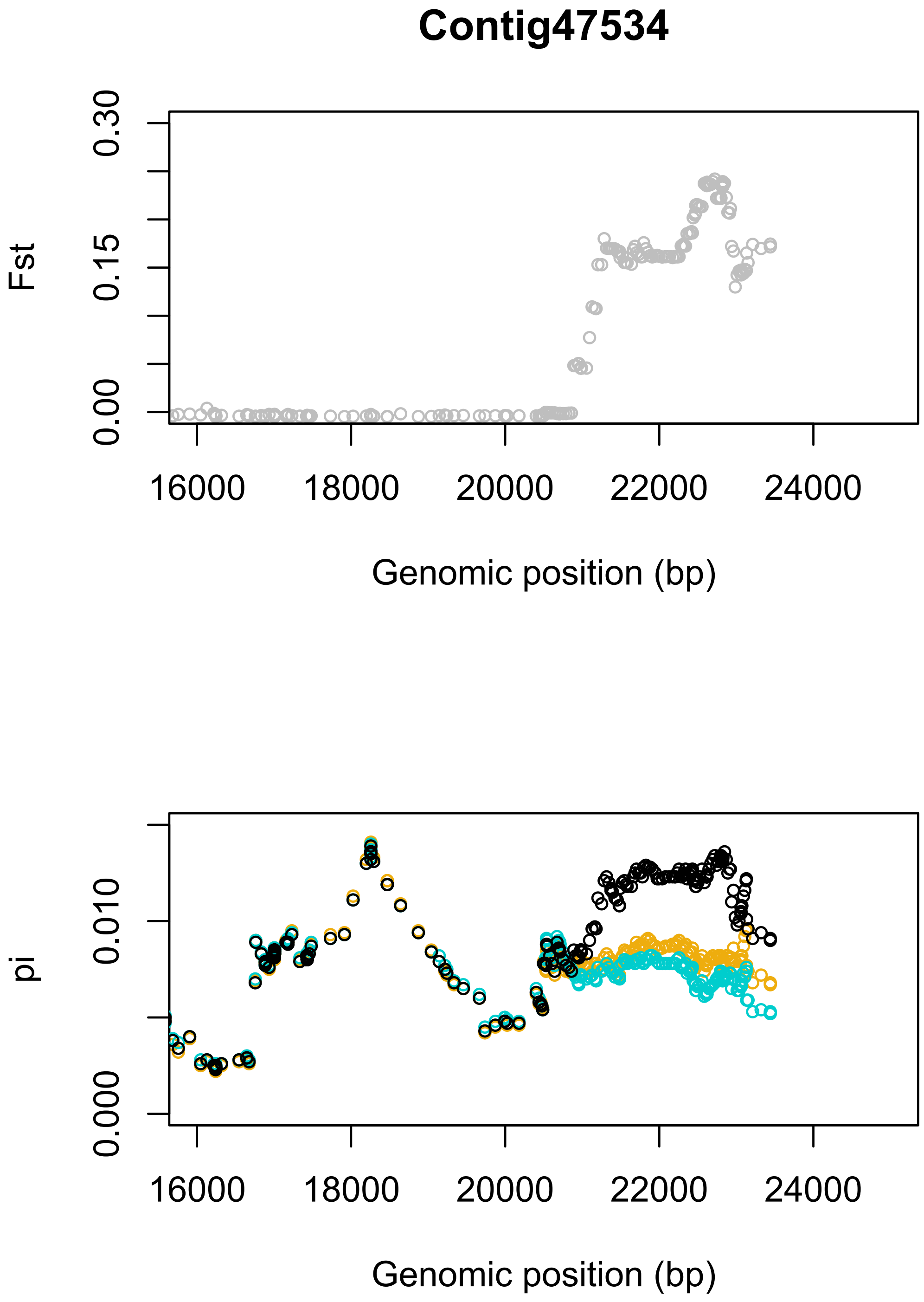
Contig47534
Fst
0.00
0.15
0.30
16000
18000
20000
22000
24000
Genomic position (bp)
pi
0.000
0.010
16000
18000
20000
22000
24000
Genomic position (bp)

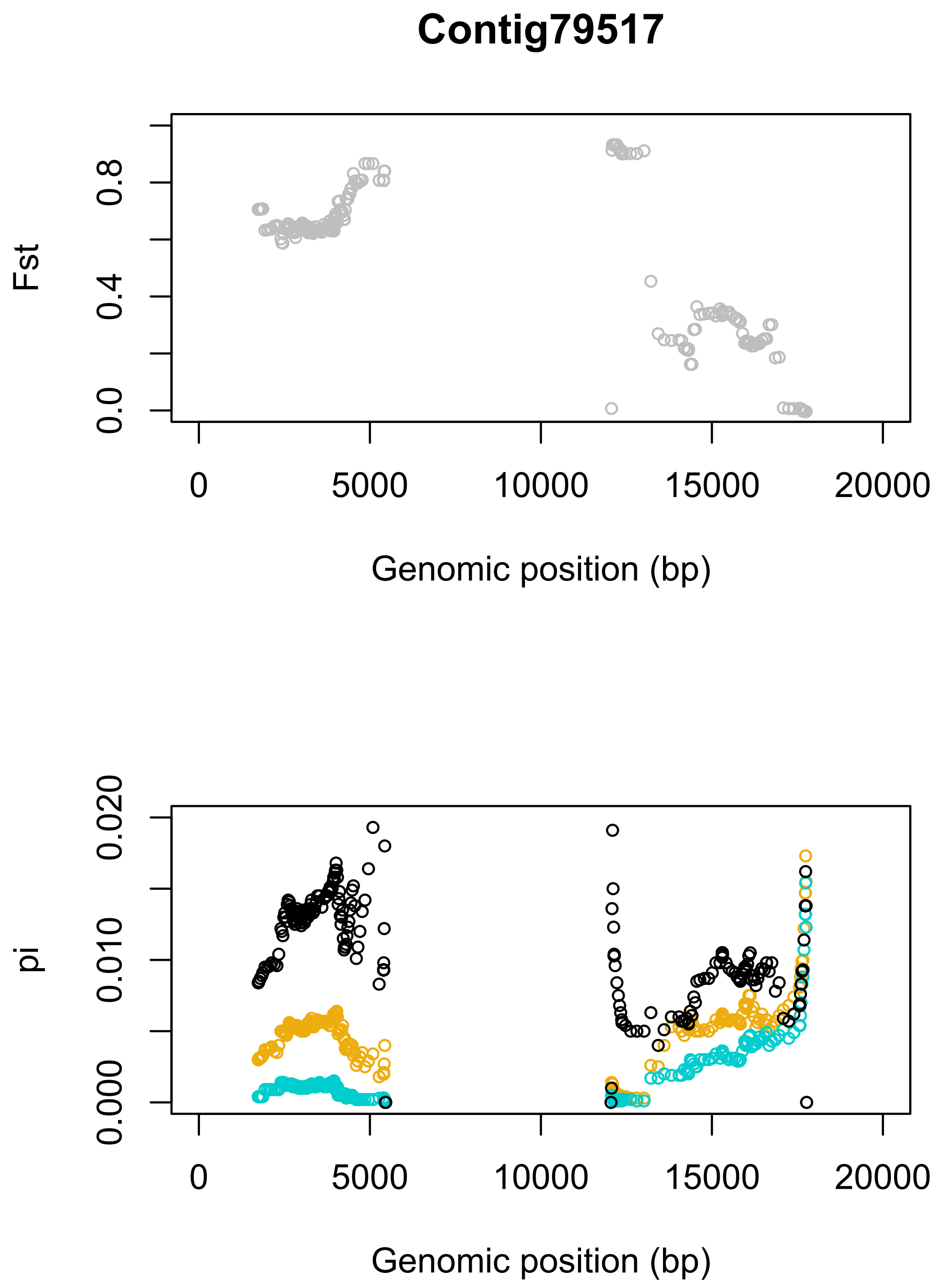
Contig79517
Fst
0.0
0.4
0.8
0
5000
10000
15000
20000
Genomic position (bp)
pi
0.000
0.010
0.020
0
5000
10000
15000
20000
Genomic position (bp)

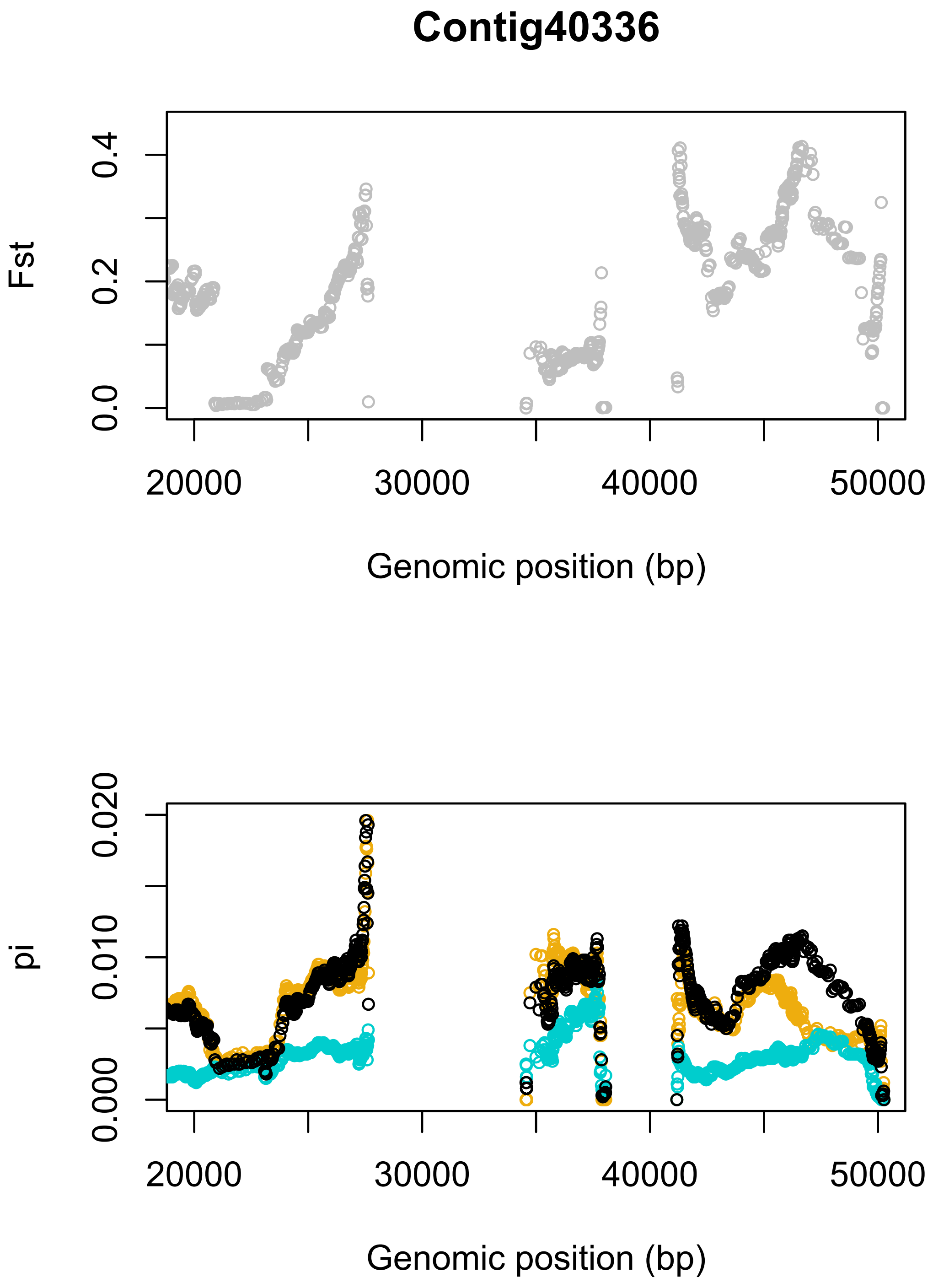
Contig40336
Fst
0.0
0.2
0.4
20000
30000
40000
50000
Genomic position (bp)
pi
0.000
0.010
0.020
20000
30000
40000
50000
Genomic position (bp)

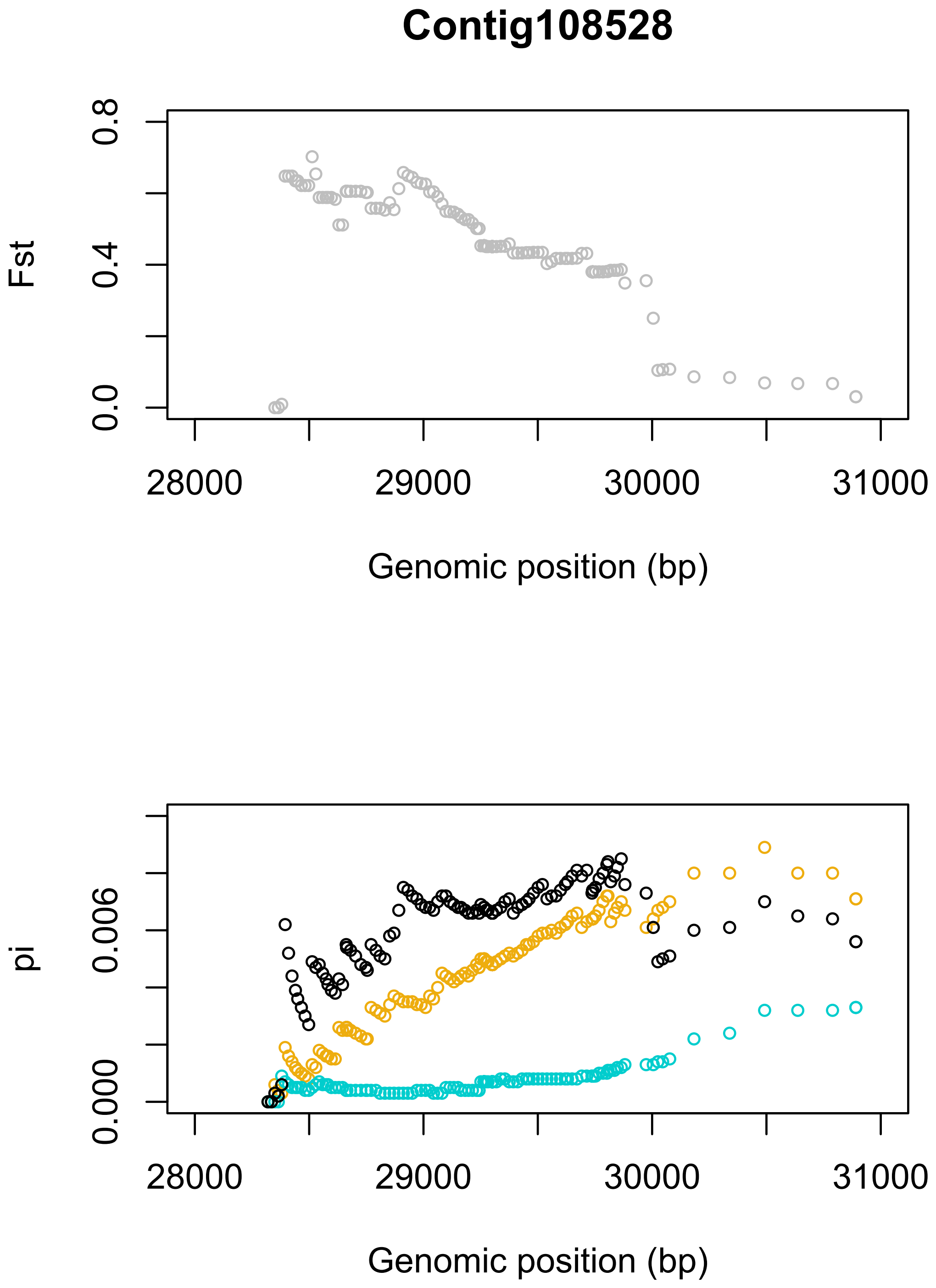
Contig108528
Fst
0.0
0.4
0.8
28000
29000
30000
31000
Genomic position (bp)
pi
0.000
0.006
28000
29000
30000
31000
Genomic position (bp)

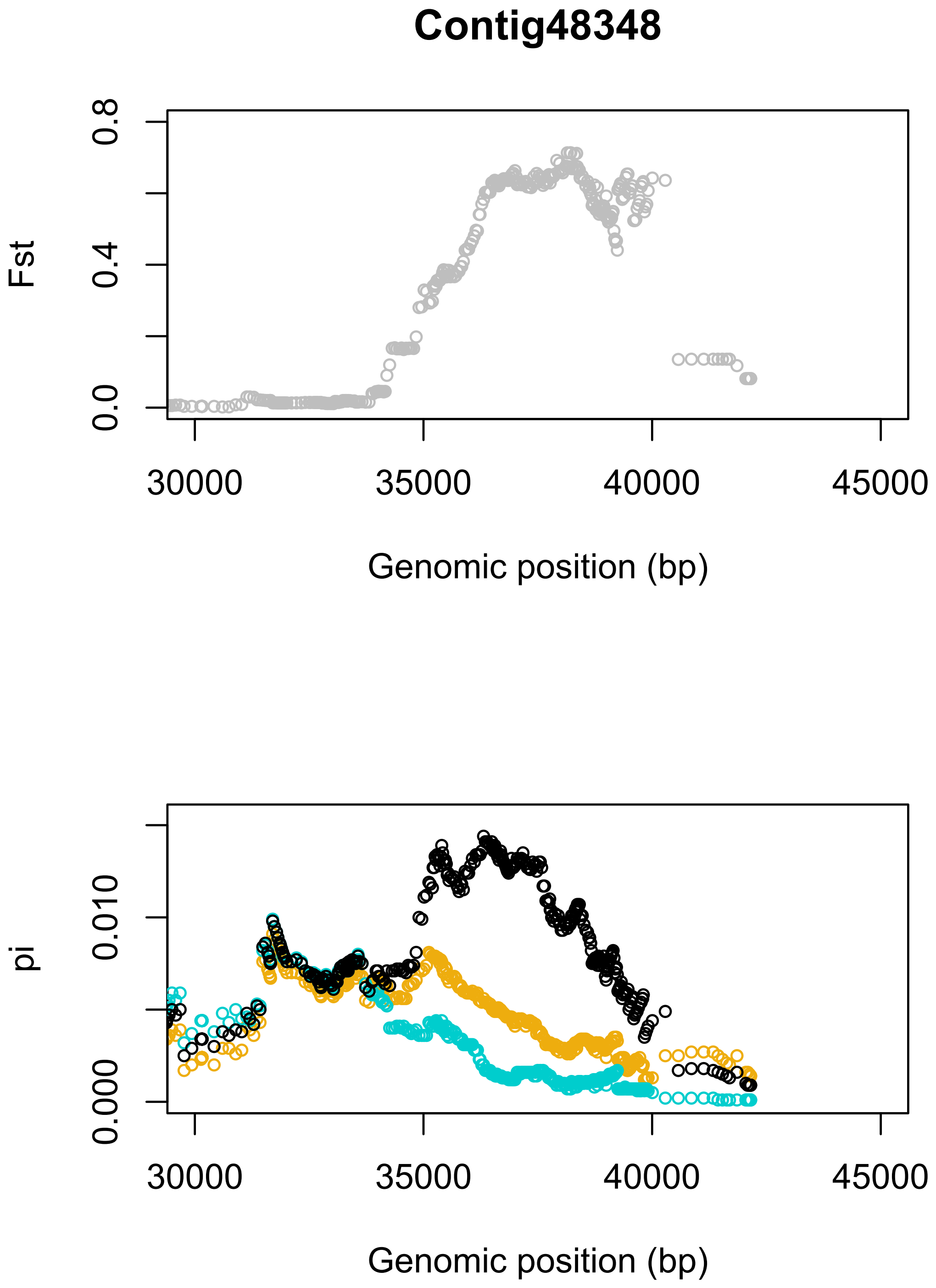
Contig48348
Fst
0.0
0.4
0.8
30000
35000
40000
45000
Genomic position (bp)
pi
0.000
0.010
30000
35000
40000
45000
Genomic position (bp)

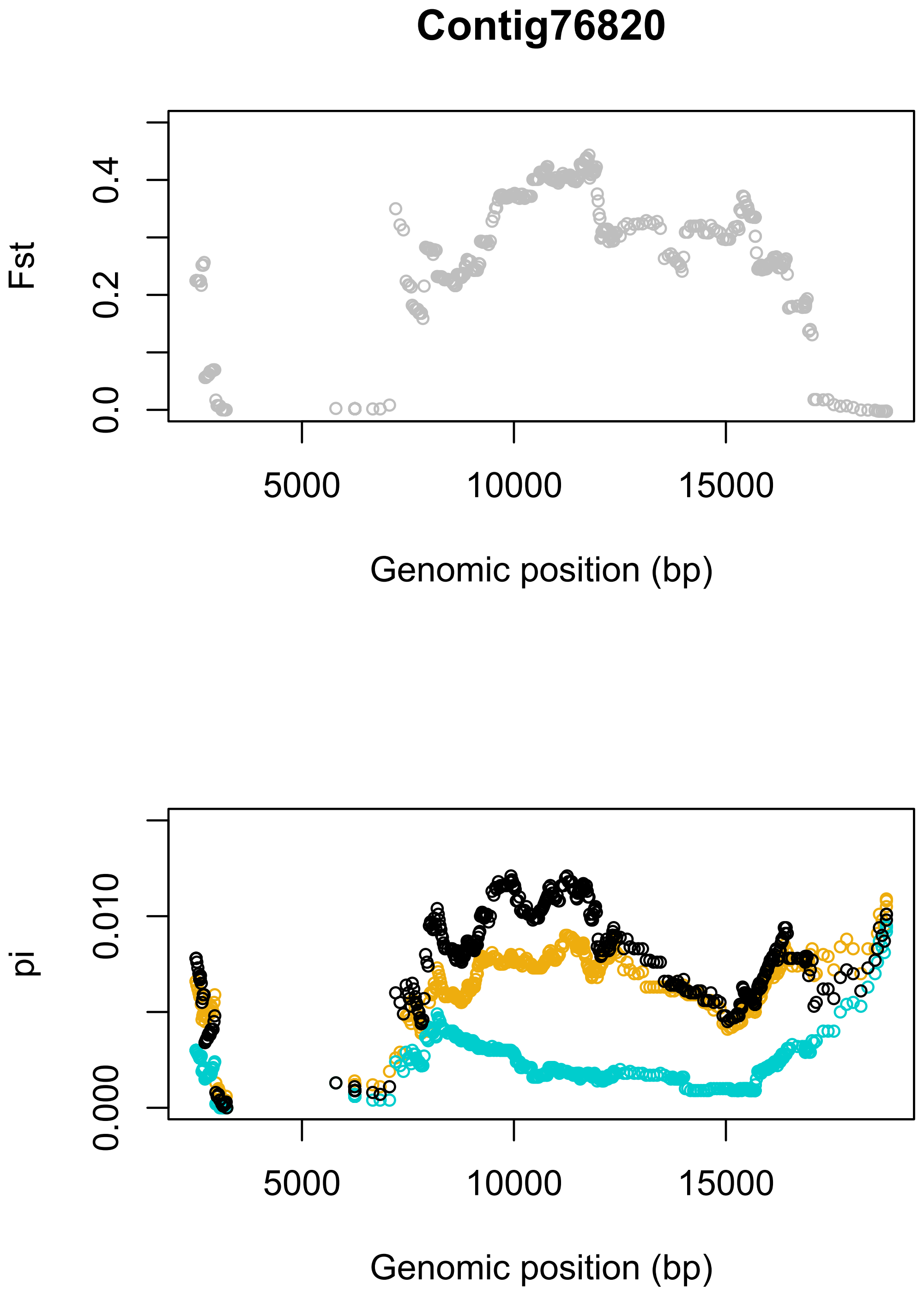
Contig76820
Fst
0.0
0.2
0.4
5000
10000
15000
Genomic position (bp)
pi
0.000
0.010
5000
10000
15000
Genomic position (bp)

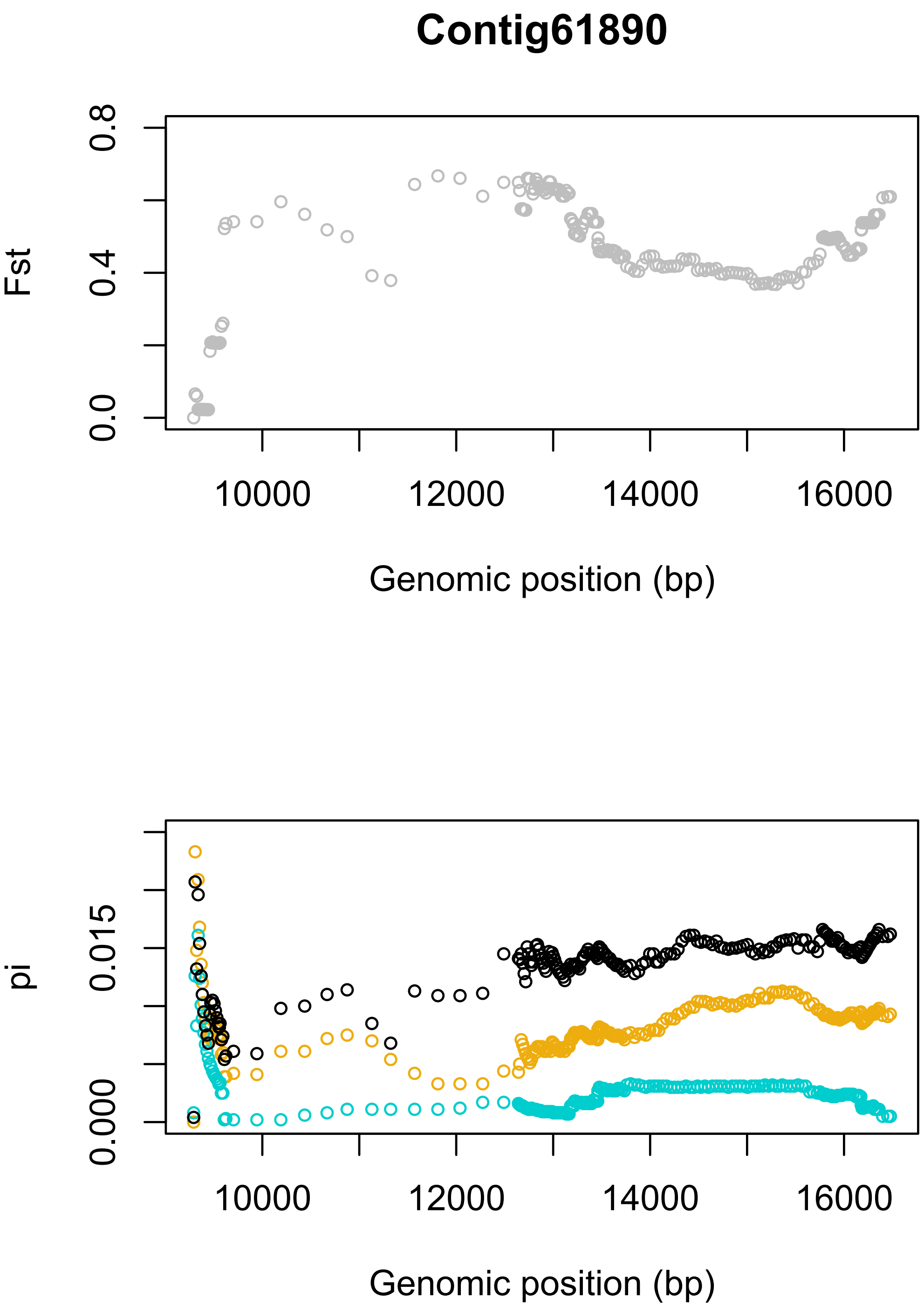
Contig61890
Fst
0.0
0.4
0.8
10000
12000
14000
16000
Genomic position (bp)
pi
0.000
0.015
10000
12000
14000
16000
Genomic position (bp)

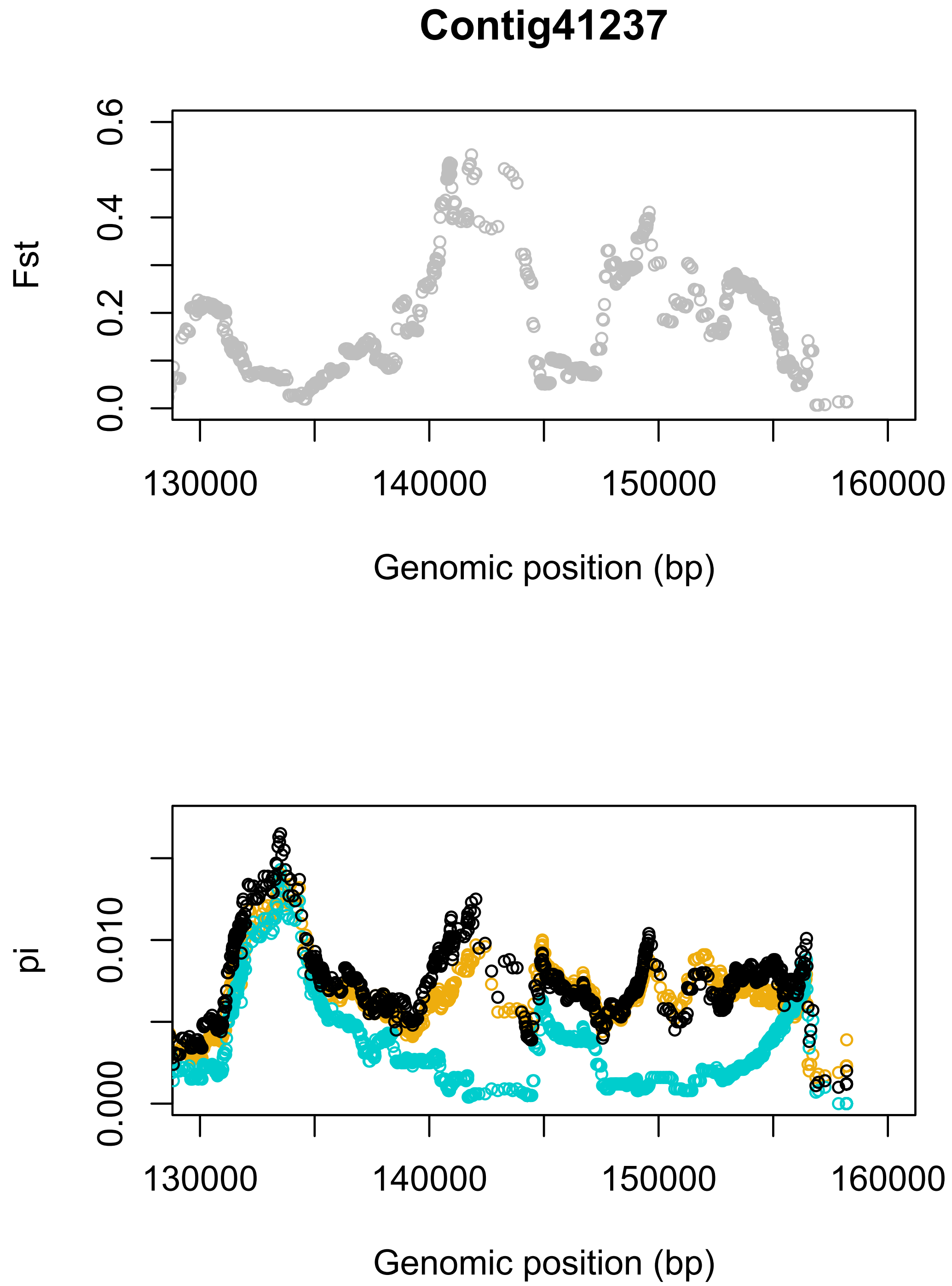
Contig41237
Fst
0.0
0.2
0.4
0.6
130000
140000
150000
160000
Genomic position (bp)
pi
0.000
0.010
130000
140000
150000
160000
Genomic position (bp)

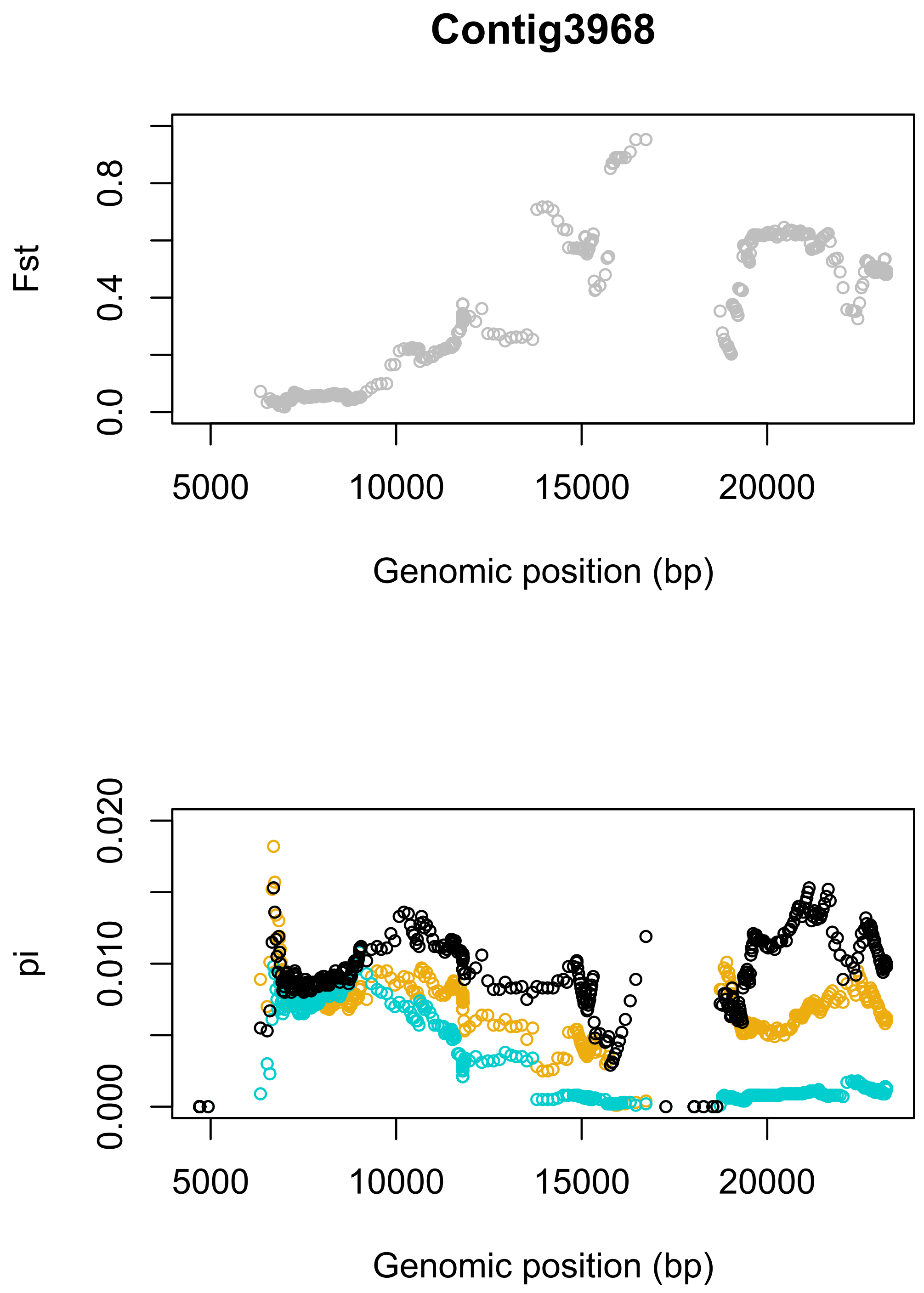
Contig3968
Fst
0.0
0.4
0.8
5000
10000
15000
20000
Genomic position (bp)
pi
0.000
0.010
0.020
5000
10000
15000
20000
Genomic position (bp)

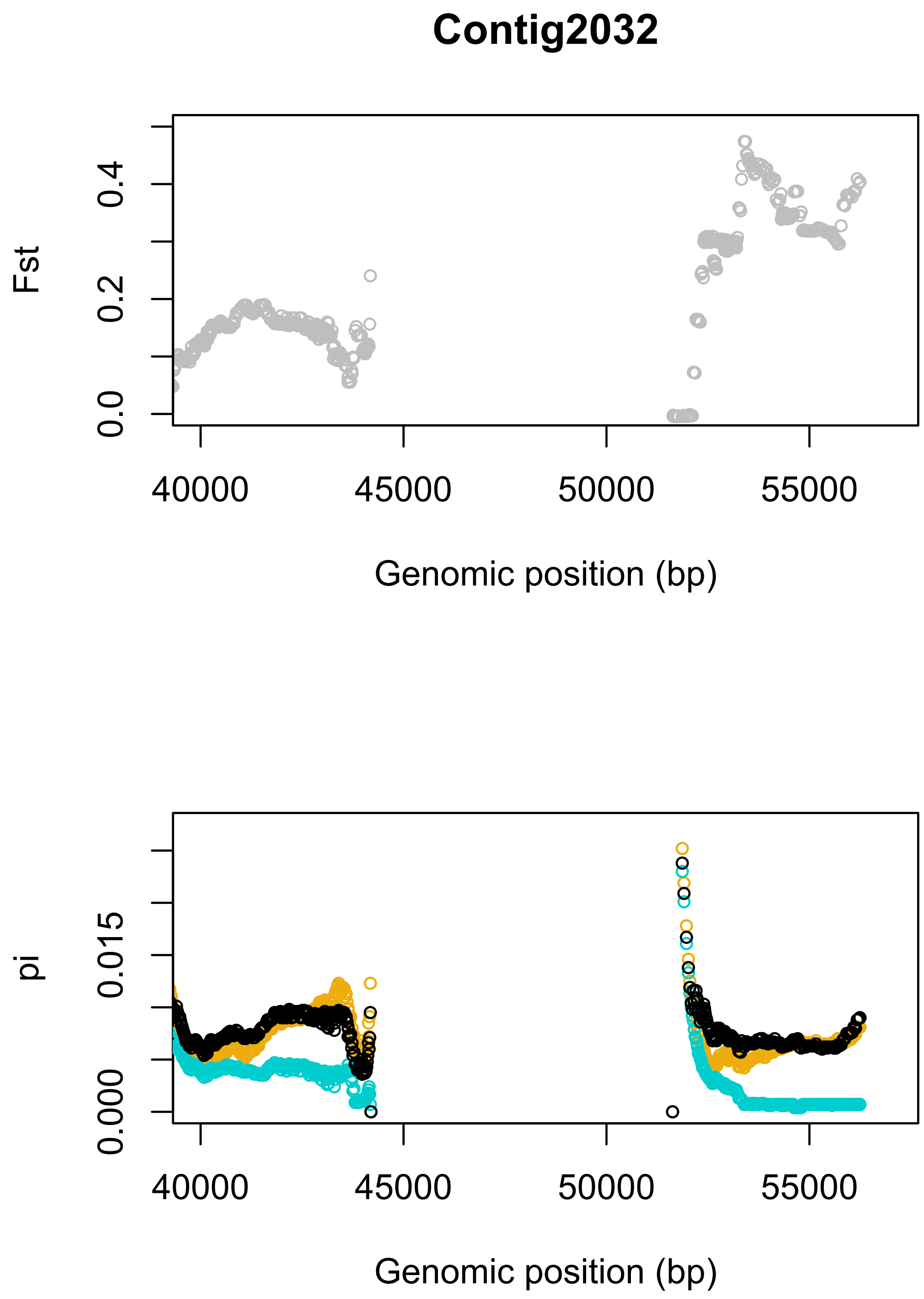
Contig2032
Fst
0.0
0.2
0.4
40000
45000
50000
55000
Genomic position (bp)
pi
0.000
0.015
40000
45000
50000
55000
Genomic position (bp)

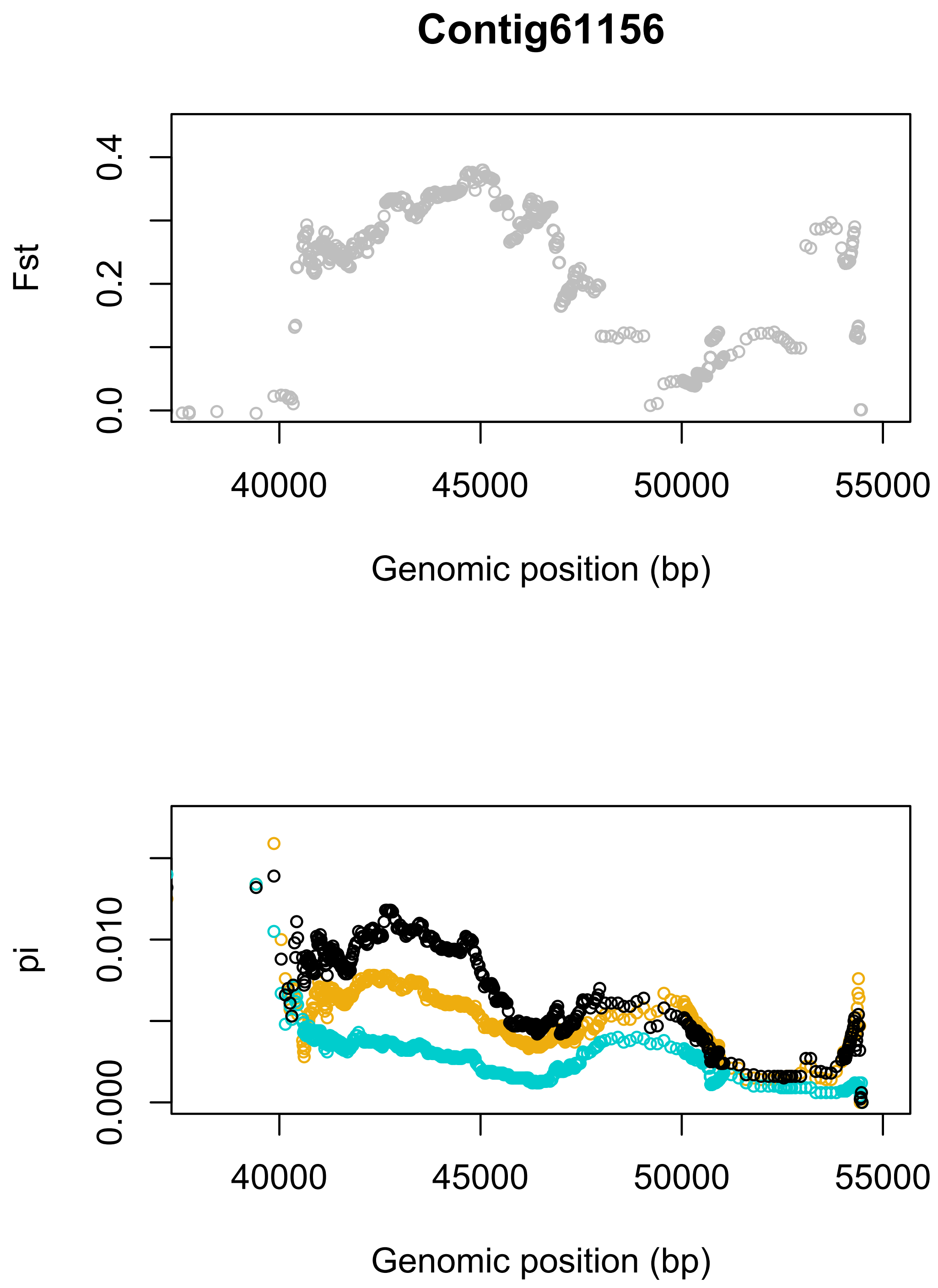
Contig61156
Fst
0.0
0.2
0.4
40000
45000
50000
55000
Genomic position (bp)
pi
0.000
0.010
40000
45000
50000
55000
Genomic position (bp)

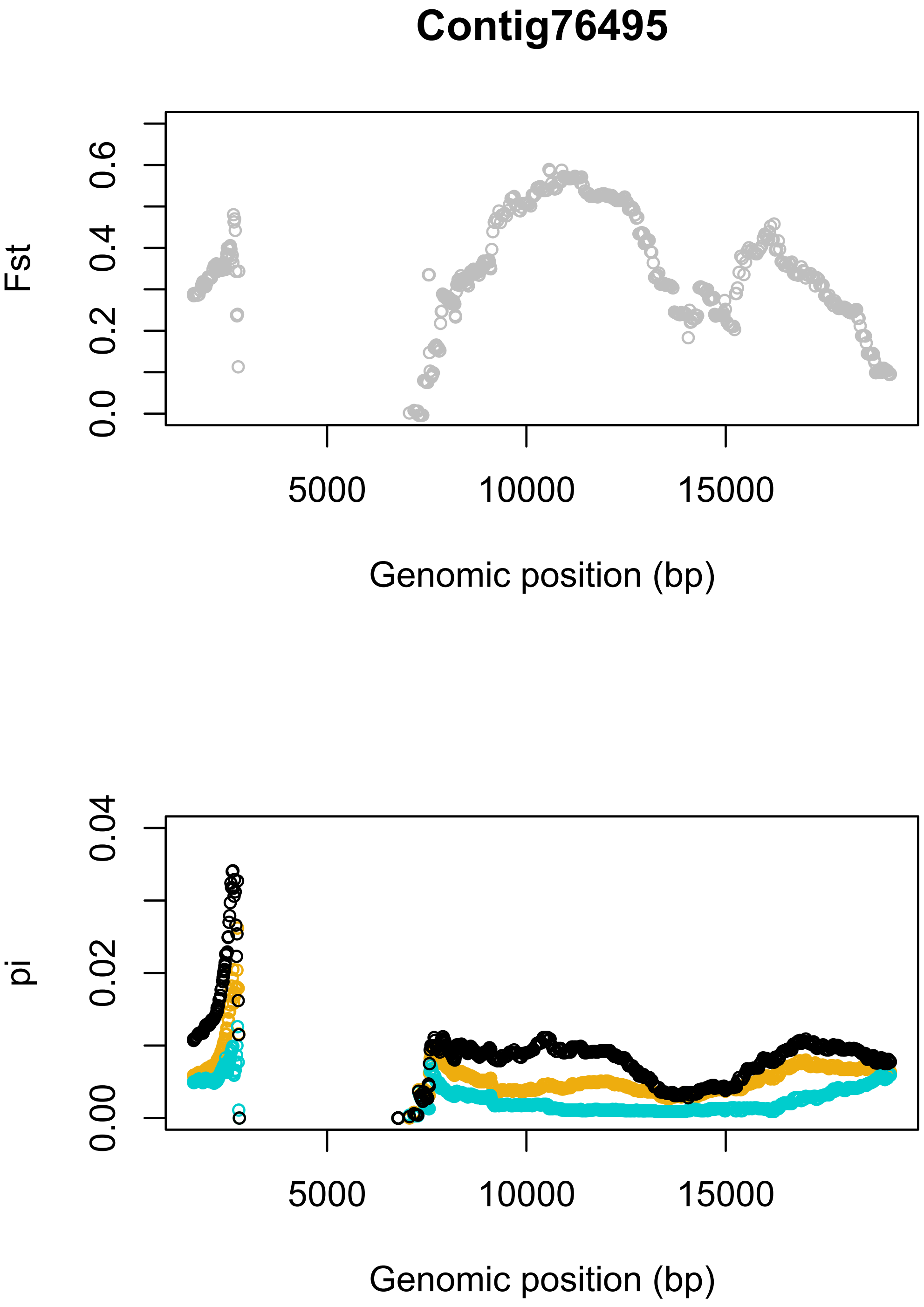
Contig76495
Fst
0.0
0.2
0.4
0.6
5000
10000
15000
Genomic position (bp)
pi
0.00
0.02
0.04
5000
10000
15000
Genomic position (bp)

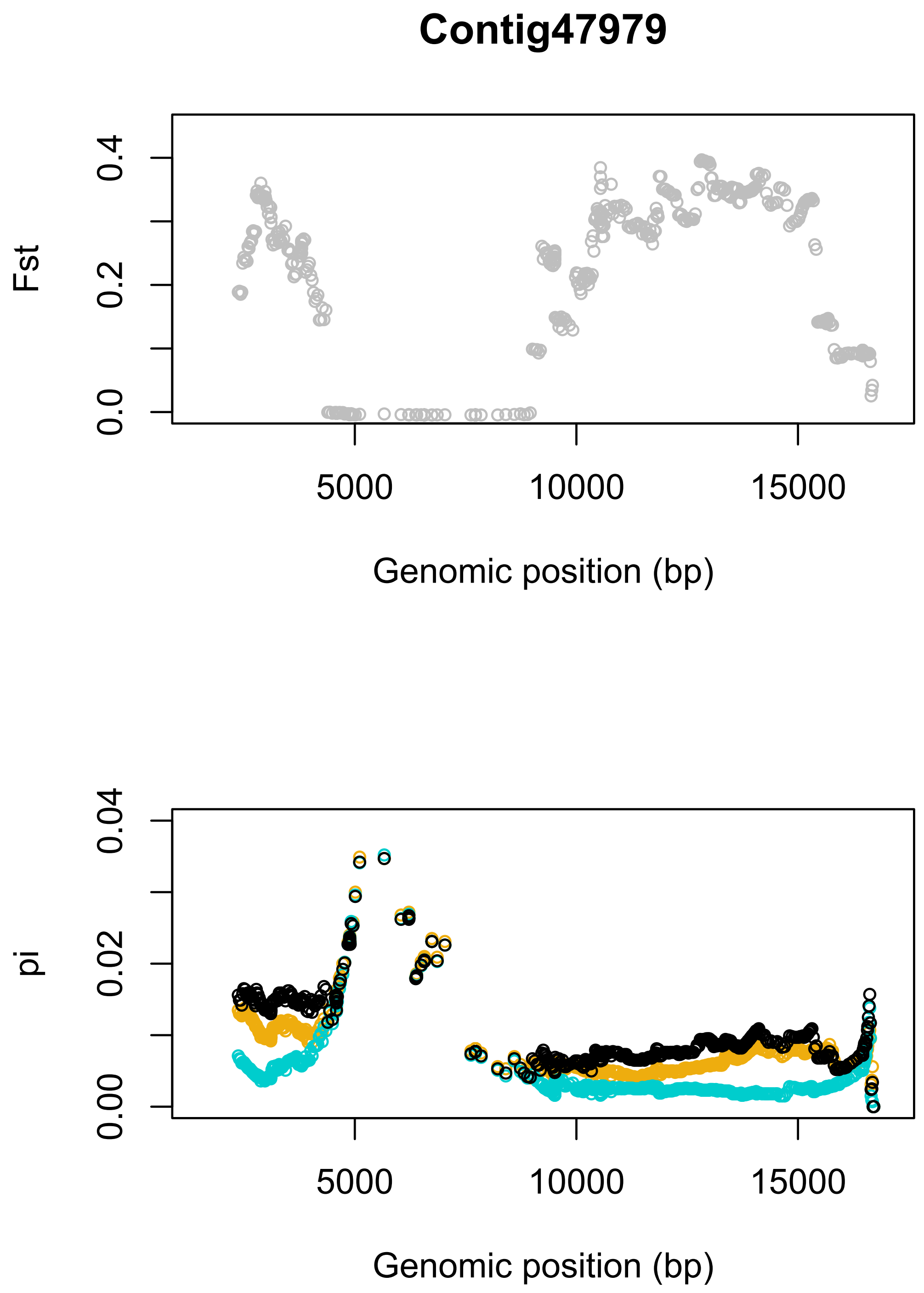
Contig47979
Fst
0.0
0.2
0.4
5000
10000
15000
Genomic position (bp)
pi
0.00
0.02
0.04
5000
10000
15000
Genomic position (bp)

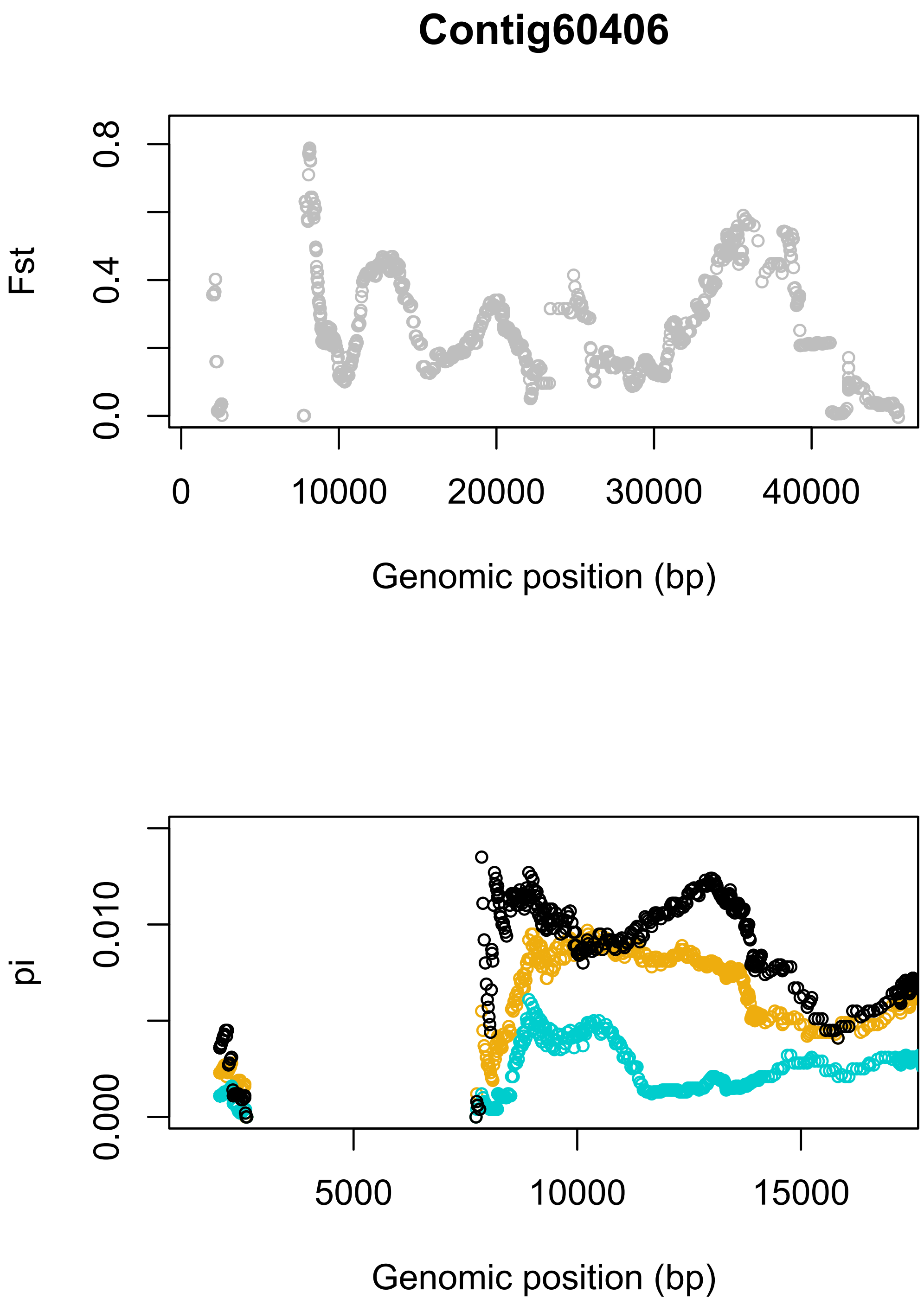
Contig60406
Fst
0.0
0.4
0.8
0
10000
20000
30000
40000
Genomic position (bp)
pi
0.000
0.010
5000
10000
15000
Genomic position (bp)

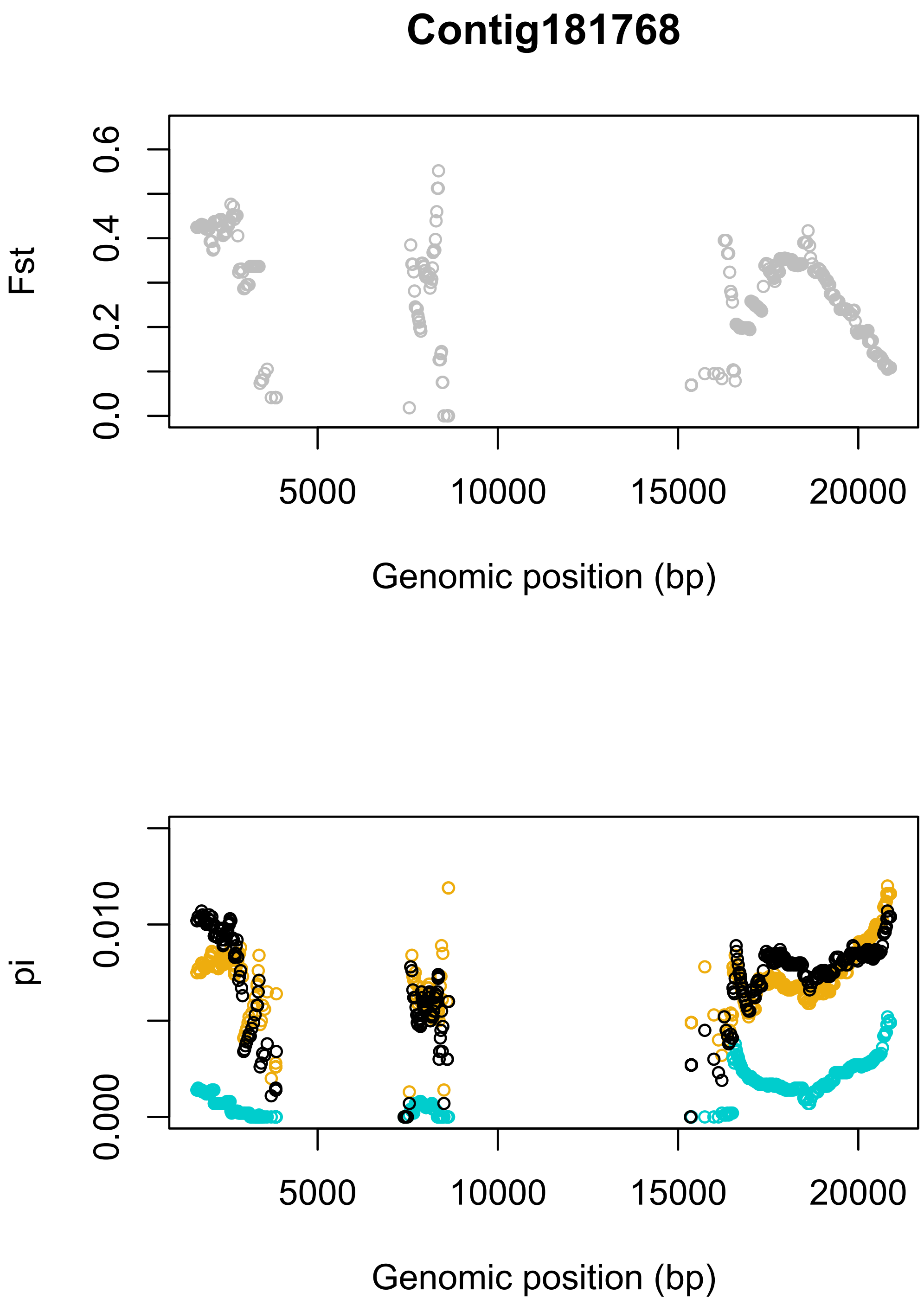
Contig181768
Fst
0.0
0.2
0.4
0.6
5000
10000
15000
20000
Genomic position (bp)
pi
0.000
0.010
5000
10000
15000
20000
Genomic position (bp)

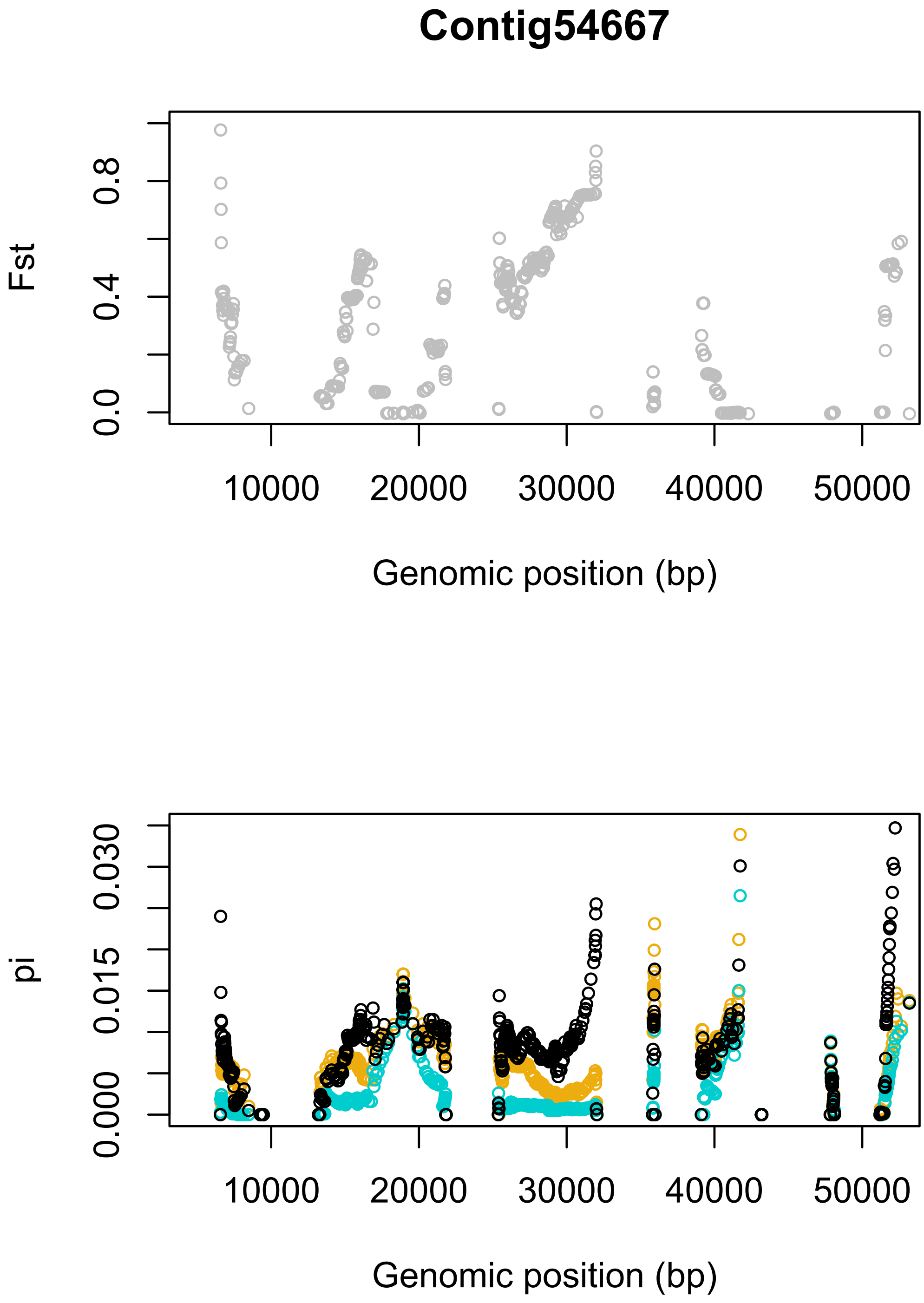
Contig54667
Fst
0.0
0.4
0.8
10000
20000
30000
40000
50000
Genomic position (bp)
pi
0.000
0.015
0.030
10000
20000
30000
40000
50000
Genomic position (bp)

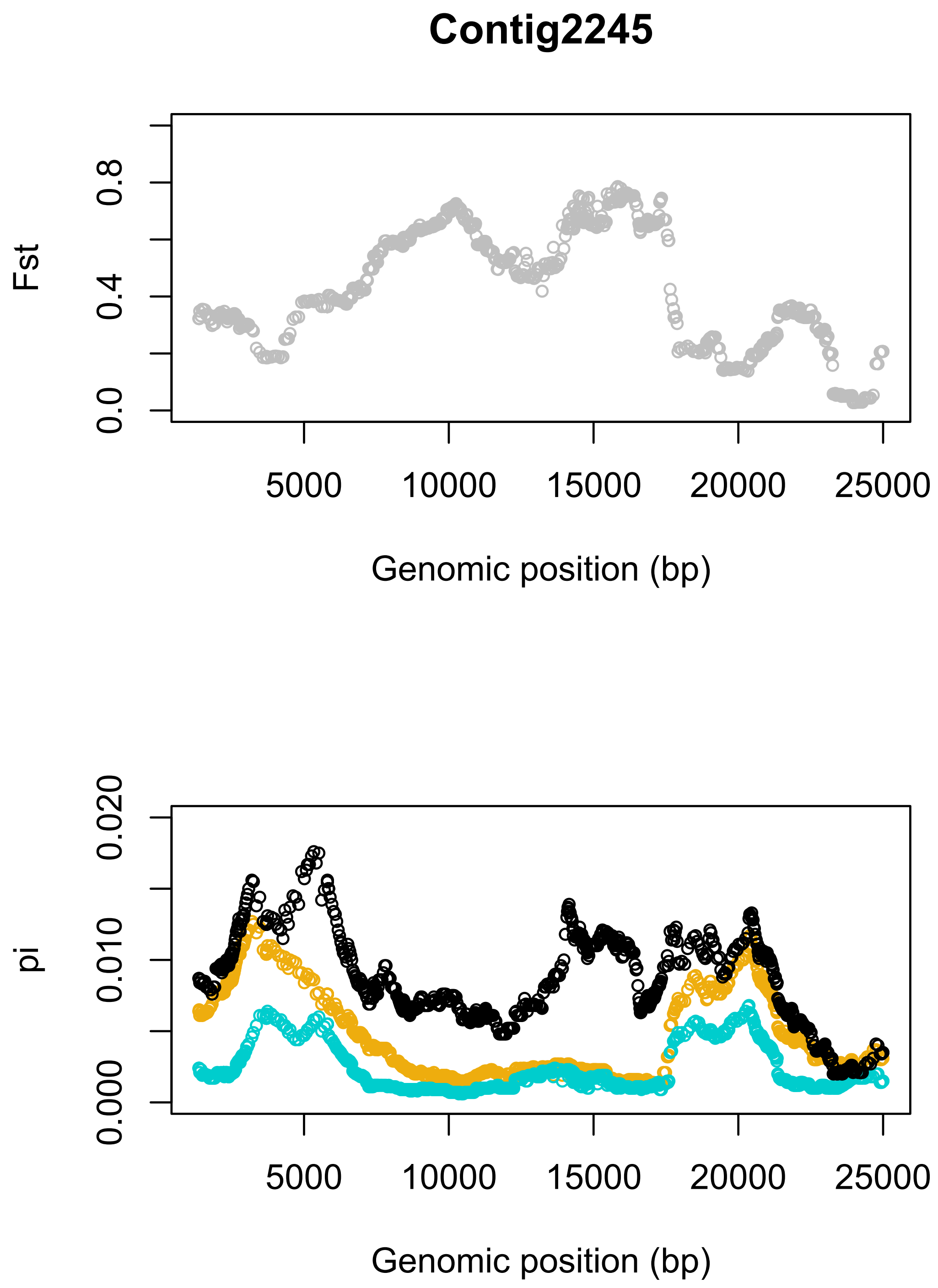
Contig2245
Fst
0.0
0.4
0.8
5000
10000
15000
20000
25000
Genomic position (bp)
pi
0.000
0.010
0.020
5000
10000
15000
20000
25000
Genomic position (bp)

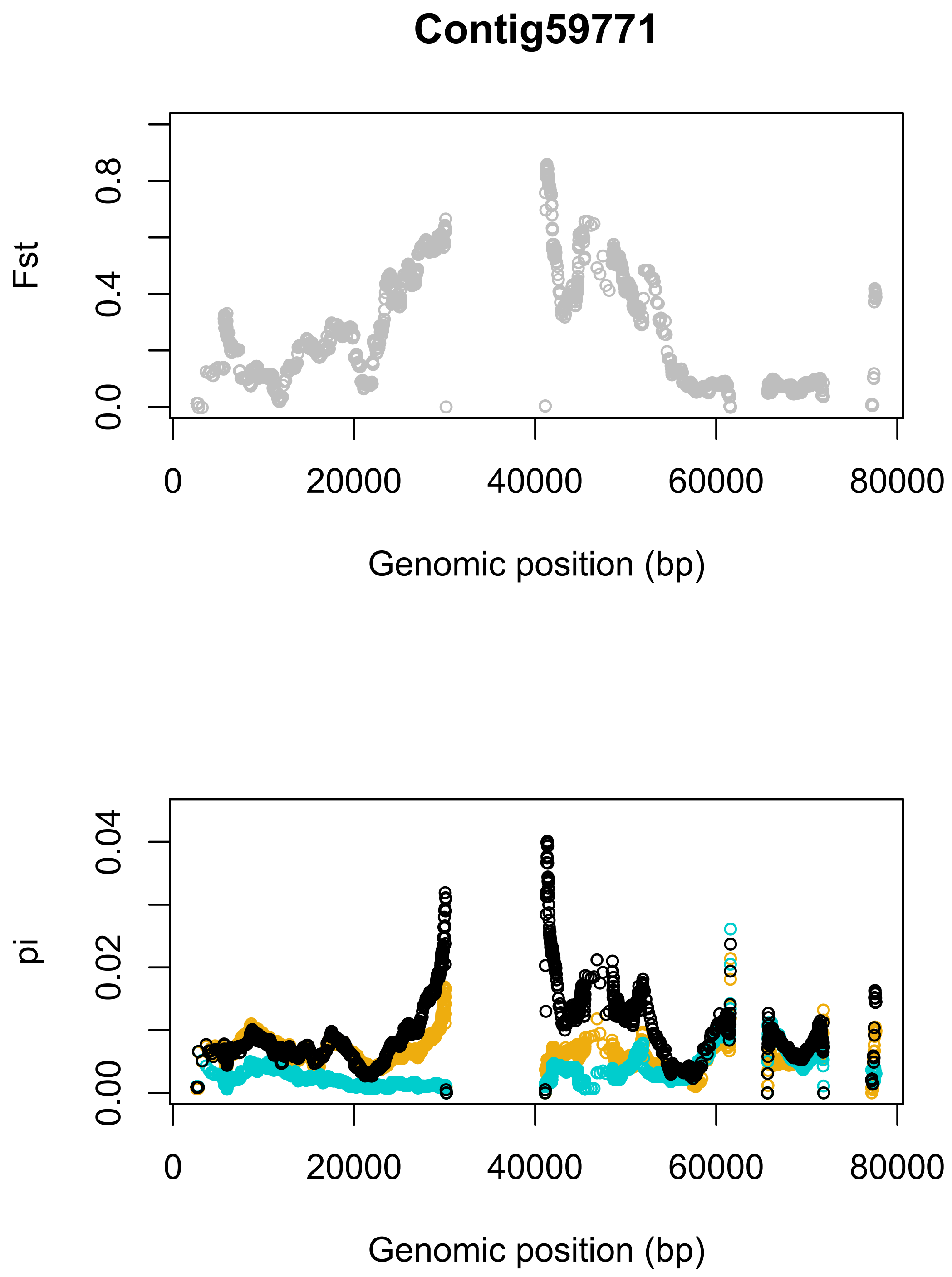
Contig59771
Fst
0.0
0.4
0.8
0
20000
40000
60000
80000
Genomic position (bp)
pi
0.00
0.02
0.04
0
20000
40000
60000
80000
Genomic position (bp)

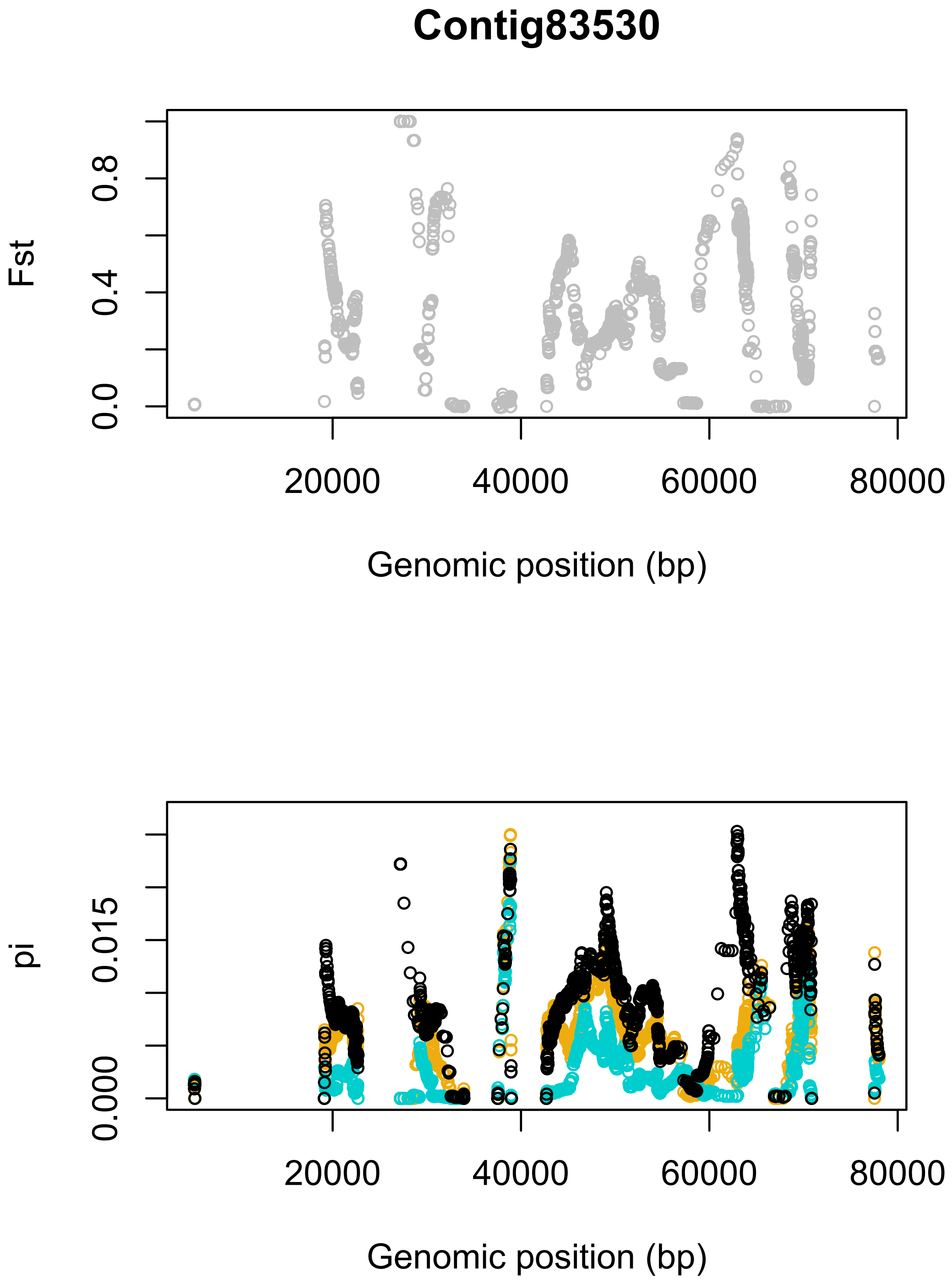
Contig83530
Fst
0.0
0.4
0.8
20000
40000
60000
80000
Genomic position (bp)
pi
0.000
0.015
20000
40000
60000
80000
Genomic position (bp)

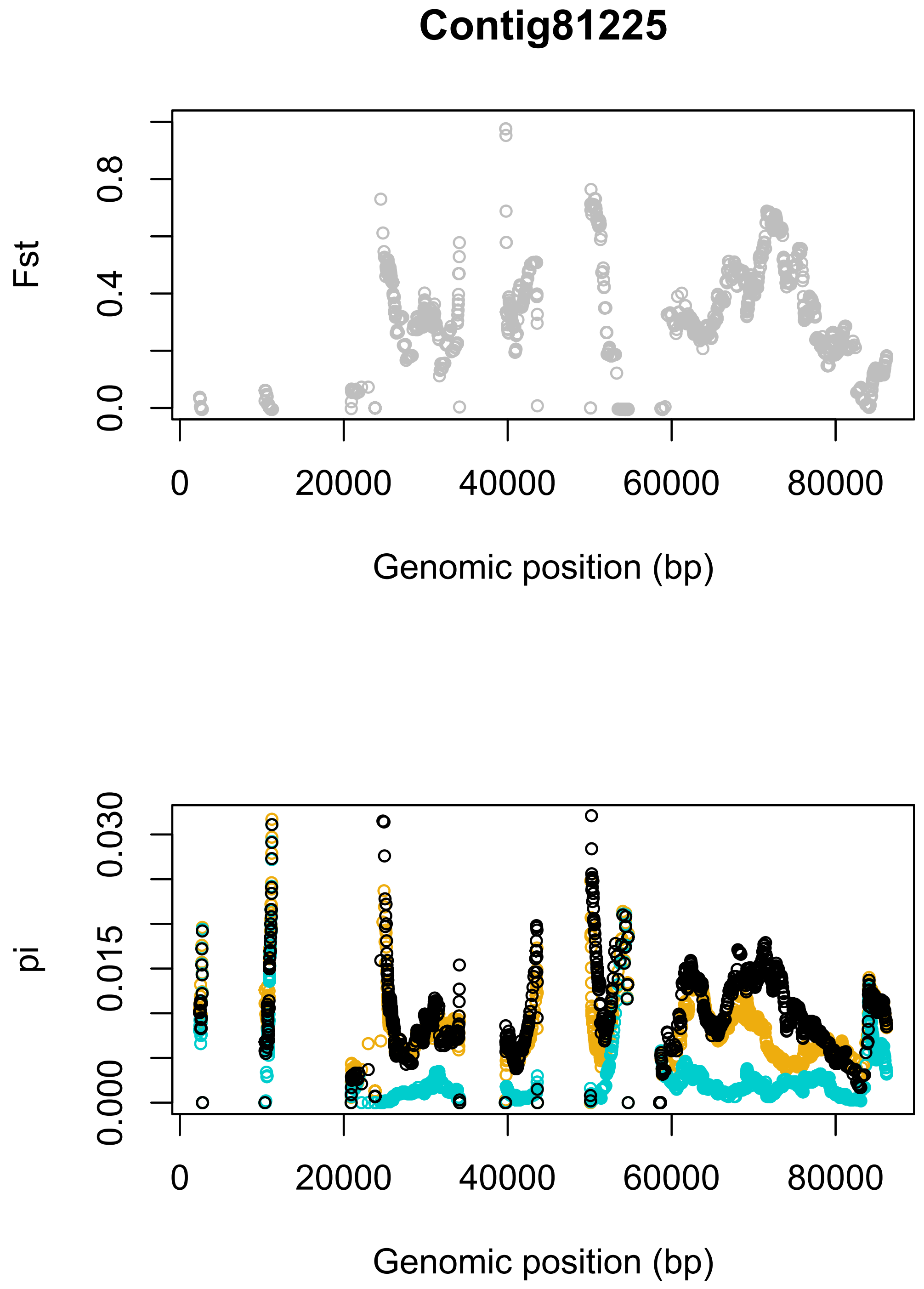
Contig81225
Fst
0.0
0.4
0.8
0
20000
40000
60000
80000
Genomic position (bp)
pi
0.000
0.015
0.030
0
20000
40000
60000
80000
Genomic position (bp)

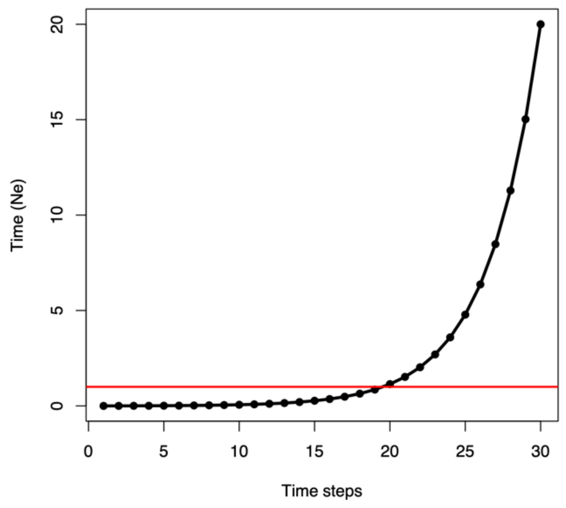

**Figure S34. Discrete time steps for ARGweaver analysis.** Coalescence and recombination events are only able to occur at these timepoints. The maximum allowed TMRCA is $20N_e$. Solid red line marks $1N_e$.

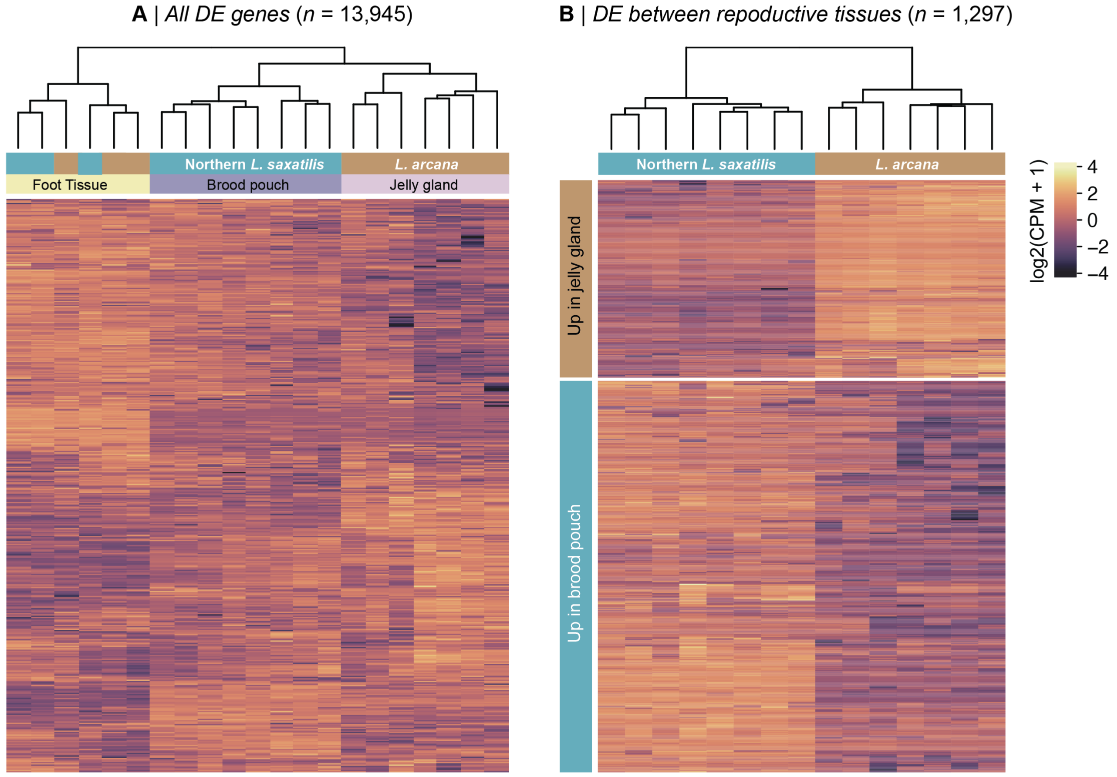

**Figure S35. Clustering and heatmap analysis of gene expression in sequenced libraries.** The left plot shows the results for all 13,945 genes that passed our filtering steps. The dendrogram shows that the libraries cluster by tissue type, but that egg-layers and live-bearers are not distinct based on the expression patterns in foot tissue. The right plot shows the result for the 1,297 genes that were differentially expressed in reproductive tissue. The brood pouch and jelly gland libraries form separate clusters, with two-thirds of genes showing higher expression in the live-bearing tissue.

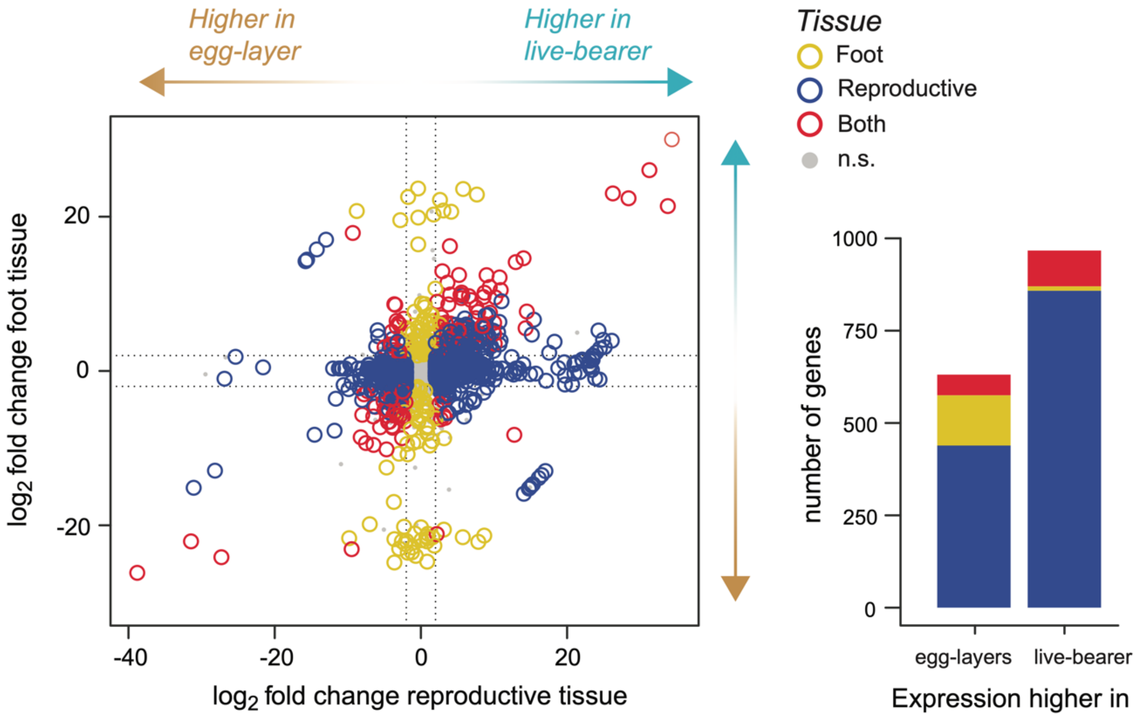

**Figure S36. Results of differential expression analysis.** The left plot shows the joint distribution of $\log_2$ fold change between the reproductive tissues (x-axis) and foot-tissues (y-axis). Arrows on each axis show the directions associated with higher expression in egg-layers or live-bearers. The color of each point indicates whether genes showed differential expression between the foot tissues only, reproductive tissues only, or in both the foot and reproductive tissues. The right plot shows the number of genes in each expression class that showed higher expression in egg-layers and live-bearers.

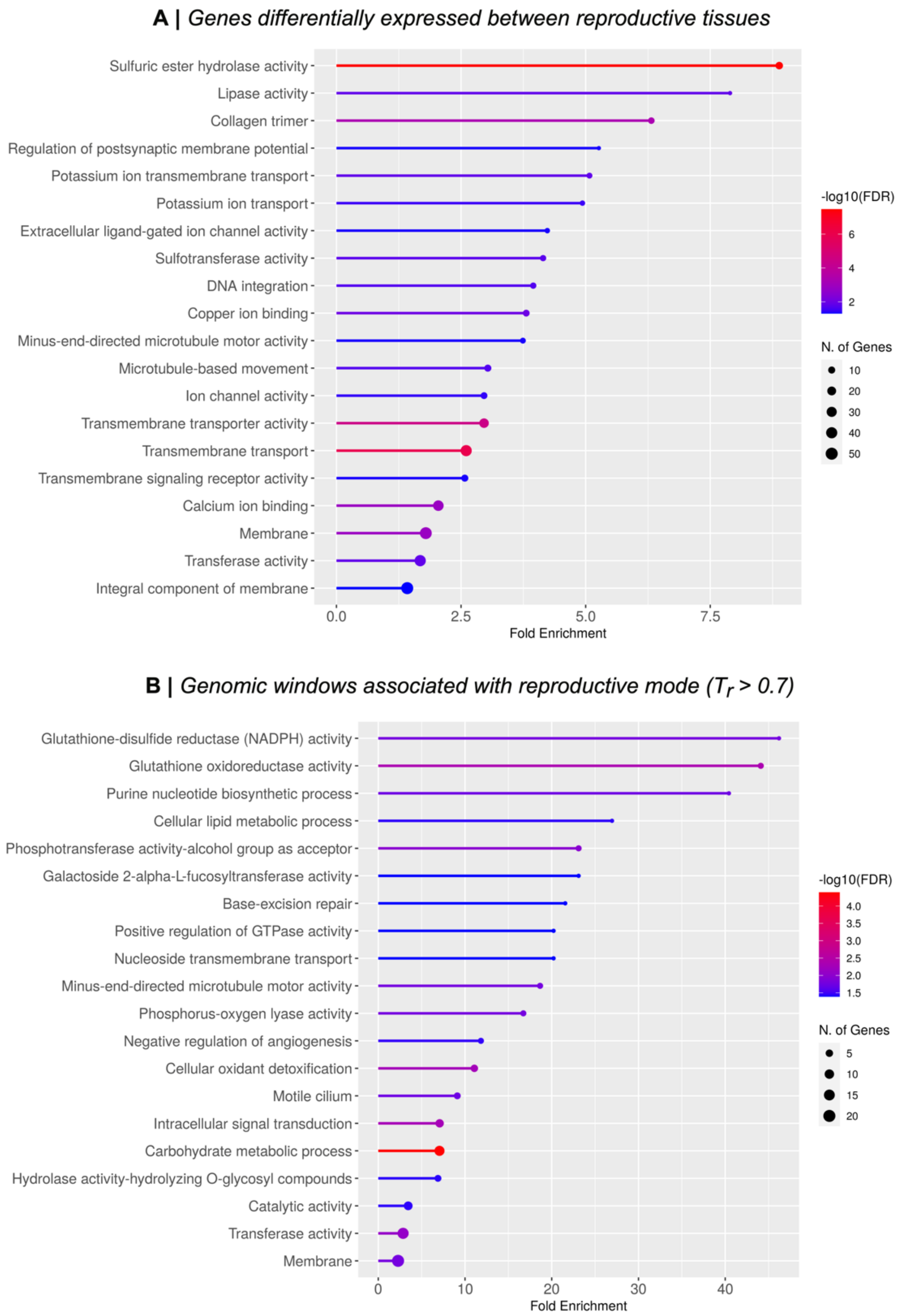

**Figure S37. Enrichment plots showing enriched gene ontology terms two reproductive mode associated gene sets.** (A) All differentially expressed genes and have a corresponding gene from *Crassostrea virginica* (*n* = 639). (B) All genes in regions that show a genealogical association with reproductive mode and that have a corresponding gene from *Crassostrea virginica* (*n* = 574).

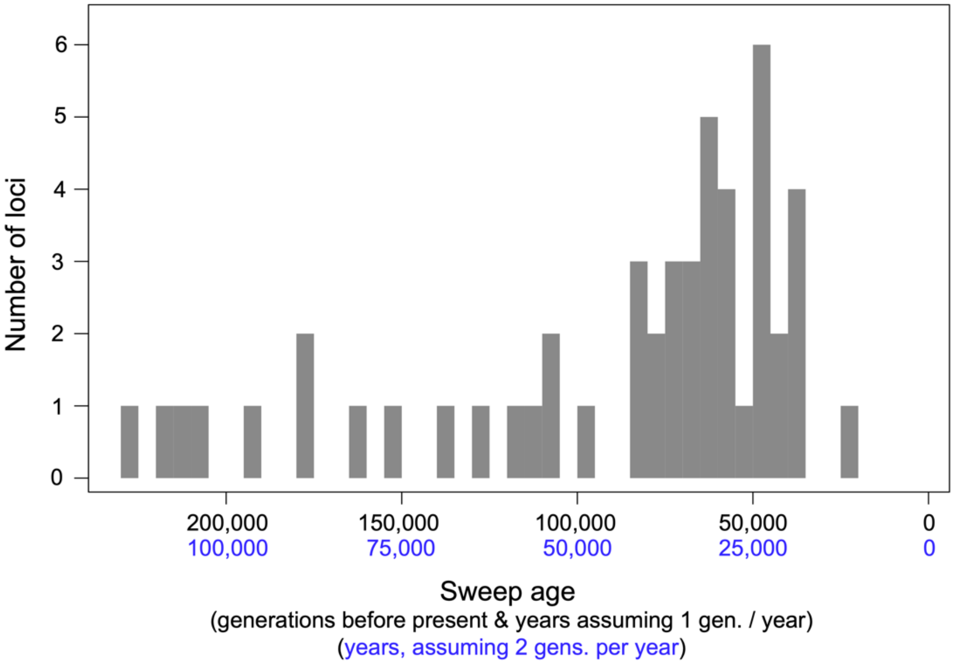

**Figure S38. Distribution of the ages of sweeps at live-bearing alleles**. For the 50 regions that are perfectly associated with live-birth (Tr = 1), we estimated the age of each sweep using the equation $T = \pi_w/2\mu$, where $\pi_w$ is the diversity in the swept segment arising from new mutation (calculated from live-bearing-specific private alleles), $\mu$ is the mutation rate, and $T$ is the time in generations. For calculating $T$, we assumed a mutation rate of $\mu = 1.5\times10^{-8}$. We converted time in generations into years using two estimates of generation time for *Littorina saxatilis*: 2 generations per year and 1 generation per year (*65*, *66*). Estimates of time in generations ranged from 23,266 to 227,094 generations before present, with a median of 69,790 generations before present. In years, this equates to a minimum of 11,633 – 23,266 years before present, and maximum of 113,547 – 227,094 years before present, with a median of 34,895 – 69,790 years before present. The distribution of ages is strongly negatively skewed, with the majority of sweeps being initiated in the last 100,000 generations.

**Table S1. Sample information.** Sample IDs, species, ecotype and sex and are provided.

| Sample ID | Species | Ecotype | Country | location | Sex | Latitude | Longitude |
|---|---|---|---|---|---|---|---|
| **AMB_5_3** | *L. arcana* | Wave | England | Amble | F | 55.33215 | -1.56292 |
| **BH_10_1** | *L. arcana* | Wave | Wales | Broad Haven | F | 51.60891 | -4.91878 |
| **BH_8_2** | *L. saxatilis* | Wave | Wales | Broad Haven | F | 51.60891 | -4.91878 |
| **BUR_Lsax_F** | *L. saxatilis* | Other | Spain | Burella | F | 43.66556 | -7.35782 |
| **BUR_Lsax_M** | *L. saxatilis* | Other | Spain | Burella | M | 43.66556 | -7.35782 |
| **CEA_Larc_F_2** | *L. saxatilis* | Wave | Ireland | Ceann Tra | F | 52.13205 | -10.36071 |
| **CEA_Larc_F_1** | *L. compressa* | N/A | Ireland | Ceann Tra | F | 52.13205 | -10.36071 |
| **CEA_Lsax_F_1** | *L. saxatilis* | Crab | Ireland | Ceann Tra | F | 52.13205 | -10.36071 |
| **DER_Lten_F** | *L. saxatilis* | Brackish | England | Dersingham | F | 52.8675 | 0.44738 |
| **DER_Lten_M** | *L. saxatilis* | Brackish | England | Dersingham | M | 52.8675 | 0.44738 |
| **W_arc_01_La** | *L. arcana* | Wave | Wales | Holyhead | F | 53.29981 | -4.67967 |
| **W_arc_02_La** | *L. arcana* | Wave | Wales | Holyhead | F | 53.29981 | -4.67967 |
| **W_arc_03_La** | *L. arcana* | Wave | Wales | Holyhead | F | 53.29981 | -4.67967 |
| **W_arc_04_La** | *L. arcana* | Wave | Wales | Holyhead | F | 53.29981 | -4.67967 |
| **W_com_01_Lc** | *L. compressa* | N/A | Wales | Holyhead | F | 53.29981 | -4.67967 |
| **83_AB1_Lsb** | *L. saxatilis* | Barnacle | Wales | Holyhead | F | 53.29981 | -4.67967 |
| **88_AB1_Lsb** | *L. saxatilis* | Barnacle | Wales | Holyhead | F | 53.29981 | -4.67967 |
| **W_brood pouch_01_Ls** | *L. saxatilis* | Wave | Wales | Holyhead | F | 53.29981 | -4.67967 |
| **W_brood pouch_02_Ls** | *L. saxatilis* | Wave | Wales | Holyhead | F | 53.29981 | -4.67967 |
| **W_brood pouch_03_Ls** | *L. saxatilis* | Wave | Wales | Holyhead | F | 53.29981 | -4.67967 |
| **W_brood pouch_04_Ls** | *L. saxatilis* | Wave | Wales | Holyhead | F | 53.29981 | -4.67967 |
| **ACF_1** | *L. saxatilis* | Crab | Wales | Holyhead | F | 53.29981 | -4.67967 |
| **ACF_2** | *L. saxatilis* | Crab | Wales | Holyhead | F | 53.29981 | -4.67967 |
| **ACF_3** | *L. saxatilis* | Crab | Wales | Holyhead | F | 53.29981 | -4.67967 |
| **ACF_4** | *L. saxatilis* | Crab | Wales | Holyhead | F | 53.29981 | -4.67967 |
| **NE_W1_331_Lsax_F** | *L. saxatilis* | Wave | Wales | Holyhead | F | 53.29981 | -4.67967 |
| **NE_W1_356_Lsax_M** | *L. saxatilis* | Wave | Wales | Holyhead | F | 53.29981 | -4.67967 |
| **IM1_6_2** | *L. saxatilis* | Crab | Scotland | Isle of Mull | F | 56.46981 | -5.70344 |
| **CZA005_Ls** | *L. saxatilis* | Crab | Sweden | Koster Area A | ? | 58.82438 | 11.06258 |
| **CZA020_Ls** | *L. saxatilis* | Crab | Sweden | Koster Area A | ? | 58.82438 | 11.06258 |
| **CZA506_Ls** | *L. saxatilis* | Wave | Sweden | Koster Area A | ? | 58.82438 | 11.06258 |
| **CZA524_Ls** | *L. saxatilis* | Wave | Sweden | Koster Area A | ? | 58.82438 | 11.06258 |
| **CZD309_Ls** | *L. saxatilis* | Crab | Sweden | Koster Area D | ? | 58.83091 | 11.13305 |
| **CZD390_Ls** | *L. saxatilis* | Crab | Sweden | Koster Area D | ? | 58.83091 | 11.13305 |
| **CZD563_Ls** | *L. saxatilis* | Wave | Sweden | Koster Area D | ? | 58.83091 | 11.13305 |
| **CZD565_Ls** | *L. saxatilis* | Wave | Sweden | Koster Area D | ? | 58.83091 | 11.13305 |
| **LR_11_Ls** | *L. saxatilis* | Other | Iceland | Laugarnes | M | 64.1525 | -21.88383 |
| **LR_14_Ls** | *L. saxatilis* | Other | Iceland | Laugarnes | F | 64.1525 | -21.88383 |
| **OBI_12_5** | *L. saxatilis* | Crab | Scotland | Oban | F | 56.42207 | -5.48392 |
| **PSM2_4_4** | *L. saxatilis* | Crab | Isle of man | Port Saint Mary | F | 54.07602 | -4.73618 |
| **NE_E1_367_La** | *L. arcana* | Wave | England | Ravenscar | F | 54.41036 | -0.49196 |
| **NE_E2_140_La** | *L. arcana* | Wave | England | Ravenscar | F | 54.41036 | -0.49196 |
| **NE_E2_158_La** | *L. arcana* | Wave | England | Ravenscar | F | 54.41036 | -0.49196 |
| **NE_E2_410_La** | *L. arcana* | Wave | England | Ravenscar | F | 54.41036 | -0.49196 |
| **NE_E2_429_La** | *L. arcana* | Wave | England | Ravenscar | F | 54.41036 | -0.49196 |
| **NE_E4_Larc_F_1** | *L. arcana* | Wave | England | Ravenscar | F | 54.41036 | -0.49196 |
| **NE_E4_Larc_F_2** | *L. arcana* | Wave | England | Ravenscar | F | 54.41036 | -0.49196 |
| **NE_E1_022_Lsb** | *L. saxatilis* | Barnacle | England | Ravenscar | F | 54.41036 | -0.49196 |
| **NE_E2_023_Lsb** | *L. saxatilis* | Barnacle | England | Ravenscar | F | 54.41036 | -0.49196 |
| **NE_E2_151_Ls** | *L. saxatilis* | Wave | England | Ravenscar | F | 54.41036 | -0.49196 |
| **NE_E2_159_Ls** | *L. saxatilis* | Wave | England | Ravenscar | F | 54.41036 | -0.49196 |
| **NE_E2_382_Ls** | *L. saxatilis* | Wave | England | Ravenscar | F | 54.41036 | -0.49196 |
| **NE_E2_439_Ls** | *L. saxatilis* | Wave | England | Ravenscar | F | 54.41036 | -0.49196 |
| **NE_E2_030_Lsax_F** | *L. saxatilis* | Wave | England | Ravenscar | F | 54.41036 | -0.49196 |
| **NE_F1_007_La** | *L. arcana* | Other | France | Roscoff | F | 48.69481 | -4.10734 |
| **NE_F1_137_La** | *L. arcana* | Other | France | Roscoff | F | 48.69481 | -4.10734 |
| **NE_F1_338_La** | *L. arcana* | Other | France | Roscoff | F | 48.69481 | -4.10734 |
| **NE_F1_371_La** | *L. arcana* | Other | France | Roscoff | F | 48.69481 | -4.10734 |
| **NE_F1_141_Lc** | *L. compressa* | N/A | France | Roscoff | F | 48.69481 | -4.10734 |
| **NE_F1_003_Ls** | *L. saxatilis* | Other | France | Roscoff | F | 48.69481 | -4.10734 |
| **NE_F1_087_Lsb** | *L. saxatilis* | Barnacle | France | Roscoff | F | 48.69481 | -4.10734 |
| **NE_F1_134_Ls** | *L. saxatilis* | Other | France | Roscoff | F | 48.69481 | -4.10734 |
| **NE_F1_263_Lsb** | *L. saxatilis* | Barnacle | France | Roscoff | F | 48.69481 | -4.10734 |
| **NE_F1_333_Ls** | *L. saxatilis* | Other | France | Roscoff | F | 48.69481 | -4.10734 |
| **NE_F1_373_Lsb** | *L. saxatilis* | Barnacle | France | Roscoff | F | 48.69481 | -4.10734 |
| **ER_EA1_073_Ls** | *L. saxatilis* | Wave | Spain | Sillerio | ? | 42.07786 | -8.89555 |
| **ER_EA1_116_Ls** | *L. saxatilis* | Wave | Spain | Sillerio | ? | 42.07786 | -8.89555 |
| **ER_EA1_463_Ls** | *L. saxatilis* | Crab | Spain | Sillerio | ? | 42.07786 | -8.89555 |
| **ER_EA1_468_Ls** | *L. saxatilis* | Crab | Spain | Sillerio | ? | 42.07786 | -8.89555 |
| **ER_EB1_100_Ls** | *L. saxatilis* | Wave | Spain | Sillerio | ? | 42.07786 | -8.89555 |
| **ER_EB1_101_Ls** | *L. saxatilis* | Wave | Spain | Sillerio | ? | 42.07786 | -8.89555 |
| **ER_EB1_266_Ls** | *L. saxatilis* | Crab | Spain | Sillerio | ? | 42.07786 | -8.89555 |
| **ER_EA1_547** | *L. saxatilis* | Crab | Spain | Sillerio | ? | 42.07786 | -8.89555 |
| **ER_EA2_029** | *L. saxatilis* | Crab | Spain | Sillerio | ? | 42.07786 | -8.89555 |
| **ER_EA2_569** | *L. saxatilis* | Wave | Spain | Sillerio | ? | 42.07786 | -8.89555 |
| **STA_Larc_F_1** | *L. arcana* | Wave | Scotland | St Abbs | F | 55.89968 | -2.13004 |
| **STA_Larc_F_2** | *L. arcana* | Wave | Scotland | St Abbs | F | 55.89968 | -2.13004 |
| **STA_Lsax_F_1** | *L. saxatilis* | Wave | Scotland | St Abbs | F | 55.89968 | -2.13004 |
| **STA_Lsax_F_2** | *L. saxatilis* | Wave | Scotland | St Abbs | F | 55.89968 | -2.13004 |
| **TCF_1** | *L. saxatilis* | Crab | England | Thornwick | F | 54.13267 | -0.11503 |
| **TCF_2** | *L. saxatilis* | Crab | England | Thornwick | F | 54.13267 | -0.11503 |
| **TCF_3** | *L. saxatilis* | Crab | England | Thornwick | F | 54.13267 | -0.11503 |
| **TCF_4** | *L. saxatilis* | Crab | England | Thornwick | F | 54.13267 | -0.11503 |
| **TWF_1** | *L. saxatilis* | Wave | England | Thornwick | F | 54.13267 | -0.11503 |
| **TWF_2** | *L. saxatilis* | Wave | England | Thornwick | F | 54.13267 | -0.11503 |
| **TWF_3** | *L. saxatilis* | Wave | England | Thornwick | F | 54.13267 | -0.11503 |
| **TWF_4** | *L. saxatilis* | Wave | England | Thornwick | F | 54.13267 | -0.11503 |
| **SWE_Lten_F** | *L. saxatilis* | Brackish | Sweden | Tjarno | F | 58.88994 | 11.13866 |
| **SWE_Lten_M** | *L. saxatilis* | Brackish | Sweden | Tjarno | M | 58.88994 | 11.13866 |
| **TF_130_La** | *L. arcana* | Other | Norway | Trondheim Fjord | F | 63.55228 | 10.46486 |
| **TF_155_La** | *L. saxatilis* | Other | Norway | Trondheim Fjord | F | 63.55228 | 10.46486 |
| **TF9_153_Ls** | *L. saxatilis* | Other | Norway | Trondheim Fjord | F | 63.55228 | 10.46486 |
| **TF9_157_Ls** | *L. saxatilis* | Other | Norway | Trondheim Fjord | F | 63.55228 | 10.46486 |
| **NOR_Larc_F_1** | *L. arcana* | Other | Norway | Varanger Fjord | F | 70.04039 | 29.58401 |
| **NOR_Larc_F_2** | *L. arcana* | Other | Norway | Varanger Fjord | F | 70.04039 | 29.58401 |
| **NOR_Larc_F_3** | *L. arcana* | Other | Norway | Varanger Fjord | F | 70.04039 | 29.58401 |
| **NOR_Larc_F_4** | *L. arcana* | Other | Norway | Varanger Fjord | F | 70.04039 | 29.58401 |
| **NOR_Lcom_F** | *L. compressa* | N/A | Norway | Varanger Fjord | F | 70.04039 | 29.58401 |
| **NOR_Lsax_F_1** | *L. saxatilis* | Other | Norway | Varanger Fjord | F | 70.04039 | 29.58401 |
| **NOR_Lsax_F_2** | *L. saxatilis* | Other | Norway | Varanger Fjord | F | 70.04039 | 29.58401 |
| **NOR_Lsax_F_3** | *L. saxatilis* | Other | Norway | Varanger Fjord | F | 70.04039 | 29.58401 |
| **NOR_Lsax_F_4** | *L. saxatilis* | Other | Norway | Varanger Fjord | F | 70.04039 | 29.58401 |
| **WS_female_Ls** | *L. saxatilis* | Other | Russia | White sea | F | 66.33082 | 33.06251 |
| **WS_male_Ls** | *L. saxatilis* | Other | Russia | White sea | M | 66.33082 | 33.06251 |
| **York_B-12_Ls** | *L. saxatilis* | Other | USA | York | M | 43.15093 | -70.62182 |
| **York_B-1_Ls** | *L. saxatilis* | Other | USA | York | F | 43.15093 | -70.62182 |
| **York_T-3_Ls** | *L. saxatilis* | Other | USA | York | F | 43.15093 | -70.62182 |
| **York_T-9_Ls** | *L. saxatilis* | Other | USA | York | M | 43.15093 | -70.62182 |

**Table S2. Change in the number of sites at each filtering step.** Key commands for filtering in BCFtools are given.

<table>
<tr><th>Dataset</th><th>Total sites</th><th>Total invariant</th><th>Total SNPs</th><th>Multallele SNPs</th></tr>
<tr><td>Full unfiltered dataset</td><td>1,658,076,957</td><td>1,521,844,025</td><td>99,448,272</td><td>17,240,263</td></tr>
<tr><td>After Hard filters<br>-e 'FS>60.0 || SOR>3 || MQ<40 || MQRankSum<-12.5 || QD<2.0 || ReadPosRankSum<-8.0'</td><td>1,527,806,671</td><td>1,429,859,187</td><td>74,224,476</td><td>8,485,685</td></tr>
<tr><td>After Drop SNPs near indels & complex vars<br>-g 5:indel,other</td><td>1,499,745,457</td><td>1,429,859,176</td><td>46,163,273</td><td>2,620,532</td></tr>
<tr><td>After retaining only bialleleic SNPs<br>--exclude-types indels,other --max-alleles 2</td><td>1,473,401,917</td><td>1,429,859,176</td><td>43,542,741</td><td>-</td></tr>
<tr><td>After droping sites above threshold DP<br>-e 'INFO/DP>3750'</td><td>1,450,599,180</td><td>1,408,460,066</td><td>42,139,114</td><td>-</td></tr>
<tr><td>After soft filtering:<br>Set low DP genotypes to missing<br>-S . -e 'FMT/DP<5 | FMT/GQ<20'<br>drop sites with > 0.2 missing data<br>-e 'F_MISSING > 0.2'</td><td>454,361,480</td><td>429,399,898</td><td>24,961,582</td><td>-</td></tr>
<tr><td>After additional hard and soft filtering<br>Drop sites below depth threshold<br>-e 'INFO/DP<1500'<br>Drop genotypes with < 10 reads<br>-e 'FMT/DP<10',<br>Drop sites with > 10% missing data<br>'F_MISSING > 0.1'</td><td>328,831,939</td><td>310,294,741</td><td>18,537,198</td><td>-</td></tr>
</table>

**Table S3. Parameters of the four-population coalescent model implemented in MSprime.** $N_e$, number of haploid sequences, *m* number of haploid sequences per generation moving from one population to another. T, time in generations.

| Parameter | Description |
|---|---|
| $N_e$P1 | The effective size of population 1 |
| $N_e$P2 | The effective size of population 2 |
| $N_e$P3 | The effective size of population 3 |
| $N_e$O | The effective size of population O |
| $N_e$P12 | The effective size of the ancestor of P1,P2 |
| $N_e$P123 | The effective size of the ancestor of P1,P2,P3 |
| $N_eAnc$ | The effective size of the ancestor of P1,P2,P3,O |
| $T_1$ | The time of the split time of P1 and P2 |
| $T_2$ | The split time of P3 and P12 |
| $T_3$ | The split time of O and P123 |
| $m_{2,3}$ | The migration rate from P2 to P3 |
| $m_{3,2}$ | The migration rate from P3 to P2 |

**Table S4. Parameter values for simulating genealogies in MSprime and estimates of $D_{LR}$ calculated from the resulting topology weights.** Each line corresponds to one simulation. Each shaded section corresponds to one of the simulation scenarios described in the text (a - i). SimID: The ID of the simulation, which corresponds with the IDs used in the plotting code and output files on GitHub; Wins: the number of windows simulated for each simulation. *n*, number of individuals simulated for each population; $N_e$, number of haploid sequences simulated in each of the lineages specified (illustrated in Fig. S9); Splits: the time of each split in generations (Illustrated in fig. S9); Migration: the rates of migration from P3 to P2 and P2 to P3 (Illustrated in fig. S12); $D_{LR}$, value of $D_{LR}$ calculated for each simulation, with negative values indicating a deficit of windows on the left side. If 'NA', $D_{LR}$ could not be calculated because there were no left or right sided topologies (*i.e.*, lineage sorting was complete). For scenarios *h* and *i*, the values outside parentheses are for the 'alternate' model (10% of simulated loci), while values inside are for the 'background' model (90% of simulated loci); p-value, the p-value for a G-test of the departure of $D_{LR}$ using 0.5 as the expected proportion of left and right sided windows. The p-value is reported as 'NA' for simulations where $D_{LR}$ could not be calculated. The table continues on the next page.

| | | | | $N_e$ | | | | | | | splits (Gens.) | | | Migration | | | |
|---|---|---|---|---|---|---|---|---|---|---|---|---|---|---|---|---|---|
| | SimID | wins | *n* | P1 | P2 | P3 | O | P12 | P123 | ANC | T1 | T2 | T3 | 3>2 | 2>3 | $D_{LR}$ | *p-value* |
| *a) Varying but equal splits* | N0 | 10000 | 20 | 500 | 500 | 500 | 500 | 500 | 500 | 500 | 1 | 2 | 3 | 0 | 0 | 0.003 | 0.734 |
| | N1 | 10000 | 20 | 500 | 500 | 500 | 500 | 500 | 500 | 500 | 10 | 20 | 30 | 0 | 0 | -0.004 | 0.674 |
| | N2 | 10000 | 20 | 500 | 500 | 500 | 500 | 500 | 500 | 500 | 30 | 60 | 90 | 0 | 0 | -0.026 | 0.008 |
| | N2.5 | 10000 | 20 | 500 | 500 | 500 | 500 | 500 | 500 | 500 | 45 | 90 | 135 | 0 | 0 | -0.004 | 0.689 |
| | N3 | 10000 | 20 | 500 | 500 | 500 | 500 | 500 | 500 | 500 | 90 | 180 | 270 | 0 | 0 | -0.003 | 0.770 |
| | N3.5 | 10000 | 20 | 500 | 500 | 500 | 500 | 500 | 500 | 500 | 135 | 270 | 405 | 0 | 0 | -0.002 | 0.838 |
| | N4 | 10000 | 20 | 500 | 500 | 500 | 500 | 500 | 500 | 500 | 270 | 540 | 810 | 0 | 0 | 0.002 | 0.890 |
| | N4.5 | 10000 | 20 | 500 | 500 | 500 | 500 | 500 | 500 | 500 | 405 | 810 | 1215 | 0 | 0 | -0.008 | 0.560 |
| | N5 | 10000 | 20 | 500 | 500 | 500 | 500 | 500 | 500 | 500 | 810 | 1620 | 2430 | 0 | 0 | -0.017 | 0.467 |
| | N6 | 10000 | 20 | 500 | 500 | 500 | 500 | 500 | 500 | 500 | 2430 | 4860 | 7290 | 0 | 0 | 0.043 | 0.768 |
| | N7 | 10000 | 20 | 500 | 500 | 500 | 500 | 500 | 500 | 500 | 3645 | 7290 | 10935 | 0 | 0 | -0.143 | 0.705 |
| | N8 | 10000 | 20 | 500 | 500 | 500 | 500 | 500 | 500 | 500 | 7290 | 14580 | 21870 | 0 | 0 | NA | NA |
| *b) 5k gens after T1* | Y0 | 10000 | 20 | 500 | 500 | 500 | 500 | 500 | 500 | 500 | 5000 | 5050 | 5100 | 0 | 0 | 0.002 | 0.897 |
| | Y1 | 10000 | 20 | 500 | 500 | 500 | 500 | 500 | 500 | 500 | 5000 | 5100 | 5200 | 0 | 0 | -0.029 | 0.034 |
| | Y2 | 10000 | 20 | 500 | 500 | 500 | 500 | 500 | 500 | 500 | 5000 | 5250 | 5500 | 0 | 0 | -0.005 | 0.775 |
| | Y3 | 10000 | 20 | 500 | 500 | 500 | 500 | 500 | 500 | 500 | 5000 | 5500 | 6000 | 0 | 0 | 0.024 | 0.230 |
| | Y4 | 10000 | 20 | 500 | 500 | 500 | 500 | 500 | 500 | 500 | 5000 | 6000 | 7000 | 0 | 0 | -0.019 | 0.569 |
| | Y5 | 10000 | 20 | 500 | 500 | 500 | 500 | 500 | 500 | 500 | 5000 | 7000 | 9000 | 0 | 0 | 0.058 | 0.524 |
| | Y6 | 10000 | 20 | 500 | 500 | 500 | 500 | 500 | 500 | 500 | 5000 | 9000 | 13000 | 0 | 0 | NA | NA |
| | Y7 | 10000 | 20 | 500 | 500 | 500 | 500 | 500 | 500 | 500 | 5000 | 12000 | 21000 | 0 | 0 | NA | NA |
| *c) Uneven splits* | U0 | 10000 | 20 | 500 | 500 | 500 | 500 | 500 | 500 | 500 | 90 | 260 | 270 | 0 | 0 | -0.002 | 0.878 |
| | U1 | 10000 | 20 | 500 | 500 | 500 | 500 | 500 | 500 | 500 | 90 | 230 | 270 | 0 | 0 | -0.004 | 0.670 |
| | U2 | 10000 | 20 | 500 | 500 | 500 | 500 | 500 | 500 | 500 | 90 | 180 | 270 | 0 | 0 | -0.002 | 0.879 |
| | U3 | 10000 | 20 | 500 | 500 | 500 | 500 | 500 | 500 | 500 | 90 | 130 | 270 | 0 | 0 | -0.010 | 0.326 |
| | U4 | 10000 | 20 | 500 | 500 | 500 | 500 | 500 | 500 | 500 | 90 | 100 | 270 | 0 | 0 | -0.003 | 0.756 |
| | U5 | 10000 | 20 | 500 | 500 | 500 | 500 | 500 | 500 | 500 | 405 | 540 | 1215 | 0 | 0 | -0.002 | 0.869 |
| | U6 | 10000 | 20 | 500 | 500 | 500 | 500 | 500 | 500 | 500 | 405 | 675 | 1215 | 0 | 0 | -0.002 | 0.837 |
| | U7 | 10000 | 20 | 500 | 500 | 500 | 500 | 500 | 500 | 500 | 405 | 810 | 1215 | 0 | 0 | 0.018 | 0.188 |
| | U8 | 10000 | 20 | 500 | 500 | 500 | 500 | 500 | 500 | 500 | 405 | 945 | 1215 | 0 | 0 | 0.034 | 0.027 |
| | U9 | 10000 | 20 | 500 | 500 | 500 | 500 | 500 | 500 | 500 | 405 | 1080 | 1215 | 0 | 0 | -0.005 | 0.794 |
| *d) Variation in Ne* | S0 | 10000 | 20 | 5 | 5 | 5 | 5 | 5 | 5 | 5 | 90 | 180 | 270 | 0 | 0 | NA | NA |
| | S1 | 10000 | 20 | 25 | 25 | 25 | 25 | 25 | 25 | 25 | 90 | 180 | 270 | 0 | 0 | NA | NA |
| | S2 | 10000 | 20 | 50 | 50 | 50 | 50 | 50 | 50 | 50 | 90 | 180 | 270 | 0 | 0 | -0.008 | 0.757 |
| | S3 | 10000 | 20 | 100 | 100 | 100 | 100 | 100 | 100 | 100 | 90 | 180 | 270 | 0 | 0 | -0.001 | 0.932 |
| | S4 | 10000 | 20 | 250 | 250 | 250 | 250 | 250 | 250 | 250 | 90 | 180 | 270 | 0 | 0 | -0.004 | 0.720 |
| | N3 | 10000 | 20 | 500 | 500 | 500 | 500 | 500 | 500 | 500 | 90 | 180 | 270 | 0 | 0 | -0.003 | 0.770 |
| | S5 | 10000 | 20 | 750 | 750 | 750 | 750 | 750 | 750 | 750 | 90 | 180 | 270 | 0 | 0 | 0.014 | 0.176 |
| | S6 | 10000 | 20 | 1250 | 1250 | 1250 | 1250 | 1250 | 1250 | 1250 | 90 | 180 | 270 | 0 | 0 | -0.001 | 0.936 |
| | S7 | 10000 | 20 | 2500 | 2500 | 2500 | 2500 | 2500 | 2500 | 2500 | 90 | 180 | 270 | 0 | 0 | -0.009 | 0.352 |
| | S8 | 10000 | 20 | 5000 | 5000 | 5000 | 5000 | 5000 | 5000 | 5000 | 90 | 180 | 270 | 0 | 0 | 0.001 | 0.912 |

| | SimID | wins | *n* | $N_e$ | | | | | | | splits (Gens.) | | | Migration | | $D_{LR}$ | *p-value* |
|---|---|---|---|---|---|---|---|---|---|---|---|---|---|---|---|---|---|
| | | | | P1 | P2 | P3 | O | P12 | P123 | ANC | T1 | T2 | T3 | 3>2 | 2>3 | | |
| *e) Varying Ne in one population* | N1 | 10000 | 20 | 500 | 500 | 500 | 500 | 500 | 500 | 500 | 10 | 20 | 30 | 0 | 0 | -0.004 | 0.674 |
| | X1 | 10000 | 20 | 500 | 500 | 250 | 500 | 500 | 500 | 500 | 10 | 20 | 30 | 0 | 0 | 0.002 | 0.857 |
| | X2 | 10000 | 20 | 500 | 500 | 50 | 500 | 500 | 500 | 500 | 10 | 20 | 30 | 0 | 0 | -0.017 | 0.087 |
| | X3 | 10000 | 20 | 500 | 500 | 5 | 500 | 500 | 500 | 500 | 10 | 20 | 30 | 0 | 0 | -0.004 | 0.682 |
| | X4 | 10000 | 20 | 500 | 500 | 5000 | 500 | 500 | 500 | 500 | 90 | 180 | 270 | 0 | 0 | -0.010 | 0.320 |
| | N3 | 10000 | 20 | 500 | 500 | 500 | 500 | 500 | 500 | 500 | 90 | 180 | 270 | 0 | 0 | -0.003 | 0.770 |
| | X5 | 10000 | 20 | 500 | 500 | 250 | 500 | 500 | 500 | 500 | 90 | 180 | 270 | 0 | 0 | -0.014 | 0.179 |
| | X6 | 10000 | 20 | 500 | 500 | 5 | 500 | 500 | 500 | 500 | 90 | 180 | 270 | 0 | 0 | 0.004 | 0.661 |
| | X7 | 10000 | 20 | 500 | 500 | 5000 | 500 | 500 | 500 | 500 | 405 | 810 | 1215 | 0 | 0 | -0.012 | 0.334 |
| | N4.5 | 10000 | 20 | 500 | 500 | 500 | 500 | 500 | 500 | 500 | 405 | 810 | 1215 | 0 | 0 | -0.008 | 0.560 |
| | X8 | 10000 | 20 | 500 | 500 | 250 | 500 | 500 | 500 | 500 | 405 | 810 | 1215 | 0 | 0 | 0.005 | 0.719 |
| | X9 | 10000 | 20 | 500 | 500 | 5 | 500 | 500 | 500 | 500 | 405 | 810 | 1215 | 0 | 0 | -0.007 | 0.608 |
| | X10 | 10000 | 20 | 500 | 500 | 5000 | 500 | 500 | 500 | 500 | 7290 | 14580 | 21870 | 0 | 0 | NA | NA |
| | X11 | 10000 | 20 | 500 | 500 | 3000 | 500 | 500 | 500 | 500 | 7290 | 14580 | 21870 | 0 | 0 | NA | NA |
| | X12 | 10000 | 20 | 500 | 500 | 1500 | 500 | 500 | 500 | 500 | 7290 | 14580 | 21870 | 0 | 0 | NA | NA |
| | N8 | 10000 | 20 | 500 | 500 | 500 | 500 | 500 | 500 | 500 | 7290 | 14580 | 21870 | 0 | 0 | NA | NA |
| *f) Uni-directional migration* | N1 | 10000 | 20 | 500 | 500 | 500 | 500 | 500 | 500 | 500 | 10 | 20 | 30 | 0 | 0 | 0.003 | 0.734 |
| | G1 | 10000 | 20 | 500 | 500 | 500 | 500 | 500 | 500 | 500 | 10 | 20 | 30 | 0.001 | 0 | 0.001 | 0.904 |
| | G2 | 10000 | 20 | 500 | 500 | 500 | 500 | 500 | 500 | 500 | 10 | 20 | 30 | 0.01 | 0 | -0.020 | 0.050 |
| | G3 | 10000 | 20 | 500 | 500 | 500 | 500 | 500 | 500 | 500 | 10 | 20 | 30 | 0.1 | 0 | -0.174 | 2.20E-16 |
| | N3 | 10000 | 20 | 500 | 500 | 500 | 500 | 500 | 500 | 500 | 90 | 180 | 270 | 0 | 0 | -0.003 | 0.770 |
| | G4 | 10000 | 20 | 500 | 500 | 500 | 500 | 500 | 500 | 500 | 90 | 180 | 270 | 0.001 | 0 | -0.094 | 2.20E-16 |
| | G5 | 10000 | 20 | 500 | 500 | 500 | 500 | 500 | 500 | 500 | 90 | 180 | 270 | 0.01 | 0 | -0.510 | 2.20E-16 |
| | G6 | 10000 | 20 | 500 | 500 | 500 | 500 | 500 | 500 | 500 | 90 | 180 | 270 | 0.1 | 0 | -0.777 | 2.20E-16 |
| | N4.5 | 10000 | 20 | 500 | 500 | 500 | 500 | 500 | 500 | 500 | 405 | 810 | 1215 | 0 | 0 | -0.008 | 0.560 |
| | G7 | 10000 | 20 | 500 | 500 | 500 | 500 | 500 | 500 | 500 | 405 | 810 | 1215 | 0.001 | 0 | -0.585 | 2.20E-16 |
| | G8 | 10000 | 20 | 500 | 500 | 500 | 500 | 500 | 500 | 500 | 405 | 810 | 1215 | 0.01 | 0 | -0.950 | 2.20E-16 |
| | G9 | 10000 | 20 | 500 | 500 | 500 | 500 | 500 | 500 | 500 | 405 | 810 | 1215 | 0.1 | 0 | -0.981 | 2.20E-16 |
| | N8 | 10000 | 20 | 500 | 500 | 500 | 500 | 500 | 500 | 500 | 7290 | 14580 | 21870 | 0 | 0 | NA | NA |
| | G10 | 10000 | 20 | 500 | 500 | 500 | 500 | 500 | 500 | 500 | 7290 | 14580 | 21870 | 0.001 | 0 | -1.000 | 2.20E-16 |
| | G11 | 10000 | 20 | 500 | 500 | 500 | 500 | 500 | 500 | 500 | 7290 | 14580 | 21870 | 0.01 | 0 | -1.000 | 2.20E-16 |
| | G12 | 10000 | 20 | 500 | 500 | 500 | 500 | 500 | 500 | 500 | 7290 | 14580 | 21870 | 0.1 | 0 | -1.000 | 2.20E-16 |
| *g) Bi-directional migration* | N1 | 10000 | 20 | 500 | 500 | 500 | 500 | 500 | 500 | 500 | 10 | 20 | 30 | 0 | 0 | -0.004 | 0.674 |
| | B1 | 10000 | 20 | 500 | 500 | 500 | 500 | 500 | 500 | 500 | 10 | 20 | 30 | 0.001 | 0.001 | -0.084 | 2.20E-16 |
| | B2 | 10000 | 20 | 500 | 500 | 500 | 500 | 500 | 500 | 500 | 10 | 20 | 30 | 0.01 | 0.01 | -0.098 | 2.20E-16 |
| | B3 | 10000 | 20 | 500 | 500 | 500 | 500 | 500 | 500 | 500 | 10 | 20 | 30 | 0.1 | 0.1 | -0.095 | 2.20E-16 |
| | N3 | 10000 | 20 | 500 | 500 | 500 | 500 | 500 | 500 | 500 | 90 | 180 | 270 | 0 | 0 | -0.003 | 0.7703 |
| | B4 | 10000 | 20 | 500 | 500 | 500 | 500 | 500 | 500 | 500 | 90 | 180 | 270 | 0.001 | 0.001 | -0.349 | 2.20E-16 |
| | B5 | 10000 | 20 | 500 | 500 | 500 | 500 | 500 | 500 | 500 | 90 | 180 | 270 | 0.01 | 0.01 | -0.346 | 2.20E-16 |
| | B6 | 10000 | 20 | 500 | 500 | 500 | 500 | 500 | 500 | 500 | 90 | 180 | 270 | 0.1 | 0.1 | -0.361 | 2.20E-16 |
| | N4.5 | 10000 | 20 | 500 | 500 | 500 | 500 | 500 | 500 | 500 | 405 | 810 | 1215 | 0 | 0 | -0.008 | 0.560 |
| | B7 | 10000 | 20 | 500 | 500 | 500 | 500 | 500 | 500 | 500 | 405 | 810 | 1215 | 0.001 | 0.001 | -0.735 | 2.20E-16 |
| | B8 | 10000 | 20 | 500 | 500 | 500 | 500 | 500 | 500 | 500 | 405 | 810 | 1215 | 0.01 | 0.01 | -0.724 | 2.20E-16 |
| | B9 | 10000 | 20 | 500 | 500 | 500 | 500 | 500 | 500 | 500 | 405 | 810 | 1215 | 0.1 | 0.1 | -0.717 | 2.20E-16 |
| | N8 | 10000 | 20 | 500 | 500 | 500 | 500 | 500 | 500 | 500 | 7290 | 14580 | 21870 | 0 | 0 | NA | NA |
| | B10 | 10000 | 20 | 500 | 500 | 500 | 500 | 500 | 500 | 500 | 7290 | 14580 | 21870 | 0.001 | 0.001 | -1.000 | 2.20E-16 |
| | B11 | 10000 | 20 | 500 | 500 | 500 | 500 | 500 | 500 | 500 | 7290 | 14580 | 21870 | 0.01 | 0.01 | -1.000 | 2.20E-16 |
| | B12 | 10000 | 20 | 500 | 500 | 500 | 500 | 500 | 500 | 500 | 7290 | 14580 | 21870 | 0.1 | 0.1 | -1.000 | 2.20E-16 |
| *h) Migration for 10% of the genome* | I1 | 1000 (9000) | 20 | 500 | 500 | 500 | 500 | 500 | 500 | 500 | 100 | 200 | 300 | 0.001 (0) | 0 | -0.013 | 0.200 |
| | I2 | 1000 (9000) | 20 | 500 | 500 | 500 | 500 | 500 | 500 | 500 | 100 | 200 | 300 | 0.005 (0) | 0 | -0.041 | 4.10E-05 |
| | I3 | 1000 (9000) | 20 | 500 | 500 | 500 | 500 | 500 | 500 | 500 | 100 | 200 | 300 | 0.01 (0) | 0 | -0.057 | 1.17E-08 |
| | I4 | 1000 (9000) | 20 | 500 | 500 | 500 | 500 | 500 | 500 | 500 | 100 | 200 | 300 | 0.05 (0) | 0 | -0.082 | 2.20E-16 |
| | I5 | 1000 (9000) | 20 | 500 | 500 | 500 | 500 | 500 | 500 | 500 | 100 | 200 | 300 | 0.1 (0) | 0 | -0.085 | 2.20E-16 |
| | I6 | 1000 (9000) | 20 | 500 | 500 | 500 | 500 | 500 | 500 | 500 | 100 | 200 | 300 | 0.5 (0) | 0 | -0.087 | 2.20E-16 |
| | I7 | 1000 (9000) | 20 | 500 | 500 | 500 | 500 | 500 | 500 | 500 | 405 | 945 | 1215 | 0.001 (0) | 0 | -0.125 | 2.20E-16 |
| | I8 | 1000 (9000) | 20 | 500 | 500 | 500 | 500 | 500 | 500 | 500 | 405 | 945 | 1215 | 0.005 (0) | 0 | -0.188 | 2.20E-16 |
| | I9 | 1000 (9000) | 20 | 500 | 500 | 500 | 500 | 500 | 500 | 500 | 405 | 945 | 1215 | 0.01 (0) | 0 | -0.197 | 2.20E-16 |
| | I10 | 1000 (9000) | 20 | 500 | 500 | 500 | 500 | 500 | 500 | 500 | 405 | 945 | 1215 | 0.05 (0) | 0 | -0.201 | 2.20E-16 |
| | I11 | 1000 (9000) | 20 | 500 | 500 | 500 | 500 | 500 | 500 | 500 | 405 | 945 | 1215 | 0.1 (0) | 0 | -0.200 | 2.20E-16 |
| | I12 | 1000 (9000) | 20 | 500 | 500 | 500 | 500 | 500 | 500 | 500 | 405 | 945 | 1215 | 0.5 (0) | 0 | -0.201 | 2.20E-16 |
| *i) Ancestral structure* | A1 | 1000 (9000) | 20 | 500 | 500 | 500 | 500 | 500 | 500 | 500 | 100 | 220 (200) | 300 | 0 | 0 | 0.055 | 4.69E-08 |
| | A2 | 1000 (9000) | 20 | 500 | 500 | 500 | 500 | 500 | 500 | 500 | 100 | 240 (200) | 300 | 0 | 0 | 0.061 | 1.56E-09 |
| | A3 | 1000 (9000) | 20 | 500 | 500 | 500 | 500 | 500 | 500 | 500 | 100 | 260 (200) | 300 | 0 | 0 | 0.064 | 2.02E-10 |
| | A4 | 1000 (9000) | 20 | 500 | 500 | 500 | 500 | 500 | 500 | 500 | 100 | 280 (200) | 300 | 0 | 0 | 0.071 | 1.70E-12 |
| | A5 | 1000 (9000) | 20 | 500 | 500 | 500 | 500 | 500 | 500 | 500 | 405 | 1005 (945) | 1215 | 0 | 0 | 0.161 | 2.20E-16 |
| | A6 | 1000 (9000) | 20 | 500 | 500 | 500 | 500 | 500 | 500 | 500 | 405 | 1065 (945) | 1215 | 0 | 0 | 0.165 | 2.20E-16 |
| | A7 | 1000 (9000) | 20 | 500 | 500 | 500 | 500 | 500 | 500 | 500 | 405 | 1125 (945) | 1215 | 0 | 0 | 0.175 | 2.20E-16 |
| | A8 | 1000 (9000) | 20 | 500 | 500 | 500 | 500 | 500 | 500 | 500 | 405 | 1185 (945) | 1215 | 0 | 0 | 0.176 | 2.20E-16 |

**Table S5. Results of symmetry analysis of sub-triangles.** *n*L and *n*R, number of windows on left and right side; *n*LR, sum of the *n*L and *n*R; *n*Expected, 0.5(*n*LR); *n*L - expected, the number of left-sided windows minus the expectation 0.5(*n*LR); $D_{LR}$ expressed as a percentage. *p*, probability of observing the observed $D_{LR}$ by chance (99,999 permutations.). The sub-triangle IDs correspond with the map in Fig. S19.

| Sub-triangle ID | *n*L | *n*R | *n*LR | *n*Expected | *n*L-expected | $D_{LR}$ (%) | *p* |
|---|---|---|---|---|---|---|---|
| 1 | 356 | 367 | 723 | 361.5 | -5.5 | -1.5214385 | 0.7066 |
| 2 | 149 | 150 | 299 | 149.5 | -0.5 | -0.3344482 | 1 |
| 3 | 695 | 585 | 1280 | 640 | 55 | 8.59375 | 0.00258 |
| 4 | 78 | 82 | 160 | 80 | -2 | -2.5 | 0.81362 |
| 5 | 277 | 267 | 544 | 272 | 5 | 1.83823529 | 0.70073 |
| 6 | 1908 | 1549 | 3457 | 1728.5 | 179.5 | 10.3847266 | 1.00E-05 |
| 7 | 62 | 59 | 121 | 60.5 | 1.5 | 2.47933884 | 0.85575 |
| 8 | 197 | 173 | 370 | 185 | 12 | 6.48648649 | 0.06486 |
| 9 | 1199 | 882 | 2081 | 1040.5 | 158.5 | 15.233061 | 1.00E-05 |
| 10 | 2940 | 2401 | 5341 | 2670.5 | 269.5 | 10.0917431 | 1.00E-05 |
| 11 | 39 | 51 | 90 | 45 | -6 | -13.333333 | 0.24686 |
| 12 | 144 | 106 | 250 | 125 | 19 | 15.2 | 0.01939 |
| 13 | 857 | 603 | 1460 | 730 | 127 | 17.3972603 | 1.00E-05 |
| 14 | 2020 | 1432 | 3452 | 1726 | 294 | 17.0336037 | 1.00E-05 |
| 15 | 8031 | 6681 | 14712 | 7356 | 675 | 9.17618271 | 1.00E-05 |
| 16 | 32 | 45 | 77 | 38.5 | -6.5 | -16.883117 | 0.16927 |
| 17 | 112 | 97 | 209 | 104.5 | 7.5 | 7.17703349 | 0.3333 |
| 18 | 645 | 477 | 1122 | 561 | 84 | 14.973262 | 1.00E-05 |
| 19 | 1500 | 1105 | 2605 | 1302.5 | 197.5 | 15.1631478 | 1.00E-05 |
| 20 | 5400 | 4424 | 9824 | 4912 | 488 | 9.93485342 | 1.00E-05 |
| 21 | 12168 | 11010 | 23178 | 11589 | 579 | 4.99611701 | 1.00E-05 |
| 22 | 14 | 45 | 59 | 29.5 | -15.5 | -52.542373 | 0.00014 |
| 23 | 78 | 110 | 188 | 94 | -16 | -17.021277 | 0.02371 |
| 24 | 546 | 424 | 970 | 485 | 61 | 12.5773196 | 0.00012 |
| 25 | 1103 | 853 | 1956 | 978 | 125 | 12.7811861 | 1.00E-05 |
| 26 | 2312 | 2120 | 4432 | 2216 | 96 | 4.33212996 | 0.00403 |
| 27 | 5127 | 4889 | 10016 | 5008 | 119 | 2.37619808 | 0.01788 |
| 28 | 4155 | 4167 | 8322 | 4161 | -6 | -0.1441961 | 0.90472 |
| 29 | 25 | 102 | 127 | 63.5 | -38.5 | -60.629921 | 1.00E-05 |
| 30 | 77 | 134 | 211 | 105.5 | -28.5 | -27.014218 | 9.00E-05 |
| 31 | 213 | 290 | 503 | 251.5 | -38.5 | -15.308151 | 0.00049 |
| 32 | 538 | 592 | 1130 | 565 | -27 | -4.7787611 | 0.11474 |
| 33 | 404 | 466 | 870 | 435 | -31 | -7.1264368 | 0.03906 |
| 34 | 860 | 880 | 1740 | 870 | -10 | -1.1494253 | 0.64965 |
| 35 | 385 | 393 | 778 | 389 | -4 | -1.0282776 | 0.80095 |
| 36 | 940 | 907 | 1847 | 923.5 | 16.5 | 1.7866811 | 0.45593 |
| 37 | 11 | 231 | 242 | 121 | -110 | -90.909091 | 1.00E-05 |
| 38 | 39 | 167 | 206 | 103 | -64 | -62.135922 | 1.00E-05 |
| 39 | 19 | 69 | 88 | 44 | -25 | -56.818182 | 1.00E-05 |
| 40 | 55 | 91 | 146 | 73 | -18 | -24.657534 | 0.00355 |
| 41 | 8 | 22 | 30 | 15 | -7 | -46.666667 | 0.01648 |
| 42 | 49 | 72 | 121 | 60.5 | -11.5 | -19.008264 | 0.04439 |
| 43 | 15 | 21 | 36 | 18 | -3 | -16.666667 | 0.40338 |
| 44 | 40 | 50 | 90 | 45 | -5 | -11.111111 | 0.34027 |
| 45 | 9 | 18 | 27 | 13.5 | -4.5 | -33.333333 | 0.12216 |

**Table S6. Detailed information for the $D_{LR}$ estimates for each chromosomal inversion.** *n*L and *n*R, number of windows on left and right side; *n*LR, sum of the nL and *n*R. $D_{LR}$ expressed as a percentage; p, p-value from a G-test.

| Inversion | *n*L | *n*R | *n*LR | $D_{LR}$ | p |
|---|---|---|---|---|---|
| **LGC1.1** | 638 | 590 | 1228 | 3.90879 | 0.170709 |
| **LGC14.1.5** | 702 | 620 | 1322 | 6.20272 | 0.024071 |
| **LGC14.3** | 264 | 226 | 490 | 7.75510 | 0.085882 |
| **LGC10.2** | 1220 | 914 | 2134 | 14.3392 | 3.23E-11 |
| **LGC7.1** | 191 | 137 | 328 | 16.4634 | 0.002803 |
| **LGC17.1** | 140 | 218 | 358 | -21.7877 | 3.49E-05 |
| **LGC4.1** | 562 | 317 | 879 | 27.8725 | 8.89E-17 |
| **LGC11.1** | 635 | 336 | 971 | 30.7930 | 3.89E-22 |
| **LGC7.2** | 133 | 58 | 191 | 39.2670 | 3.78E-08 |
| **LGC6.1.5** | 366 | 143 | 509 | 43.8113 | 8.78E-24 |
| **LGC1.2** | 115 | 41 | 156 | 47.4359 | 1.48E-09 |
| **LGC9.1** | 1215 | 425 | 1640 | 48.1707 | 2.70E-88 |
| **LGC10.1** | 115 | 38 | 153 | 50.3268 | 1.89E-10 |
| **LGC2.1** | 344 | 37 | 381 | 0.80577 | 5.17E-64 |

**Table S7. Physical locations for the 100 SNP windows with Tr = 1.** Window ID, Contig ID, start and end positions (bp), midpoint (bp) and length of the window are provided.

| Window ID | Contig | Start | End | Midpoint | Length (bp) |
|---|---|---|---|---|---|
| 6509 | Contig1261 | 126474 | 128742 | 127504 | 2268 |
| 9765 | Contig1808 | 11041 | 15738 | 13749 | 4697 |
| 11034 | Contig2032 | 42723 | 53508 | 47136 | 10785 |
| 11035 | Contig2032 | 53513 | 56923 | 54959 | 3410 |
| 11478 | Contig2118 | 102328 | 107302 | 104273 | 4974 |
| 12105 | Contig2245 | 3318 | 8772 | 6955 | 5454 |
| 12106 | Contig2245 | 8774 | 12651 | 10623 | 3877 |
| 12107 | Contig2245 | 13335 | 15794 | 14675 | 2459 |
| 12108 | Contig2245 | 15795 | 20413 | 17867 | 4618 |
| 17128 | Contig3201 | 56577 | 61149 | 59075 | 4572 |
| 19501 | Contig3802 | 3800 | 13562 | 7480 | 9762 |
| 20194 | Contig3968 | 8478 | 15092 | 11235 | 6614 |
| 20195 | Contig3968 | 15127 | 20380 | 19290 | 5253 |
| 20198 | Contig3970 | 114 | 2486 | 1323 | 2372 |
| 29732 | Contig7286 | 3835 | 6853 | 5315 | 3018 |
| 33677 | Contig9868 | 626 | 4887 | 2727 | 4261 |
| 35486 | Contig12341 | 104 | 8623 | 4996 | 8519 |
| 35532 | Contig12403 | 20722 | 24153 | 22488 | 3431 |
| 41475 | Contig38747 | 33289 | 36757 | 34788 | 3468 |
| 43427 | Contig39155 | 6308 | 9114 | 7508 | 2806 |
| 48810 | Contig40336 | 46155 | 65216 | 50176 | 19061 |
| 52694 | Contig41237 | 137578 | 144181 | 139999 | 6603 |
| 52696 | Contig41237 | 148590 | 153114 | 150635 | 4524 |
| 62064 | Contig43562 | 38079 | 42770 | 39672 | 4691 |
| 73401 | Contig47534 | 10949 | 25863 | 20767 | 14914 |
| 75068 | Contig47979 | 1979 | 9357 | 3107 | 7378 |
| 75069 | Contig47979 | 9361 | 13538 | 11593 | 4177 |
| 75070 | Contig47979 | 13549 | 16551 | 14532 | 3002 |
| 76238 | Contig48348 | 35646 | 37461 | 36466 | 1815 |
| 76239 | Contig48348 | 37489 | 39610 | 38534 | 2121 |
| 85306 | Contig51247 | 433 | 4747 | 2719 | 4314 |
| 87582 | Contig52055 | 6787 | 17879 | 15996 | 11092 |
| 89070 | Contig52509 | 28533 | 30226 | 29300 | 1693 |
| 95372 | Contig54667 | 6589 | 16281 | 11706 | 9692 |
| 95373 | Contig54667 | 16283 | 27260 | 24632 | 10977 |
| 95374 | Contig54667 | 27273 | 51824 | 33881 | 24551 |
| 106684 | Contig59771 | 2623 | 9459 | 7372 | 6836 |
| 106685 | Contig59771 | 9530 | 16568 | 12499 | 7038 |
| 106686 | Contig59771 | 16569 | 19644 | 18183 | 3075 |
| 106688 | Contig59771 | 23960 | 26382 | 24953 | 2422 |
| 106689 | Contig59771 | 26394 | 29344 | 27719 | 2950 |
| 106690 | Contig59771 | 29347 | 42145 | 35808 | 12798 |
| 107721 | Contig60406 | 1641 | 9707 | 6823 | 8066 |
| 107723 | Contig60406 | 12770 | 14708 | 13659 | 1938 |
| 107726 | Contig60406 | 28962 | 34175 | 30860 | 5213 |
| 109076 | Contig61156 | 74402 | 80097 | 77266 | 5695 |
| 109077 | Contig61156 | 80144 | 90066 | 86359 | 9922 |
| 110564 | Contig61890 | 9294 | 14307 | 12512 | 5013 |
| 110565 | Contig61890 | 14319 | 16927 | 15871 | 2608 |
| 113482 | Contig63723 | 406 | 6958 | 3058 | 6552 |
| 117438 | Contig66365 | 57216 | 63741 | 61151 | 6525 |
| 119150 | Contig67554 | 442 | 6257 | 2334 | 5815 |
| 123173 | Contig70615 | 50027 | 54603 | 52359 | 4576 |
| 129459 | Contig76495 | 139 | 8081 | 3103 | 7942 |
| 129460 | Contig76495 | 8089 | 11196 | 9341 | 3107 |
| 129462 | Contig76495 | 14083 | 17244 | 15818 | 3161 |
| 129755 | Contig76820 | 9113 | 11164 | 10189 | 2051 |
| 129756 | Contig76820 | 11165 | 15705 | 13333 | 4540 |
| 131863 | Contig79517 | 160 | 15210 | 5063 | 15050 |
| 132998 | Contig81225 | 2204 | 26071 | 14975 | 23867 |
| 133001 | Contig81225 | 31125 | 40681 | 34667 | 9556 |
| 133002 | Contig81225 | 40683 | 51618 | 45968 | 10935 |
| 133003 | Contig81225 | 51640 | 63572 | 60534 | 11932 |
| 133004 | Contig81225 | 63575 | 68667 | 65626 | 5092 |
| 133005 | Contig81225 | 68686 | 70837 | 69690 | 2151 |
| 133006 | Contig81225 | 70858 | 74927 | 72820 | 4069 |
| 133007 | Contig81225 | 74935 | 79268 | 77021 | 4333 |
| 133259 | Contig81547 | 7493 | 15204 | 10367 | 7711 |
| 134775 | Contig83530 | 5330 | 22231 | 20271 | 16901 |
| 134776 | Contig83530 | 22241 | 38928 | 28437 | 16687 |
| 134777 | Contig83530 | 38933 | 44610 | 43565 | 5677 |
| 134778 | Contig83530 | 44623 | 49443 | 46388 | 4820 |
| 134779 | Contig83530 | 49444 | 53568 | 51581 | 4124 |
| 134780 | Contig83530 | 53573 | 59680 | 54913 | 6107 |
| 134781 | Contig83530 | 59867 | 69596 | 64675 | 9729 |
| 136627 | Contig86117 | 2927 | 9608 | 6826 | 6681 |
| 138308 | Contig89329 | 6746 | 9600 | 7531 | 2854 |
| 145224 | Contig108528 | 615 | 29659 | 11105 | 29044 |
| 145551 | Contig110028 | 504 | 3054 | 1950 | 2550 |
| 146704 | Contig115331 | 7993 | 9268 | 8542 | 1275 |
| 147068 | Contig116989 | 10306 | 12707 | 11517 | 2401 |
| 149106 | Contig128971 | 4271 | 18449 | 13562 | 14178 |
| 152639 | Contig173651 | 983 | 12024 | 3601 | 11041 |
| 152984 | Contig181579 | 2208 | 8526 | 5578 | 6318 |
| 153003 | Contig181768 | 96 | 7912 | 3225 | 7816 |
| 153004 | Contig181768 | 7916 | 18521 | 15056 | 10605 |
| 153005 | Contig181768 | 18524 | 20699 | 19574 | 2175 |
| 154083 | Contig217335 | 1777 | 4453 | 2839 | 2676 |

**Table S8. The 50 contigs with 1 or more windows where Tr = 1**. Contig, the ID of the assembly contig; *n* wins where Tr =1, the number of windows where Tr = 1 on that contig. Start, start of the region where Tr = 1, End, end of the region where Tr = 1. Cumulative length, the sum of lengths of windows of the same contig (bp).

| Contig ID | *n* wins where Tr = 1 | Start | End | Cumulative length |
|---|---|---|---|---|
| Contig115331 | 1 | 7993 | 9268 | 1275 |
| Contig52509 | 1 | 28533 | 30226 | 1693 |
| Contig1261 | 1 | 126474 | 128742 | 2268 |
| Contig3970 | 1 | 114 | 2486 | 2372 |
| Contig116989 | 1 | 10306 | 12707 | 2401 |
| Contig110028 | 1 | 504 | 3054 | 2550 |
| Contig217335 | 1 | 1777 | 4453 | 2676 |
| Contig39155 | 1 | 6308 | 9114 | 2806 |
| Contig89329 | 1 | 6746 | 9600 | 2854 |
| Contig7286 | 1 | 3835 | 6853 | 3018 |
| Contig12403 | 1 | 20722 | 24153 | 3431 |
| Contig38747 | 1 | 33289 | 36757 | 3468 |
| Contig9868 | 1 | 626 | 4887 | 4261 |
| Contig51247 | 1 | 433 | 4747 | 4314 |
| Contig3201 | 1 | 56577 | 61149 | 4572 |
| Contig70615 | 1 | 50027 | 54603 | 4576 |
| Contig43562 | 1 | 38079 | 42770 | 4691 |
| Contig1808 | 1 | 11041 | 15738 | 4697 |
| Contig2118 | 1 | 102328 | 107302 | 4974 |
| Contig67554 | 1 | 442 | 6257 | 5815 |
| Contig181579 | 1 | 2208 | 8526 | 6318 |
| Contig66365 | 1 | 57216 | 63741 | 6525 |
| Contig63723 | 1 | 406 | 6958 | 6552 |
| Contig86117 | 1 | 2927 | 9608 | 6681 |
| Contig81547 | 1 | 7493 | 15204 | 7711 |
| Contig12341 | 1 | 104 | 8623 | 8519 |
| Contig3802 | 1 | 3800 | 13562 | 9762 |
| Contig173651 | 1 | 983 | 12024 | 11041 |
| Contig52055 | 1 | 6787 | 17879 | 11092 |
| Contig128971 | 1 | 4271 | 18449 | 14178 |
| Contig47534 | 1 | 10949 | 25863 | 14914 |
| Contig79517 | 1 | 160 | 15210 | 15050 |
| Contig40336 | 1 | 46155 | 65216 | 19061 |
| Contig108528 | 1 | 615 | 29659 | 29044 |
| Contig48348 | 2 | 35646 | 39610 | 3964 |
| Contig76820 | 2 | 9113 | 15705 | 6592 |
| Contig61890 | 2 | 9294 | 16927 | 7633 |
| Contig41237 | 2 | 137578 | 153114 | 15536 |
| Contig3968 | 2 | 8478 | 20380 | 11902 |
| Contig2032 | 2 | 42723 | 56923 | 14200 |
| Contig61156 | 2 | 74402 | 90066 | 15664 |
| Contig76495 | 3 | 139 | 17244 | 17105 |
| Contig47979 | 3 | 1979 | 16551 | 14572 |
| Contig60406 | 3 | 1641 | 34175 | 32534 |
| Contig181768 | 3 | 96 | 20699 | 20603 |
| Contig54667 | 3 | 6589 | 51824 | 45235 |
| Contig2245 | 4 | 3318 | 61149 | 57831 |
| Contig59771 | 6 | 2623 | 42145 | 39522 |
| Contig83530 | 7 | 5330 | 69596 | 64266 |
| Contig81225 | 8 | 2204 | 79268 | 77064 |

**Table S9. Estimates of Tajima's D obtained for each group along with bias and variance estimates obtained using jackknife resampling.** *n*, sample size for each group. Estimate, the point estimate of Tajima's D obtained for the whole dataset. Jackknife est., the average Tajima's D over all of the *n* - 1 estimates. Bias, the adjusted difference between the point and jackknife estimates of Tajima's D. S.E., the estimated standard error for the estimates of Tajima's D. The 95% confidence intervals around the estimates of Tajima's D.

| Group | *n* | Estimate $\hat{\theta}$ | Jackknife est. $\bar{\theta}$ | Bias $(n-1)(\hat{\theta}-\bar{\theta})$ | S.E. $\left[\frac{n-1}{n}\sum_{i=1}^{n}(\hat{\theta}_i-\bar{\theta})^2\right]^{1/2}$ | 95% CI $\bar{\theta} \pm 1.96SE$ |
|---|---|---|---|---|---|---|
| Egg-layers | 28 | -0.241 | 0.237 | 0.105 | 0.102 | -0.037 — -0.437 |
| Live-bearers | 80 | -1.893 | -1.889 | 0.303 | 0.061 | -1.770 — -2.007 |
| North | 68 | -1.747 | -1.742 | 0.349 | 0.066 | -1.612 — -1.872 |
| Spain | 12 | -0.590 | -0.542 | 0.526 | 0.134 | -0.279 — -0.804 |

**Table S10. Mean estimate of the number, proportion, and difference in proportion of private alleles between groups.** The means are calculated from 10000 iterations, where the larger group was randomly down sampled without replacement to match the size of the smaller group. *n*, number of samples of each group compared, *n* private, the number of loci with an allele private to one group. *P* private, the proportion of private alleles ($P_{priv} = n_{private} / n_{polymorphic}$). Difference $P_{priv}$, the difference in the proportion of private alleles between groups.

| Group | *n* | *n* private | | *P* private | | Difference $P_{priv}$ |
|---|---|---|---|---|---|---|
| | | Egg-layers | Live-bearers | Egg-layers | Live-bearers | Egg-layers - Live-bearers |
| | | Spain | North | Spain | North | Spain - North |
| Egg-layer vs Live-bearer | 28 | 4965 | 2063 | 0.907 | 0.801 | 0.105 |
| Spain vs North | 12 | 1066 | 834 | 0.864 | 0.832 | 0.318 |

**Table S11. Details of the number of high Tr windows on each linkage group.** LG, linkage group; n wins, the total number of windows associated with each LG; Tr 0.7-0.8, the number of windows on each LG with a Tr score between 0.7 & 0.8; Tr 0.8-0.9, the number of windows on each LG with a Tr score between 0.8 & 0.9; Tr 0.9-1.0, the number of windows on each LG with a Tr score between 0.9 & 1.0; Tr 1, the number of windows on each LG with a Tr score of 1. Prop hTr, the proportion of windows on each LG that high Tr (*i.e.* Tr > 0.7); *n* Prop hTr$_r$, the number of windows on each LG that are hTr ; Exp, the expected number of high Tr windows on each LG. P, the probability of observing *n* hTr by chance, as determined by a permutation test (9,999 permutations), adjusted for multiple testing (Benjamini-Hochberg correction at α = 0.05).

| LG | *n* wins | Tr 0.7-0.8 | Tr 0.8-0.9 | Tr 0.9-1 | Tr 1 | Prop. hTr | *n* hTr | Exp | *P* |
|---|---|---|---|---|---|---|---|---|---|
| **1** | 10587 | 40 | 20 | 19 | 7 | 0.130 | 86 | 54.67 | 0.00085 |
| **2** | 10799 | 31 | 11 | 9 | 3 | 0.132 | 54 | 55.77 | 0.47385 |
| **3** | 6390 | 14 | 1 | 5 | 0 | 0.078 | 20 | 33.00 | 0.0204 |
| **4** | 5970 | 21 | 4 | 0 | 1 | 0.073 | 26 | 30.83 | 0.27292 |
| **5** | 6449 | 6 | 6 | 6 | 1 | 0.079 | 19 | 33.30 | 0.01496 |
| **6** | 5228 | 11 | 4 | 8 | 2 | 0.064 | 26 | 26.99 | 0.4591 |
| **7** | 4551 | 11 | 13 | 10 | 1 | 0.056 | 35 | 23.50 | 0.03251 |
| **8** | 2597 | 3 | 1 | 3 | 0 | 0.032 | 7 | 13.41 | 0.06953 |
| **9** | 3646 | 4 | 2 | 3 | 3 | 0.044 | 12 | 18.82 | 0.0867 |
| **10** | 3289 | 4 | 1 | 0 | 0 | 0.040 | 5 | 16.98 | 0.00383 |
| **11** | 2365 | 3 | 3 | 3 | 0 | 0.029 | 9 | 12.21 | 0.26581 |
| **12** | 5279 | 12 | 3 | 3 | 0 | 0.065 | 18 | 27.26 | 0.06724 |
| **13** | 3476 | 8 | 5 | 1 | 0 | 0.043 | 14 | 17.95 | 0.28872 |
| **14** | 2597 | 14 | 4 | 6 | 3 | 0.032 | 27 | 13.41 | 0.00453 |
| **15** | 2901 | 16 | 10 | 0 | 0 | 0.035 | 26 | 14.98 | 0.01615 |
| **16** | 1046 | 12 | 5 | 0 | 0 | 0.013 | 17 | 5.402 | 0.0017 |
| **17** | 4542 | 15 | 7 | 0 | 0 | 0.056 | 22 | 23.45 | 0.46028 |

**Table S12. Counts of genes from the differential expression analysis and their association with Tr weight.**

| Tr | counts | | | | | Proportions | | | | Total Repro |
|---|---|---|---|---|---|---|---|---|---|---|
| | None | Foot | Repro | Both | Total | None | Foot | Repro | Both | |
| < 0.1 | 469 | 3 | 33 | 4 | 509 | 0.921 | 0.006 | 0.065 | 0.008 | 0.073 |
| 0.1-0.2 | 2530 | 19 | 165 | 15 | 2729 | 0.927 | 0.007 | 0.060 | 0.005 | 0.066 |
| 0.2-0.3 | 7405 | 58 | 536 | 27 | 8026 | 0.923 | 0.007 | 0.067 | 0.003 | 0.070 |
| 0.3-0.4 | 7539 | 37 | 526 | 21 | 8123 | 0.928 | 0.005 | 0.065 | 0.003 | 0.067 |
| 0.4-0.5 | 2337 | 13 | 190 | 12 | 2552 | 0.916 | 0.005 | 0.074 | 0.005 | 0.079 |
| 0.5-0.6 | 722 | 6 | 56 | 4 | 788 | 0.916 | 0.008 | 0.071 | 0.005 | 0.076 |
| 0.6-0.7 | 246 | 1 | 24 | 6 | 277 | 0.888 | 0.004 | 0.087 | 0.022 | 0.108 |
| 0.7-0.8 | 68 | 1 | 11 | 2 | 82 | 0.829 | 0.012 | 0.134 | 0.024 | 0.159 |
| 0.8-0.9 | 27 | 0 | 9 | 3 | 39 | 0.692 | 0.000 | 0.231 | 0.077 | 0.308 |
| > 0.9 | 16 | 0 | 13 | 5 | 34 | 0.471 | 0.000 | 0.382 | 0.147 | 0.529 |

**Table S13. Best blast protein hits for the 27 genes.** These were both differentially expressed between reproductive tissues and in windows associated with reproductive mode (Tr > 0.7). $Tr_{gene}$ if the highest Tr value of a window overlapping the gene. $Tr_{max}$ is the highest Tr value for a window on the same assembly contig.

| ***L. saxatilis* gene ID** | **Tissue** | **Expression higher in** | **Blast % Identity** | **Blast % e-value** | **$Tr_{gene}$** | **$Tr_{max}$** | **Description** |
|---|---|---|---|---|---|---|---|
| Lsa_00004286 | reproductive | brood pouch | 52.96 | 4.07E-131 | 0.733 | 0.733 | non-neuronal intermediate filament protein B [*Cornu aspersum*] |
| Lsa_00006231 | reproductive | brood pouch | 57.86 | 2.82E-48 | 0.741 | 0.741 | cartilage matrix protein-like [*Pomacea canaliculata*] |
| Lsa_00012399 | reproductive | Jelly gland | 68.03 | 5.36E-96 | 0.789 | 0.789 | adenosine 3'-phospho 5'-phosphosulfate transporter 1-like isoform X2 [*Pomacea canaliculata*] |
| Lsa_00014176 | reproductive | brood pouch | 59.56 | 2.82E-48 | 0.726 | 0.726 | putative polypeptide N-acetylgalactosaminyltransferase 10 [Pomacea canaliculata] |
| Lsa_00014813 | reproductive | Jelly gland | 44.54 | 1.63E-30 | 0.962 | 0.962 | hemicentin-1-like isoform X2 [*Haliotis rubra*] |
| Lsa_00025155 | reproductive | brood pouch | 65.80 | 3.74E-56 | 0.803 | 0.803 | proton myo-inositol cotransporter-like [*Pomacea canaliculata*] |
| Lsa_00028176 | reproductive | brood pouch | 50.19 | 4.58E-148 | 0.968 | 1.000 | cytochrome P450 26A1-like [*Haliotis rufescens*] |
| Lsa_00033754 | reproductive | Jelly gland | 69.47 | 2.74E-31 | 0.801 | 0.801 | capsule gland specific secretory protein [*Reishia bronni*] |
| Lsa_00035649 | reproductive | Jelly gland | 38.97 | 1.87E-21 | 0.956 | 0.956 | lectin L6-like protein [*Crepidula fornicata*] |
| Lsa_00037480 | reproductive | Jelly gland | 36.27 | 8.67E-117 | 0.786 | 0.786 | uncharacterized protein LOC112573002 [*Pomacea canaliculata*] |
| Lsa_00039211 | reproductive | Jelly gland | 50.00 | 9.23E-46 | 0.731 | 0.731 | capsule gland specific secretory protein [*Reishia bronni*] |
| Lsa_00040601 | reproductive | Jelly gland | 41.81 | 4.69E-30 | 1.00 | 1.000 | semaphorin-5A-like [*Haliotis rufescens*] |
| Lsa_00043868 | reproductive | Jelly gland | 52.40 | 4.53E-53 | 0.9063 | 1.000 | hypothetical protein BaRGS_026586, partial [*Batillaria attramentaria*] |
| Lsa_00000885 | reproductive | No blast hit | | | 0.8279 | 0.828 | |
| Lsa_00005839 | reproductive | No blast hit | | | 0.887 | 0.914 | |
| Lsa_00015408 | reproductive | No blast hit | | | 0.838 | 0.838 | |
| Lsa_00021287 | reproductive | No blast hit | | | 0.755 | 0.755 | |
| Lsa_00021345 | reproductive | No blast hit | | | 0.800 | 0.800 | |
| Lsa_00032450 | reproductive | No blast hit | | | 1.000 | 1.000 | |
| Lsa_00035674 | reproductive | No blast hit | | | 0.802 | 0.802 | |
| Lsa_00048035 | reproductive | No blast hit | | | 0.913 | 0.913 | |
| Lsa_00039796 | reproductive | No blast hit | | | 0.971 | 0.971 | |
| Lsa_00016725 | reproductive & foot | No blast hit | | | 0.781 | 0.927 | |
| Lsa_00018835 | reproductive & foot | No blast hit | | | 0.898 | 0.898 | |
| Lsa_00026229 | reproductive & foot | No blast hit | | | 0.725 | 0.725 | |
| Lsa_00031598 | reproductive & foot | No blast hit | | | 1.000 | 1.000 | |
| Lsa_00042681 | reproductive & foot | No blast hit | | | 0.988 | 0.988 | |

**Table S14. Ratio of windows that show perfect concordance with the species tree and the alternative tree Tr in simulations with differing levels of migration.** We conducted 12 simulations where migration only occurs for a fraction of the genome. We defined four histories with an $N_e$ of 500 for all populations and equal split times, but with the duration between splits varying between the two scenarios (T1 = 100, T2 = 200 T3 = 300; T1 = 405, T2 = 945, T3 = 1215). For each scenario, we modelled heterogenous migration by combining two simulations together. 90% of genealogies were simulated under a 'background' demography without gene flow, and the remaining 10% were simulated under the same model, but one of seven rates of migration from P3 to P2: *m* = 0, 0.001, 0.005, 0.010, 0.05, 0.01, or 0.05. For each simulation we determined the number of windows where T = 1 (*n* Tr = 1) and divided this by the number of windows that perfectly fit the background topology (Tb). In our simulation, we always observed a ratio of Tr:Tb much less than 1, with maximum of 0.37 at an extreme migration rate of 50% between P3 and P2. In our observed data, we observed a positive ratio of 1.41, indicating that windows with Tr = 1 are more abundant than Tb = 1.

| Split times | $m_{p3 > p2}$ | *n* Tr = 1 | Tr:Tb |
|---|---|---|---|
| T100, 200, 300 | 0.000 | 0 | 0.00000000 |
| T100, 200, 300 | 0.001 | 0 | 0.00000000 |
| T100, 200, 300 | 0.005 | 0 | 0.00000000 |
| T100, 200, 300 | 0.010 | 1 | 0.01538462 |
| T100, 200, 300 | 0.050 | 9 | 0.13846154 |
| T100, 200, 300 | 0.100 | 22 | 0.33846154 |
| T100, 200, 300 | 0.500 | 24 | 0.36923077 |
| T405, 945, 1215 | 0.000 | 0 | 0.00000000 |
| T405, 945, 1215 | 0.001 | 54 | 0.01084337 |
| T405, 945, 1215 | 0.005 | 359 | 0.07208835 |
| T405, 945, 1215 | 0.010 | 580 | 0.11646586 |
| T405, 945, 1215 | 0.050 | 715 | 0.14357430 |
| T405, 945, 1215 | 0.100 | 695 | 0.13955823 |
| T405, 945, 1215 | 0.500 | 713 | 0.14317269 |
